# SingleCellSignalR: Inference of intercellular networks from single-cell transcriptomics

**DOI:** 10.1101/2019.12.11.872895

**Authors:** Simon Cabello-Aguilar, Fabien Kon Sun Tack, Mélissa Alame, Caroline Fau, Matthieu Lacroix, Jacques Colinge

## Abstract

Single-cell transcriptomics offers unprecedented opportunities to infer the ligand-receptor interactions underlying cellular networks. We introduce a new, curated ligand-receptor database and a novel regularized score to perform such inferences. For the first time, we try to assess the confidence in predicted ligand-receptor interactions and show that our regularized score outperforms other scoring schemes while controlling false positives. SingleCellSignalR is implemented as an open-access R package accessible to entry-level users and available from https://github.com/SCA-IRCM. Analysis results come in a variety of tabular and graphical formats. For instance, we provide a unique network view integrating all the intercellular interactions, and a function relating receptors to expressed intracellular pathways. A detailed comparison with related tools is conducted. Among various examples, we demonstrate SingleCellSignalR on mouse epidermis data and discover an oriented communication structure from external to basal layers.

## INTRODUCTION

In multicellular organisms, cells engage in a large number of interactions with adjacent or remote partners. They do so to coordinate their fate and behavior from early development stages to mature tissues (1–4), in healthy and diseased (5) conditions. Although other mechanisms may play a role such as tunneling nanotubes, secreted vesicles, or ion fluxes, a significant part of cellular communications is carried over by secreted ligand and cell surface receptor physical interactions (6). In the particular case of tumors, cancer cells can reprogram their microenvironment through secreted factors, turning neutral or anti-tumor cells into tumor supportive elements (7, 8). The emergence of single-cell RNA sequencing (scRNA-seq) technologies (9–11) has provided researchers with powerful means of learning which cells compose specific tissues. The different cell populations present in a sample can be determined by applying unsupervised clustering (12). Further tools exist to infer intracellular pathway activity (13–15), i.e., internal cell states. While such compositional descriptions are essential, deciphering individual cell contributions in tissues requires the unraveling of cellular interactions. Recent scRNA-seq-based studies illustrated how ligand-receptor (LR) interaction mapping might provide a better insight in tissue development and homeostasis or tumor biology. For instance, Puram *et al*. (16) studied head and neck squamous cell carcinomas. They were able to identify an LR interaction, TGFB3-TGFBR2, involved in the communication between cancer cells undergoing (partial) epithelial to mesenchymal transition, at the leading edge of the tumor and cancer-associated fibroblasts. The latter interaction was showed to be necessary for invasiveness. Additional examples illustrating the discovery potential of LR maps can be found in the literature (17–21). Those results highlight the need for a systems biology tool that would assist investigators in portraying cellular networks efficiently and rigorously.

SingleCellSignalR is the first such tool available in R. It relies on a comprehensive database of known LR interactions, which we called LR*db*. It also introduces a new regularized product score aimed at adapting to variable levels of depth in single-cell data sets, i.e., the prevalence of censored read counts or dropouts. LR*db* is the result of integrating and curating existing sources plus manual additions; to the best of our knowledge, it is the largest database of this kind. The new scoring approach has the advantage of facilitating the use of thresholds on LR interaction scores to control false positives and not only rank LR interactions. SingleSignalR can start from raw read count matrices and use integrated data normalization, clustering, and cell type calling solutions before inferring LR interactions between cell populations, i.e., cell clusters. Alternatively, those preliminary steps can be substituted by any other tools or framework, and SingleSignalR used for LR interaction inference only, its primary purpose. In order to facilitate the interpretation of LR interactions and put them in context, a range of visualization and complementary analysis tools are provided, e.g., the inference of intracellular networks rooted at the receptors – or ligands – expressed by a particular cell type. LR*db* contains human genes, but we can accommodate murine data sets by translating murine genes to their human orthologs. An application example illustrates this functionality on mouse skin data.

Several authors proposed tools for mapping LR interactions (18, 22–26) from scRNA-seq data. They exploited different reference lists of potential LR interactions and LR interaction scoring schemes to rank those interactions. A systematic comparison of scoring performance and tool features is presented.

## MATERIALS AND METHODS

### LR*db* – A curated database of ligand-receptor interactions

We decided to emphasize interpretable inferences that are supported by the literature or experimental data. Accordingly, we compiled the content of existing databases that contain such LR pairs: FANTOM5 (6), HPRD (27), HPMR (28), the IUPHAR/BPS Guide to Pharmacology (29), and UniProtKB/Swissprot (30) annotations related to families of ligands or receptors covered in the previous databases. We also extracted LR pairs, with respective participants annotated as ligand or receptor in GO (31), from Reactome (32) pathways. We required GO Cellular Compartment (GOCC) annotation “receptor complex” (GO:0043235) for receptors, and “extracellular space” (GO:0005615) or “extracellular region” (GO:0005576) for ligands. Reactome pathways were downloaded from Pathway Commons (33). That yielded 3,191 pairs. Inspection of the latter revealed 106 dubious cases, i.e., ligand-ligand or receptor-receptor, or non-secreted gene products. After elimination, we obtained 3,085 reliable LR pairs, which we extended with 166 pairs extracted from cellsignaling.com maps and related literature manually to reach 3,251 LR pairs (Fig. 1A-B). All the gene symbols were converted into the latest HUGO version as part of the curation process.

**Figure 1.**
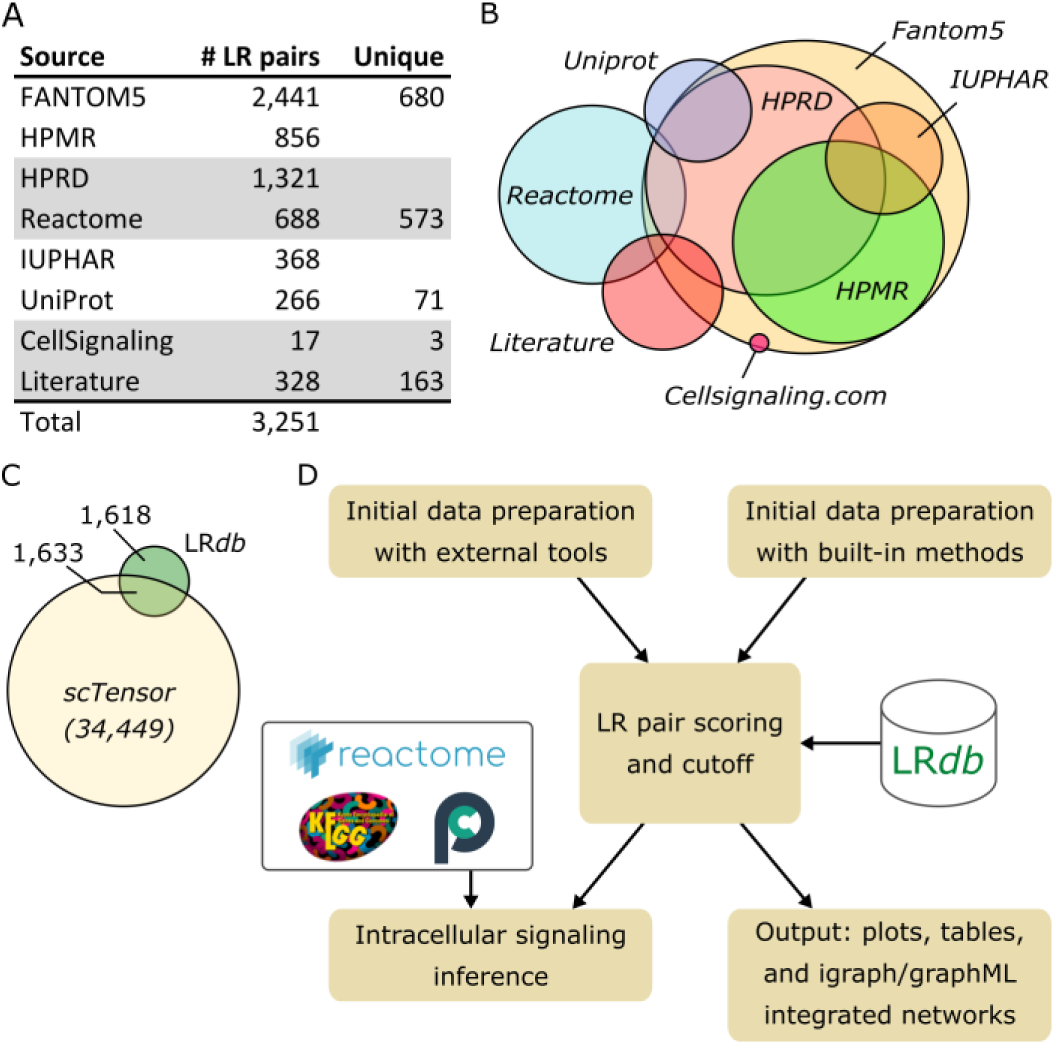
SingleCellSignalR databases and workflow. **A.** Ligand-receptor database (LR*db*) sources. **B.** Approximate overlap of sources. **C.** Overlap (50.2%) of LR*db* with an example of LR database derived from STRING (here the database of scTensor). **D.** SingleCellSignalR general workflow with input transcript expression matrix both normalized and clustered by either independent tools or by SingleCellSignalR basic built-in algorithms.

A proper comparison of SingleCellSignalR with existing tools is presented in Results and Discussion. At this stage, it is nonetheless interesting to note that some tools (23, 24) relied on LR pairs deduced from STRING with principles analogous to what we did with Reactome. That yielded a large number of reference LR pairs, >30,000 typically, but the overlap with LR*db* extensive collection of known LR interactions remained modest (50.2%) (Fig. 1C). Although that approach might provide additional discovery potential, we decided to stay with literature-derived LR interactions.

### SingleCellSignalR implementation

The software package was implemented in R, following Bioconductor standards. Basic usage examples are provided in **Supplemental Material** and in the package documentation. In principle, we encourage users to apply advanced data normalization, clustering, and cell type calling tools (12, 34), and to submit preprocessed count matrices to SingleCellSignalR to perform LR interaction inference and visualization. For convenience, we implemented simple data preparation steps enabling users to start from a raw read count matrix. We normalize individual cell transcriptomes according to their 99^th^ read count percentile. In case a cell has its 99^th^ percentile equal to zero, it is discarded. Normalized read counts (+1 to avoid zeros) are log-transformed. Clustering can be obtained either by chaining principal component analysis for dimension reduction and K-means (35), or by using the advanced SIMLR model (36). Cell type calling is implemented based on a list of gene signatures following a format identical to PanglaoDB (37) exports such that users can easily add cell types from this rich source or provide their own. Our algorithm computes average gene signature expression across all the cells and for all the signatures to obtain a matrix signatures × cells. This matrix is normalized and a threshold is iteratively adjusted to maximize the number of cells assigned to a single cell type. Full details and an example are in **Supplemental Material**.

It is possible to infer paracrine or autocrine only interactions, or both types (**Suppl. Fig. 1**); details in **Supplemental Material**. For annotation purposes, differentially expressed genes between each cluster and all the other clusters pooled, are successively searched with edgeR functions glmFit and glmRT (38). LR interactions with both the ligand and the receptor significantly enriched in their respective cell types are labeled “specific” (**Suppl. Fig. 1**).

In order to relate receptors to intracellular signaling, we make use of Reactome and KEGG (39) interactions downloaded from Pathway Commons. Interactions are assigned to several types that we simplified to facilitate the display of networks afterward. Interaction types “interacts-with” and “in-complex-with” were assigned to the simplified type “complex.” The interaction types “chemical-affects”, “’consumption-controlled-by”, “controls-expression-of”, “controls-phosphorylation-of”, “controls-production-of”, “controls-state-change-of”, “controls-transport-of”, and “controls-transport-of-chemical” were simplified as “control”. The interaction types “catalysis-precedes,” “’reacts-with,” and “used-to-produce” were simplified as “reaction.” The simplified type “control” was considered directional whereas “complex,” and “reaction” were considered undirected.

### Data sets

Different single-cell data sets were used to illustrate and benchmark SingleCellSignalR. MELANOMA is a metastatic melanoma data set covering several patients (40), SMART-Seq2 protocol, GEO GSE72056. 10xPBMC is a peripheral blood monoclonal cell data set from healthy donor (41), Chromium 2 protocol. 10xT is a pan T-cell data set from healthy donor (42), Chromium 2 protocol. 10xPBMC and 10xT were downloaded from 10x Genomics web site. HNSCC is a head and neck squamous cell carcinoma (primary and metastatic) data set (16), SMART-Seq2 protocol, GEO GSE103322. PBMC is a second, deeper peripheral blood monoclonal cell data set (43), SCRB-seq protocol, GEO GSE103568.

### Mouse skin immunolabeling

Fresh adult mouse skin samples were embedded in OCT (Sakura) and cryosectioned. Immunolabeling was performed on unfixed 10μm cryosections using the following antibodies: anti-PSEN1 (SAB4502423, Sigma) and CD44 (MA1-10225, ThermoFisher). For IF, we used anti-rat conjugated 488 (A-11006, ThermoFisher) and anti-rabbit conjugated 594 (A-21207, ThermoFisher) antibodies.

## RESULTS AND DISCUSSION

### Workflow overview

Independent of the chosen scRNA-seq platform, data come as a table of read or unique molecule identifier (UMI) counts, one column per individual cell, and one row per gene. The prediction of LR interactions between cells requires that scRNA-seq data are normalized and clustered, with each cluster corresponding to a cell type (12). Ideally, the cell types should be called, e.g., using gene signatures, to facilitate the interpretation of the LR interactions. The design of SingleCellSignalR is such that those preliminary steps can be accomplished within the package, using built-in or integrated solutions, or realized with other tools according to the user preference. The latter option is preferable to benefit from the latest or most advanced techniques. Once data have been prepared, the inference of LR interactions is performed by successively considering each possible cell type – or cluster – couple, e.g., CD8+ T cells *versus* macrophages (Fig 1D). The reality of each potential LR pair according to LR*db* is assessed by the computation of a score based on gene expression in the respective cell types. It is also possible to infer autocrine interactions. The output interaction lists are provided in various formats (tables, different plots, and networks), and complementary functions were designed to help to interpret these lists, e.g., linking receptors to intracellular signaling pathways from Reactome and KEGG.

### Scoring ligand-receptor pairs

To determine reliable LR interactions between cell types *A* and *B*, we interrogate LR*db* and score each LR pair found with average ligand expression *l* > 0 in *A*, and average receptor expression *r* > 0 in *B*, or *vice versa*. Existing tools often consider the product *lr* and sometimes the average 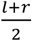. Alone, such scores can rank LR pairs, but users are left without any clue where to cut off likely false positives. The computation of a score should hence be accompanied by a procedure to determine a threshold below which scores are deemed unreliable. A common choice is to shuffle cell type assignments multiple times and to obtain a score null distribution to estimate score P-values. Although intuitive and statistically sound, this solution does not address the real question. It rather identifies LR pairs that are significantly specific to a given couple of cell types. That should imply that the interaction is likely to exist but it might result in poor P-values for real LR interactions that are shared by many couples of cell types. For instance, in tumors, it is common that many immune cell populations express immune checkpoints and their ligands, e.g. PD-1/PD-L1, forming an immunosuppressive microenvironment. In such a case, many combinations of cell types would involve the PD-1/PD-L1 interaction resulting in bad P-values if an insufficient number of other cell populations, not expressing these molecules, would be present in the data set. To address the points above, we introduced a regularized product score:

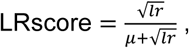

where *μ* = mean(*C*) and *C* is the normalized read count matrix. In this empirical score, the mean *μ* acts as a scaling factor, and the square roots are meant to keep the *lr* products and *μ* on the same scale. The LRscore is bounded by 0 and 1, independent of the data set depth.

Remains the problem of defining a score threshold. This is not a simple task since, by essence, all LR pairs listed in LR*db* are correct in a particular context, i.e., defining false LR interactions must rely on external knowledge specific to the couple of cell types considered each time. This knowledge does not exist for all possible LR pairs and cell type combinations. To estimate an appropriate LRscore threshold and to compare LRscore to other scoring schemes, we hence needed an *ad hoc* benchmark that would be unbiased regarding the scoring schemes and accurate enough biologically. A first opportunity was provided by Ramilowski *et al*. (6) data, which are part of FANTOM5 and included a table reporting the expression (in TPM) of many ligands and receptors over 144 primary cell types. This table, which we denote *T*_ref_, was obtained by sequencing techniques that were much deeper than scRNA-seq, i.e., virtually devoid of dropout for our purpose. The authors considered that expression above 10 TPM could be taken as a conservative gene expression basal limit. We thus designed an evaluation where candidate LR*db* pairs, restricted to ligands, receptors, and cell types covered in *T*_ref_, could be considered correct if both the ligand and the receptor TPM were above 10. If both were below 10 TPM, the pair was deemed false, and mixed cases (one >10 and the other ≤10) were ignored. Admittedly, ligand and receptor concomitant expression does not guarantee a functional interaction, but since those ligands and receptors are known to interact for at least one combination of cell types, we considered the benchmark above sufficiently accurate for our purpose. A second opportunity was provided by a proteomics study of 28 primary human hematopoietic cell populations in steady and activated states (44), which enabled a similar benchmark relating censored scRNA-seq data to their proteomics counterpart. Namely, each cell population was measured in quadruplicate and we stored the average peptide counts in a table *P*_ref_ (similar to *T*_ref_). This second evaluation was conducted as above with a threshold of an average spectral counts ≥ 2 for an LR pair to be true.

We used ROC curves to compare methods and determine score thresholds at 5% false positives (FPs). We already mentioned LR pair scores equal to the product *lr* or the mean 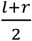 as well as their P-values. A further proposed scheme was to select in each cell type, those genes whose expression was specific. Subsequently, whether the reference database would contain pairs of such genes was checked (22, 24). In the case of Zhou *et al*. (22), gene selection relied on a z-score calculation. To obtain a ROC curve, we varied the coefficient applied to the standard deviation at that gene selection stage. Another selection-based tool (24) did not report any score. It is thus not included here but covered in the tool comparison below only. scTensor (23) applied non-negative Tucker decomposition to model ligand and receptor read counts as sums over the many LR pairs they might contribute to. Despite considerable efforts, we could not use it beyond its pre-packaged example. It is therefore absent from the presented comparison.

We applied the two benchmarks above to five data sets covering several cell types in *T*_ref_ and in *P*_ref_ (Materials and Methods). Every couple of cell types, both in *T*_ref_ or *P*_ref_, yielded ROC curves such as the example featured in Fig. 2A for 10xPBMC data against *T*_ref_. Areas under the curves (AUCs) over all the data sets and all the cell type couples represented in *T*_ref_ are presented in Fig. 2B; individual curves and data set specific AUCs are in **Suppl. Figs 2-6**. We see that the best AUCs were achieved by LRscore, the product, and the average. As expected, P-value calculations yielded inferior performance. Zhou *et al.* selection mechanism was also inferior to the best scores. We next asked how variable a threshold aimed at cutting at a certain FP rate would be; we imposed 5% FPs (Fig 2C and **Suppl. Figs 2-6** for data set specific plots). Naturally, due to its design, LRscore thresholds turned out to be much more stable. In Fig. 2D, we show how a threshold common to all the single-cell data sets could be determined, imposing less than 5% FPs in 75% of all the ROC curves of all the 5 data sets. We found LRscore > 0.4. Repeating the analysis for the proteomics reference *P*_ref_ gave the results in Figs. 2E-F (and **Suppl. Figs 7-11**) that are very similar. The computation of a common threshold to achieve less than 5% FPs in 75% of the ROC curves resulted in LRscore > 0.6 (**Suppl. Fig. 12**). This more stringent threshold can be explained by a potential lower sensitivity (~10,000 proteins) compared to FANTOM5 transcriptomics and by the obvious differences between transcriptome and proteome. These considerations indicate that a universal threshold that would guarantee, e.g., 5% FPs cannot be determined, but score regularization is already a major step towards more rigorous cutoffs. To illustrate this, we imposed LRscore > 0.5 to all the data sets against both *T*_ref_ and *P*_ref_. From Fig. 2G, we see that we managed to maintain the FPs in a reasonable range, and, most importantly, the variability of the FP rate within each data set is modest. That is, a threshold can be set and FP rates on LR interactions between any combinations of two cell types remain comparable. Repeating this analysis with the product score and common threshold 10^−2^ found as above, we note large discrepancies between data sets.

**Figure 2.**
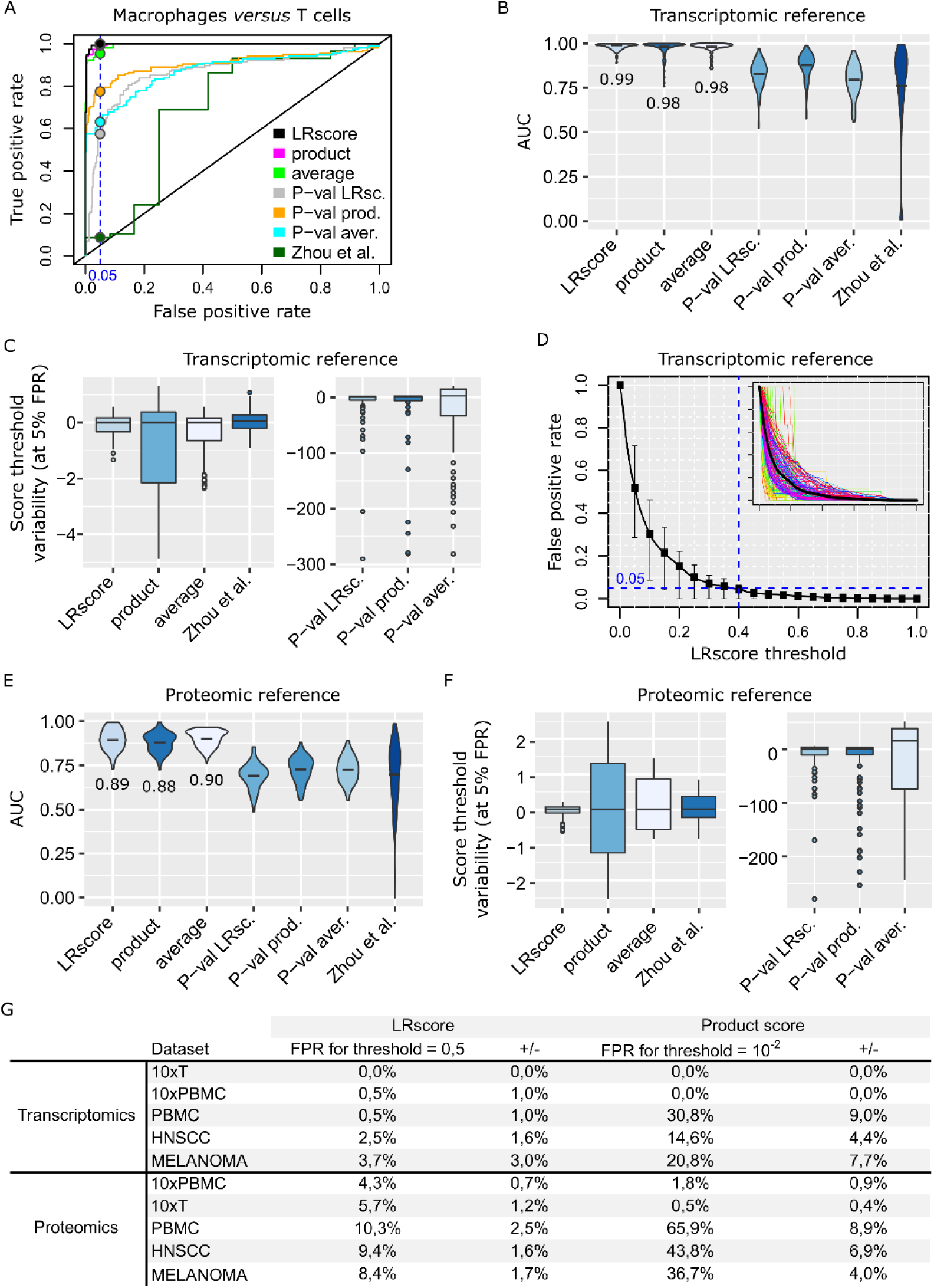
Statistical analysis. **A.** Representative ROC curve. **B.** Areas under the curve (AUCs) over all the ROC curves of the five data sets; transcriptomics reference. **C.** Relative variability (with respect to the median value) of the thresholds required to achieve 5% FPs in each ROC curve (all couples of cell types, all the data sets). **D.** FP rate upon various LRscore thresholds. A small box plot is featured for each threshold value and the figure inset shows all the ROC curves. LRscore threshold (blue dashed line) such that 75% of the ROC curves would yield FPs below 5%. **E,F.** Same as B,C, but for the proteomics reference. **G.** FP rates (+/− sd) on each data set against the transcriptomic and the proteomic references when imposing a consensus LRscore threshold of 0.5. Same results for an equivalent product score threshold consensus in the two right-most columns.

### Representing ligand-receptor interactions

After scoring and application of a threshold, SingleCellSignalR outputs LR interactions in various formats. 10xPBMC raw UMI counts were submitted to our default pipeline, which clustered cells in six populations: B cells, T cells, regulatory T cells (Tregs), neutrophils, cytotoxic cells, and macrophages (Fig. 3A). A summary chord diagram can be generated that indicate the number of LR interactions between each cell population couple (Fig. 3B). Chord diagrams for individual interactions between two populations are possible as well (Figs. 3C). Inspection of the expression of the ligand and the receptor involved in a LR interaction is implemented in mixed or dual 2D projections (Fig. 3D-E).

**Figure 3.**
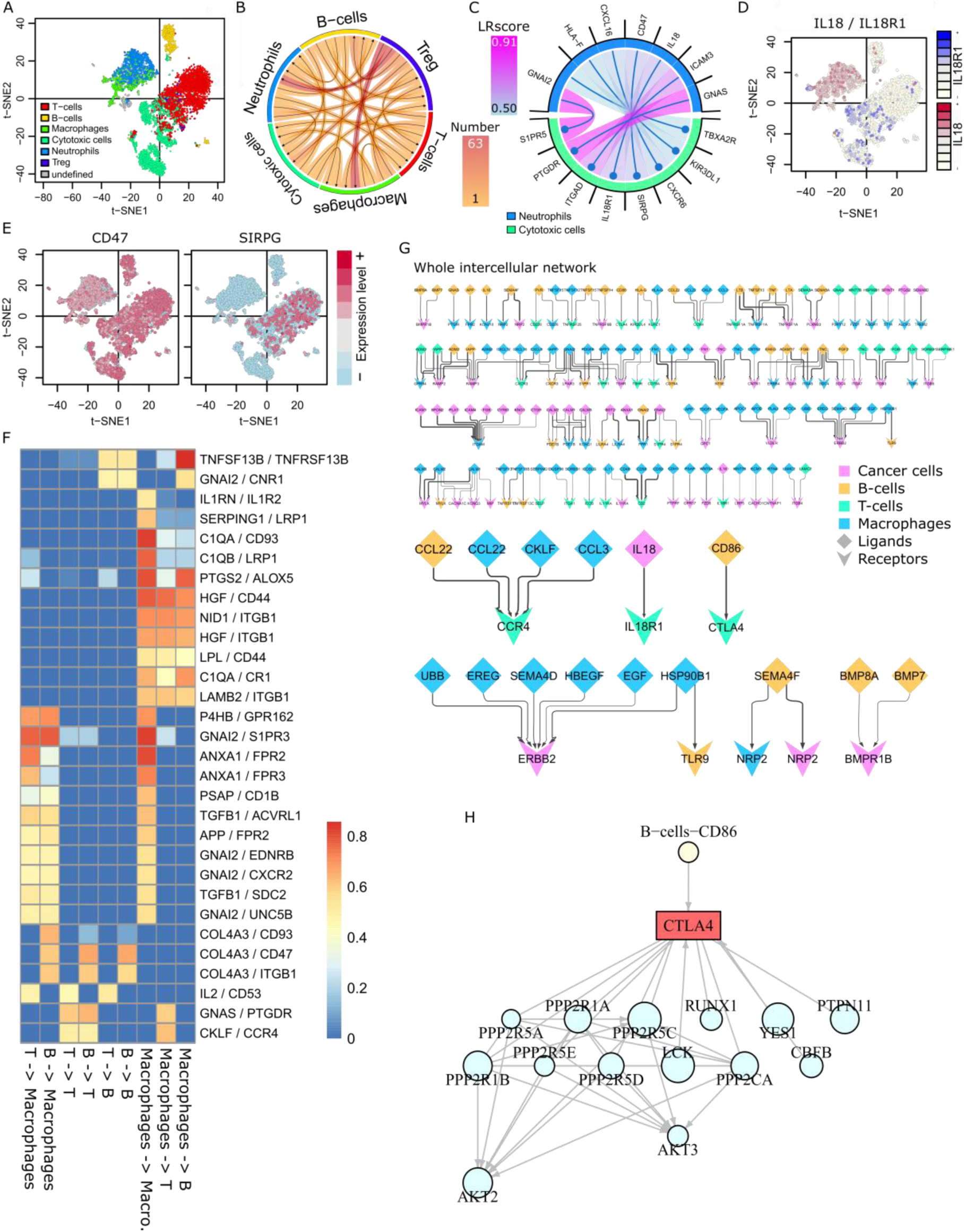
Graphical representations. **A.** 10xPBMC data with SIMLR clusters. **B.** Summary chord diagram of the paracrine interactions; largest number of interactions from regulatory T cells towards macrophages and neutrophils. **C.** Paracrine interactions from neutrophils towards cytotoxic cells. **D, E.** Joined, and separated expression plots over the t-SNE map to assess LR interaction specificity and prevalence. **F.** Integrated tabular view of the *N* most variable LR pairs with LRscore > 0.5 in one cell type couple at least. **G.** Integrated network of MELANOMA data patient 89 intercellular interactions. Overview and chosen interactions. The full network is in **Suppl. Fig. 13**.

Chord diagrams are convenient for looking at specific couples of cell types but not to account for the whole data set. We hence implemented two integrated views as either a tabular plot (Fig. 3F) or a network exported in graphML format. The latter can be imported in tools such as Cytoscape (45). We exemplify the network view with MELANOMA patient 89 data. In Fig. 3G, we picture an overview of the intercellular network (full details in **Suppl. Fig. 13**). The complexity and partial redundancy of this network is particularly noteworthy, with multiple cell types expressing the same ligands or receptors. That is typical of immunosuppressive TMEs.

### Putting ligand-receptor interactions in context

To help understanding the functional consequences of LR interactions, we relate the receptors in each cell type with downstream biological pathways (Fig. 3H). This requires a global reference of all possible pathways, including links from the receptors, and a method to generate receptor-related, intracellular regulatory networks that are restricted to each cell type. We decided to use KEGG (39) and Reactome (32) pathways taken from Pathway Commons (33) as global reference. This choice guaranteed interpretable, literature-supported network interactions. Surprisingly, we found 100 receptors in LR*db* that were devoid of downstream interactions in this pathway ensemble. By manually checking their respective UniProtKB/Swissprot entries, we found 1-12 known interactions for 61 of them, which were added along with literature references (176 interactions in total).

Construction of a ligand-related internal network for each cell type was obtained by the following algorithm. Given a list of receptors, e.g., all the receptors involved in LR interactions for the cell type at hand or a subset, we first identify all the pathways including these receptors. The union of all such pathways is intersected with the set of genes expressed by the cell type. That gene lists is obtained by selecting the genes expressed by more than a proportion *p* of the individual cells after normalization of the raw read count matrix, that is in matrix *C* (*p* default is 20%; it can be adjusted by the user). To the intersected pathways, we add direct receptor/expressed gene interaction that would not be included in Reactome or KEGG, which are those we manually added. The functionality can be applied to ligands as well to obtain information about their pathways of origin. In every case, the resulting network edges are annotated with Reactome/KEGG data plus a simplified interaction typology that is convenient for visualization (Materials and Methods).

The neutral and straightforward procedure above for calling expressed genes was retained to be independent of the cell population size, on the one hand, and to avoid interference with raw read count preprocessing, on the other hand. In particular, users could apply imputation strategies (46) to reduce dropouts and/or advanced data normalization and regularization (12, 47).

### Mouse interfollicular epidermis application

Mouse interfollicular epidermis (IFE) is a multilayered epithelium in which proliferating cells reside in the basal layer (IFE B), where they undergo regulated cell division. Their daughter cells move upwards into the suprabasal layers while further differentiating until they reach the outermost layer (IFE K2) (Fig. 4A). In this application, we demonstrate SingleCellSignalR mouse data functionality, which is implemented by internally mapping mouse genes to their human orthologues according to Ensembl (48) to exploit LR*db*. Murine gene names are preserved in the different outputs.

**Figure 4.**
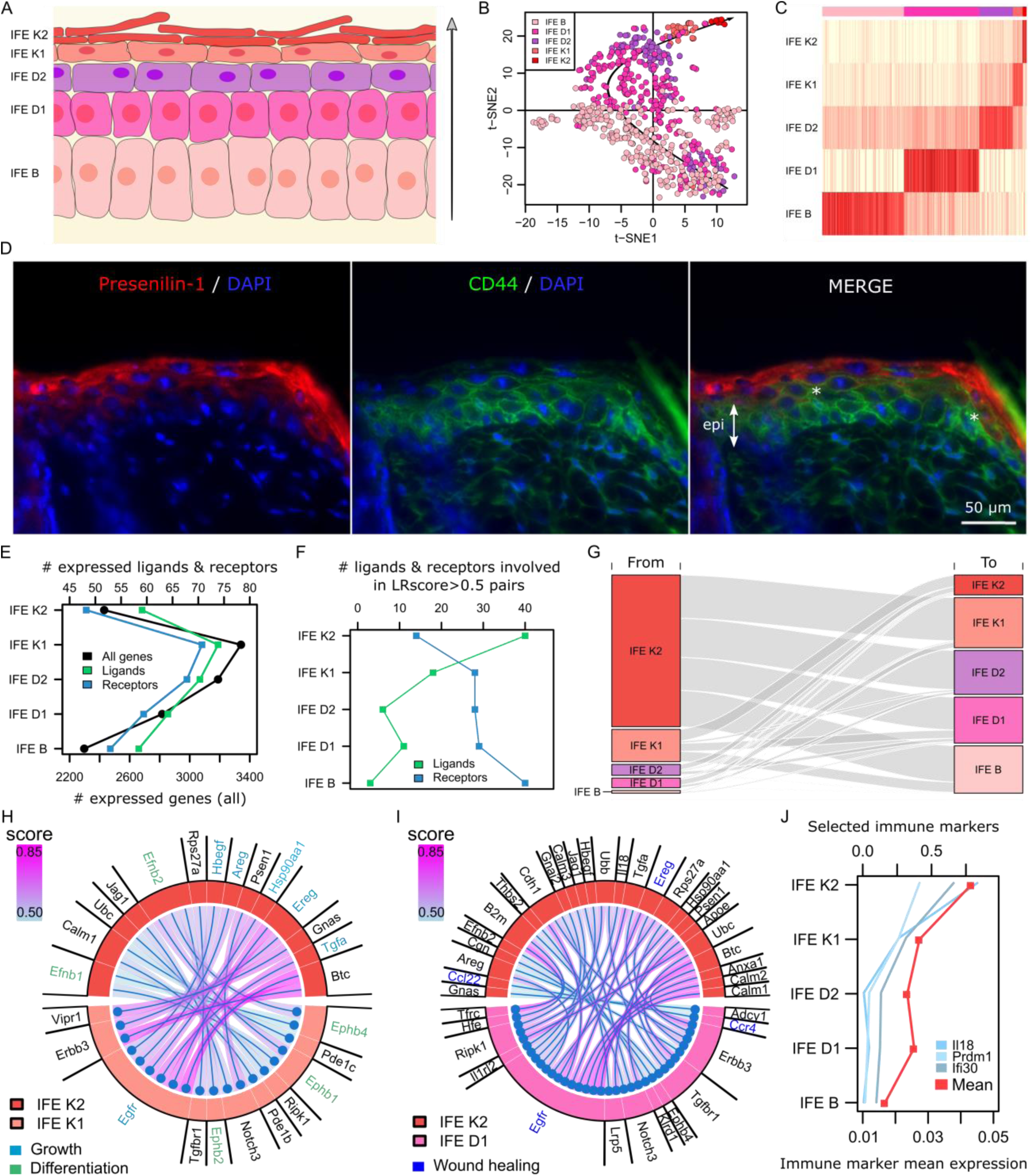
Mouse interfollicular epidermis. **A.** Schematics of the interfollicular epidermis (arrow=axis of differentiation). **B.** t-SNE plot of the IFE cells with the underlying black arrow representing the axis of differentiation. Some incompletely differentiated non IFE B cells lie among the IFE B subpopulation. **C.** Cell type calling with actual subpopulations in the top color band. **D.** Presenilin-1 (left panel) and CD44 (middle panel) immunostainings of mouse skin sections. Sections were counterstained with DAPI (epi=epidermis, *=co-localization). **E.** Number of genes with average expression different from 0 in each IFE layer. **F.** Number of ligands and receptors involved in LRscore>0.5 interactions. **G.** Flow diagram representing the number of interactions between the cell subpopulations. **H, I.** Chord diagrams of adjacent and remote IFE K2 cell-cell interactions. **J.** Immune marker gene expression. Il18 is expressed by macrophages, Ifi30 by antigen-presenting cells, and Prdm1 by T cells (genecards.org).

From Joost *et al*. data (49), we selected IFE cells exclusively (658 cells, Fig. 4B-C). To confront these data with our FP analysis above, we considered LR interactions between the keratinized layer (IFE K2 and K1 cells pooled) and the suprabasal layer (IFE D1 and D2 cells pooled). The pooling was motivated by minor differences between cells and a facilitated comparison with human epidermis (see below). We found 248 LR interactions with LRscore > 0.5 (**Suppl. Table 1**). Among the top-ranked interactions, several cases involved Presenilin-1 (Psen1). Based on novelty and specific antibody availability, we decided to validate in mouse epidermis the interactions Presenilin-1/CD44, which is likely involved in cell differentiation and tissue organization. From Fig. 4D, we observe correct localization of those two proteins in the IFE layers (left and middle pictures), Presenilin-1 being both cytoplasmic and secreted in agreement with the literature. In the merged picture (right), we see an overlap between the diffusive, extracellular Presenilin-1 signal and suprabasal cell plasma membranes harboring CD44 in some regions, e.g., at the two locations indicated by white asterisks. Using the Human Protein Atlas (HPA) (50) as a systematic resource to search human orthologs expression in the corresponding epidermis layers, literature (51–53) and the above experimental validation, we confirmed 158 out of the 176 inferred LR interactions for which we had data (**Suppl. Table 1**). That is 10.2% FPs, in line with our ROC curve analyses above compared to the proteomics reference.

Top layer IFE K2 cells supposedly maintain elementary activities only, which was supported by the limited number of expressed genes found in those cells (Fig. 4E). The number of expressed genes coding for ligands or receptors was directly correlated with the total number of genes in Fig. 4E, reflecting the proportion they occupy in the mouse genome. This configuration changed considering ligands and receptors that were involved in reliable LR interactions only (LRScore > 0.5). The largest numbers of ligands were found in upper layers and the receptors followed a reversed pattern (Fig. 4F). The overall LR interactions are quantified in Fig. 4G, which shows an IFE K2 layer sending most signals and receiving the least. The number of LR interactions with a ligand contribution diminish from the outer layer to the most basal one. That is, the application of SingleCellSignalR unraveled upper IFE layer cells that produce much information but receive little. The basal layers behaved anti-symmetrically. LR interactions between IFE K2 and K1 and more remotely between IFE K2 and D1 are reported in Fig. 4H-I. Pathway analysis showed that the interactions issued from IFE K2 cells were mostly involved in growth (*e.g.* GAB1 signalosome, or signaling by EGFR) and differentiation (EPH-ephrin mediated repulsion of cells, or Ephrin signaling). That illustrates the notion of a tight long-distance connection between outer and basal layers, which might be of particular interest in understanding the regulation of the epidermis constant self-renewal.

Interestingly, we noticed that LR pairs involved in wound healing regulation (54) were expressed by IFE K2 and D1 (Fig. 4I). That suggests a mechanism for rapid reaction upon wounding, bringing K2 and D1 cells in contact. Although we excluded the immune cells present in Joost *et al*. data, we found 146 genes associated with immune cells. Selected genes with a clear pattern, as well as the average of all immune-related genes, are reported in Fig. 4J. IL18 and PRDM1 were found with epidermis keratinocyte expression in human skin according to HPA (**Suppl. Fig. 14**). This striking observation strengthens the potential for an immune function of the epidermis (55, 56) and may have essential implications in inflammatory skin diseases such as psoriasis or atopic dermatitis.

### Comparison with other tools

The general characteristics of SingleCellSignalR and related tools appear in Table 1. iTALK (25) infers LR pairs from a compilation of public LR databases using a product score limited to the most abundantly expressed genes. It requires a read count matrix, where cell type calling was performed beforehand. iTALK includes a feature to deal with multiple data sets (time course, different conditions) such that variability and trends in ligand and receptor expression across data sets can be added to the LR plots. CellPhoneDB (26) is both an online tool and a Python package that can be downloaded. It scores LR pairs after P-values of the mean score. A modification is operated in case of a multimeric receptor and/or ligand, requiring that all the subunits are expressed. This unique and biologically sound feature might cause false negatives due to dropouts in scRNA-seq data. Kumar *et al*. (18) employed a product score. Instead of simple data shuffling, they applied a statistical test (Wilcoxon) to assess that over several tumors, the score was significantly different from zero. That procedure that cannot be reproduced when analyzing a single data set. Zhou *et al*. (22) proposed to select LR pairs based on specific expression of the ligand and the receptor in two cell subpopulations, referring to a database of known LR pairs. Specific gene expression was tested, requiring that the average expression of a gene, in a subpopulation, was above its mean expression over the whole count matrix plus 3 standard deviations. scTensor (23) aims at inferring LR pairs through non-negative Tucker decomposition, explaining the observed ligand and receptor read counts as sums of the contributions of all the interactions they would engage in. It exploits a database of potential LR pairs generated from STRING interactions and Swissprot annotations (secreted/membrane) automatically, which yields a considerable number of putative LR pairs and does not cover many known interactions (Fig. 1C). One attractive scTensor feature is the LR reference available for multiple organisms. PyMINEr (24) is a Python program implementing a complete pipeline to process raw read counts. Cell type calling is based on subpopulation specific gene pathway enrichment; application of characteristic gene signatures is not available. It is not possible to prepare and cluster data externally. LR pairs between two cell subpopulations are inferred from pairs of genes found to be specific to each subpopulation, in interaction in STRING (57), and respectively classified as receptor and ligand in GO. There is no scoring *per se*, only this selection process. PyMINEr further infers a so-called co-expression graph, linking genes correlated over the whole read count matrix. This graph can be exploited to project different data such as the expression of genes in one specific subpopulation. Like scTensor, PyMINEr relies on automatically generated LR references from STRING.

**Table 1.** Software tool comparison.

|  | SingleCellSignalR | iTALK | PyMINER | CellPhoneDB | scTensor | Zhou *et al.*<sup>1</sup> | Kumar *et al.* |
| --- | --- | --- | --- | --- | --- | --- | --- |
| LR database size | 3,251 | 2,648 | NA<sup>2</sup> | 1,396 | 34,449 | 2,558 | 1,800 |
| Complete pipeline | Y | N | Y | N |  | N | N |
| Accept preprocessed data | Y | Y | N | Y |  | Y | Y |
| Multiple samples | N | Y | N | N | N | N | N |
| Intracellular signaling | Y | N | Y | N | N | N | N |
| Scoring approach | regularized product | product | selection | modified<sup>3</sup> mean P-value | linear decomposition | selection | product<sup>4</sup> |
| *Export types* |  |  |  |  |  |  |  |
| tables | Y | Y | Y | Y |  | NA | Y |
| circular plots | Y | Y | N | N |  | NA | N |
| graphML or equiv. | Y | N | Y | N |  | NA | N |
| Num. species | 2 | 1 | 1 | 1 | 12 | 1 | 2 |
| Platform/language | R | R | Python | web site or Python | R | NA | MATLAB |

<sup>1</sup>Not available as a software package.
<sup>2</sup>Large reference database of LR interactions derived from STRING, comparable to scTensor, actual size not known.
<sup>3</sup>In case a complex is considered, all subunits must be selected in a significant interaction with the partner.
<sup>4</sup>The authors added a Wilcoxon test to assess that the LR pair score was different from zero in multiple tumors.

As explained above, we could not use scTensor beyond its prepackaged example data set (58) despite considerable efforts. We nonetheless tried to obtain some elements of comparison by processing the same data with SingleCellSignalR. We found 2 reliable LR pairs whereas scTensor returned 14 pairs, none in common, see **Suppl. Table 2**. Not being able to test scTensor on more examples did not allow us to perform a real comparison. We analyzed 10xPBMC data with PyMINEr and observed a substantial discrepancy with our results. Without imposing any threshold on PyMINEr output (no guidance provided by the authors), PyMINEr typically retrieves 10-20 times more paracrine interactions, and twice the number of pooled paracrine and autocrine interactions, with little interactions with our LRscore > 0.5 results (**Suppl. Table 3-4**). The ROC curve analysis we conducted and such enormous lists of interactions inferred by PyMINEr suggest contamination with FPs. Obviously, PyMINEr output should be imposed a threshold in practice, which would reduce the FPs but would not improve the overlap with the known interactions in LR*db*. We next submitted the same 10xPBMC data to CellPhoneDB web site. The smaller LR reference database, strict rules regarding multimeric ligands or receptors, and the impact of a re-shuffling-based P-value calculation caused reduced sensitivity compared to our solution by a factor 8.5 on average (**Suppl. Table 5**). Inspection of CellPhoneDB output showed that the interactions we missed were absent from LR*db* and imported in CellPhoneDB from sources that we did not consider, such as InnateDB, MINT, IntAct, MatrixDB, or manual curation. Finally, we installed iTALK R package and applied it to 10xPBMC data as downloaded from iTALK GitHub repository. As for PyMINEr, iTALK scores and orders the LR interactions but does not propose any cutoff. Contrary to PyMINEr, iTALK relies on a selective database of LR pairs. We hence decided to consider the 118 top-scoring pairs of its output since LRscore > 0.5 gave us an average of 118 selections. Results are in **Suppl. Table 6** and we observe a good overlap with our larger selections. Dissecting the data showed that all of iTALK unique LR interactions were caused by LR pairs with deprecated HUGO gene symbols; they were present in LR*db* with the current symbols. For CellPhoneDB and iTALK, it was not possible to clearly separate paracrine from autocrine, and we hence compared results on the pooled paracrine and autocrine predictions. Overall, we found significant discrepancies with PyMINEr and scTensor, while CellPhoneDB and iTALK were more comparable with our tool. In the latter two cases, differences originated from reference databases and scoring.

## CONCLUSION

In the nascent topic of intercellular network mapping, SingleCellSignalR is an R software package that facilitates the transformation of complex data into higher-order information. The package comes with a large variety of graphical representations and export formats to accommodate users and to support the downstream analysis of data. Of particular note are the abilities to represent a complete intercellular network and to import the latter in systems biology tools such as Cytoscape, and to explore receptor downstream signaling by integrating Reactome and KEGG pathways.

For the first time, we discussed the question of the significance of inferred LR interactions. We showed that SingleCellSignalR regularized score achieves better control of the false positives, independent of the retained single-cell platform. That is an important feature to explore intercellular communication networks, where each cell type might be involved in a different number of interactions. It allowed us to evidence cells that emit more signals than they receive in mouse interfollicular epidermis (Fig. 4).

A detailed comparison with existing tools revealed the specific features contributed by each solution. The particular design choices we made, e.g., to rely on well-documented sources of interaction data and to control false inferences, clearly produced more or more reliable information. SingleCellSignalR is open to other software packages; UMI or read count matrices prepared with other tools can be imported, or preparatory steps can be accomplished with SingleCellSignalR built-in procedures. Although SingleCellSignalR was designed for scRNA-seq data, it could be used with emerging single-cell proteomics technologies such as CyTOF (59) or SCoPE-MS (60). Finally, our R library does not require advanced R skills from users.

Supplementary Data are available at NAR online.

## Supporting information

Supplemental file 2 - Human Protein Atlas (HPA) images

## AVAILABILITY

The R package and LR*db* are available from https://github.com/SCA-IRCM under the GPL v3 license; submitted to Bioconductor.

## SUPPLEMENTARY DATA

Supplementary figures, tables and methods are in SupplementalMaterial.pdf. Human skin images taken from HPA are in HPA.pdf.

## ACKNOWLEDGEMENT

We thank our colleagues Emmanuel Cornillot and Andreï Turtoi for their insightful comments.

