## Supplemental file 2 - Human Protein Atlas (HPA) images for "SingleCellSignalR: Inference of intercellular networks from single-cell transcriptomics"

- Confirmed ligand-receptor interactions names in black, e.g., Cdh1/Ptprf.
- Not confirmed in HPA, names in red, e.g., Ptn/Sdc1.
- Not confirmed in HPA, but with literature or IF names in green, e.g., Hbegf/Cd9.
- Mouse gene names are provided, human ortholog used in HPA.
- Low expression considered as the minimum required.

Cdh1 Ptprf

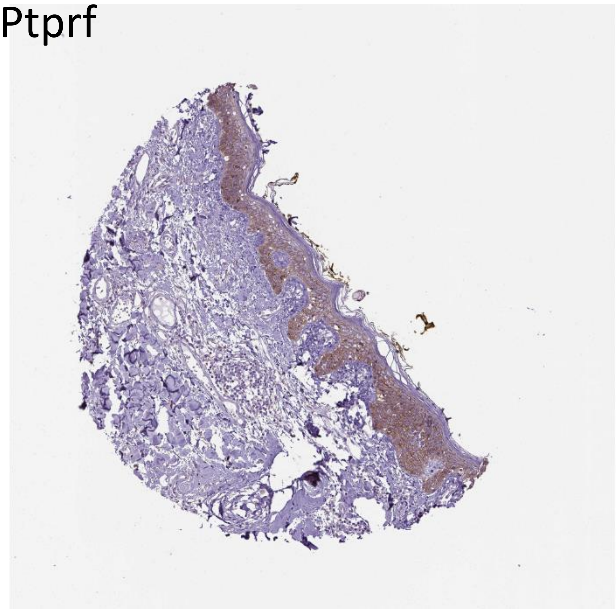
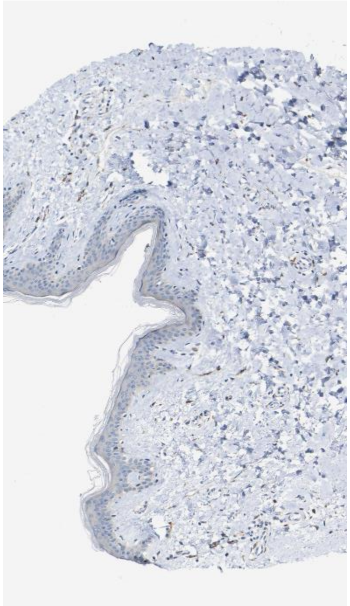

Hbegf Cd9

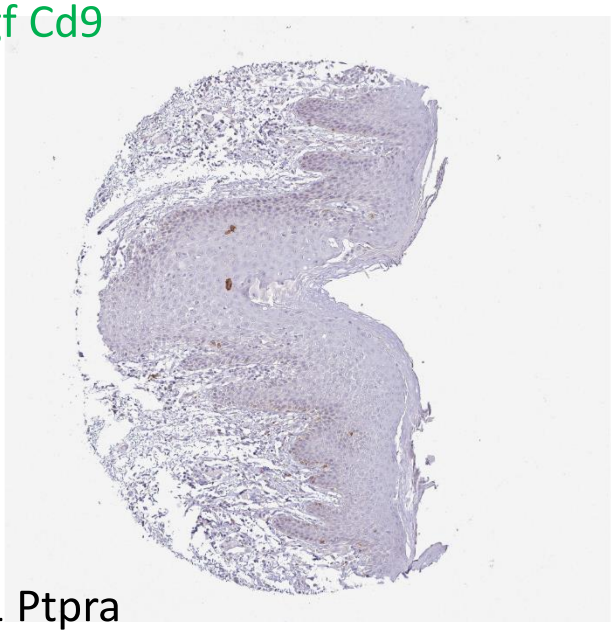
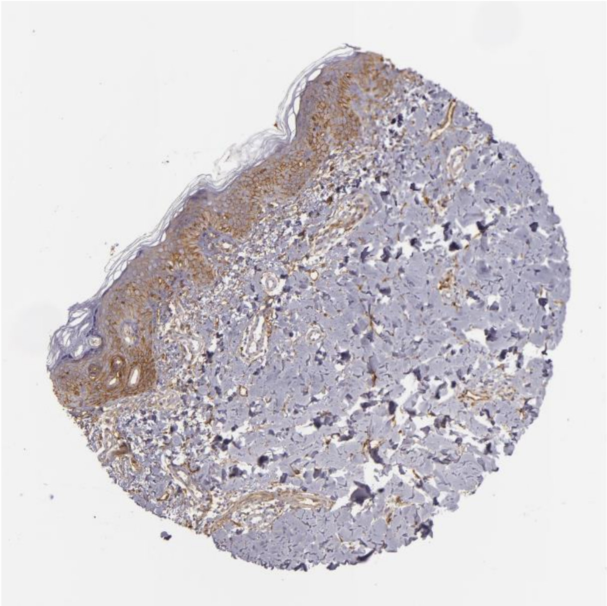

Calm1 Ptpra

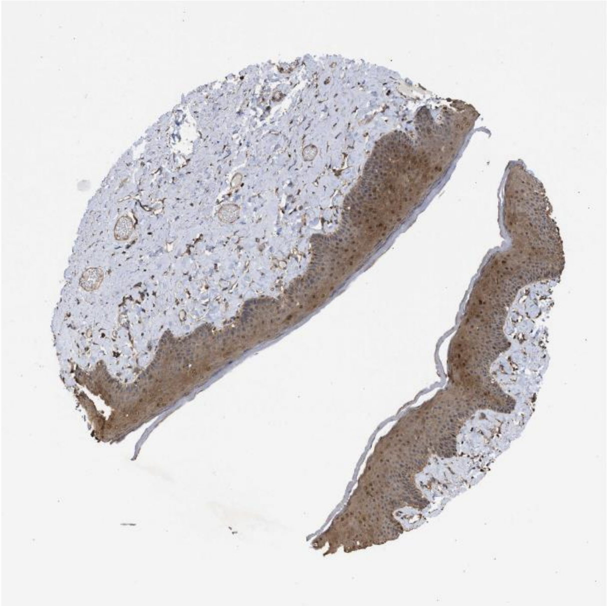
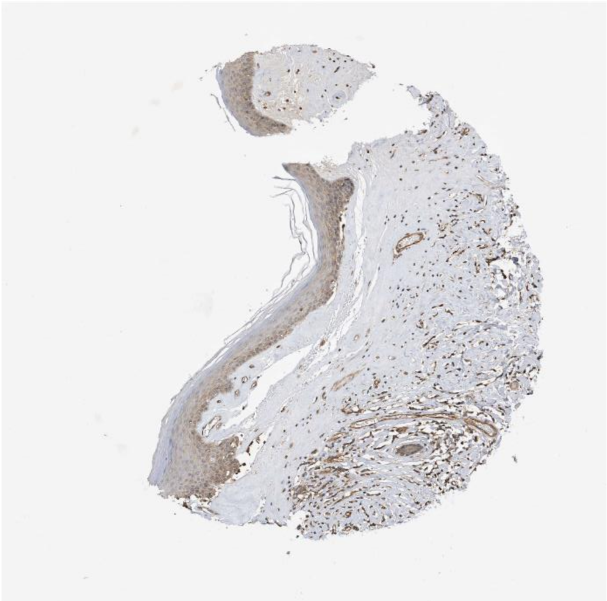

Calm1 Egfr

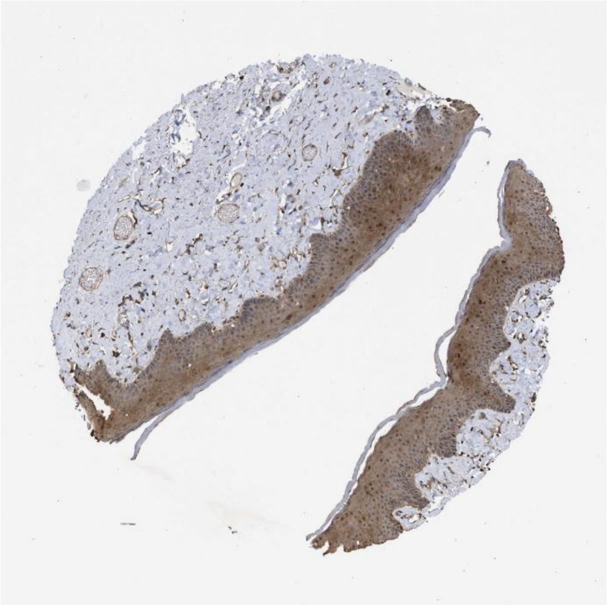

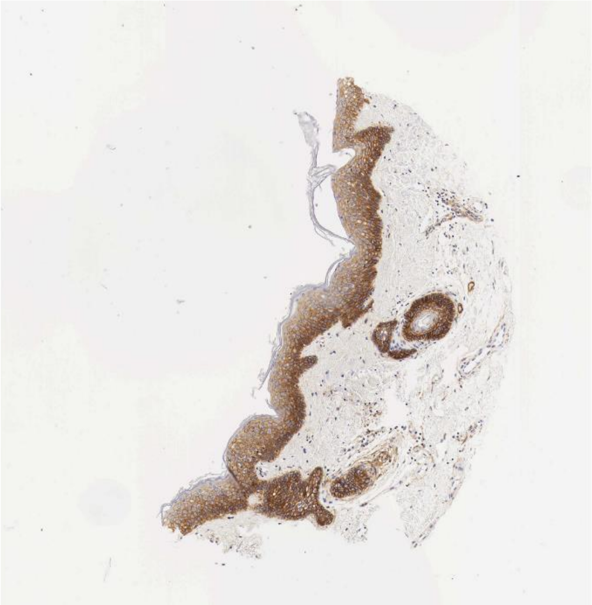

Ubc Ldlr

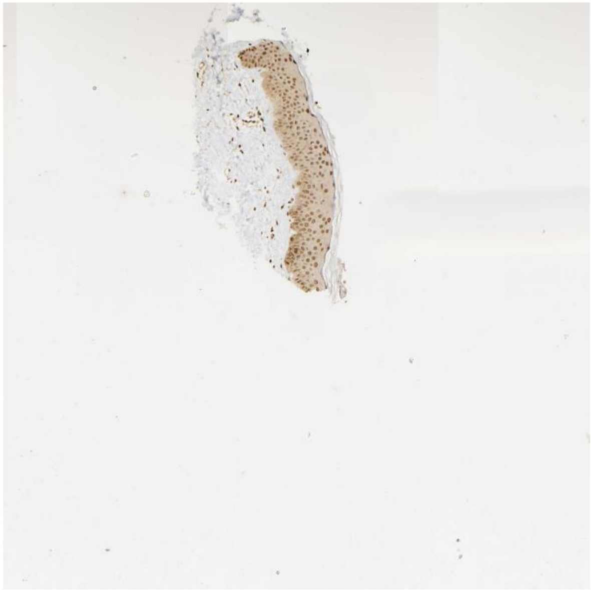

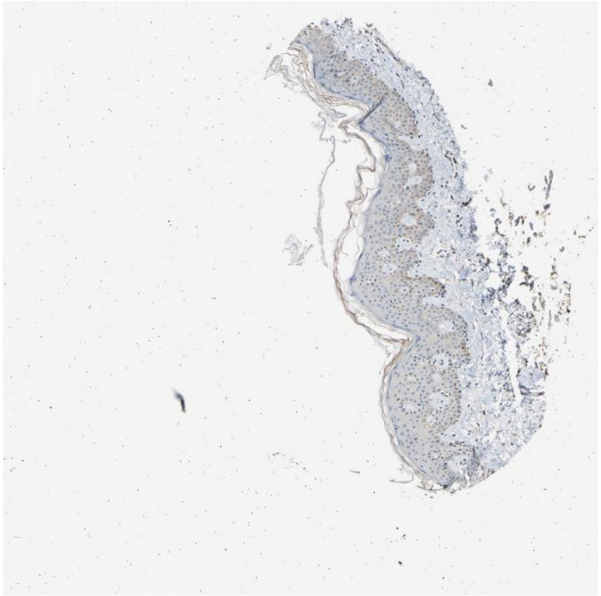

Calm2 Egfr

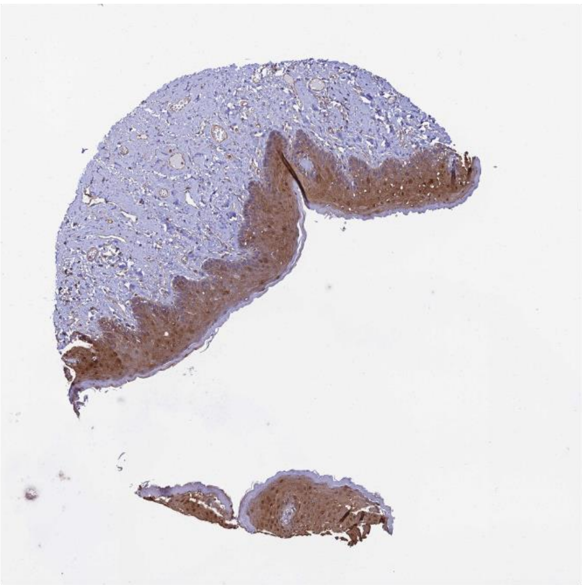

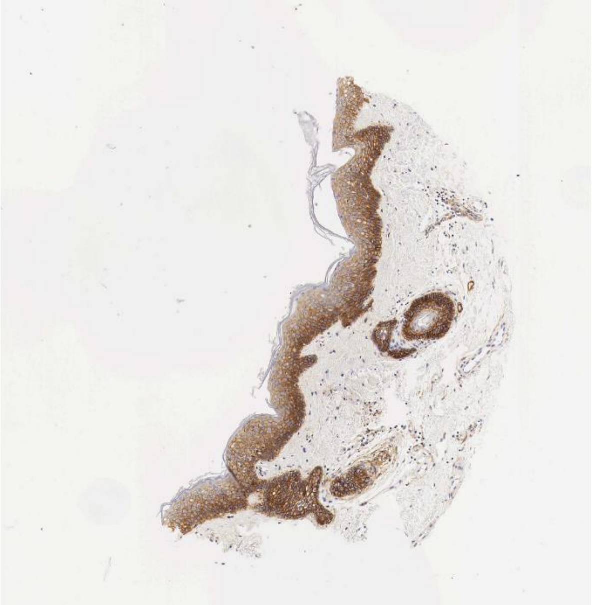

Ubc Erbb2

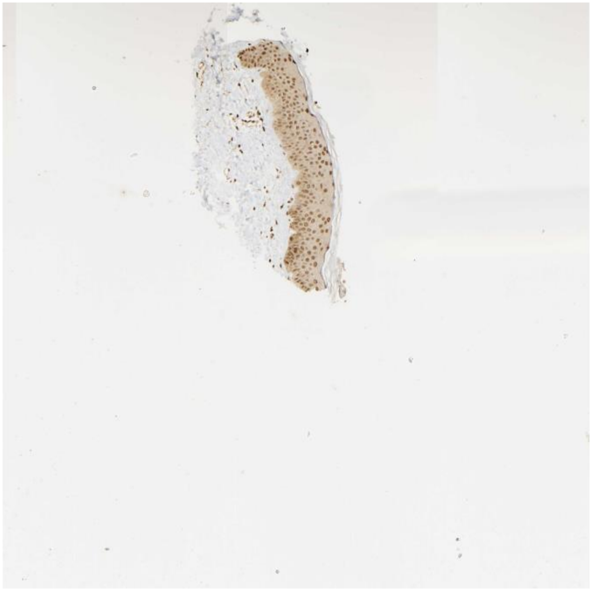

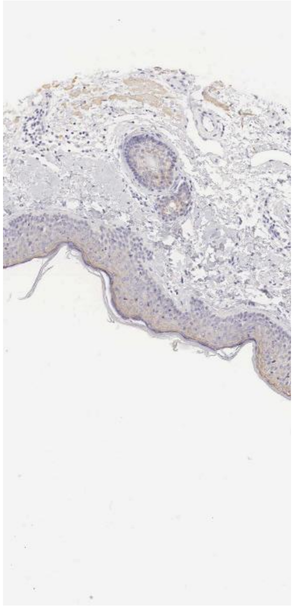

Rps27a Ldlr

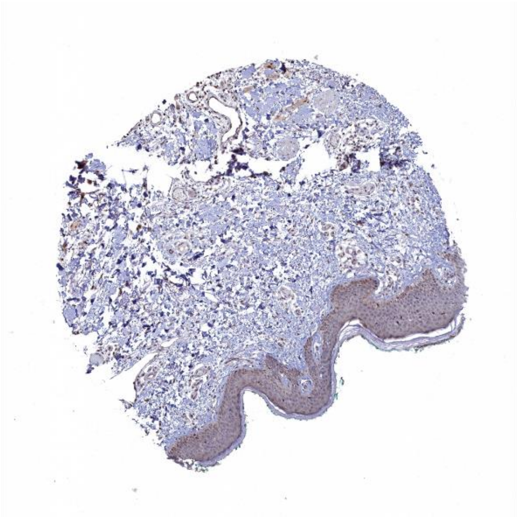

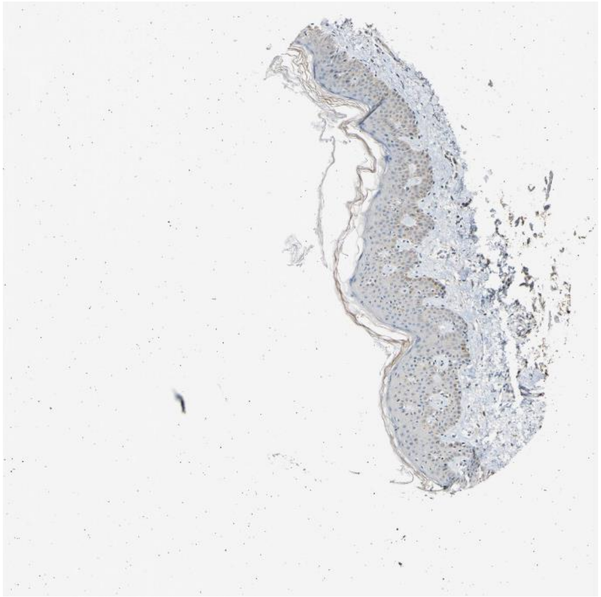

B2m Tfrc

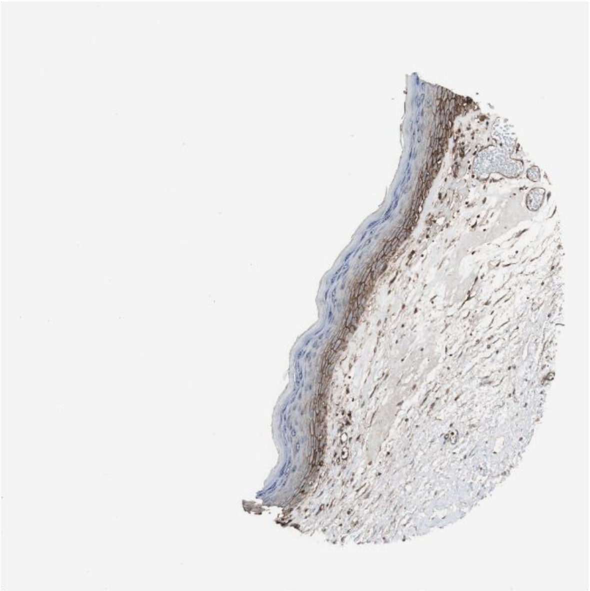

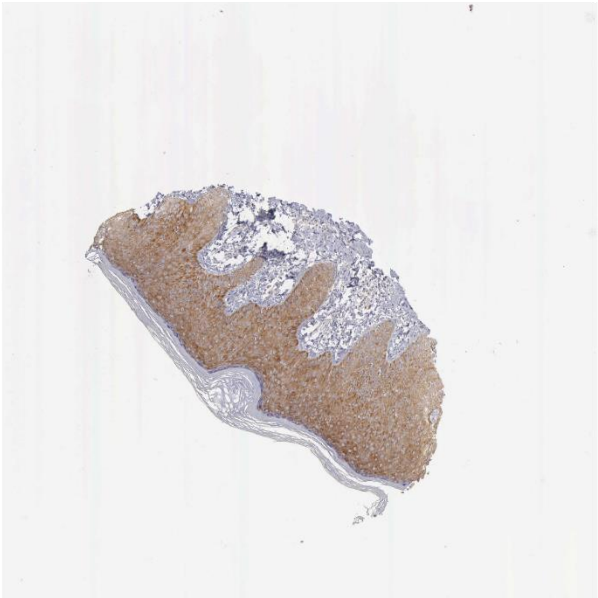

Apoe Ldlr

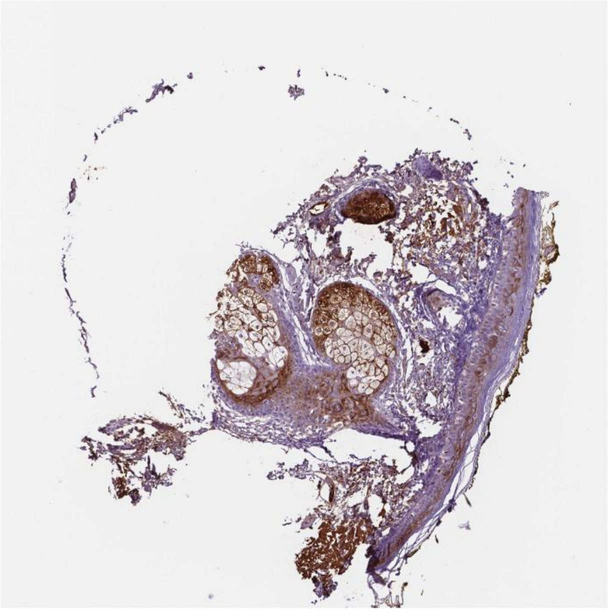

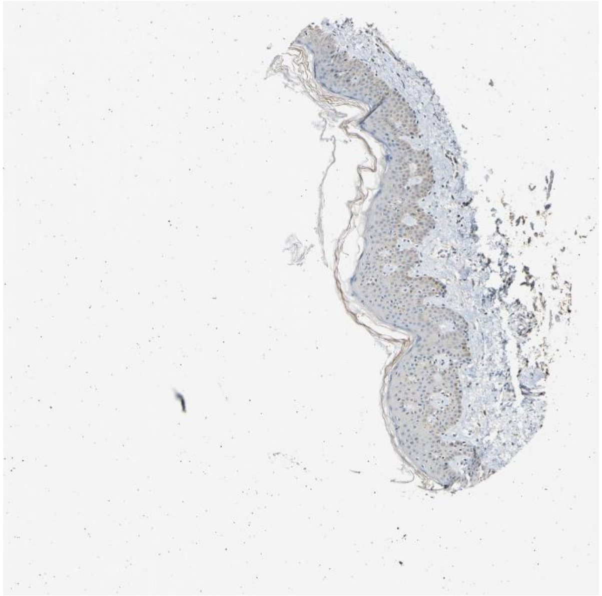

Calm1 Insr

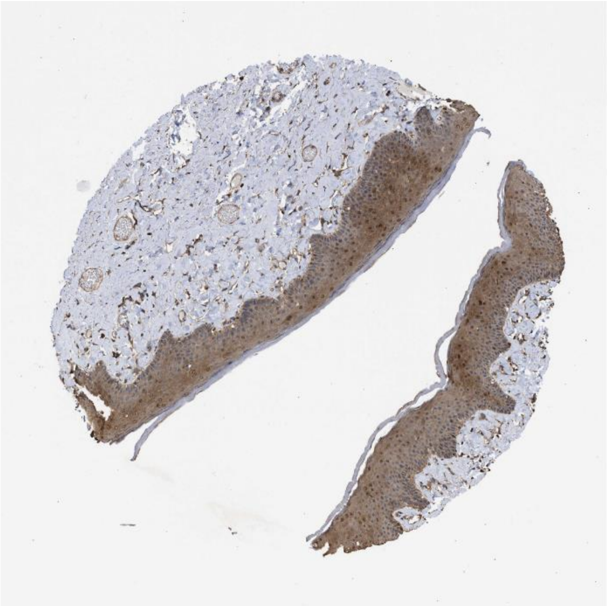

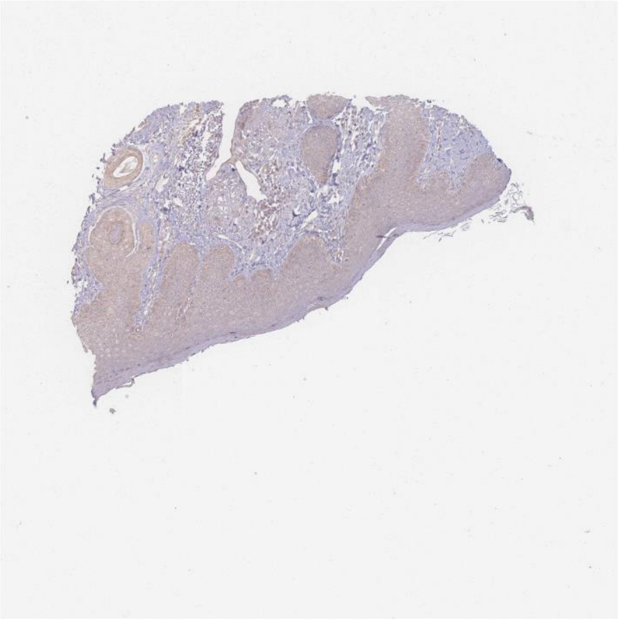

Anxa1 Egfr

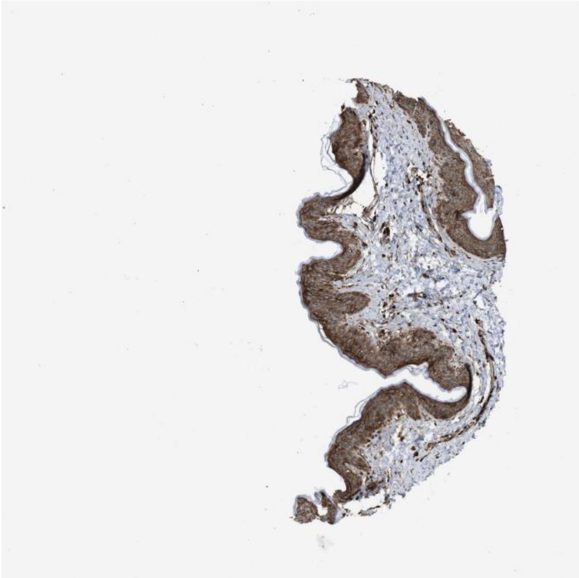

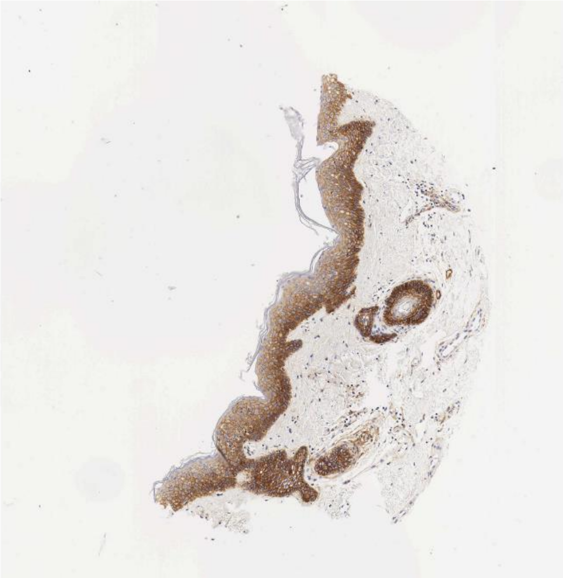

Ptn Sdc1

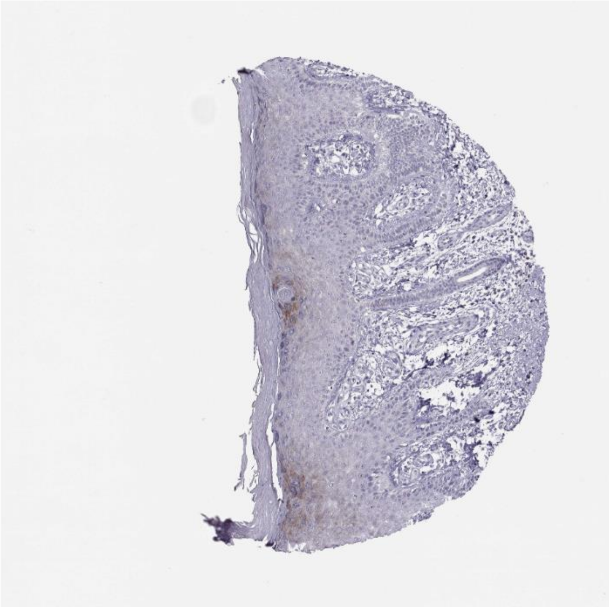

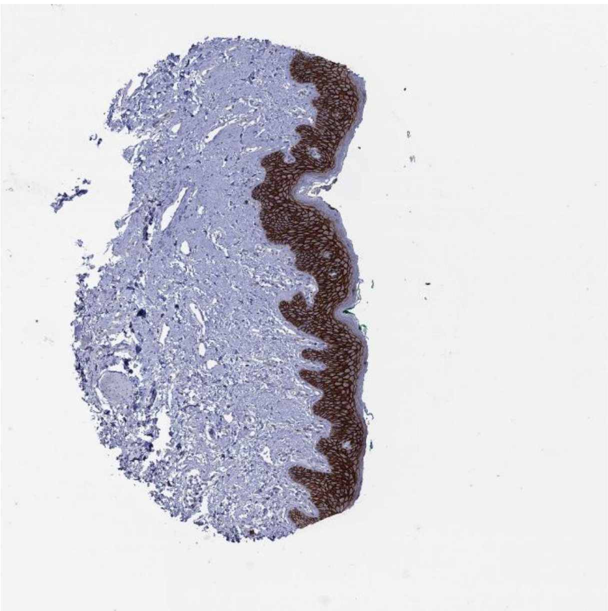

Rps27a Erbb2

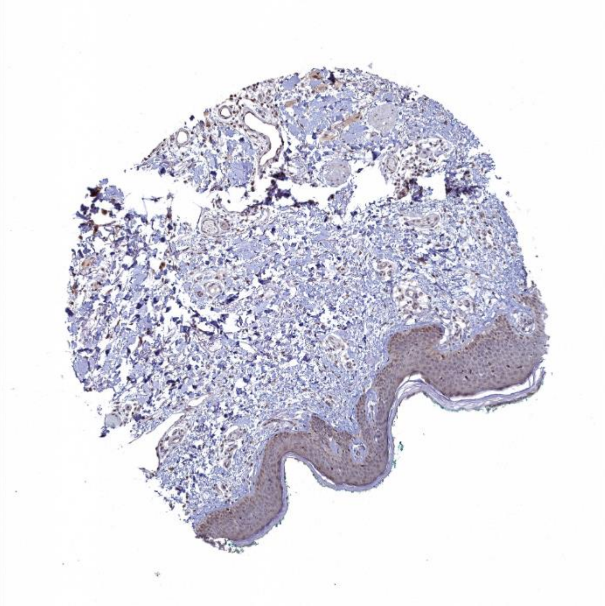

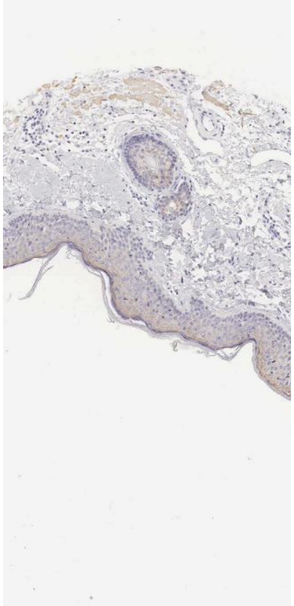

Cgn F11r

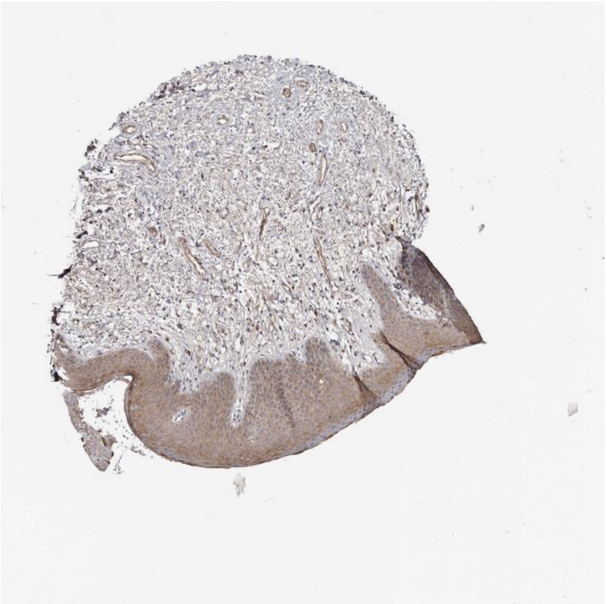

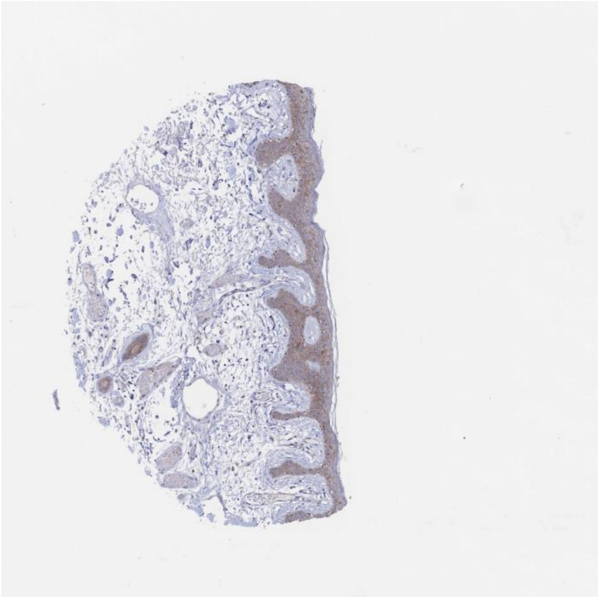

Hspa8 Ldlr

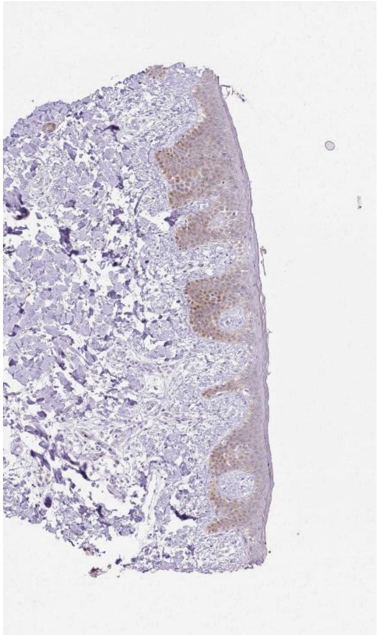

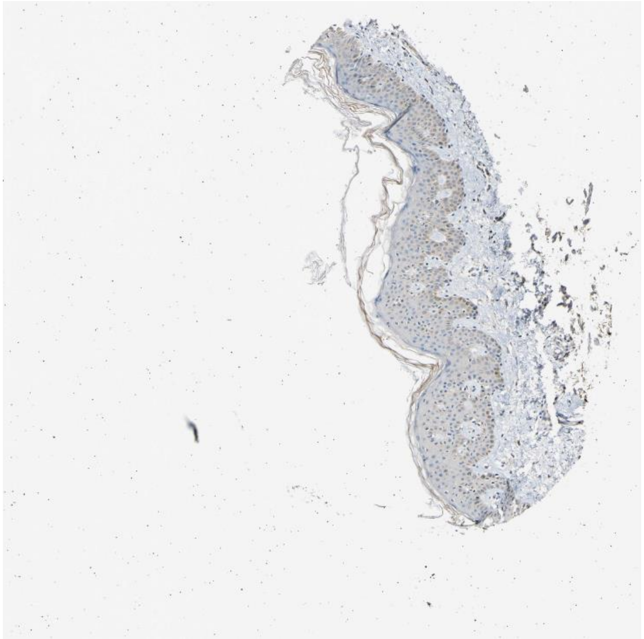

Calm2 Insr

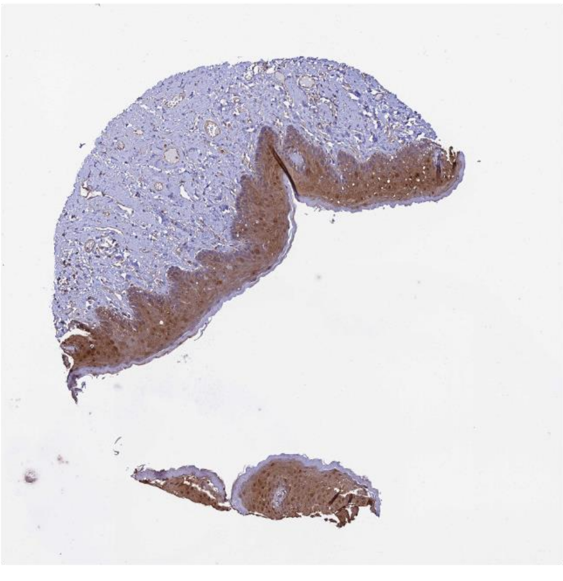

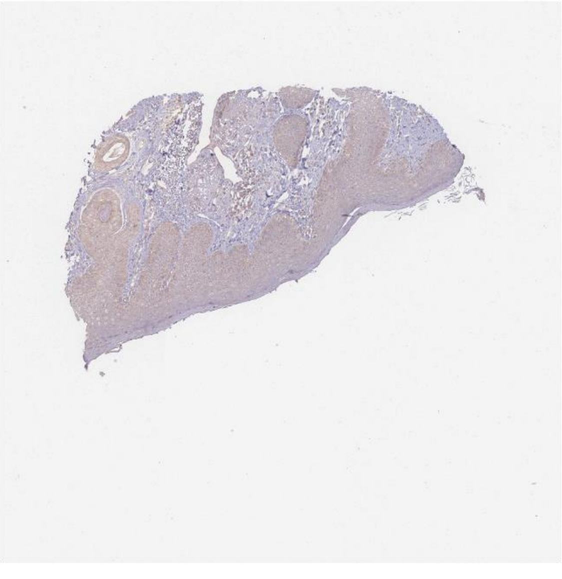

Efnb2 Epha4

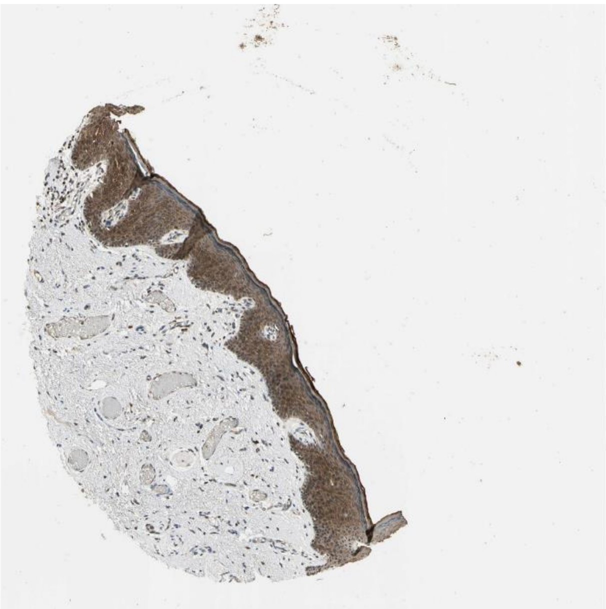

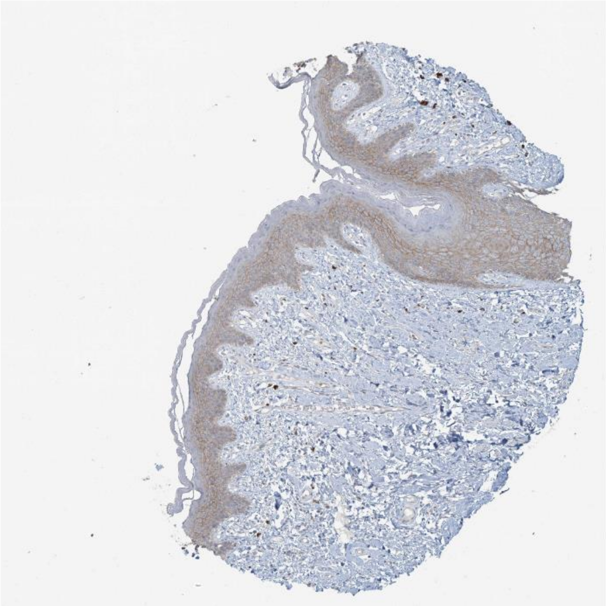

Ubc Fgfr2

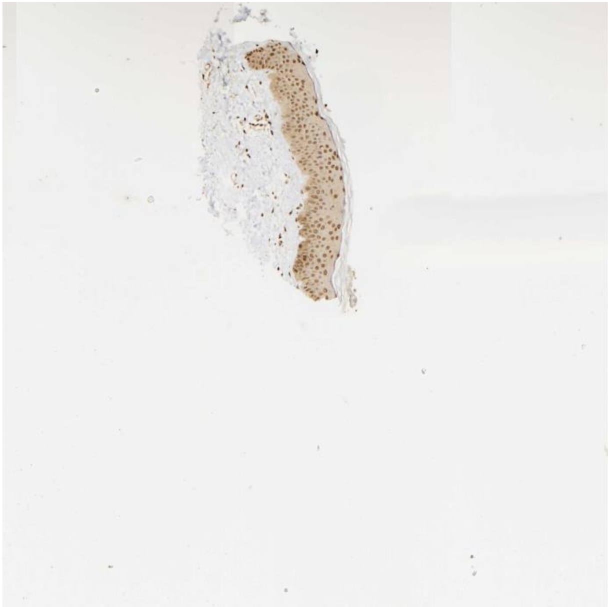

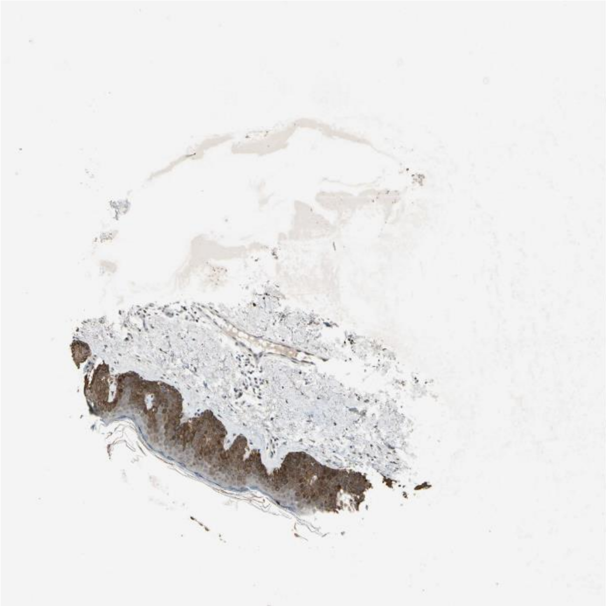

Hsp90b1 Erbb2

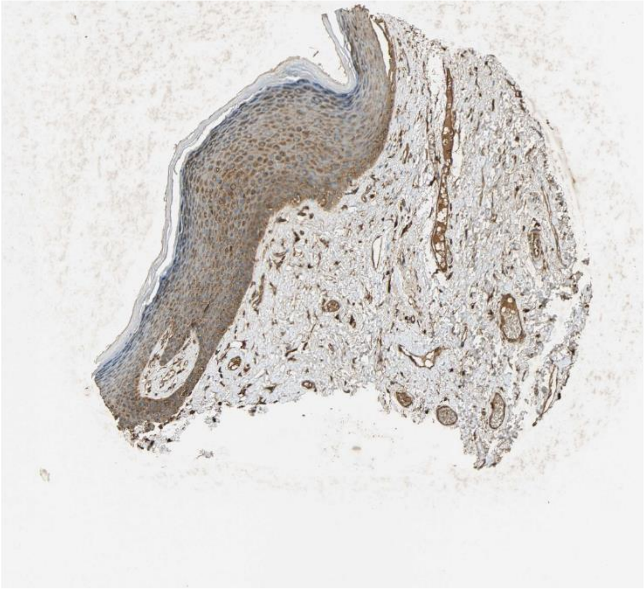

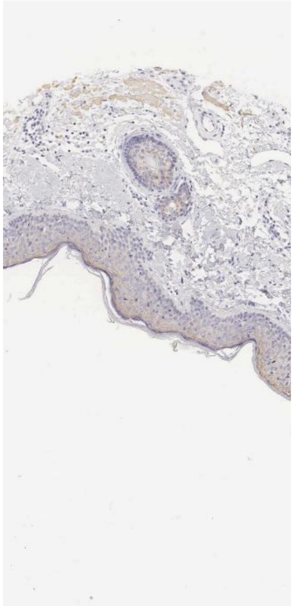

Fabp5 Rxra

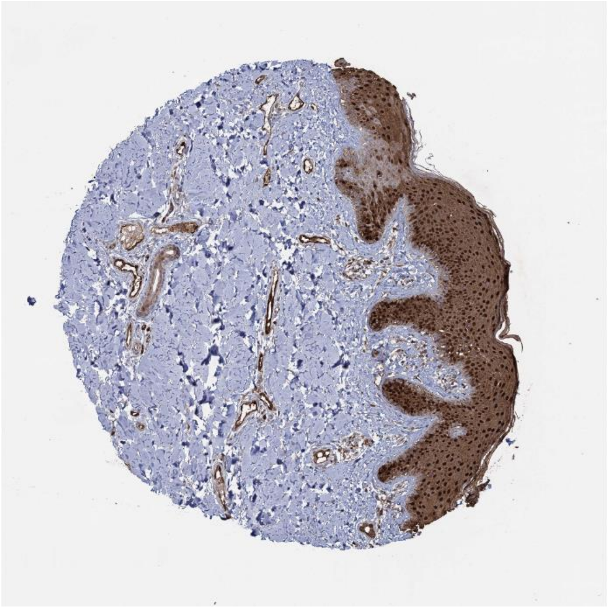

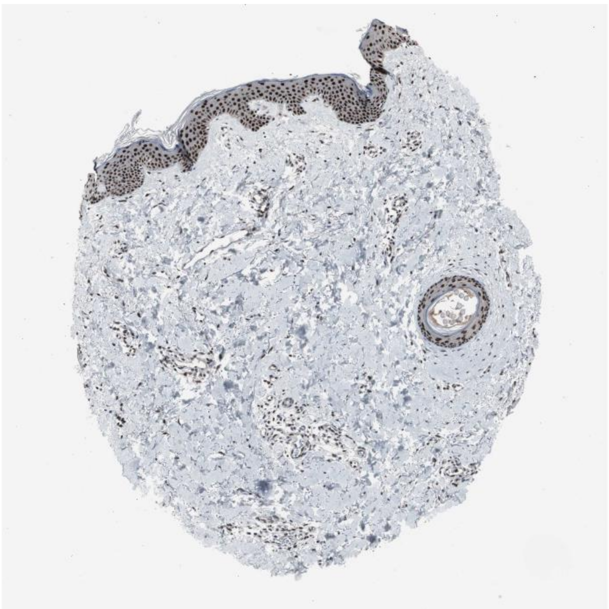

Ubb Ldlr

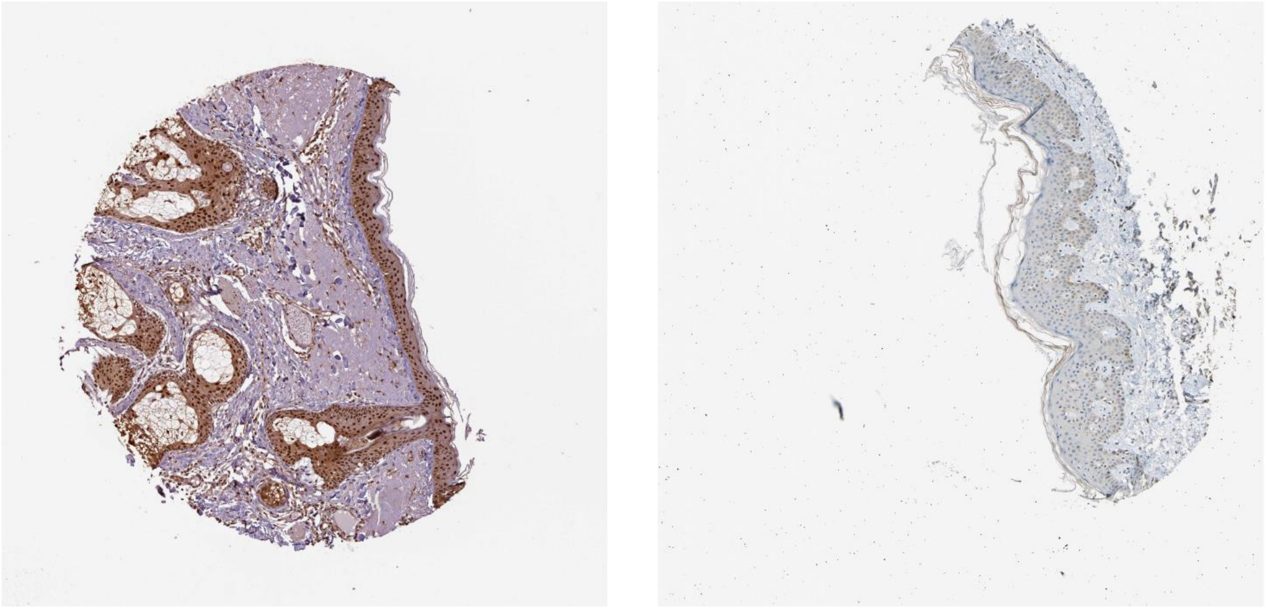

Actr2 Ldlr

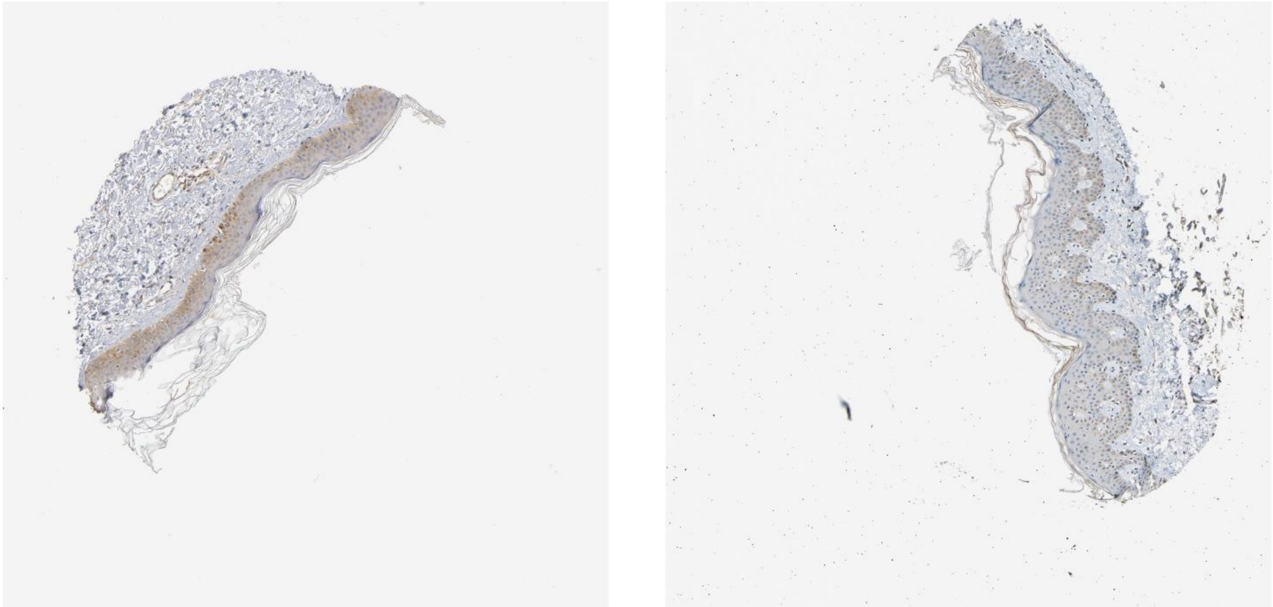

Hbegf **Cd44**

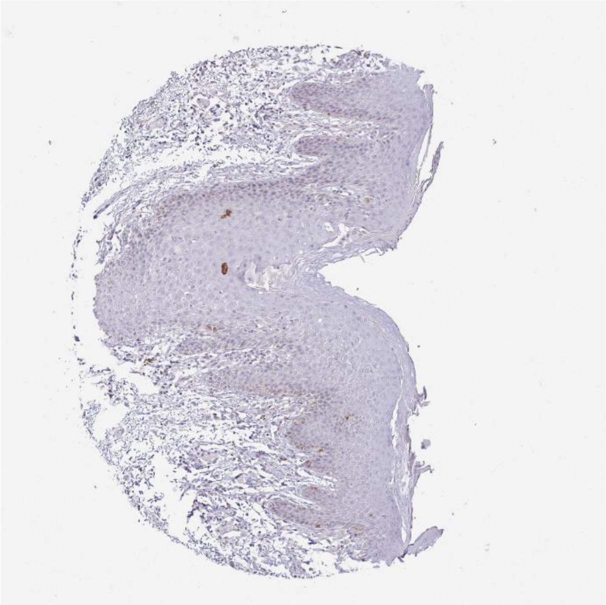

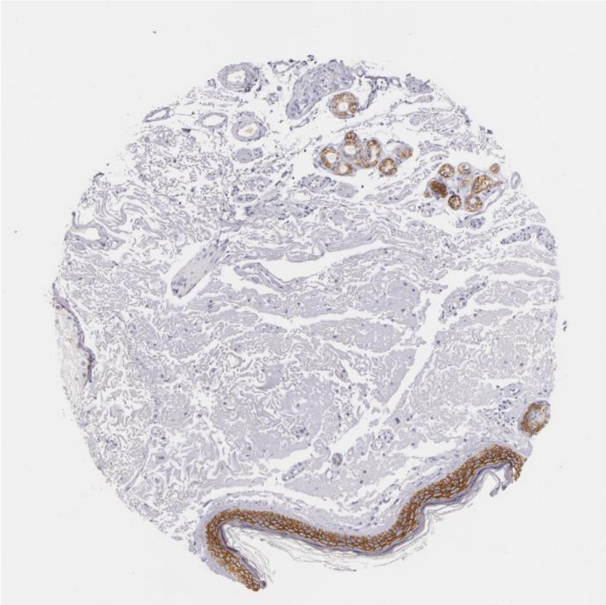

Hbegf Cd82

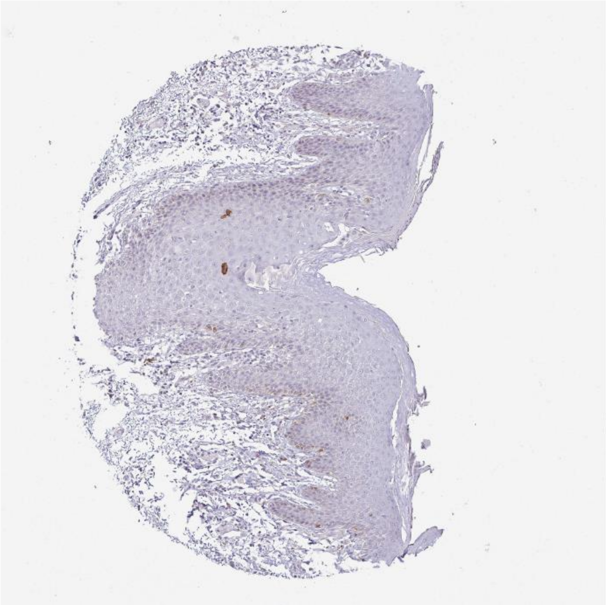

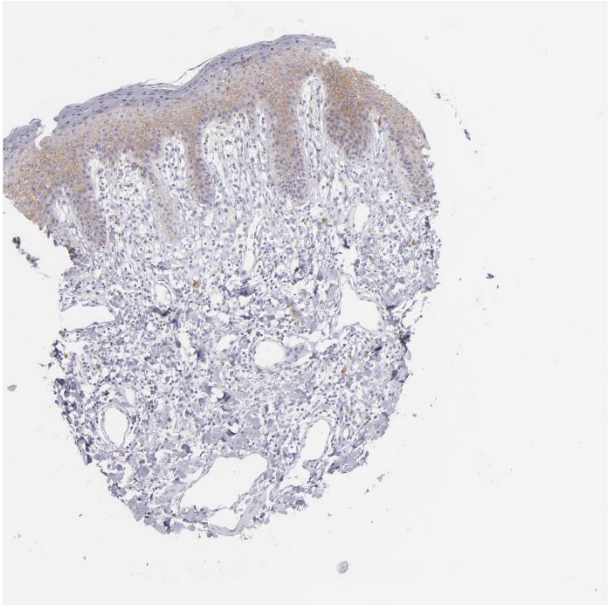

Rps27a Fgfr2

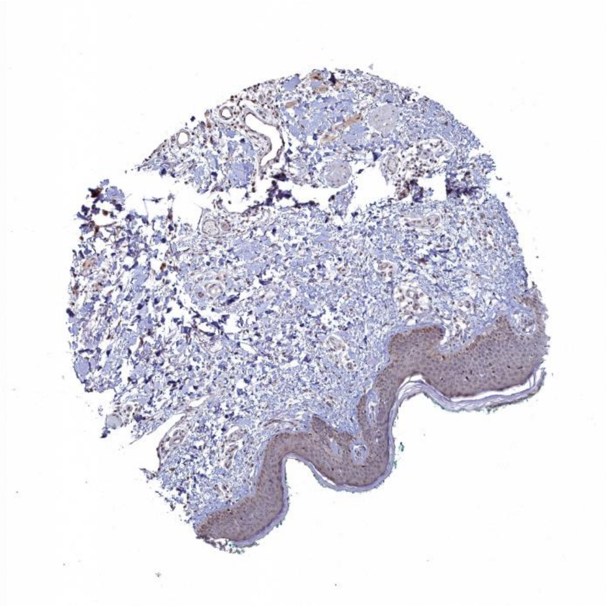

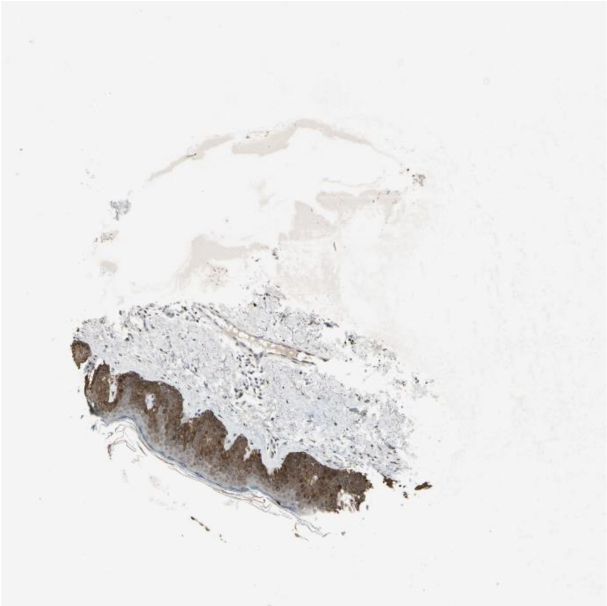

Ubb Erbb2

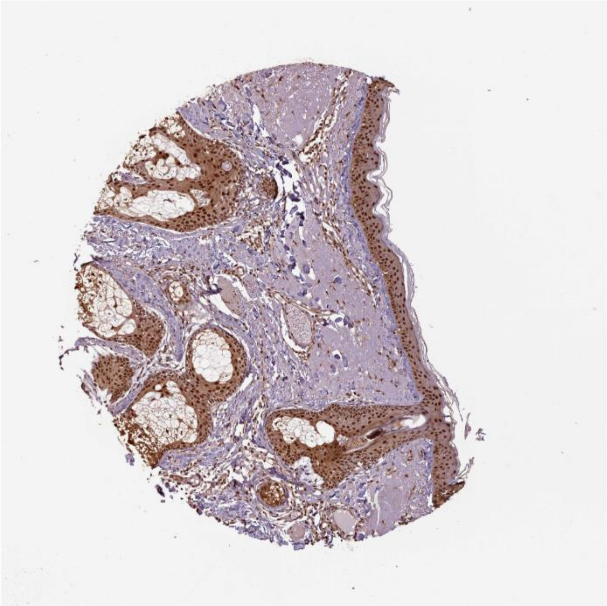

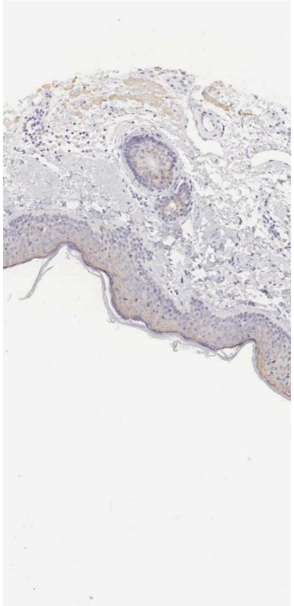

**Psen1** Notch1

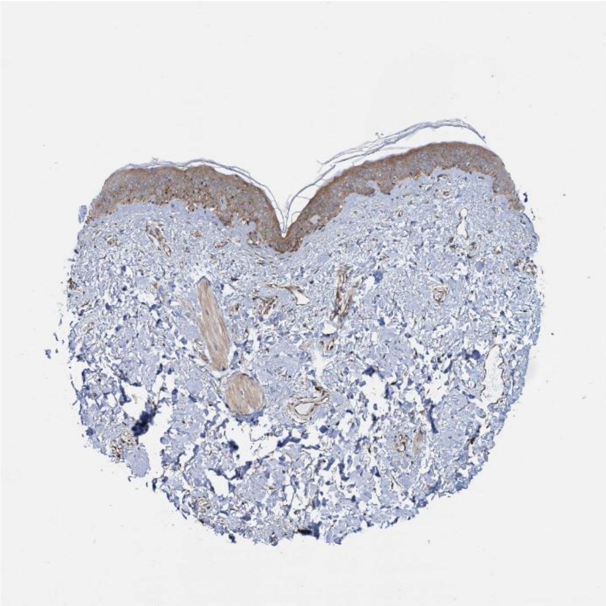

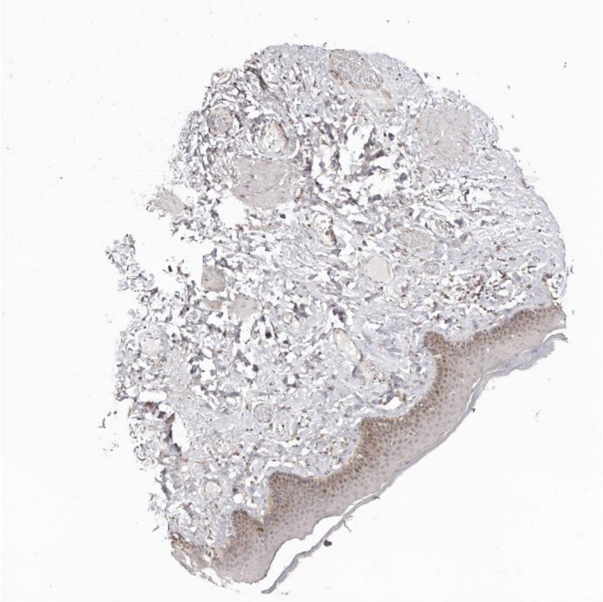

Dusp18 Itga6

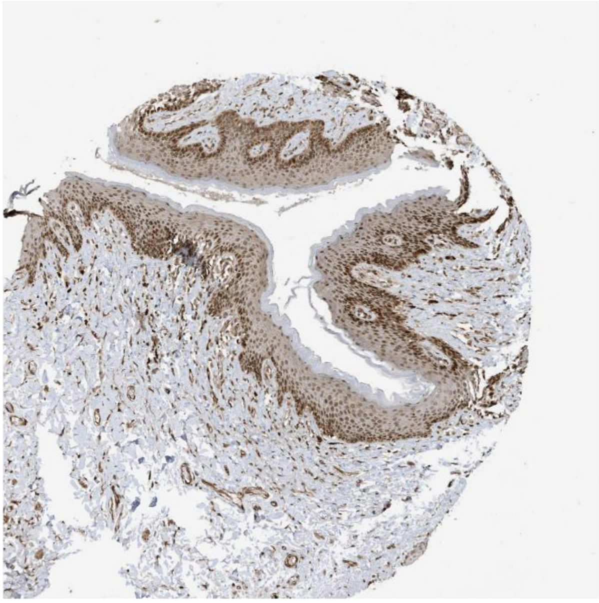

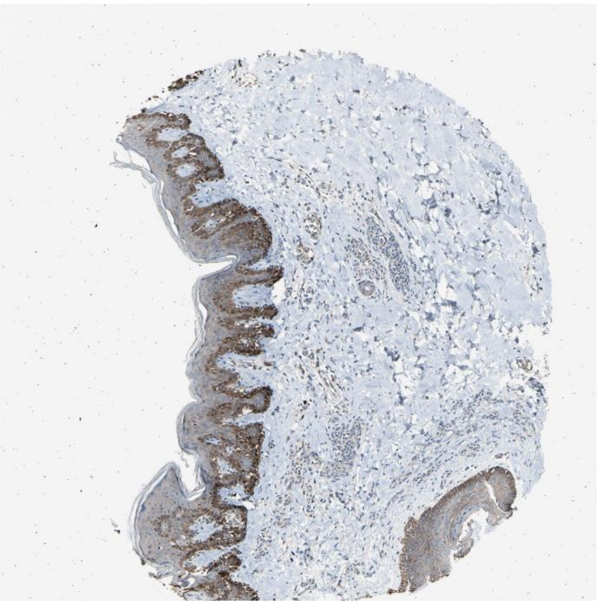

#### Hbegf Erbb2

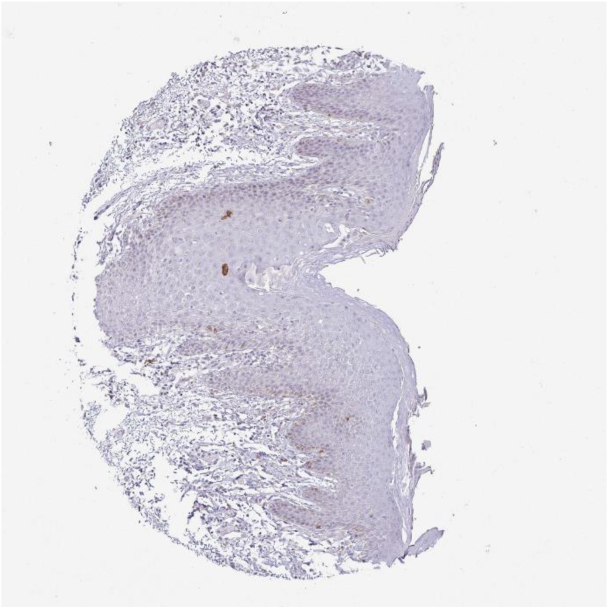

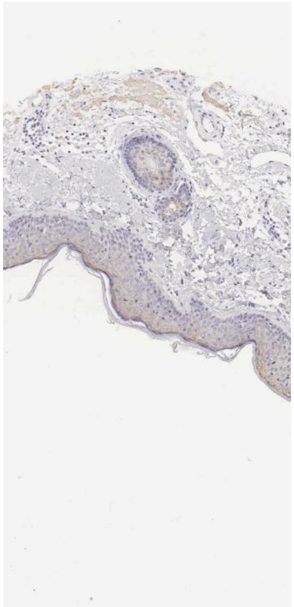

#### Cgn Ocln

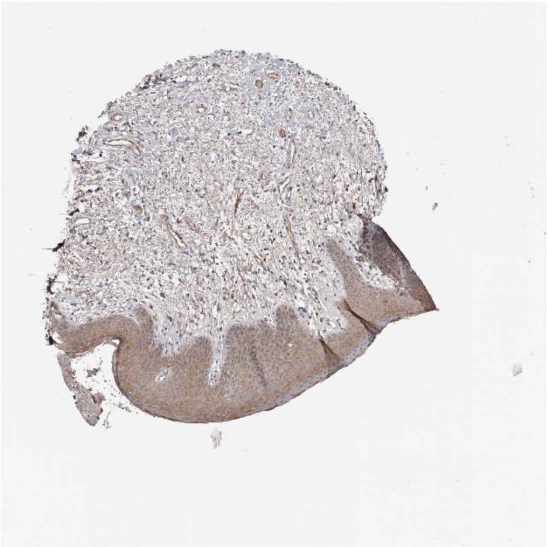

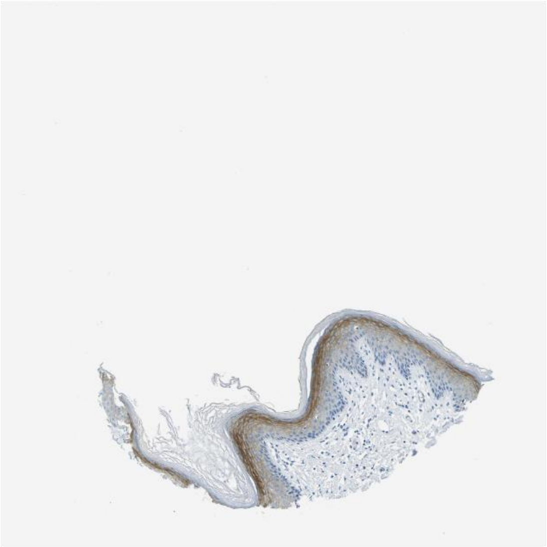

Efna1 Epha4

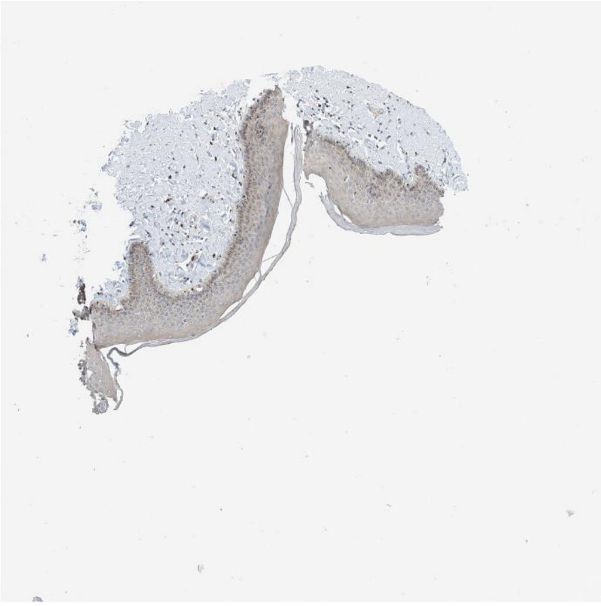

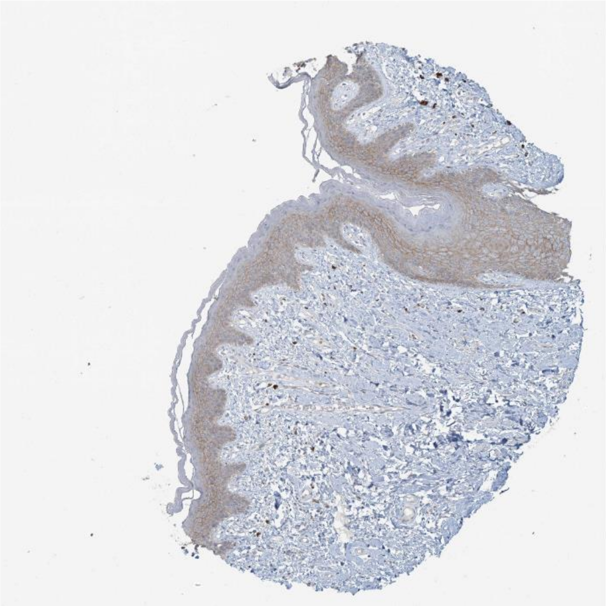

Calm1 Abca1

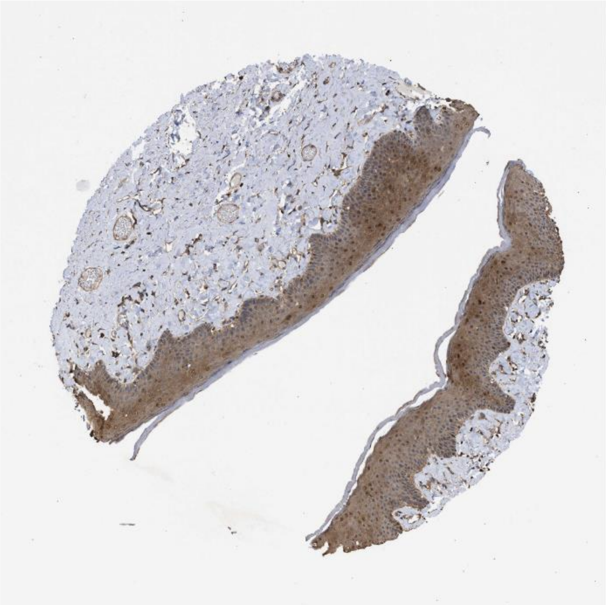

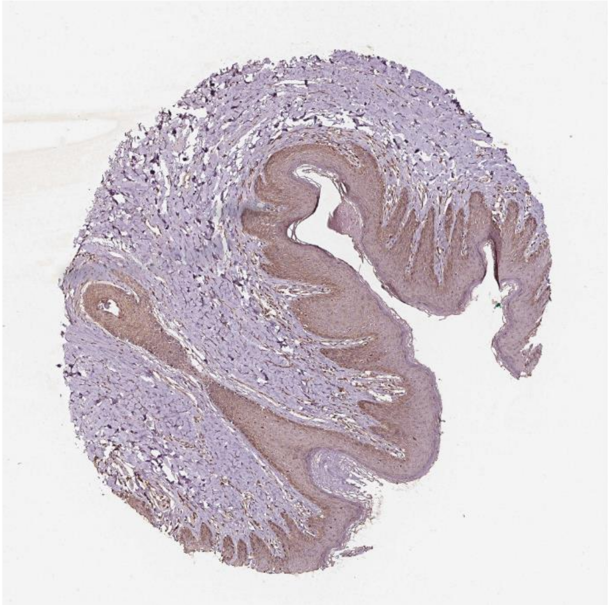

Cdh1 Egfr

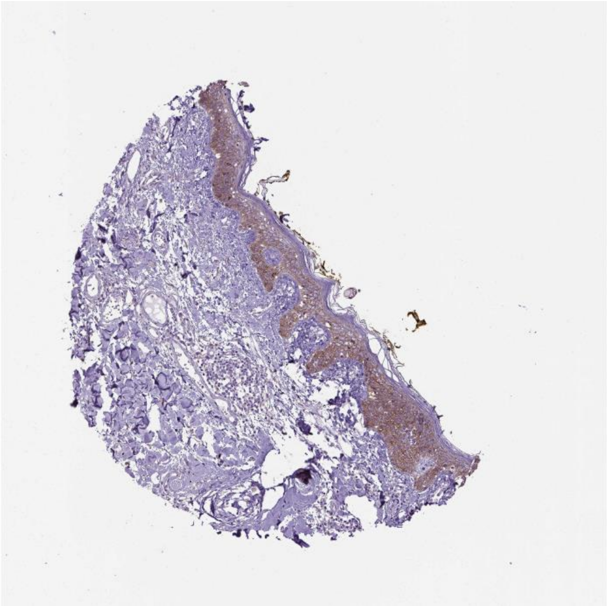 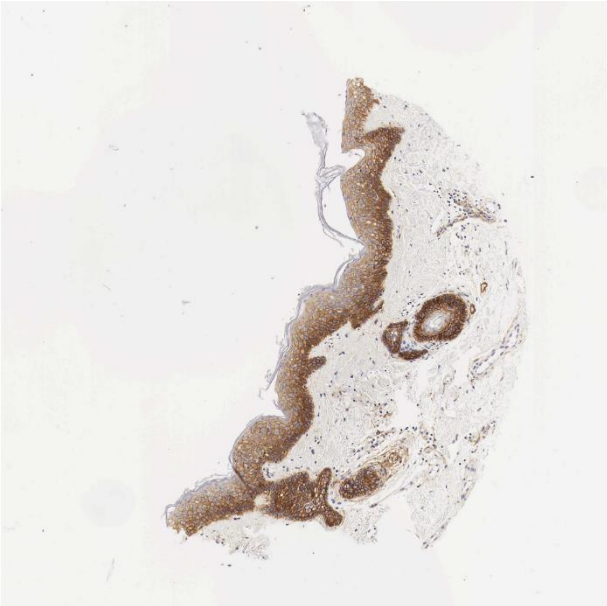

Psen1 Cd44

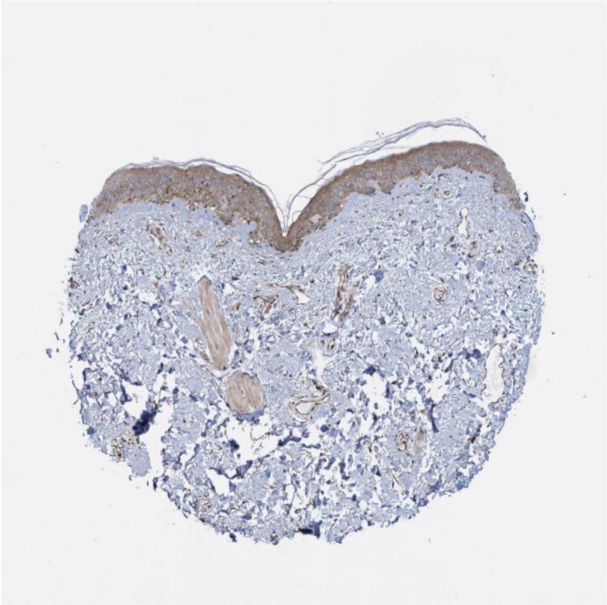 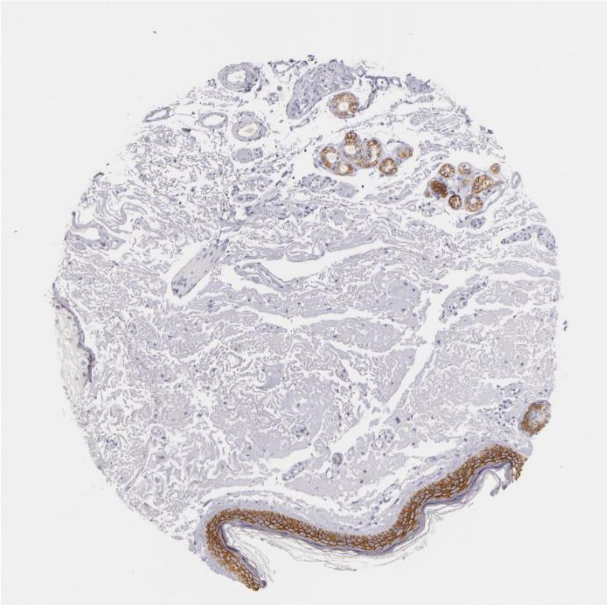

Adam17 Notch1

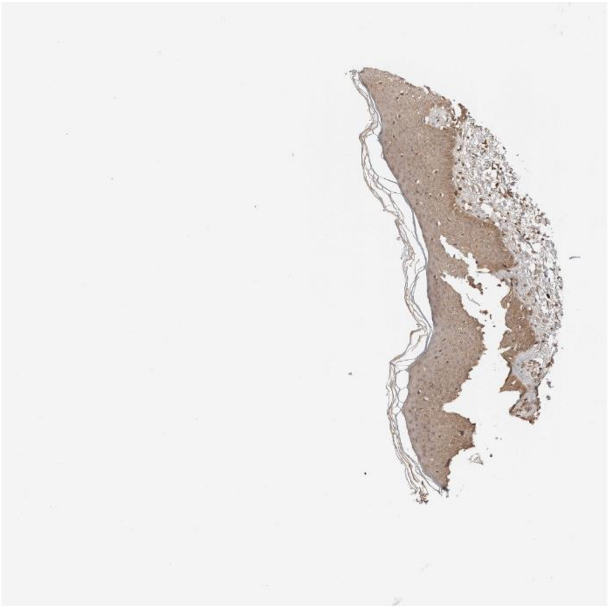

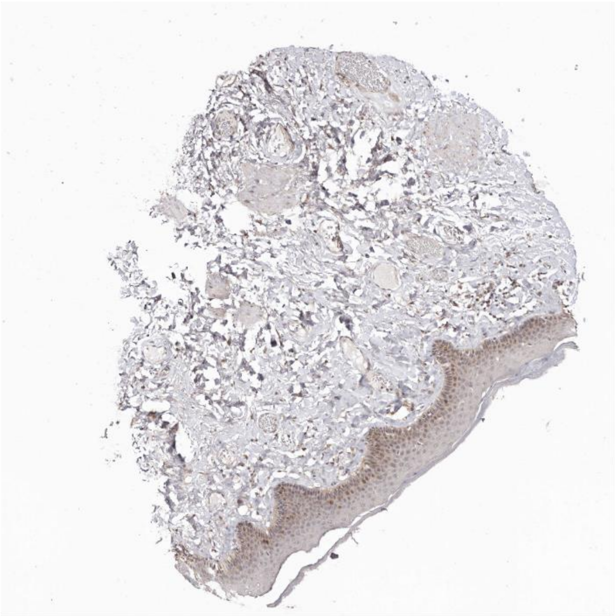

Apoe Lrp1

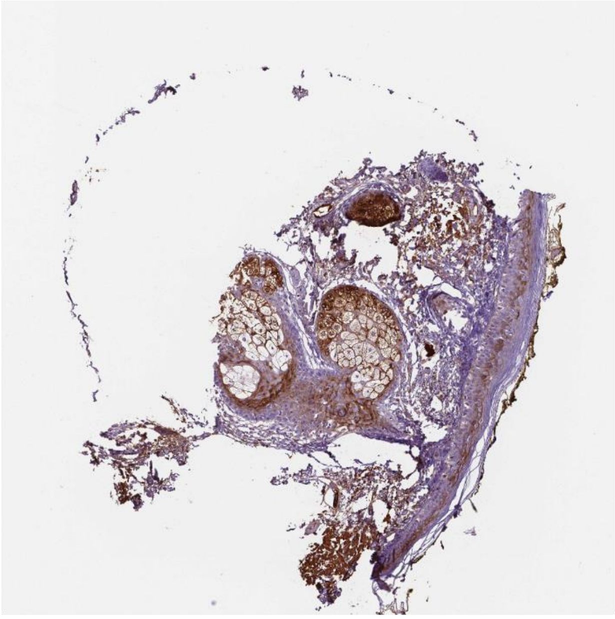

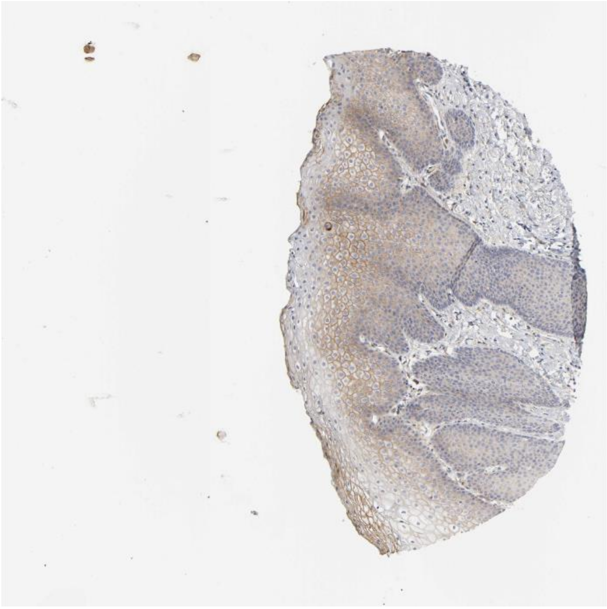

Calr Itgav

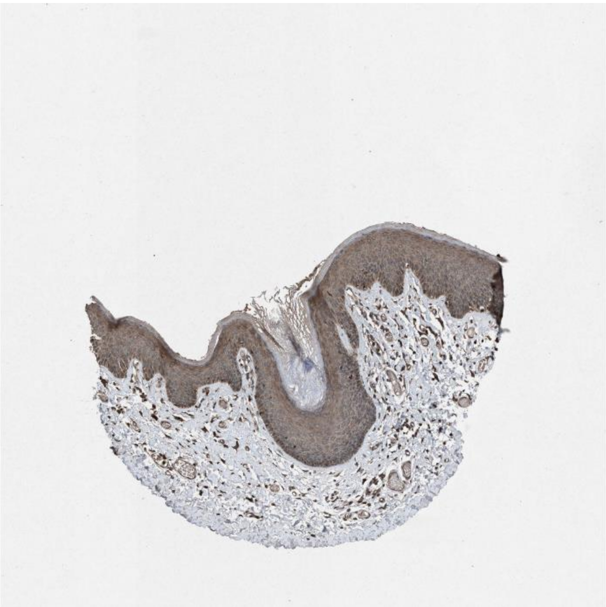

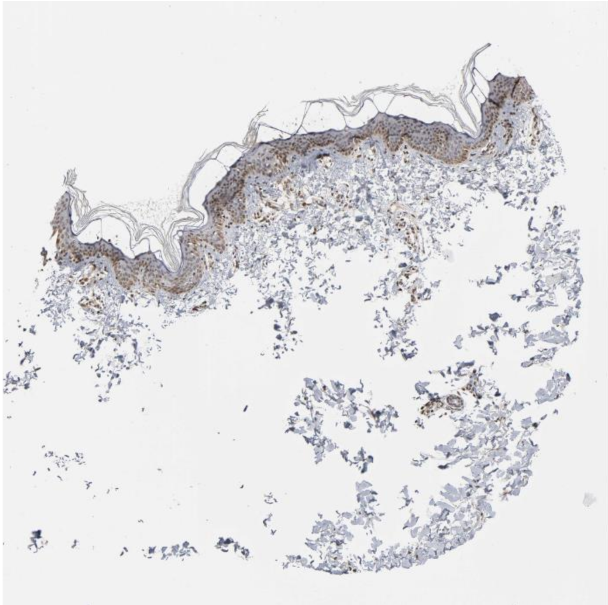

Cdh1 Igf1r

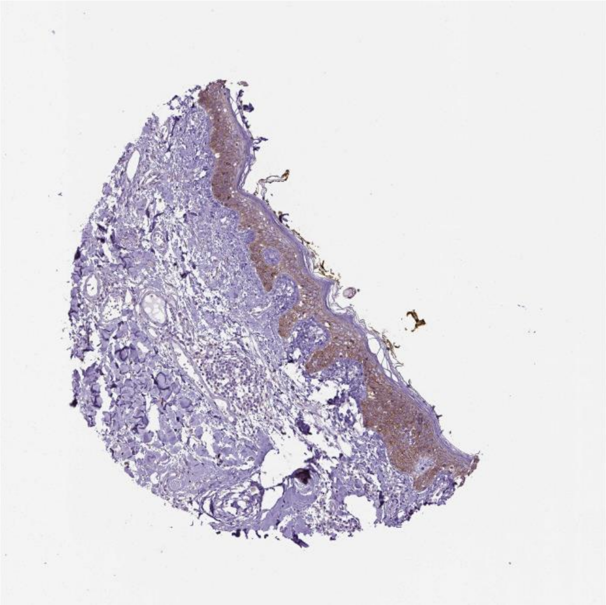

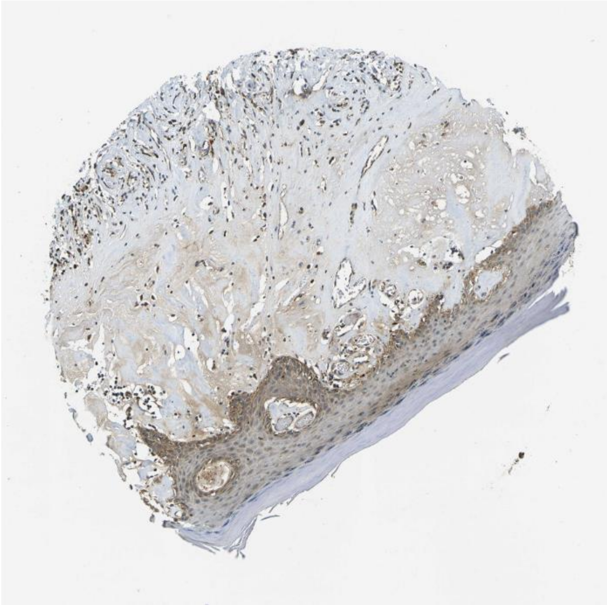

Sorbs1 Itgb5

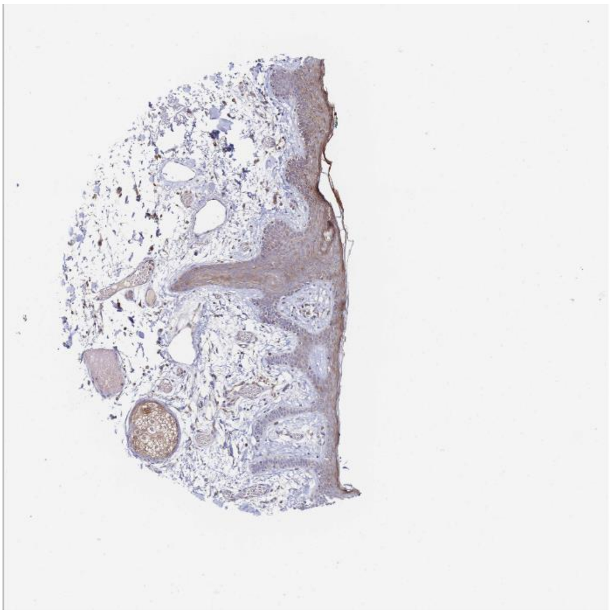

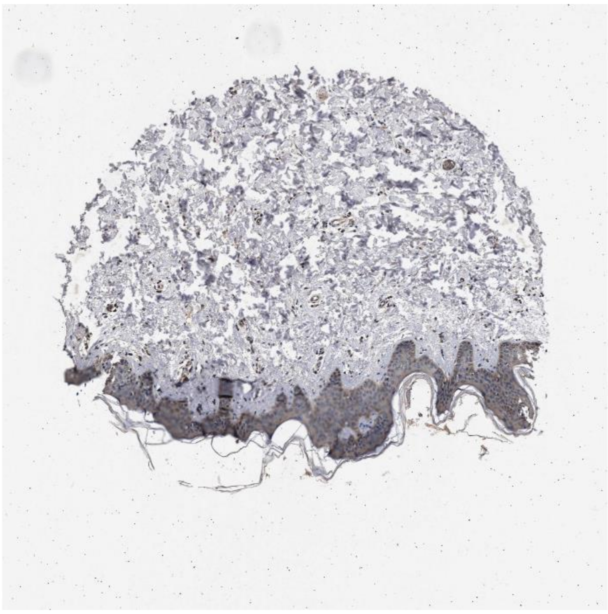

Ubb Fgfr2

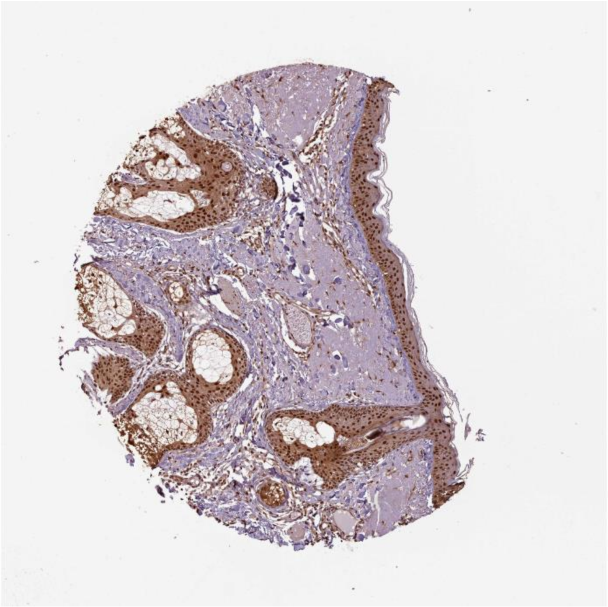

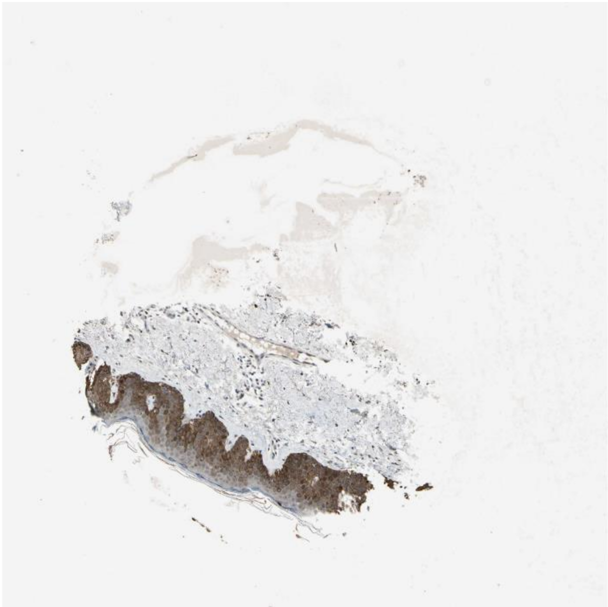

Calm2 Abca1

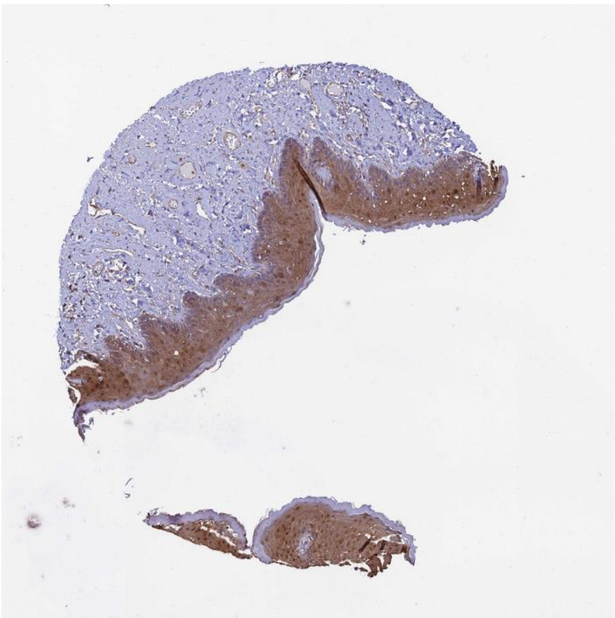 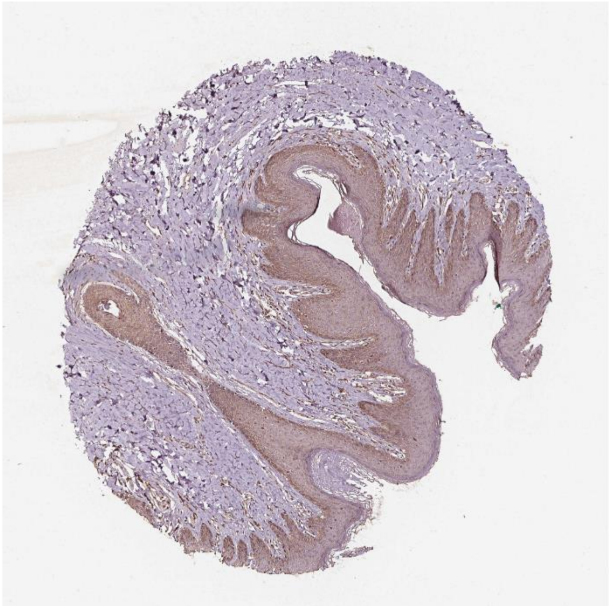

Pkm **Cd44**

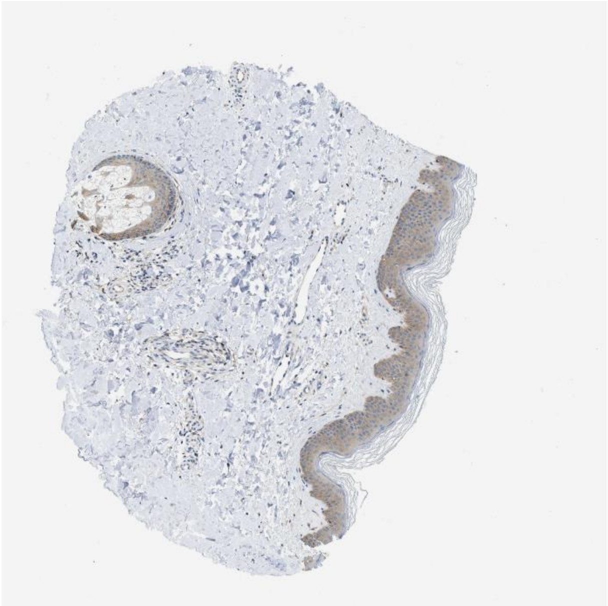 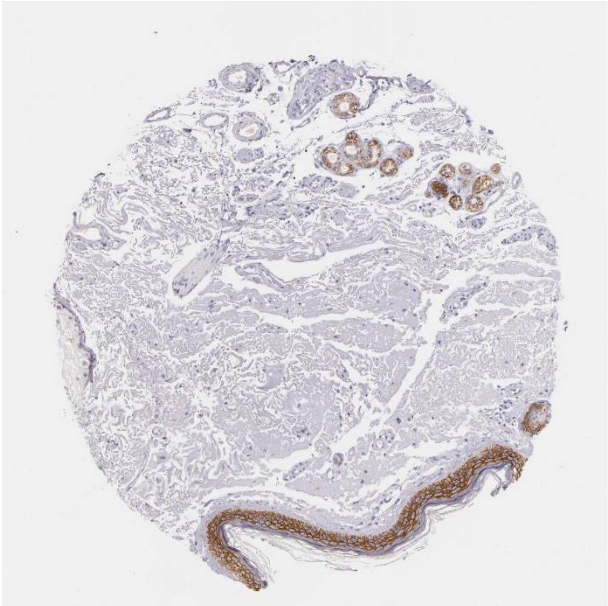

Hsp90b1 Lrp1

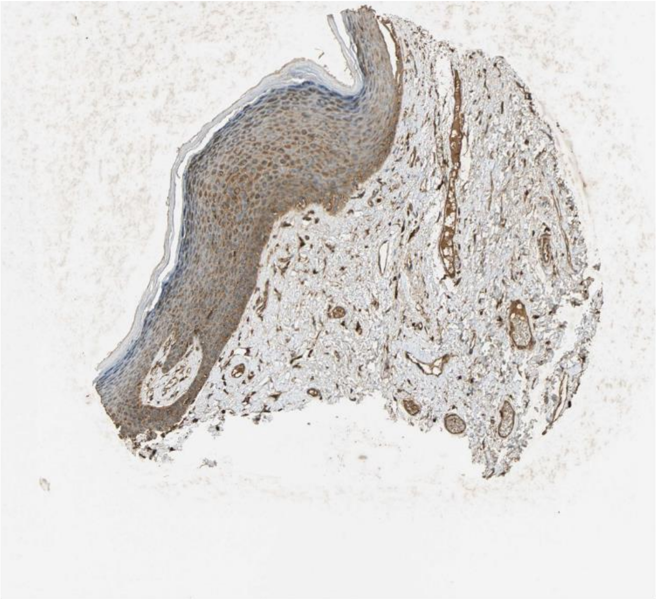

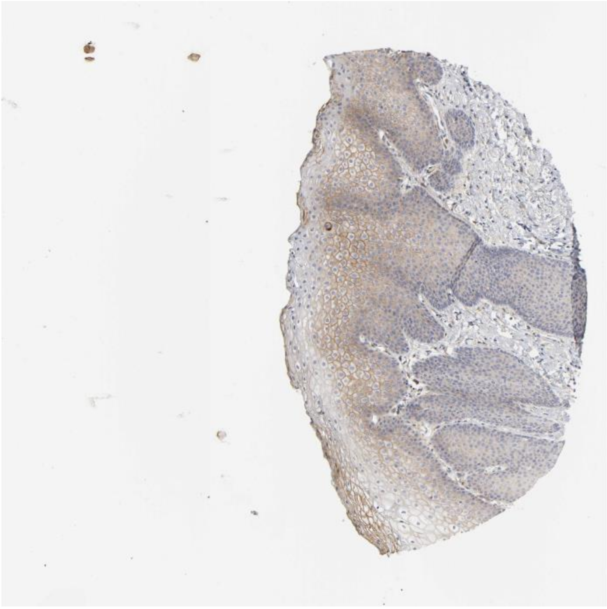

Tgfa Erbb2

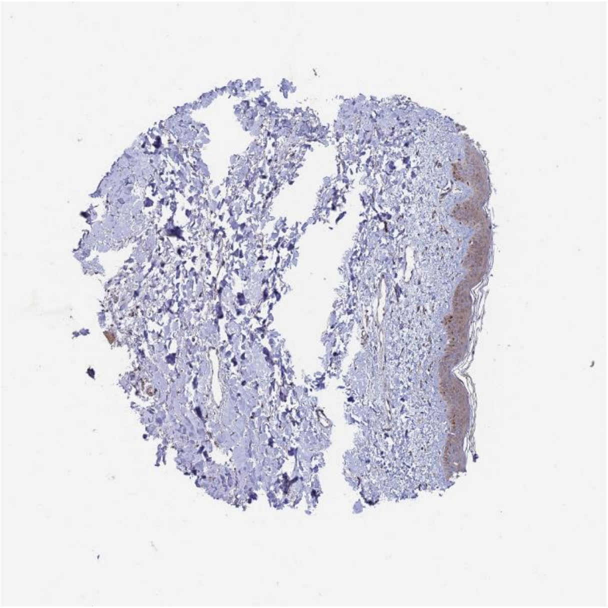

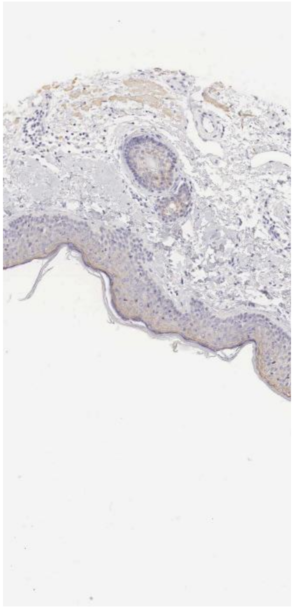

Sptan1 Ptpra

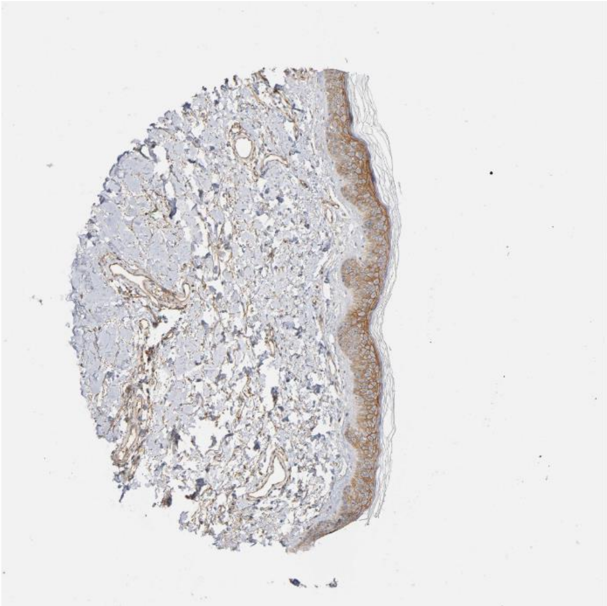

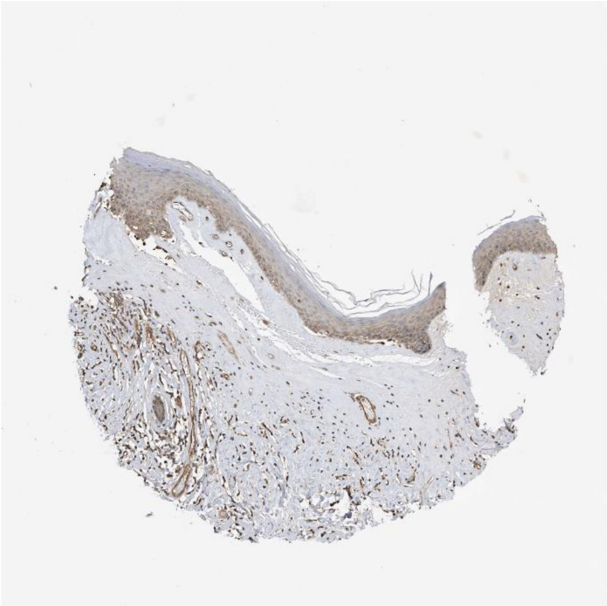

Arf1 Insr

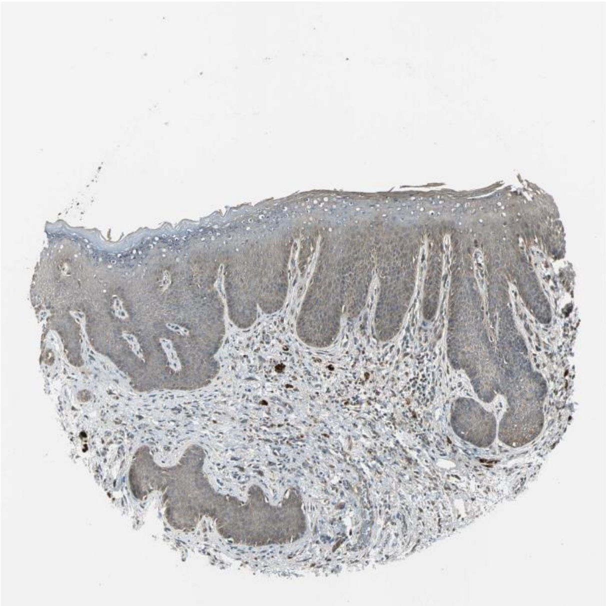

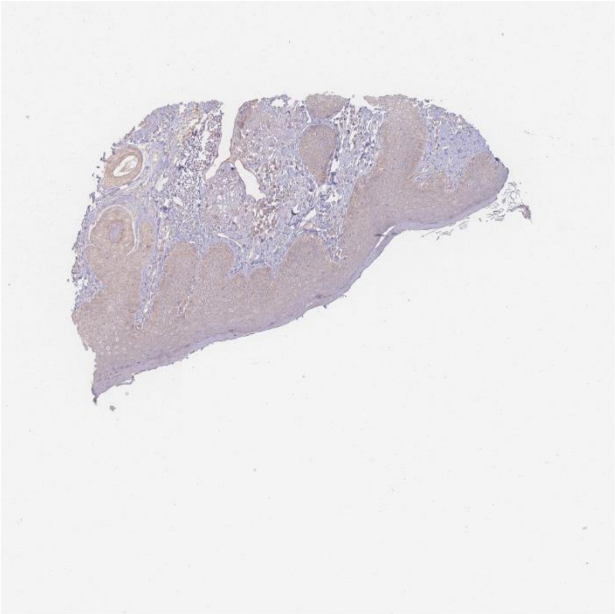

Sptbn2 Ptpra

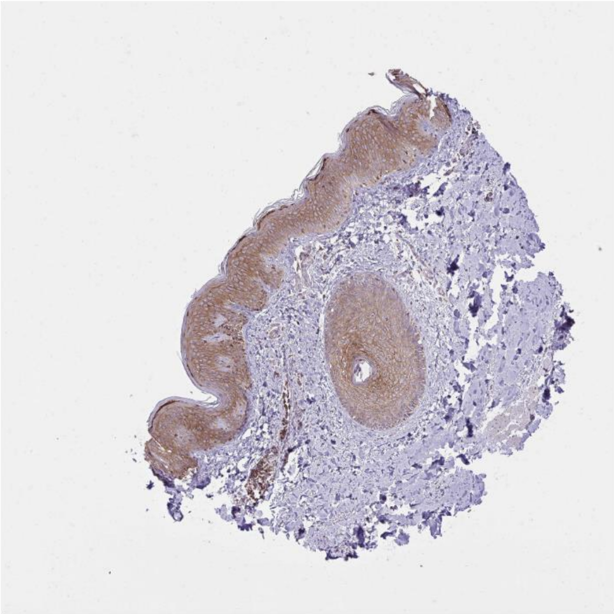

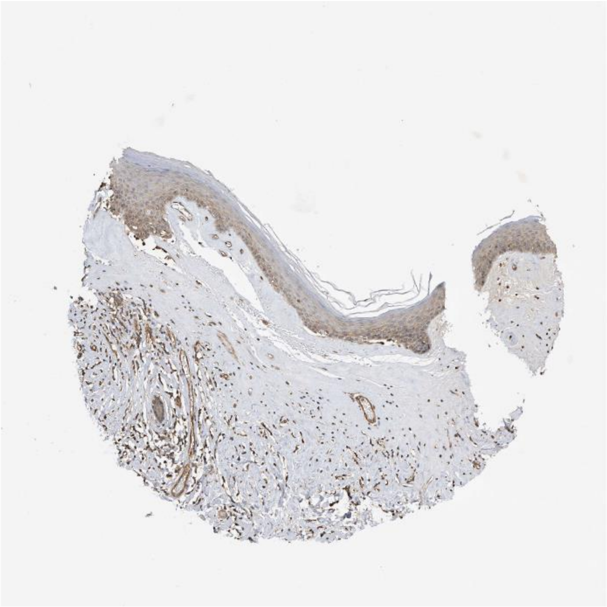

Lamb3 Col17a1

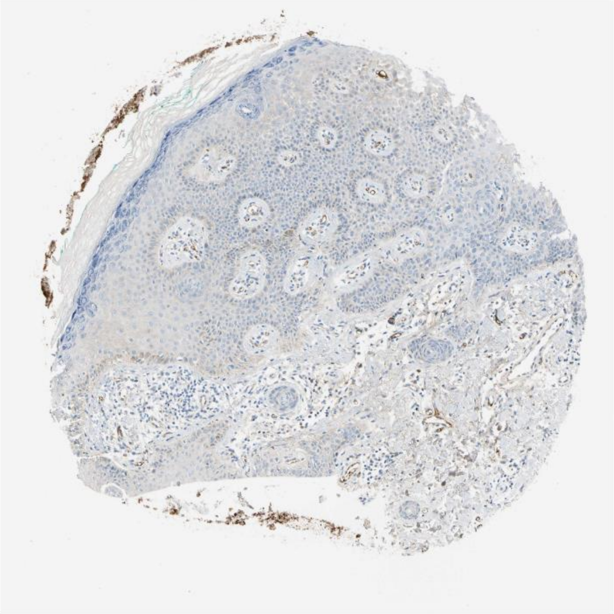

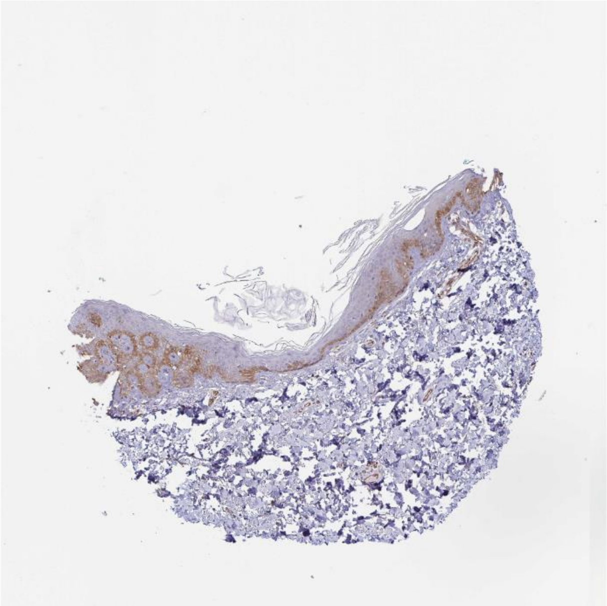

Hbegf Egfr

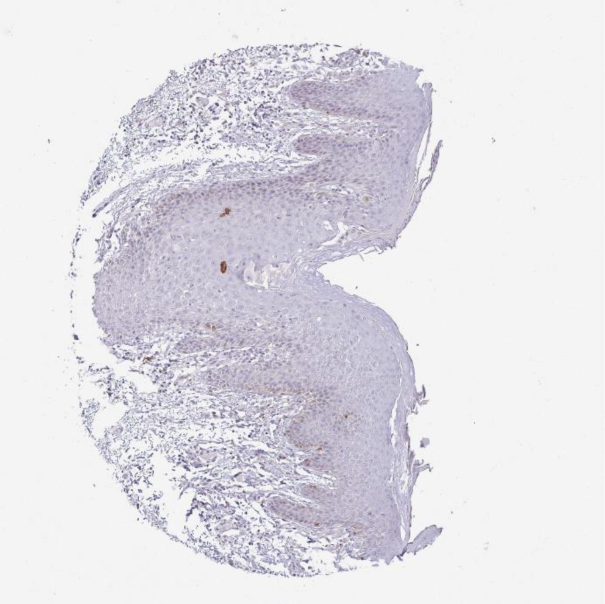

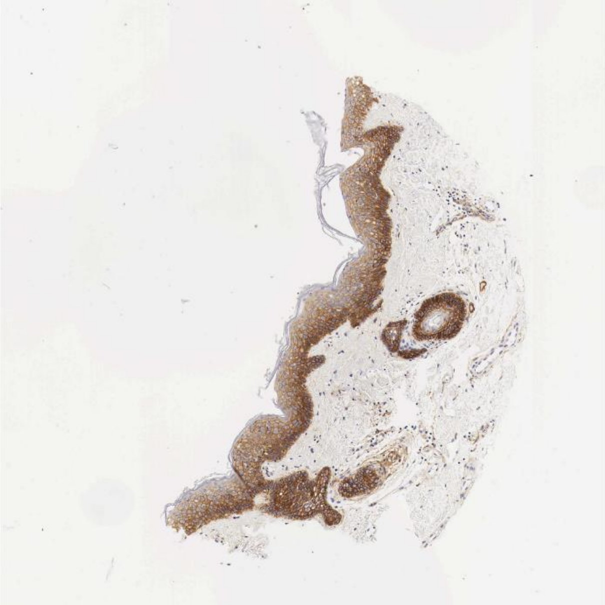

Ecm1 Cachd1

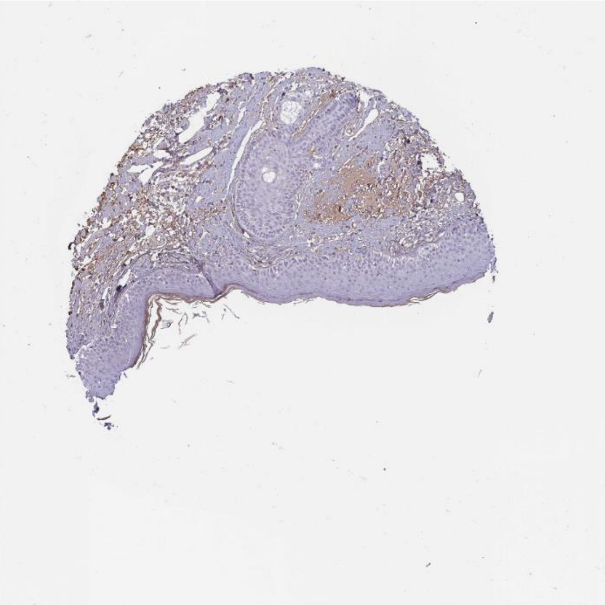

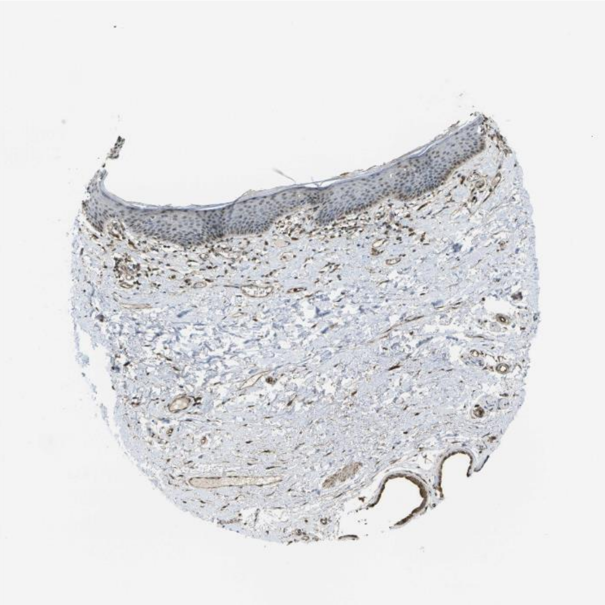

**Psen1** Notch2

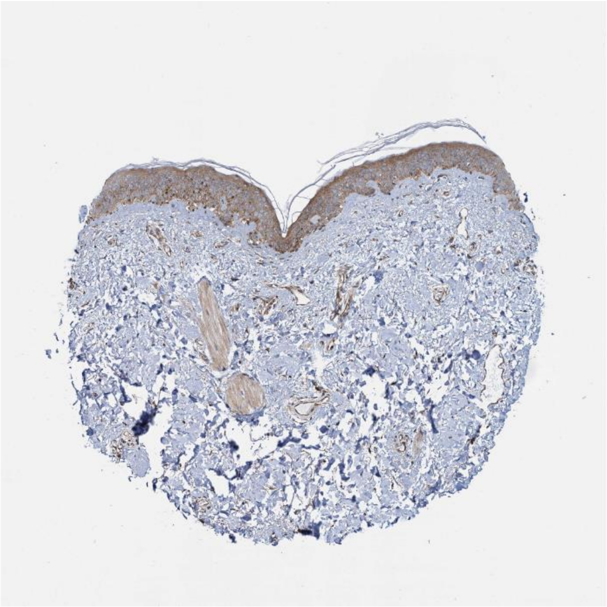

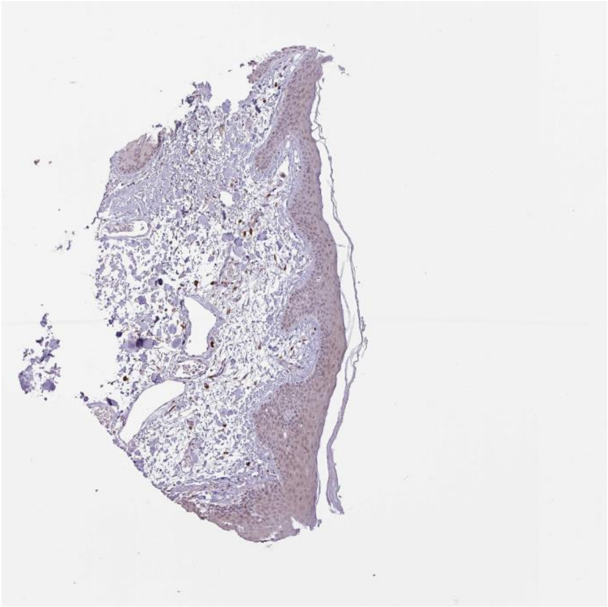

Hspg2 Sdc1

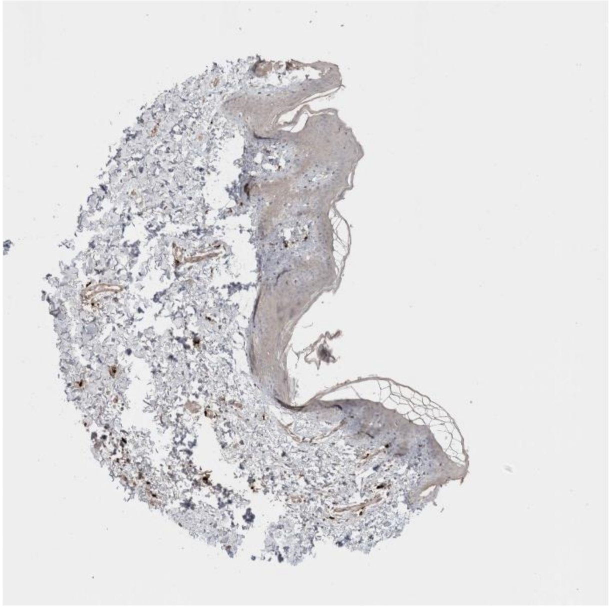

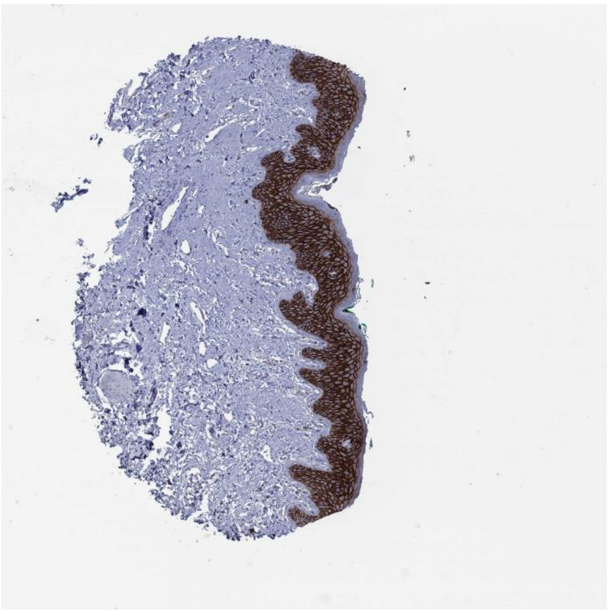

Ubc Ripk1

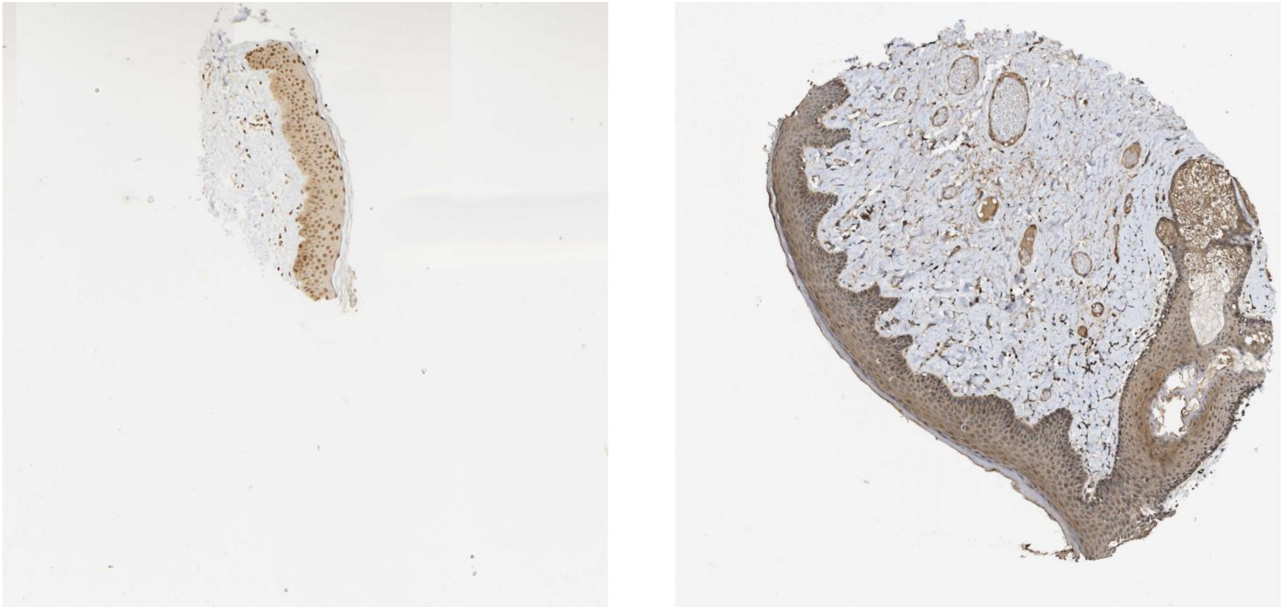

Adam10 Notch1

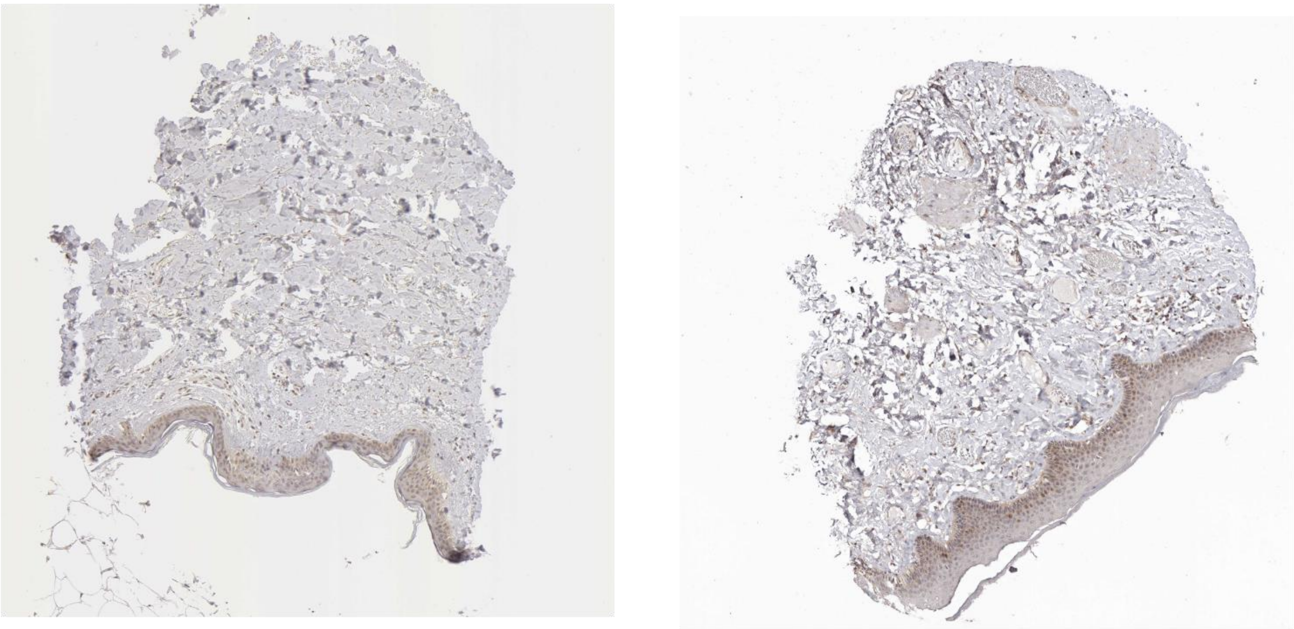

Calr Lrp1

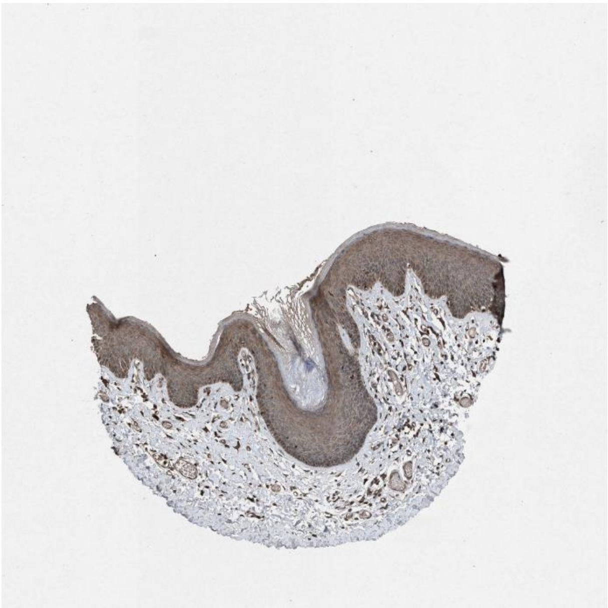

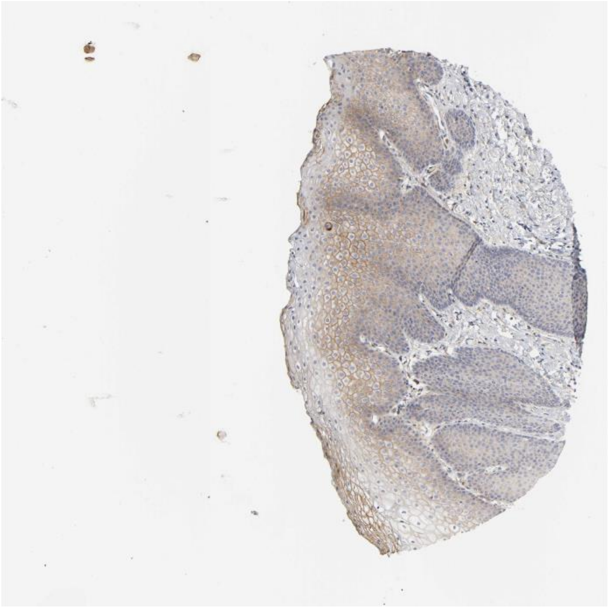

Gas6 Tyro3

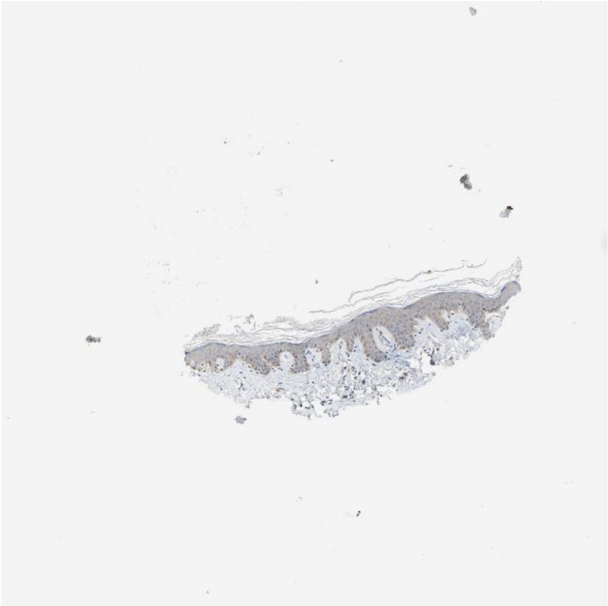

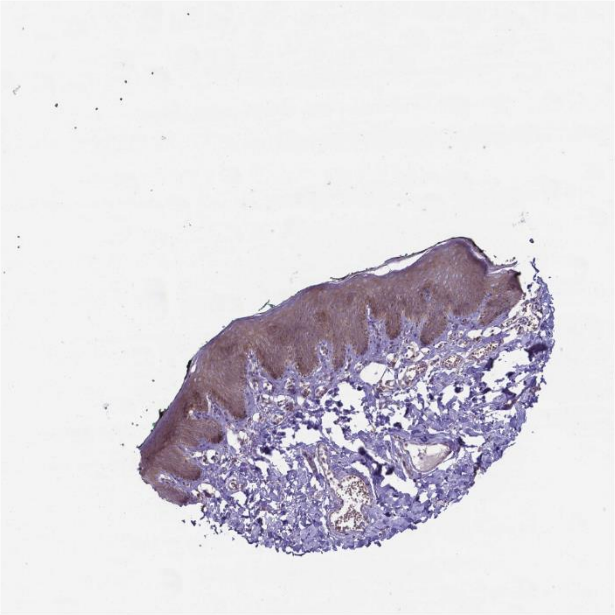

Dusp18 Cd151

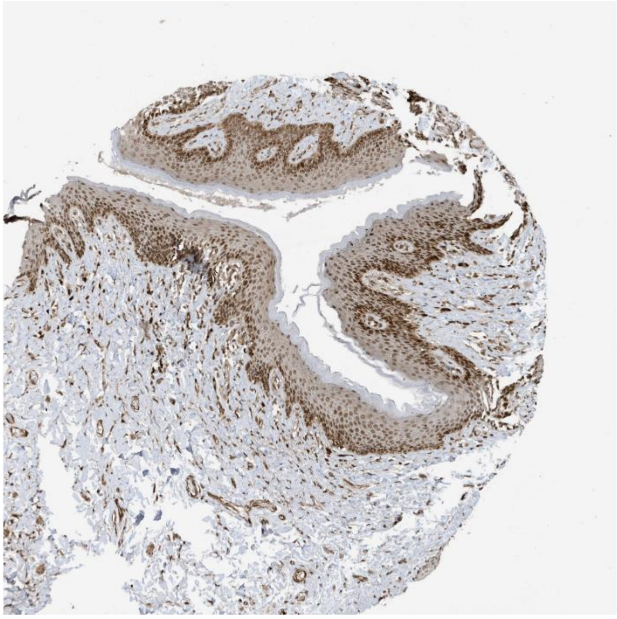
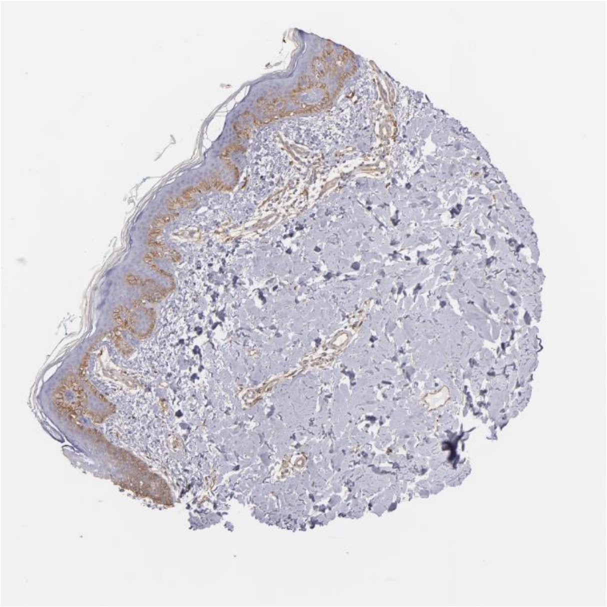

Dusp18 Itgb4

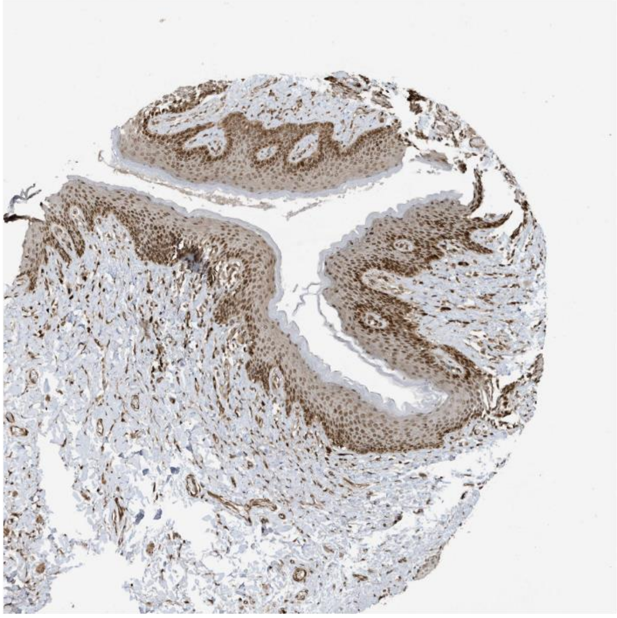
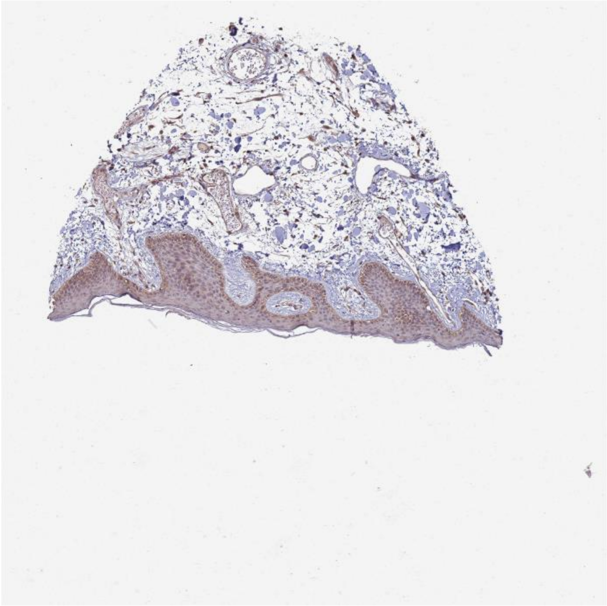

Efnb2 Ephb3

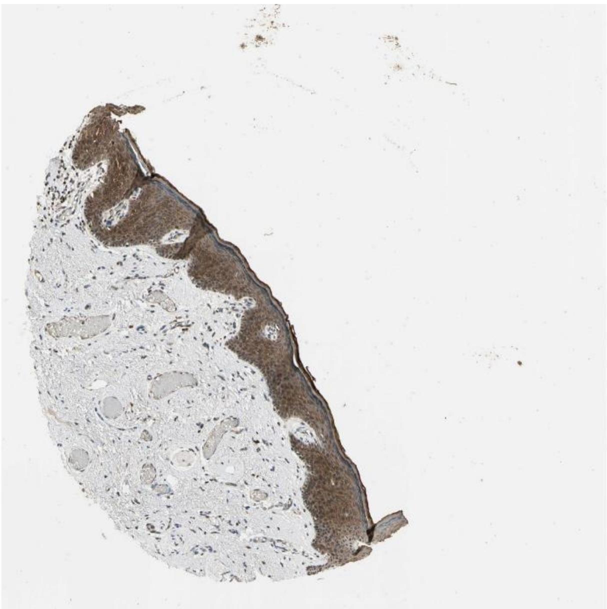

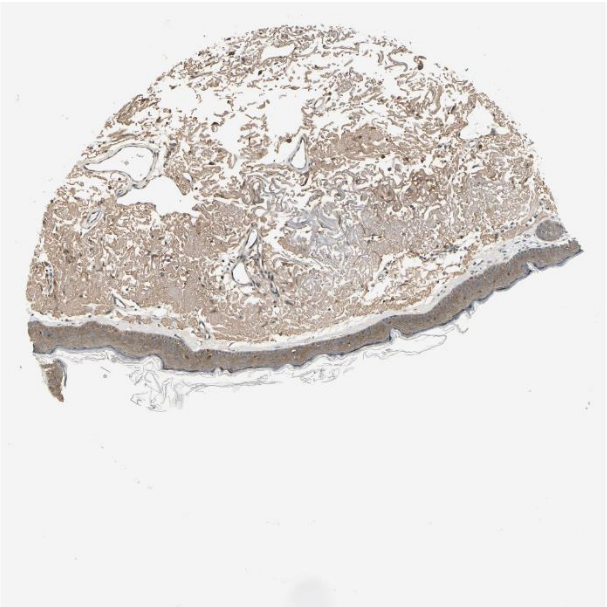

Sorbs1 Insr

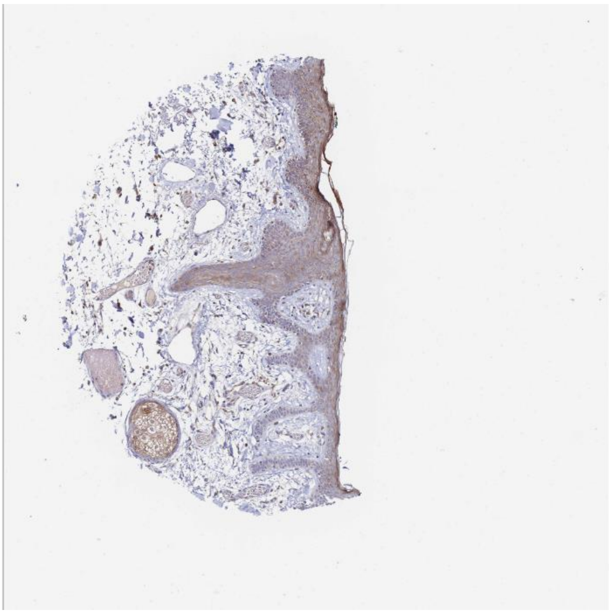

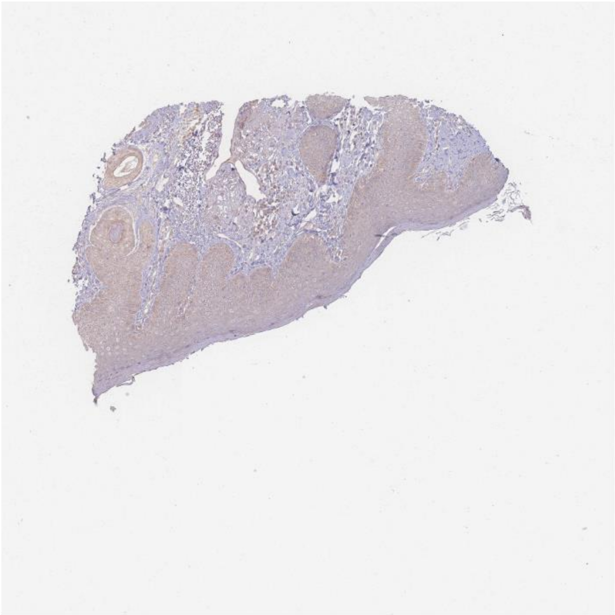

Rps27a Ripk1

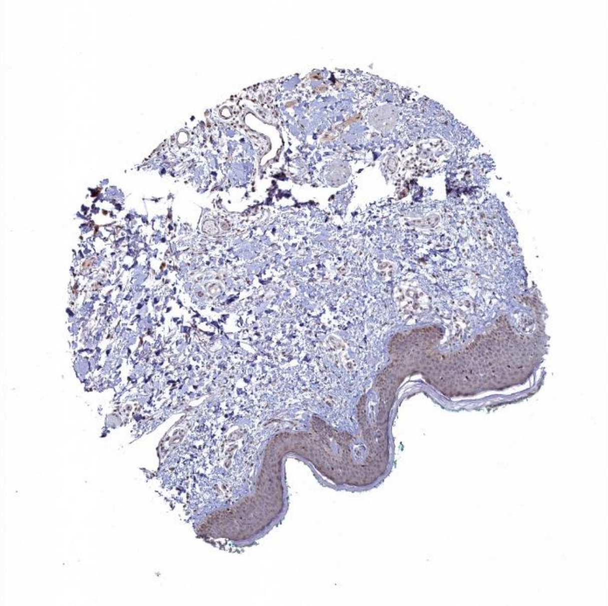

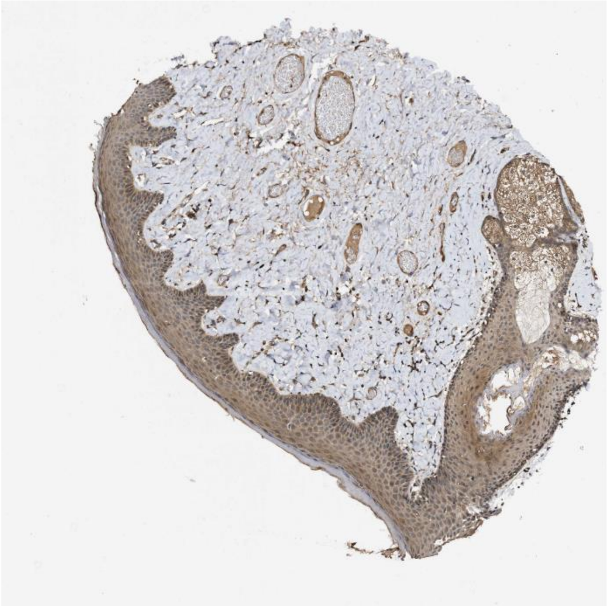

Efna1 Epha1

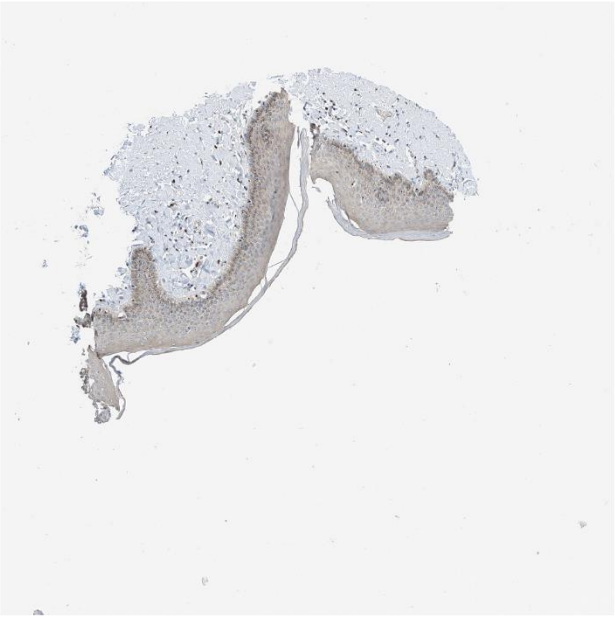

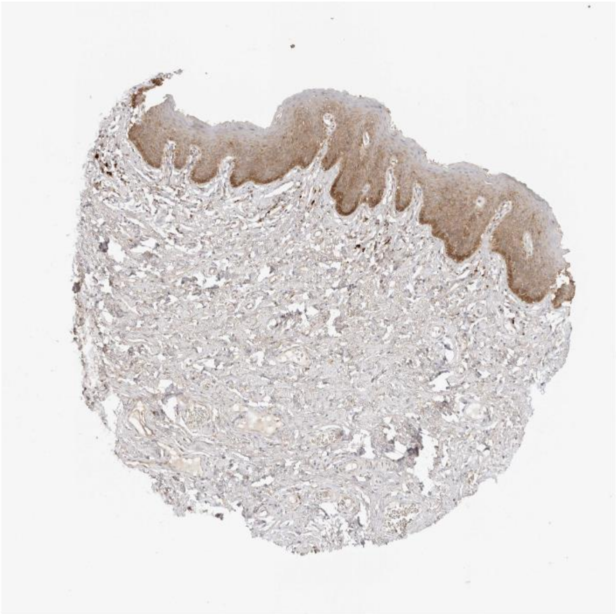

Tgfa Egfr

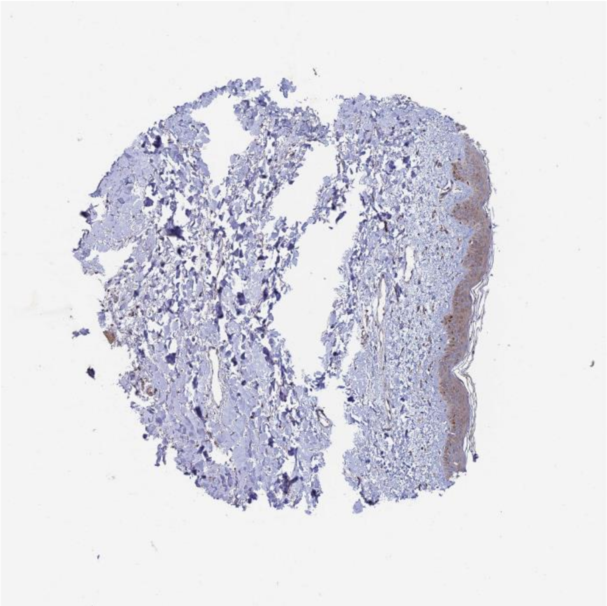

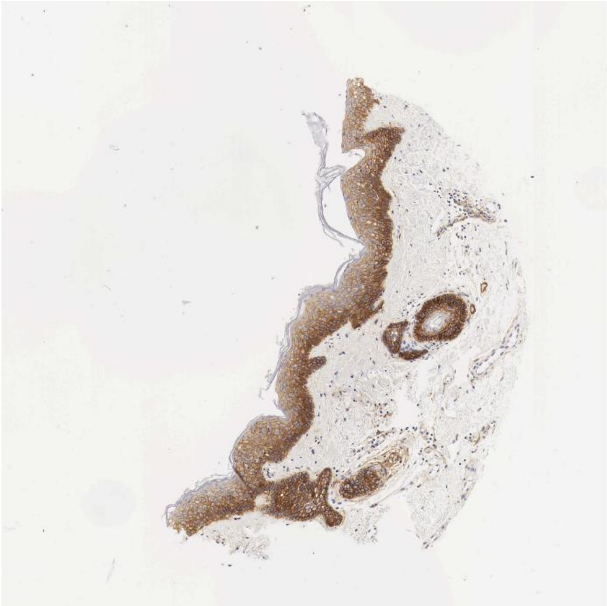

Efnb1 Erbb2

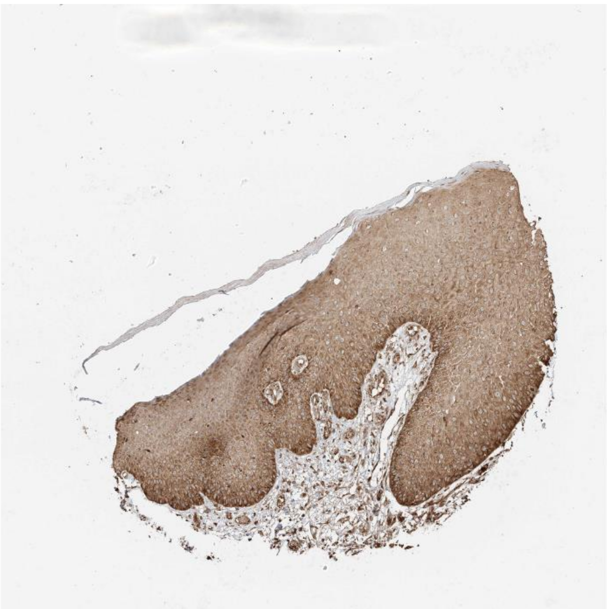

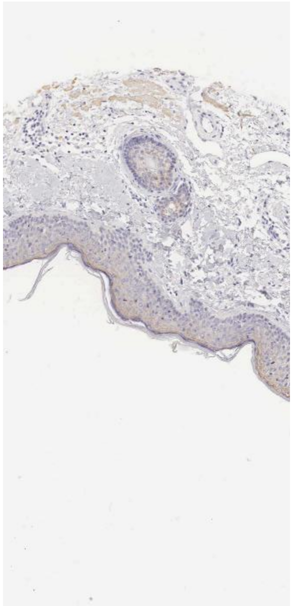

Mapk1 Fgfr2

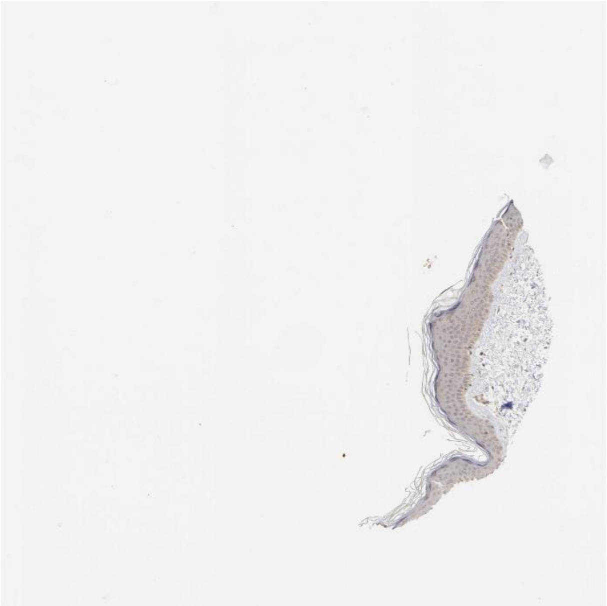

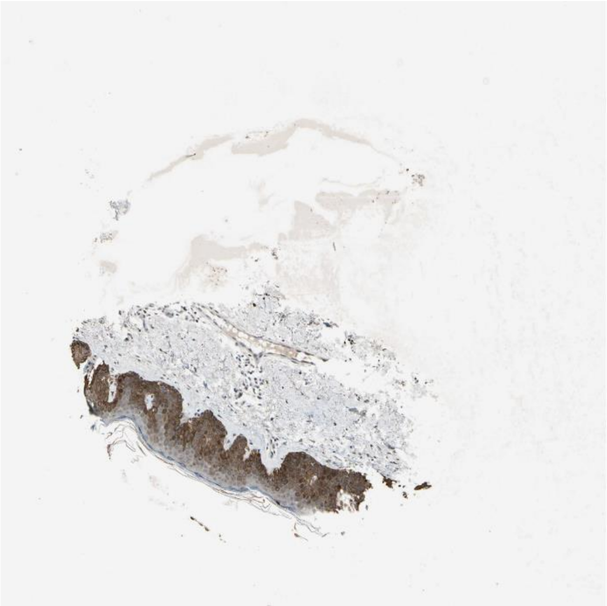

Jag1 Notch1

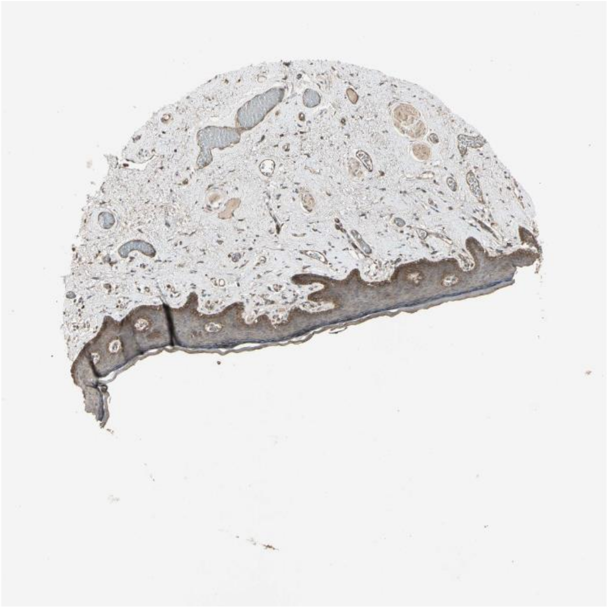

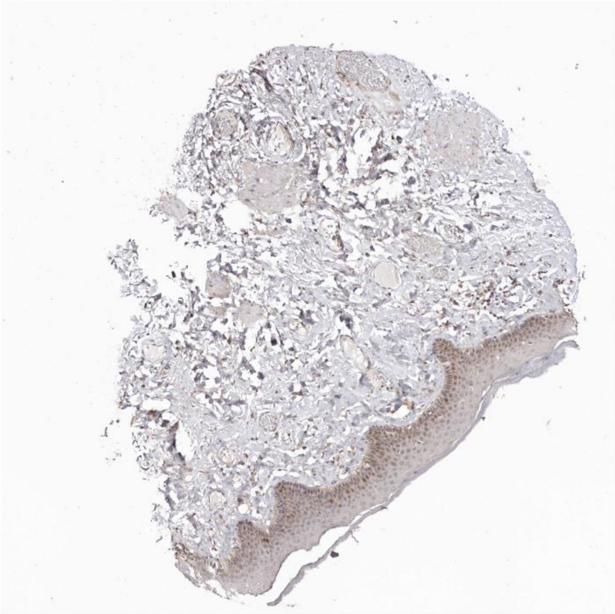

Il18 Il18r1

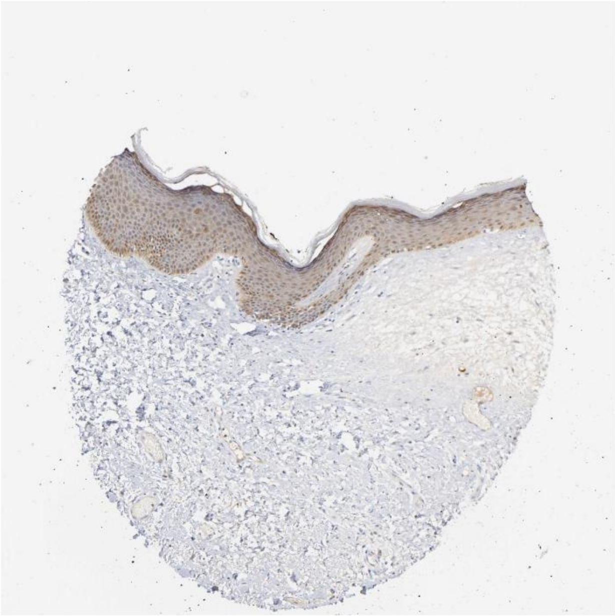

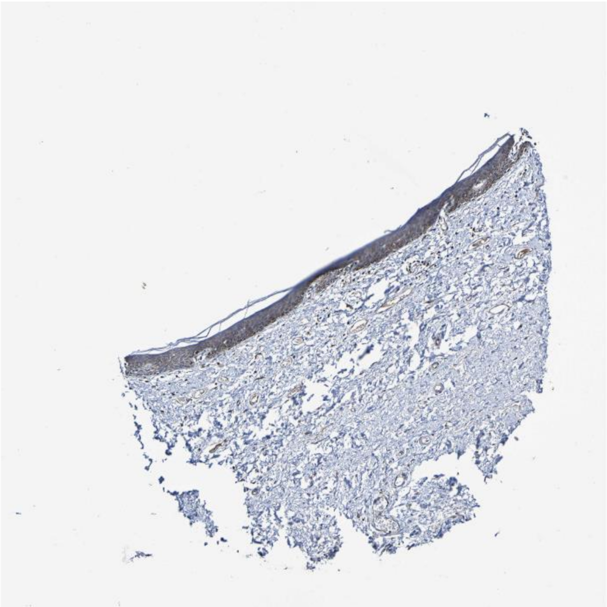

**Psen1** Ncstn

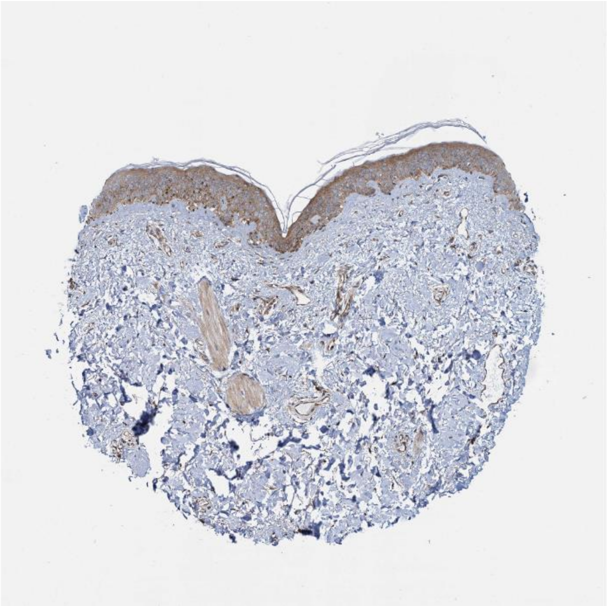

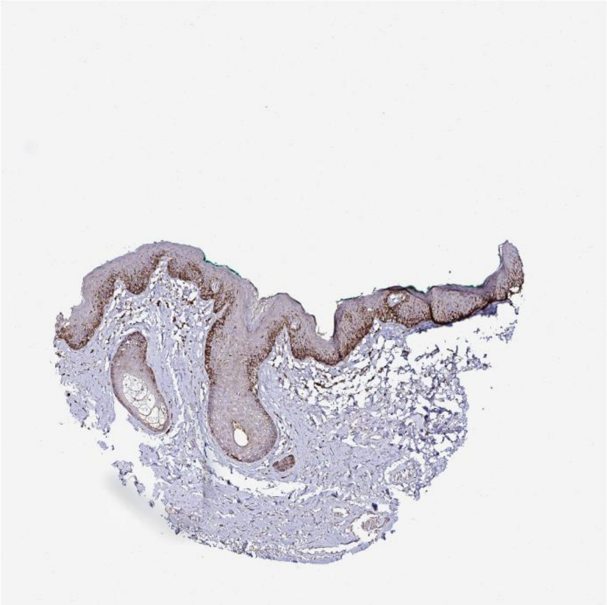

B2m Cd3g

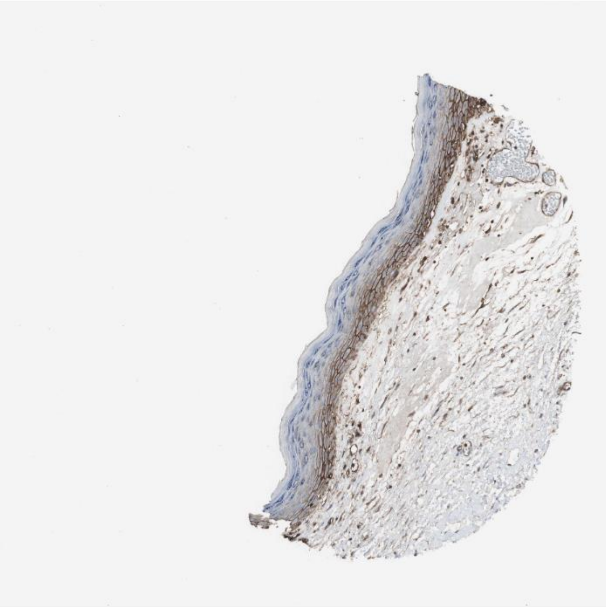

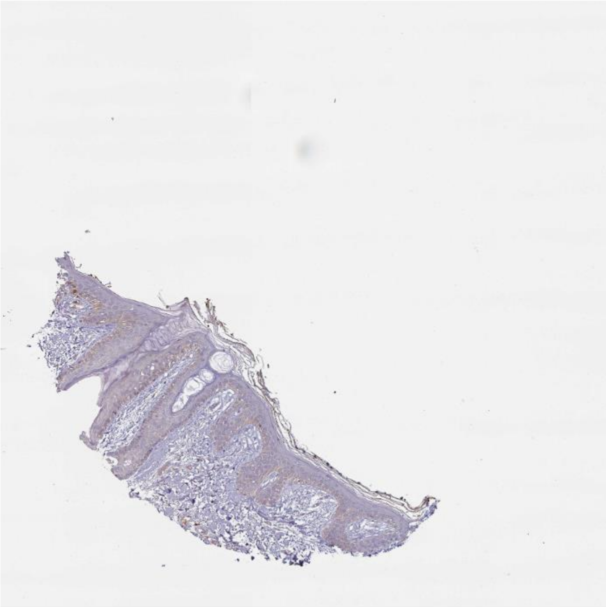

Ubc Tgfbr1

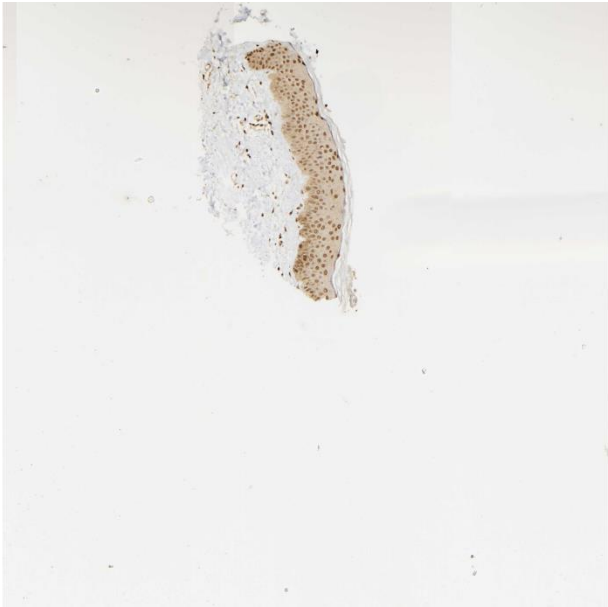

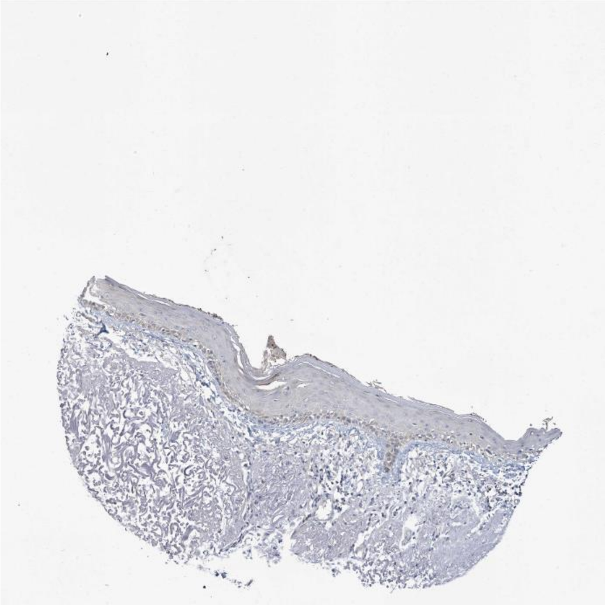

Fgf18 Fgfr3

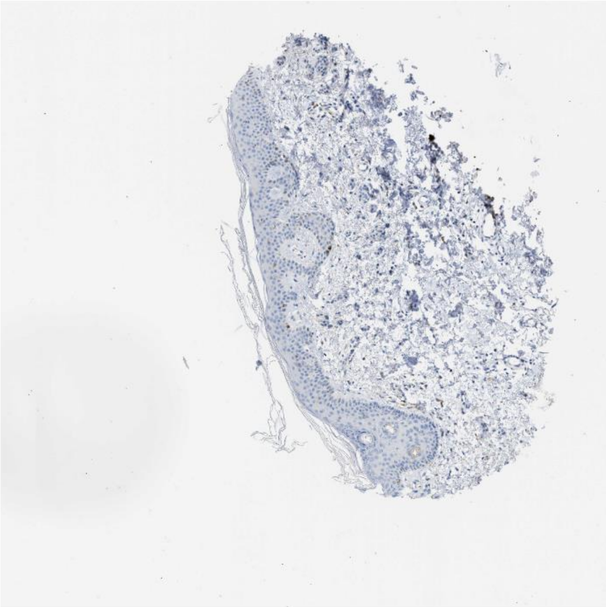

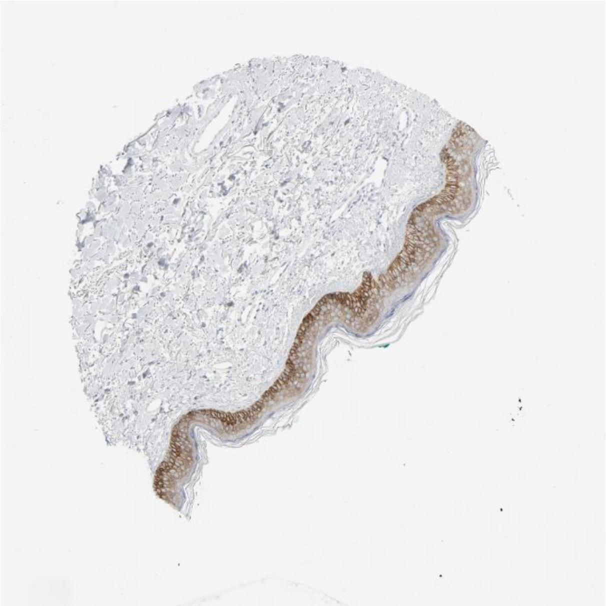

Gpi1 Amfr

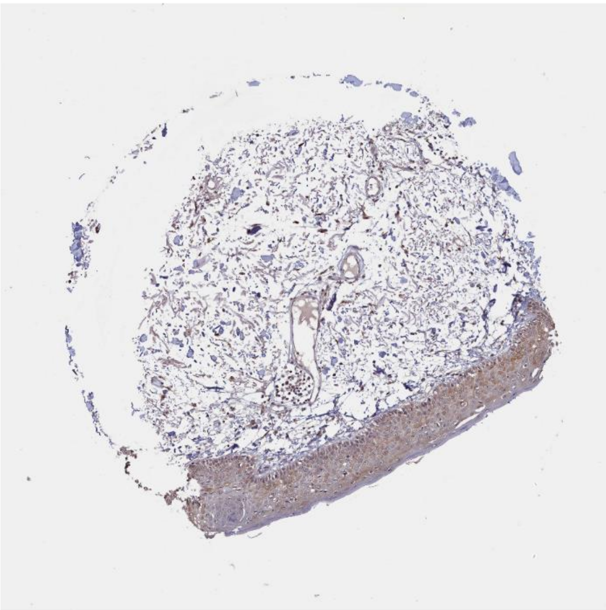

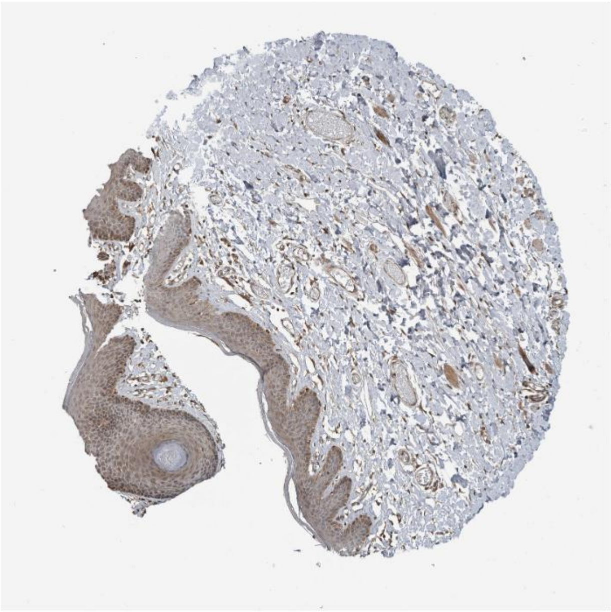

Cdh1 Erbb3

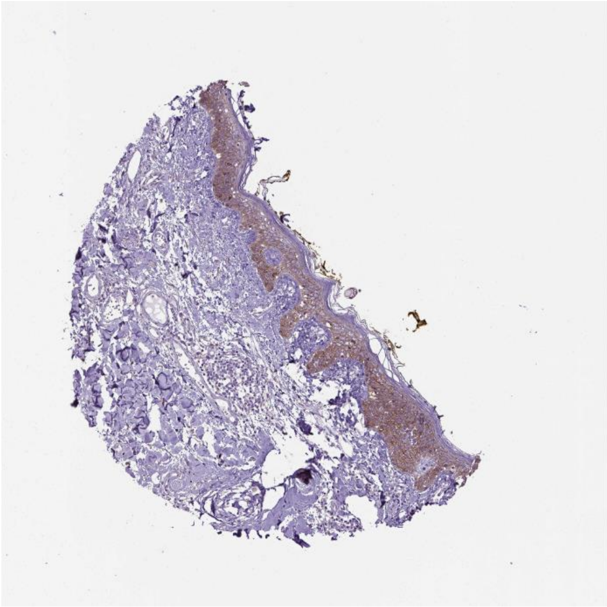

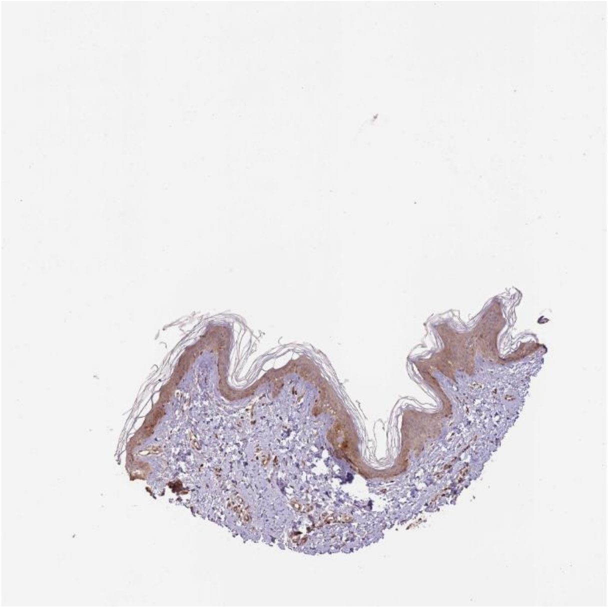

Il1rn Il1r2

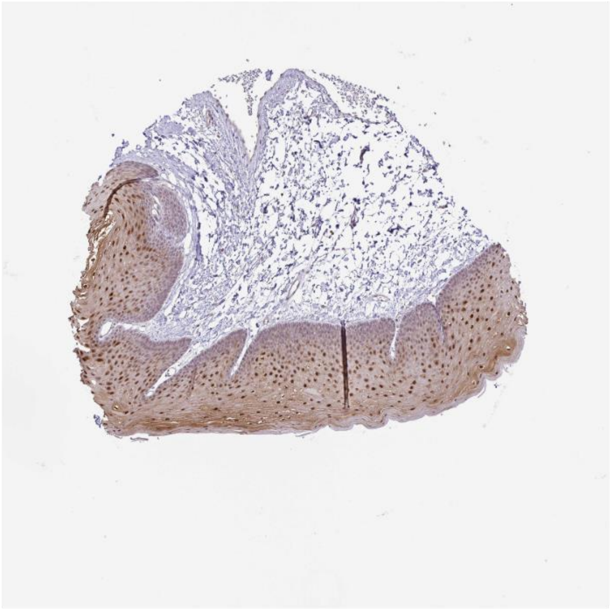

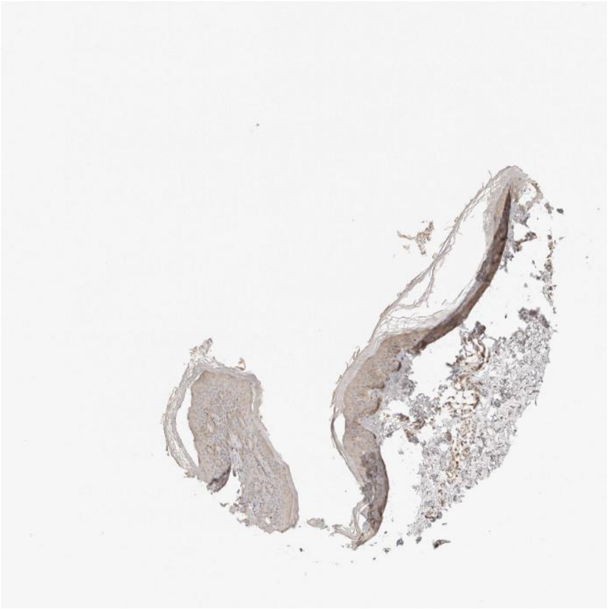

Efna5 Epha1

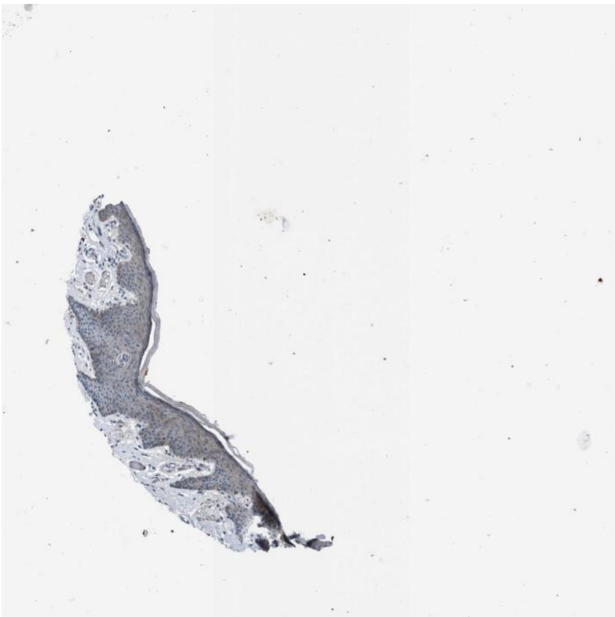
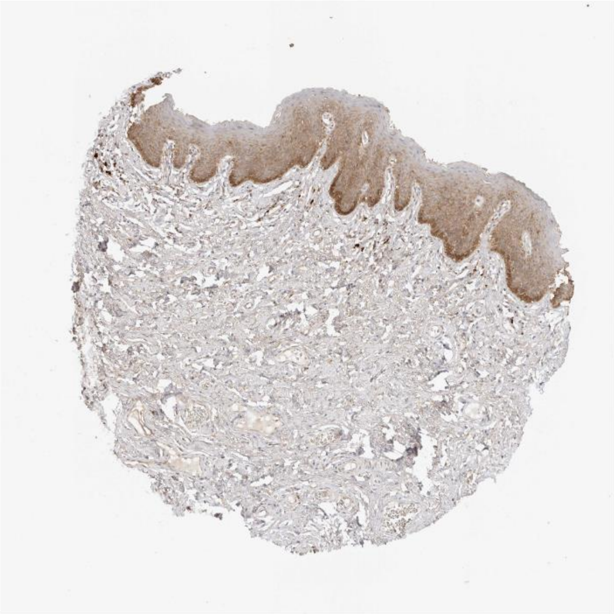

Ubb Ripk1

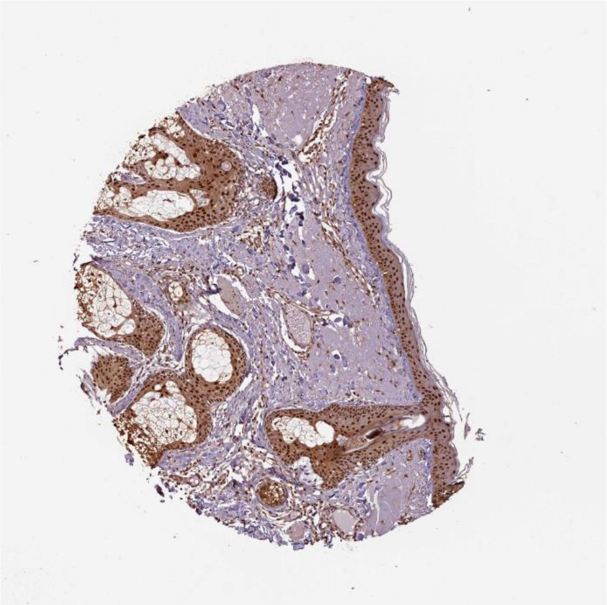
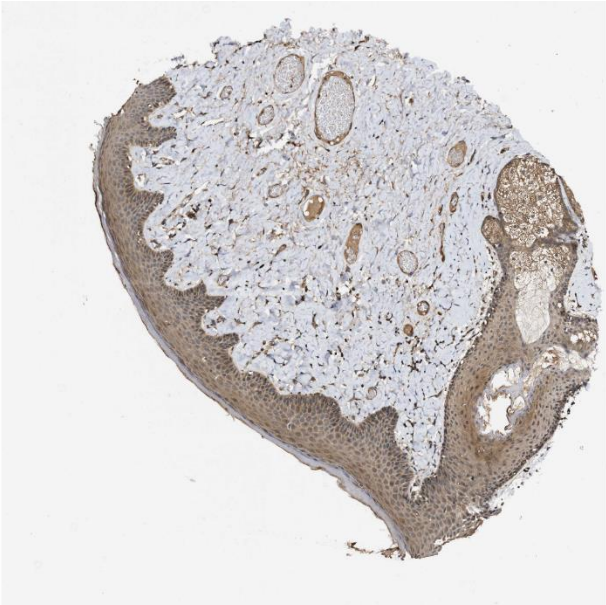

Thbs1 Sdc1

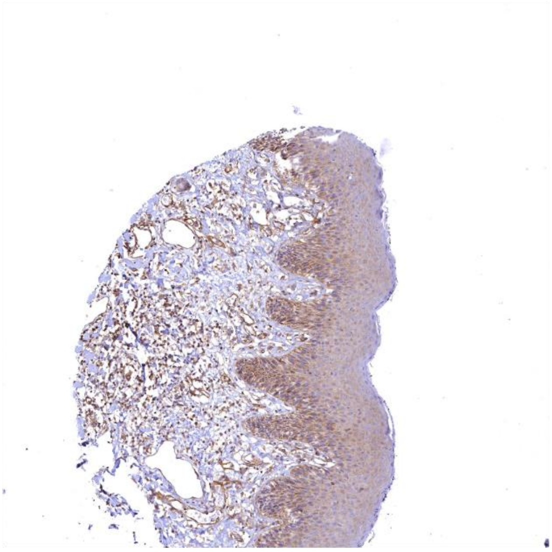

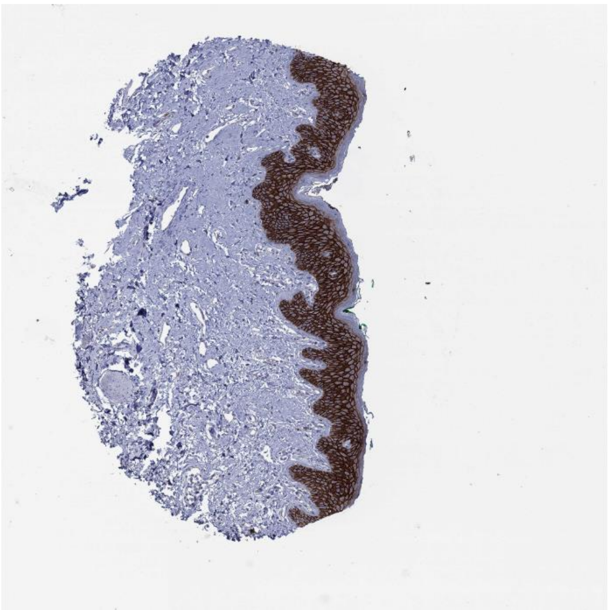

Apoe Lrp5

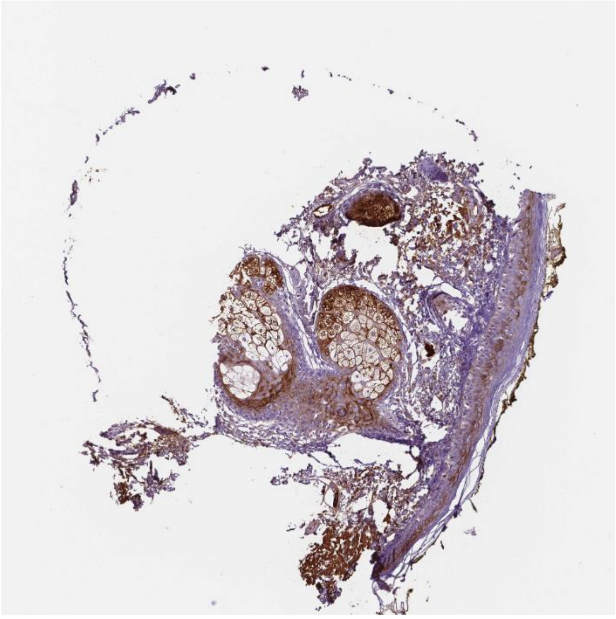

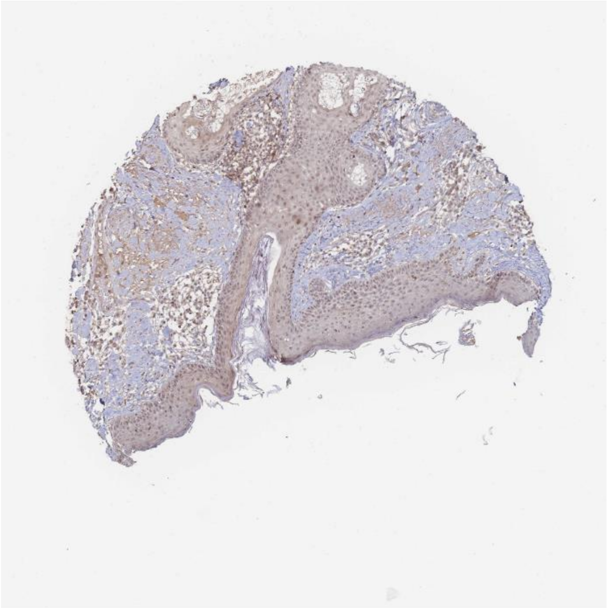

Apoe Sorl1

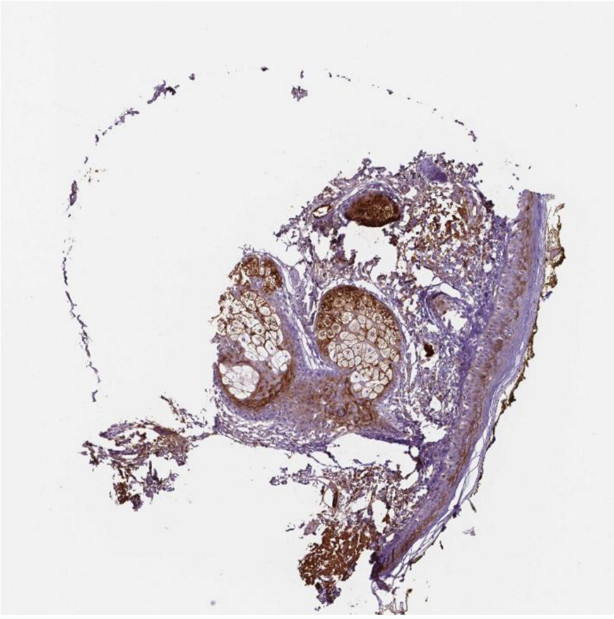

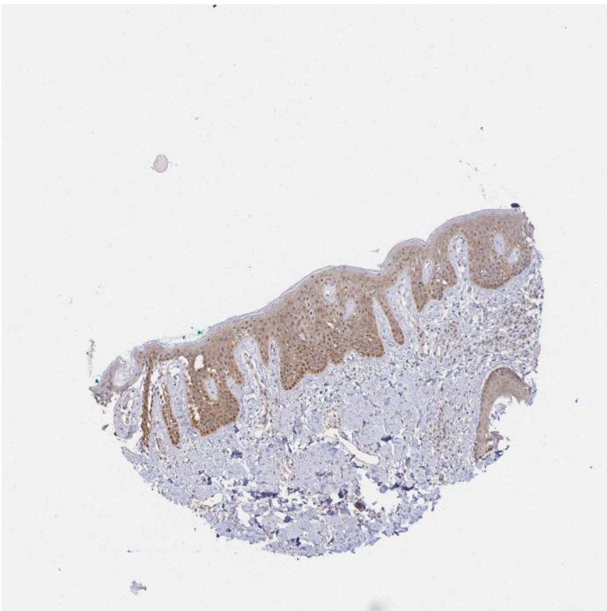

Rps27a Tgfbr1

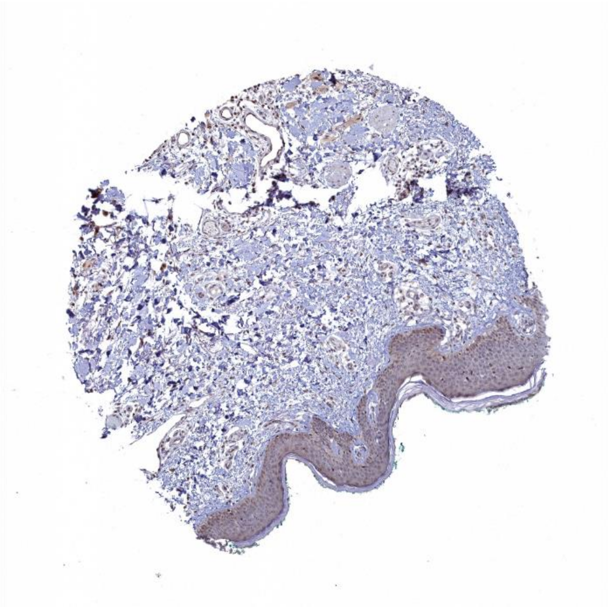

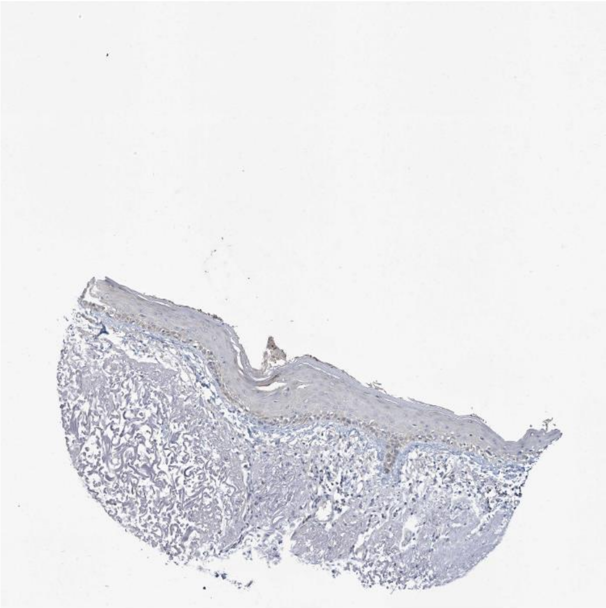

Efnb2 Rhbdl2

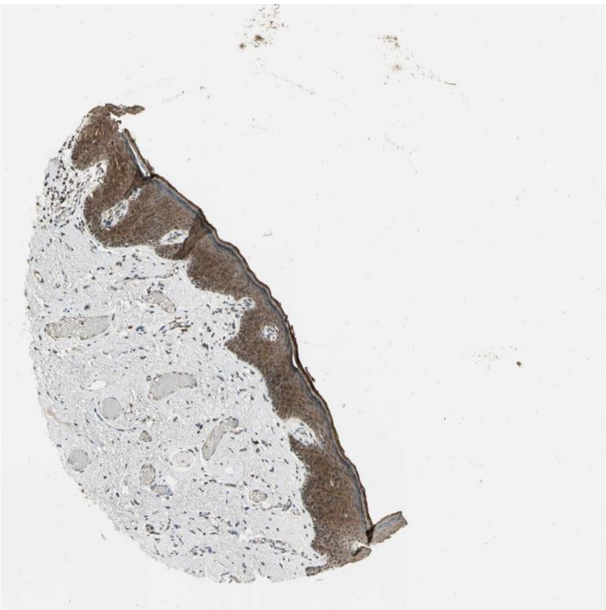
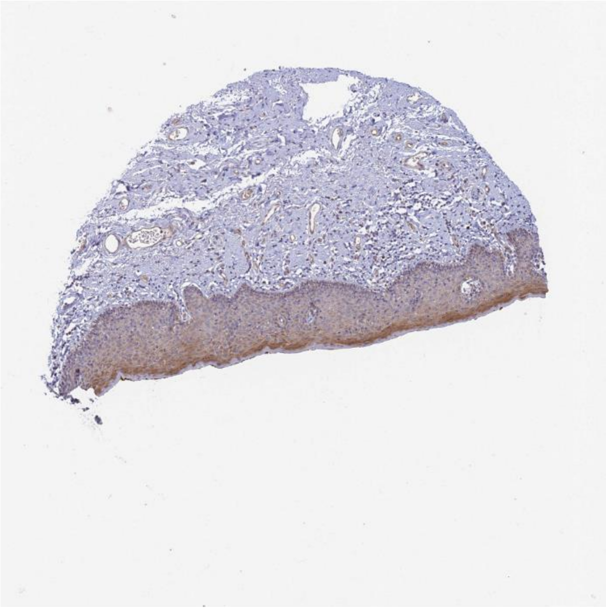

Gas6 Axl

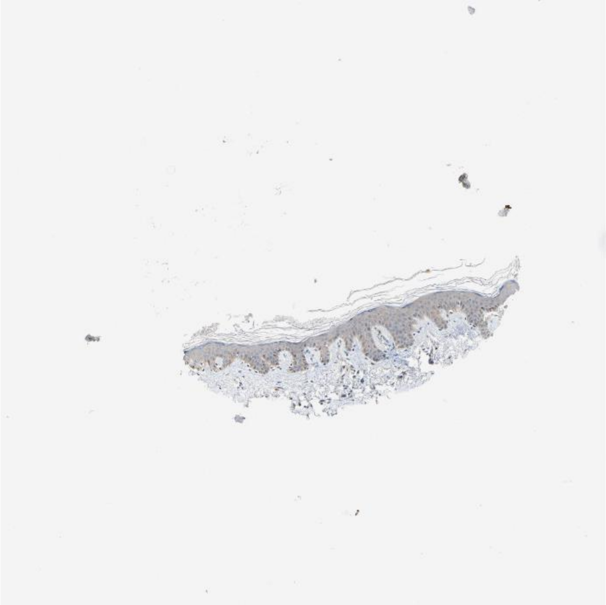
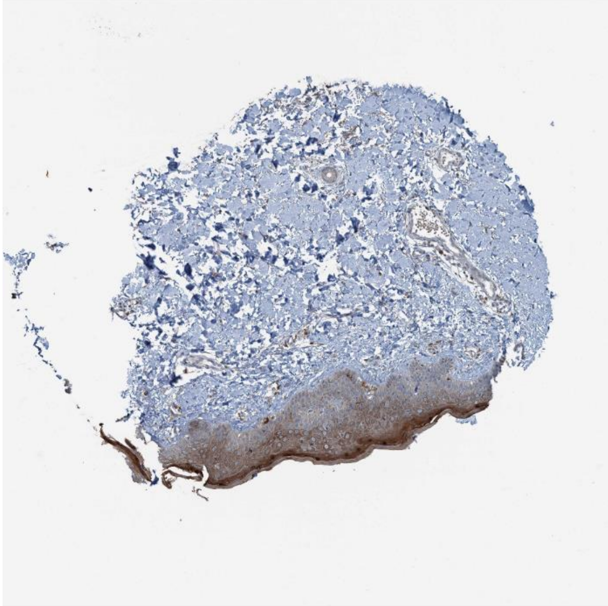

Dusp18 Itgb1

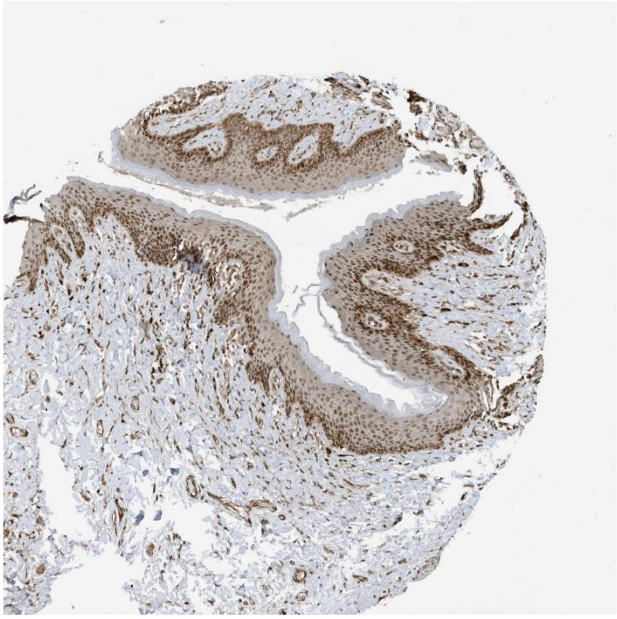

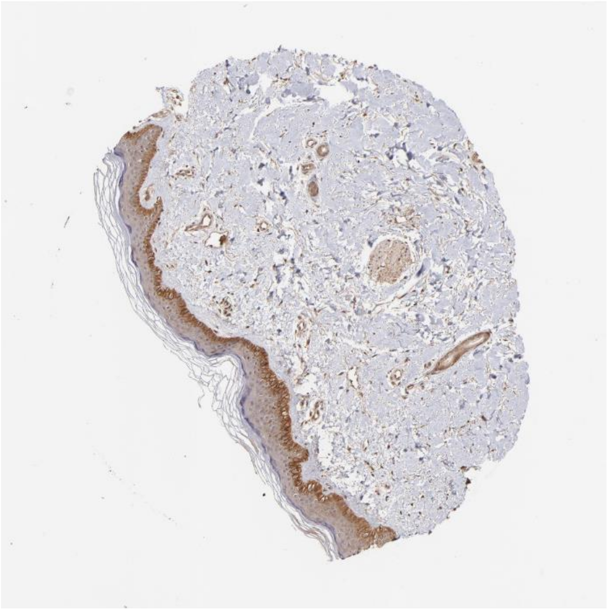

Efnb2 Ephb4

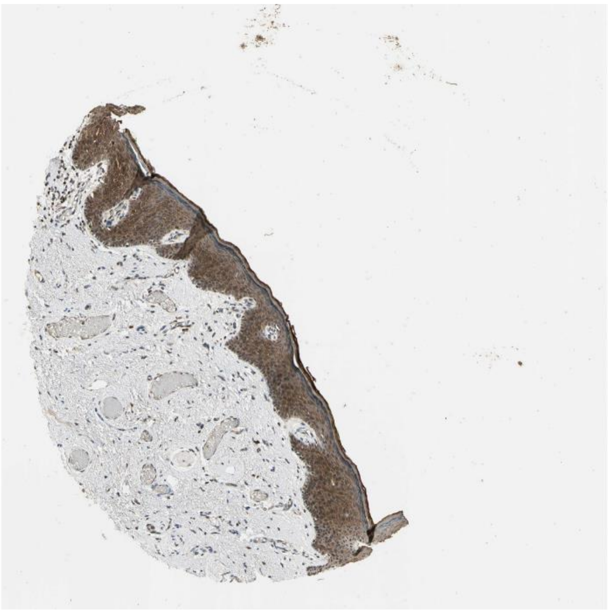

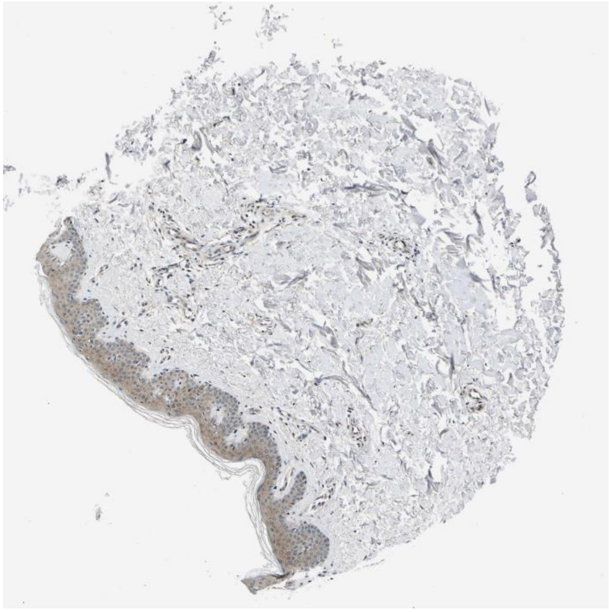

Tgs1 Rxra

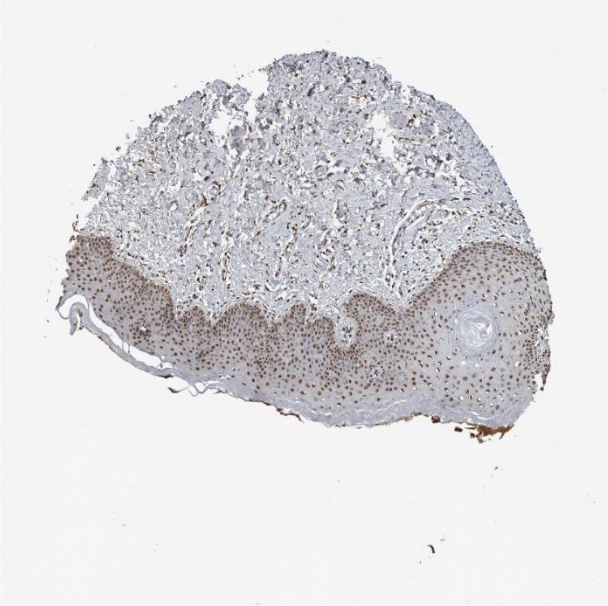

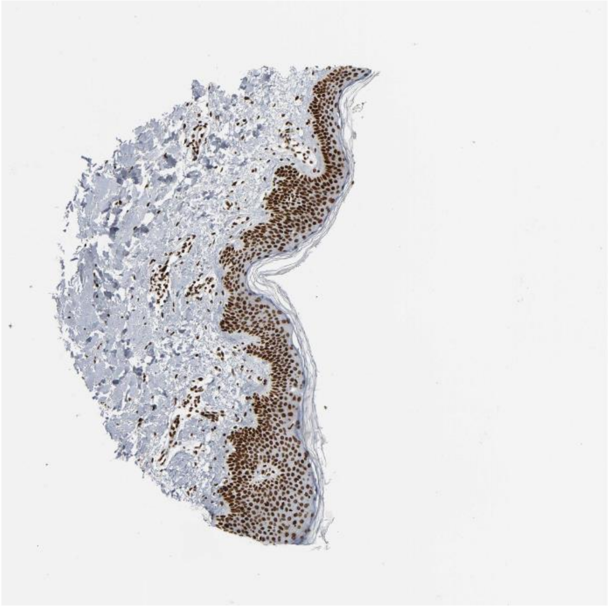

Gnai2 Egfr

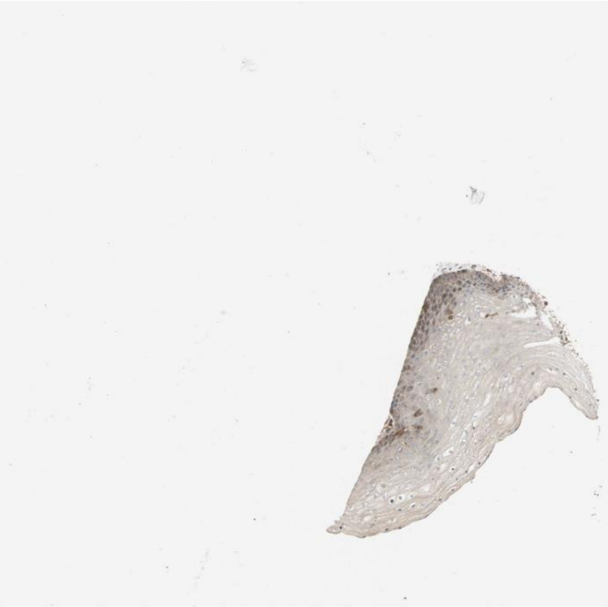

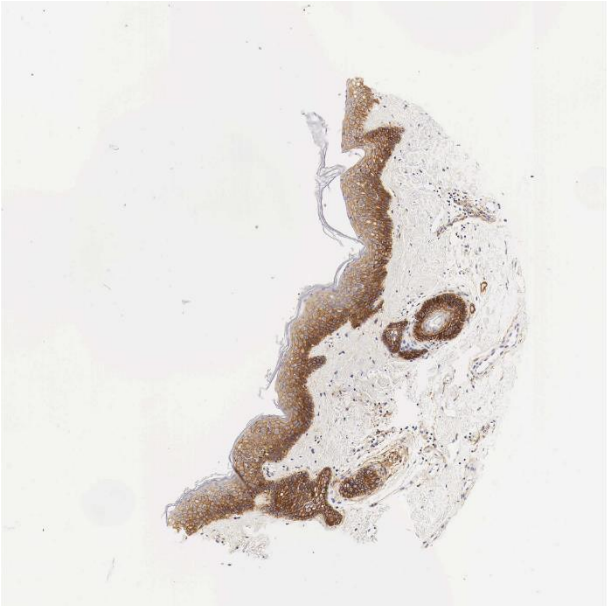

Lama5 Sdc1

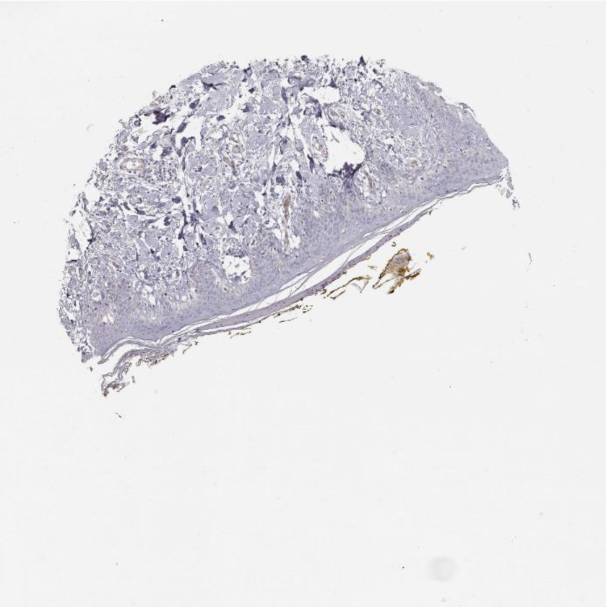

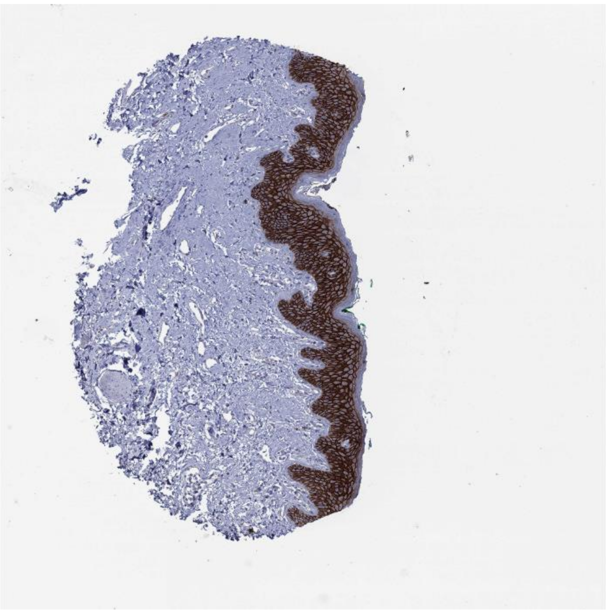

Cgn Tgfbr1

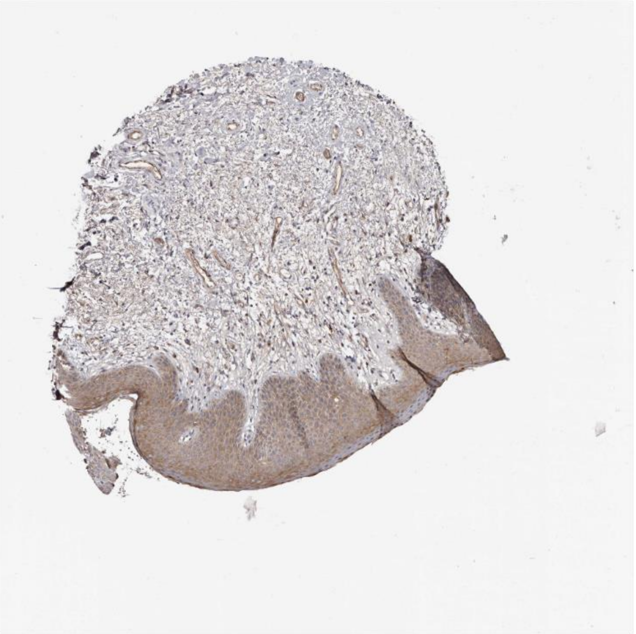

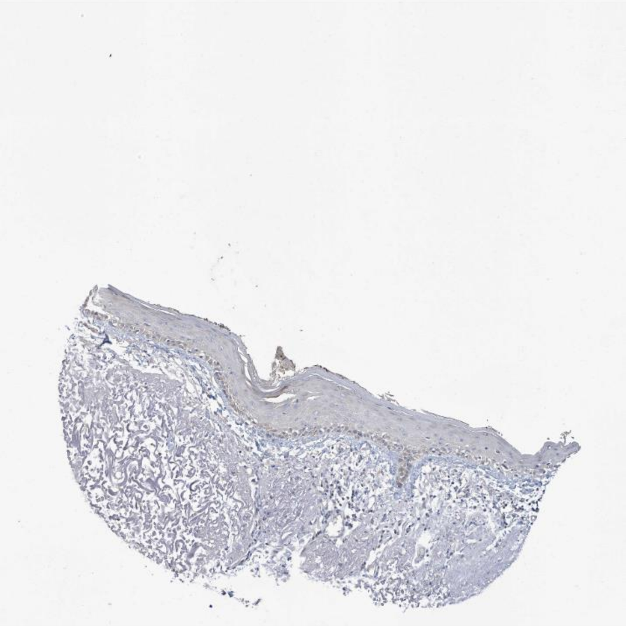

Areg Egfr

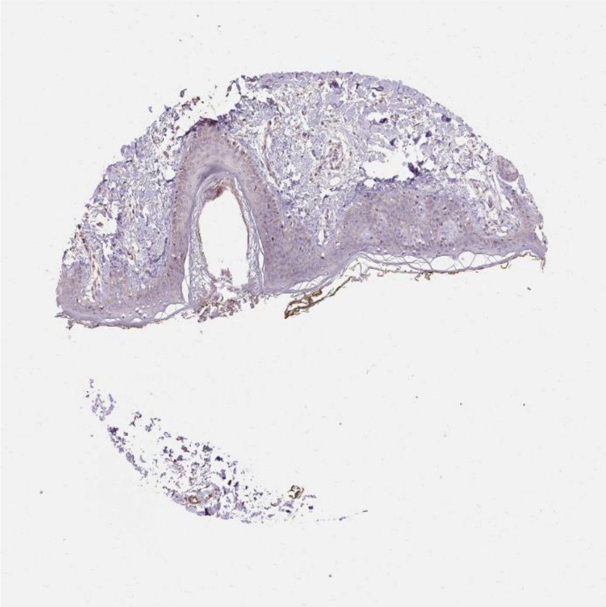

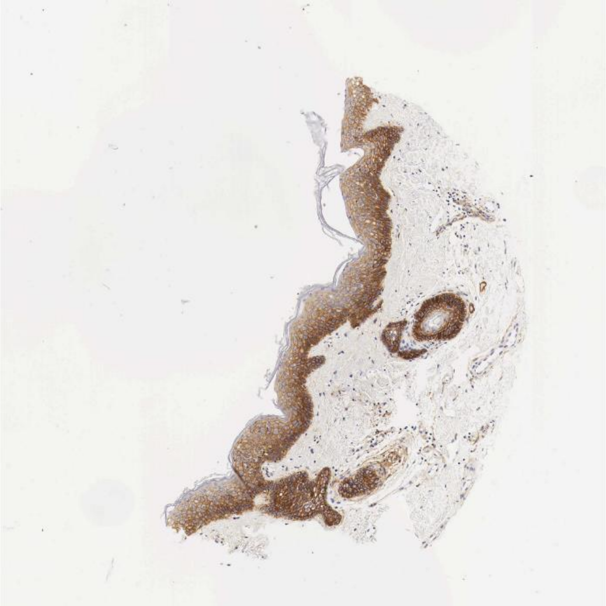

Lrpap1 Ldlr

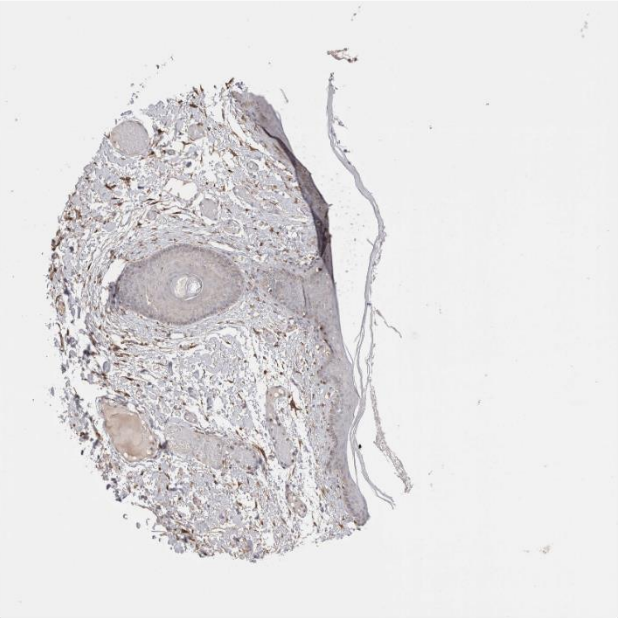

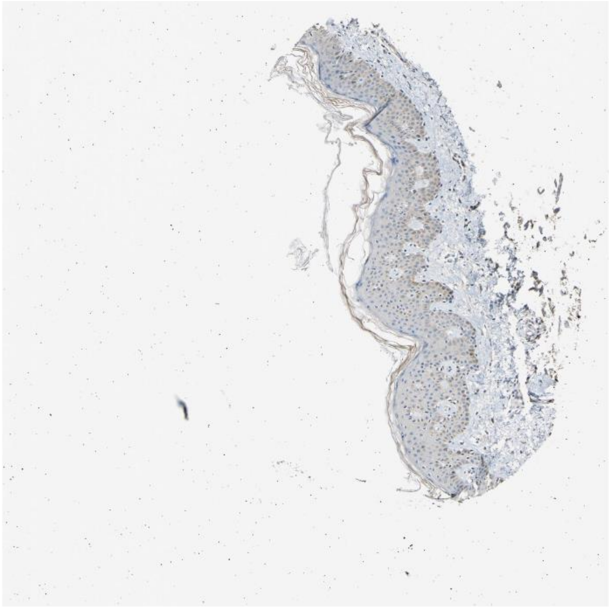

B2m Hfe

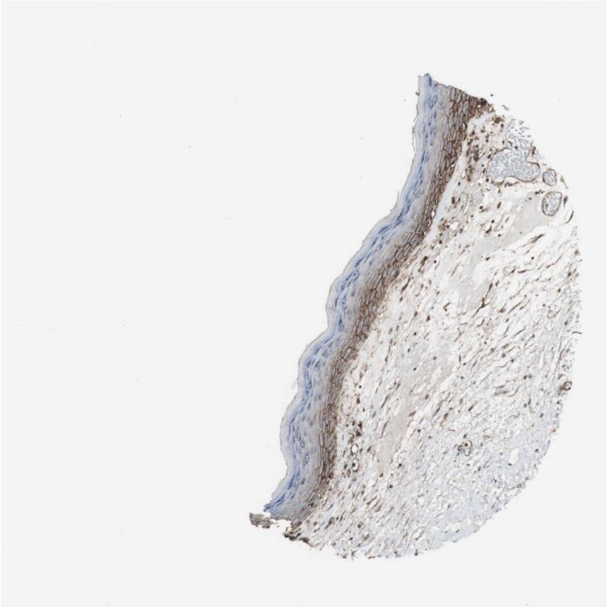

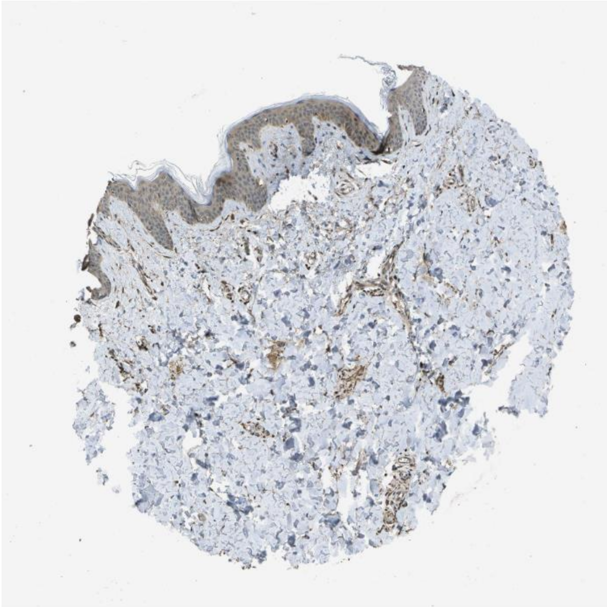

Cdh1 Lrp5

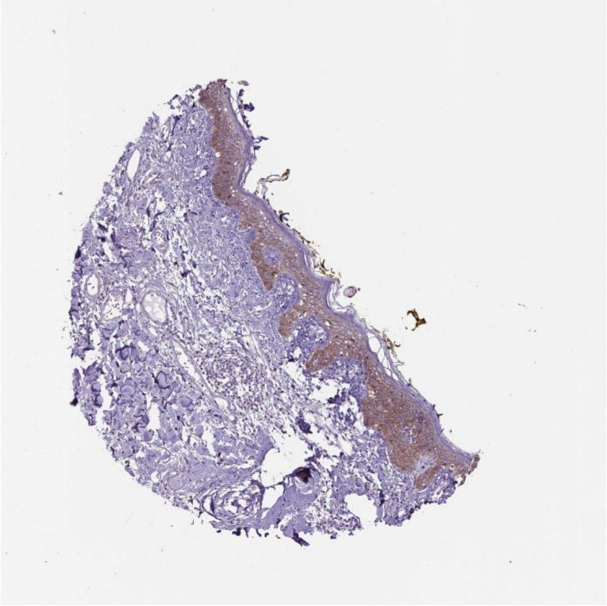

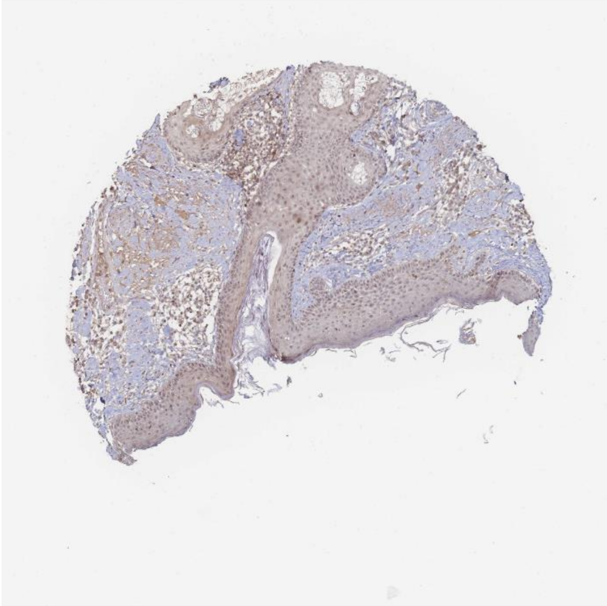

Ptdss1 Jmjd6

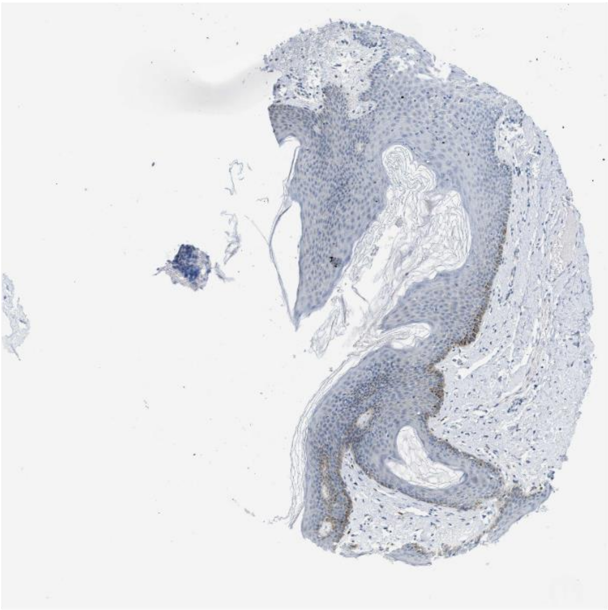

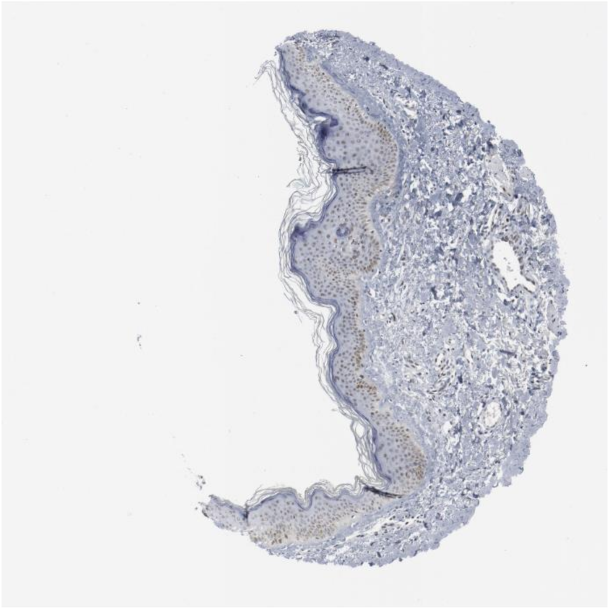

Gnai2 Igf1r

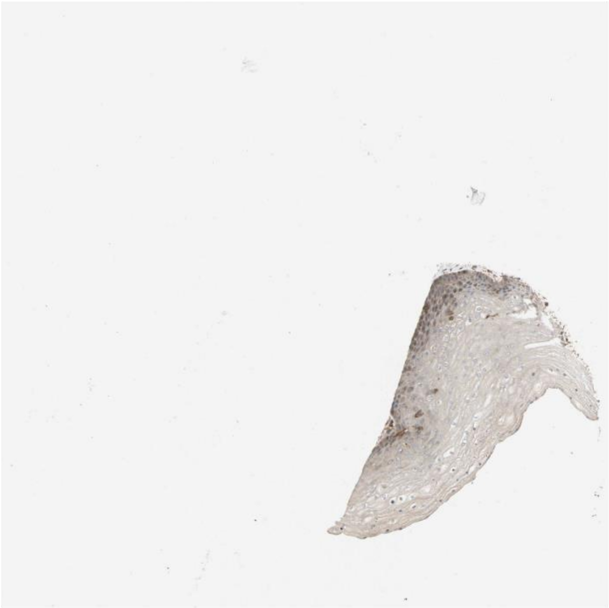

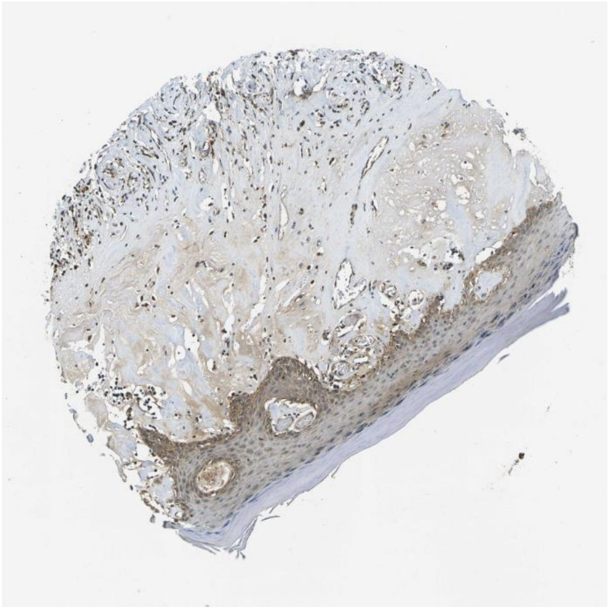

Gnai2 Cav1

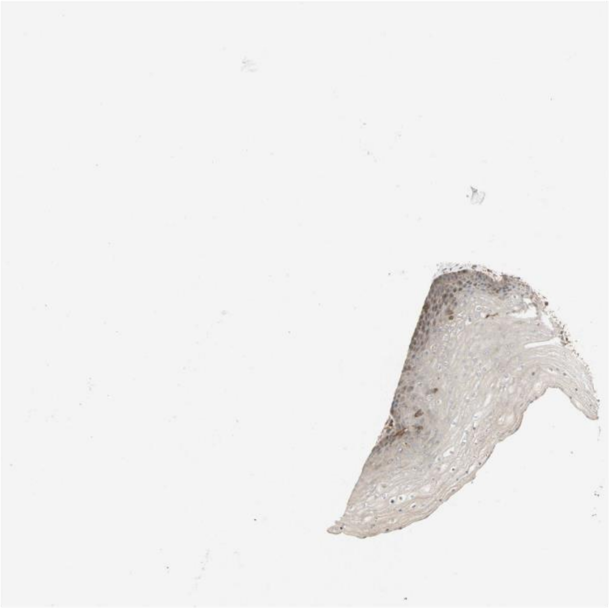

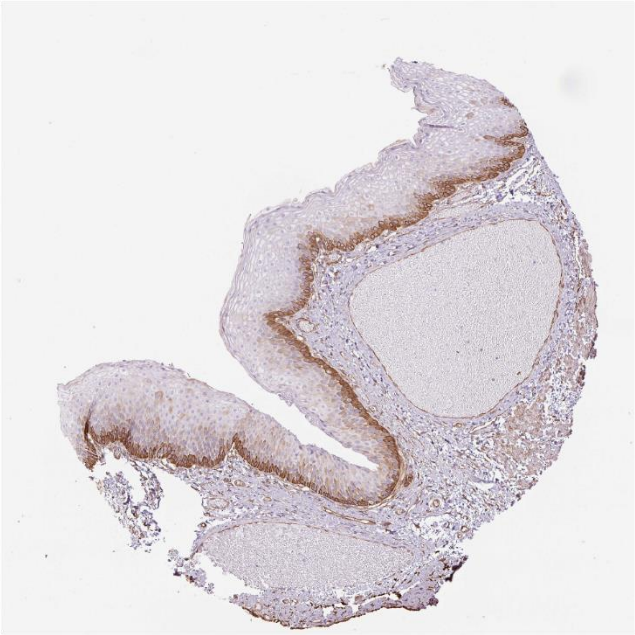

Lamb3 Itga6

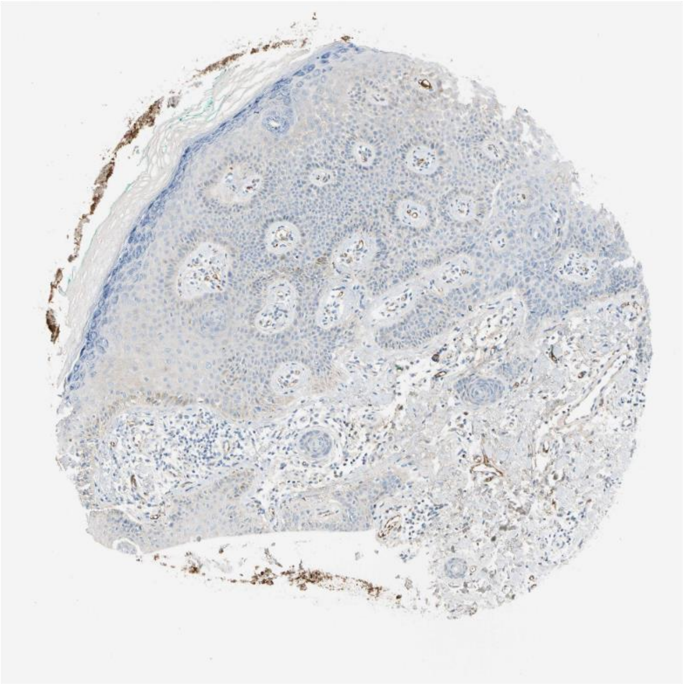

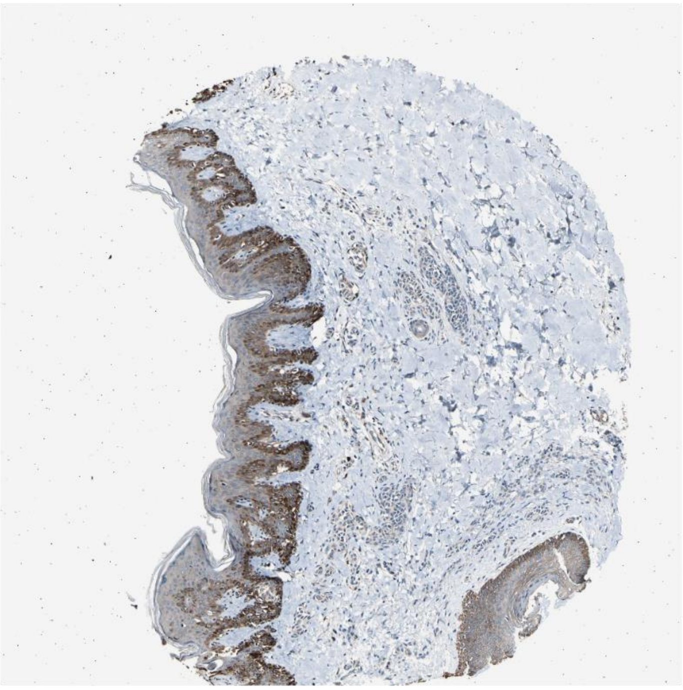

Agrn Lrp4

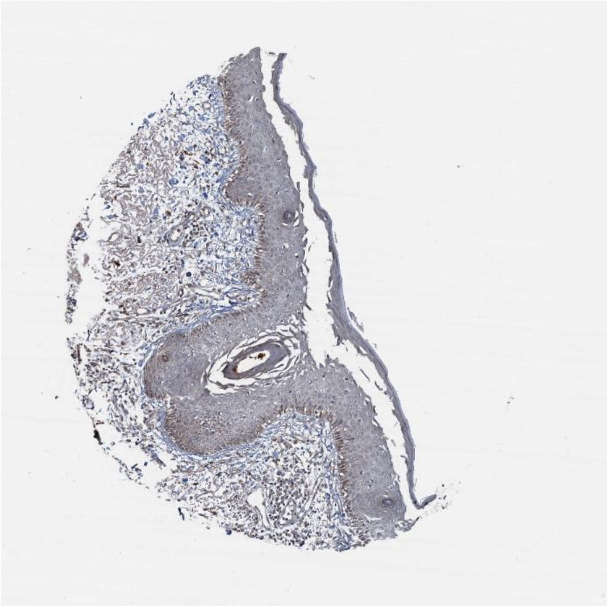

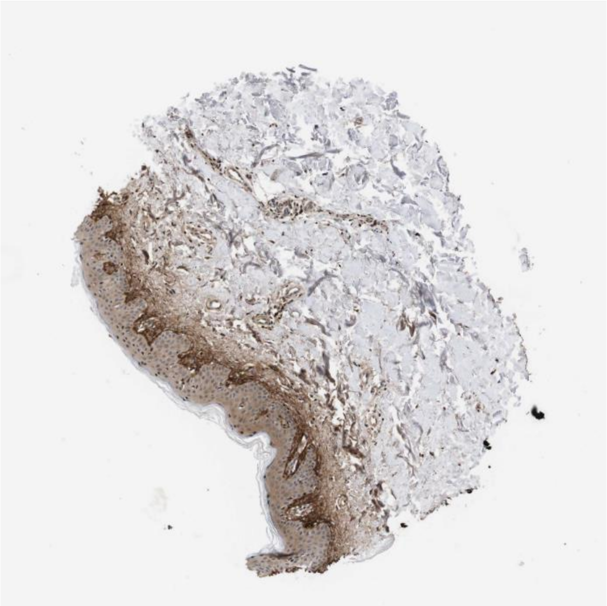

Jag1 Notch2

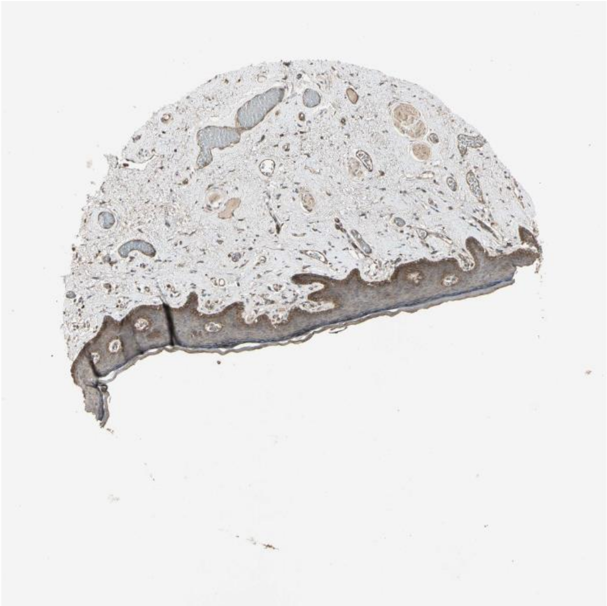

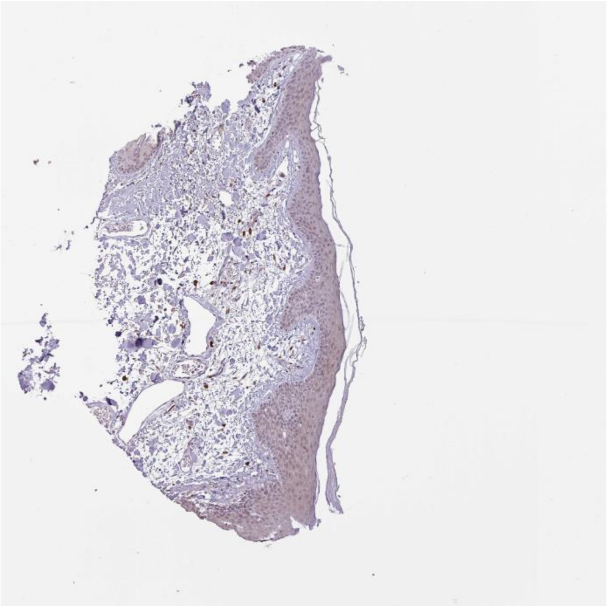

Fgf18 Fgfr2

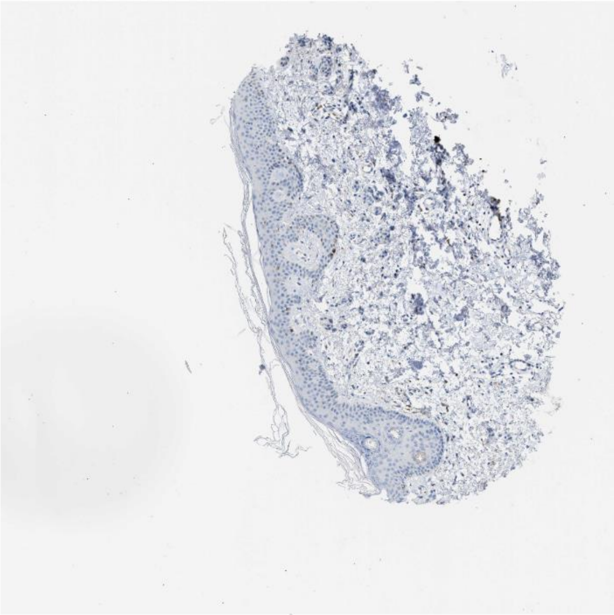

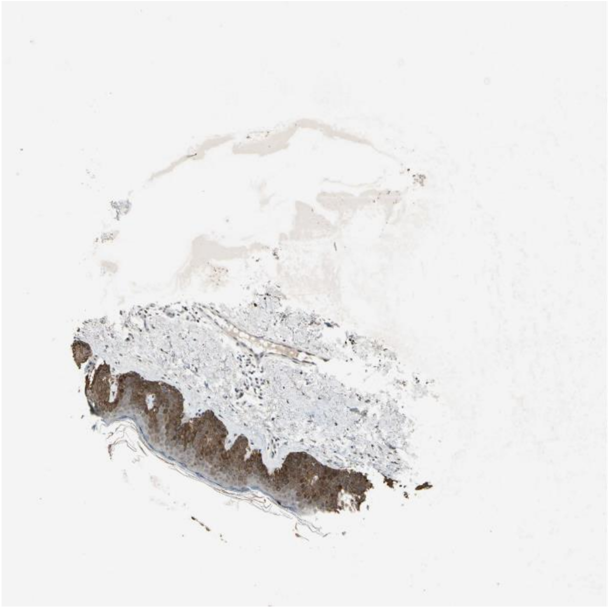

Psap Sort1

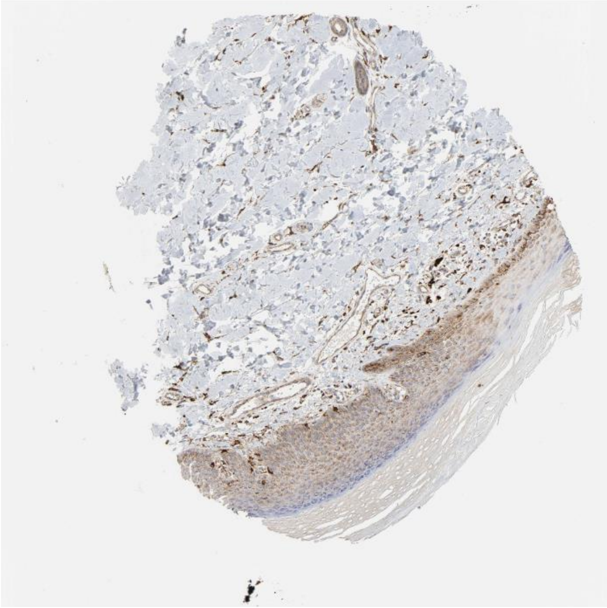

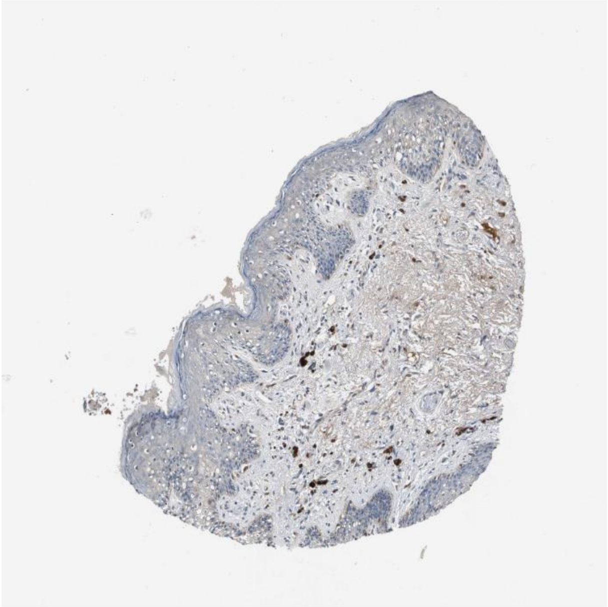

B2m Cd247

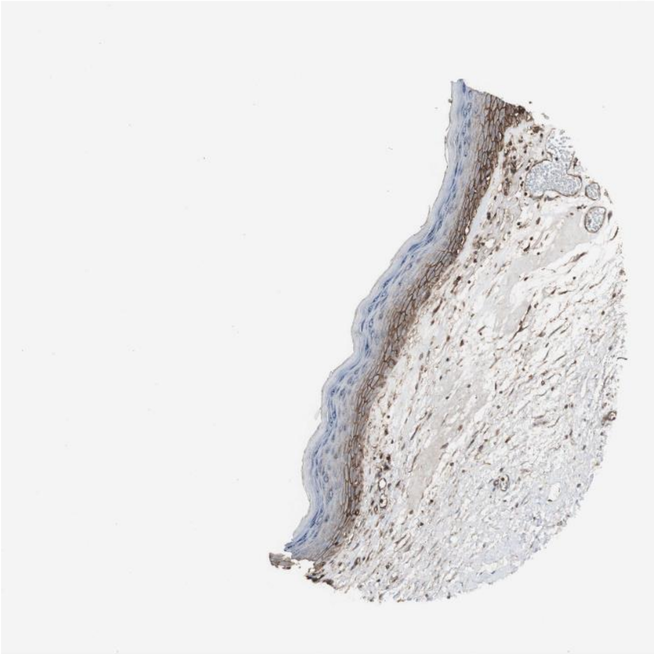

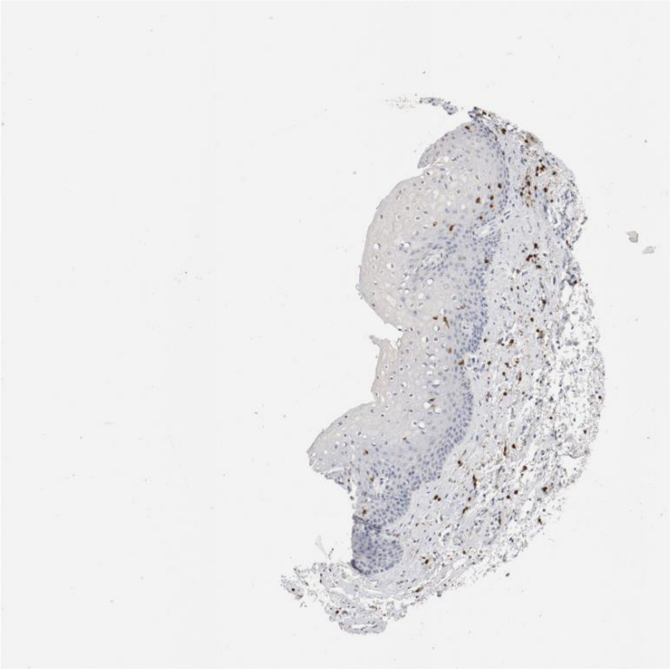

Gnai2 S1pr5

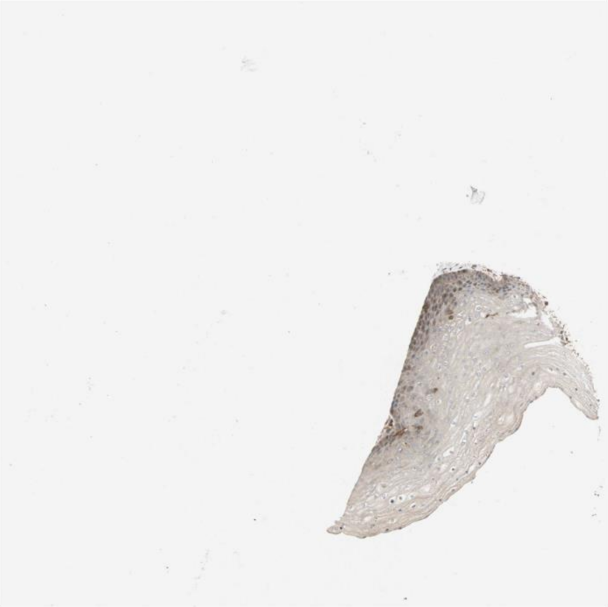

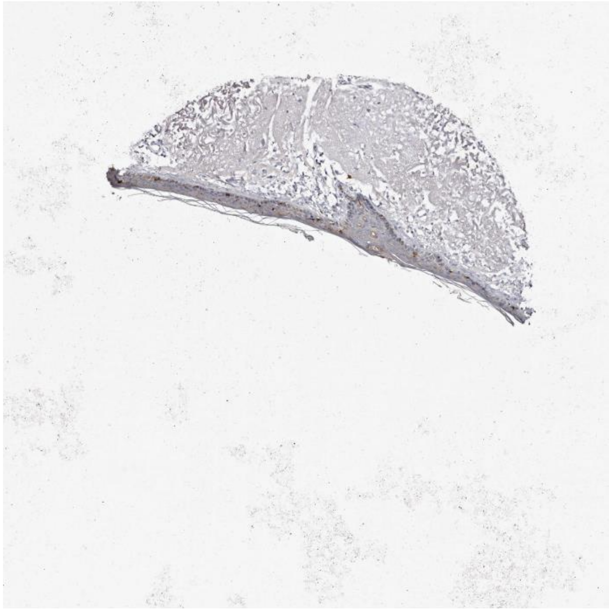

Il1rn Il1r1

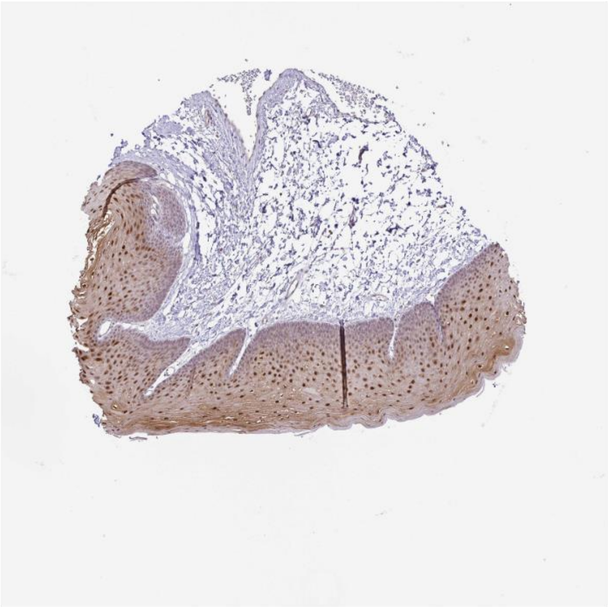

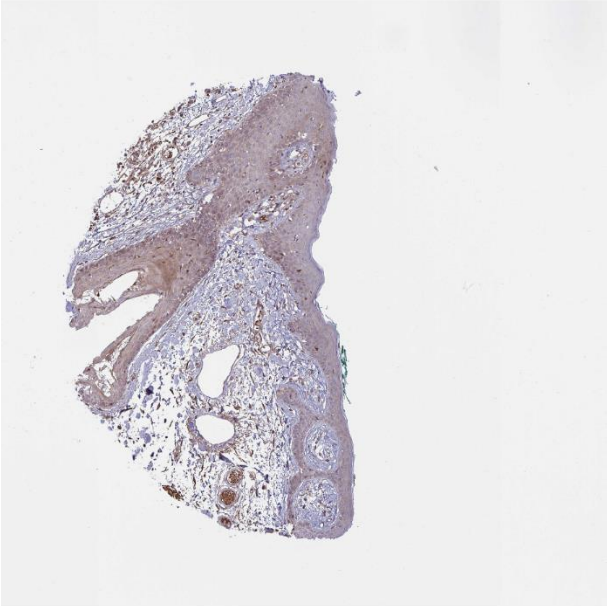

Ubb Tgfbr1

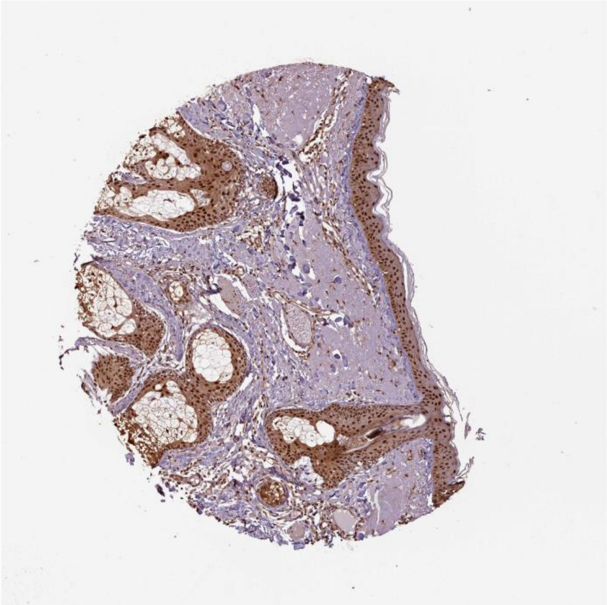

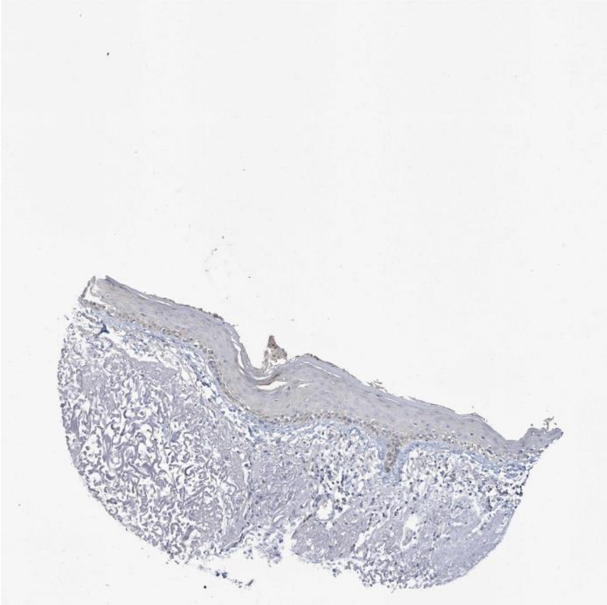

Lin7c Abca1

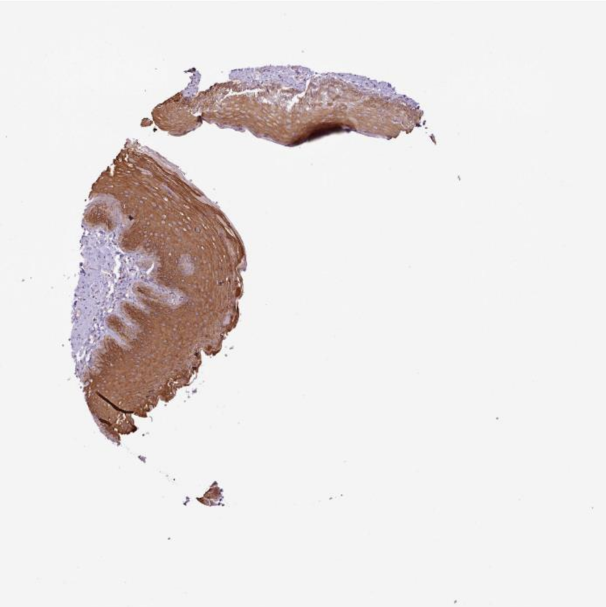

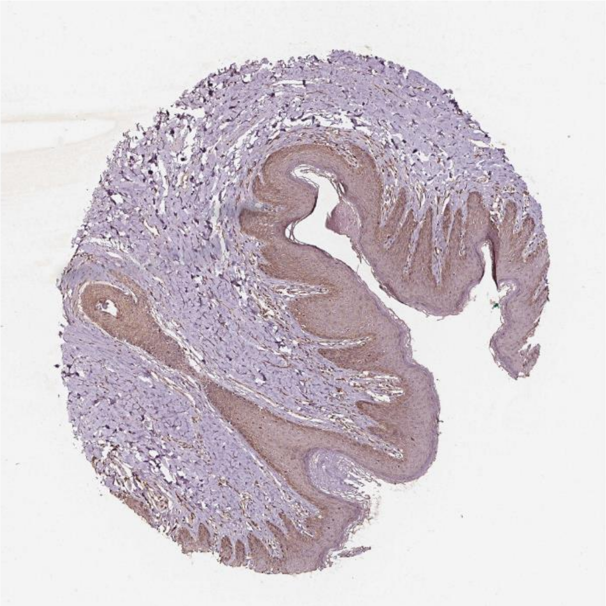

Vim **Cd44**

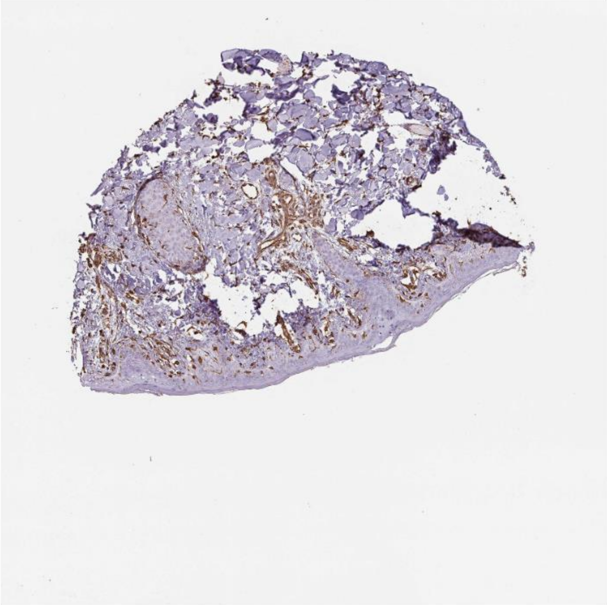

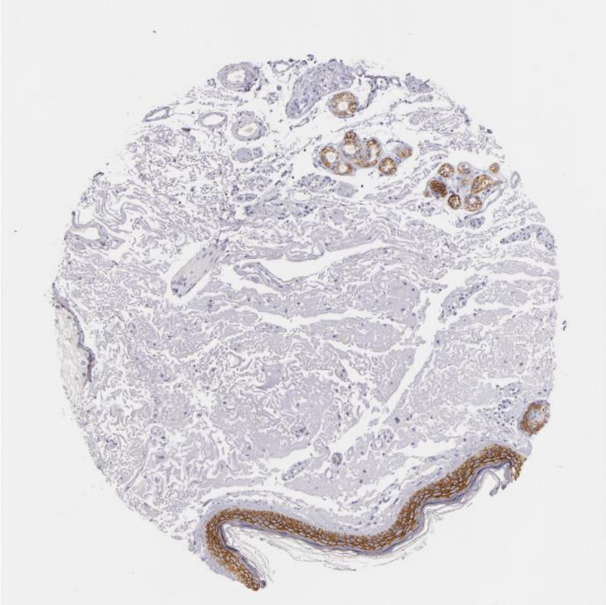

Col6a1 Itga6

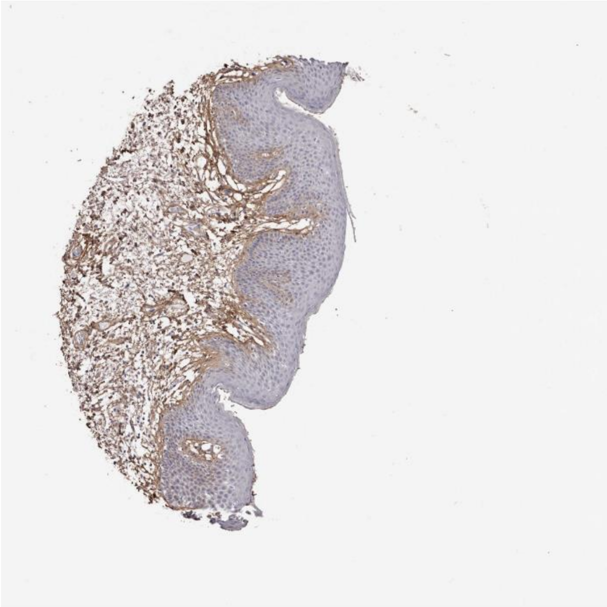

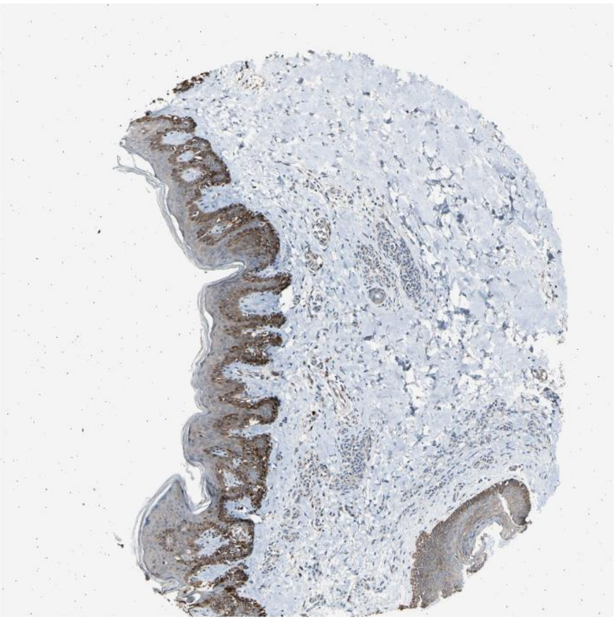

Rgma Bmpr2

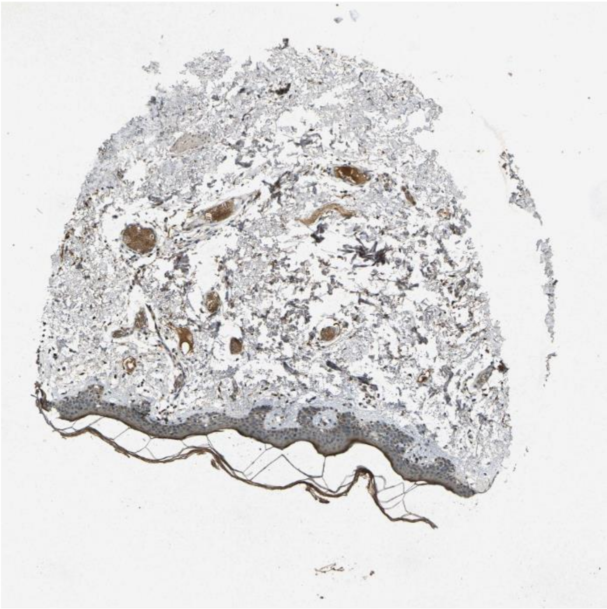

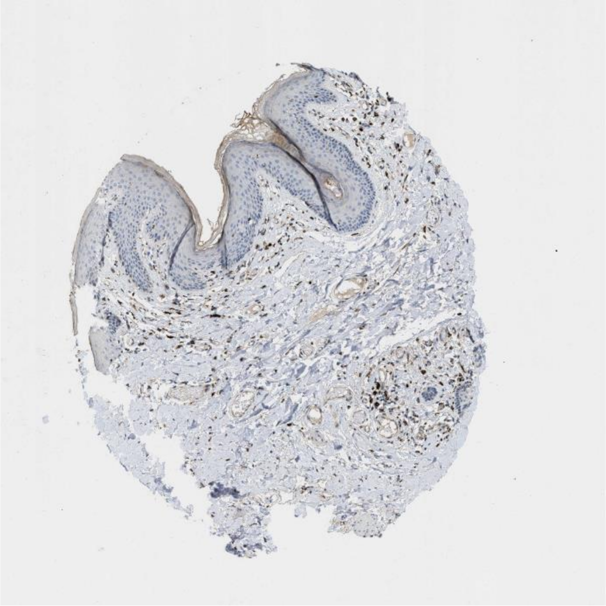

Rtn4 Rtn4rl1

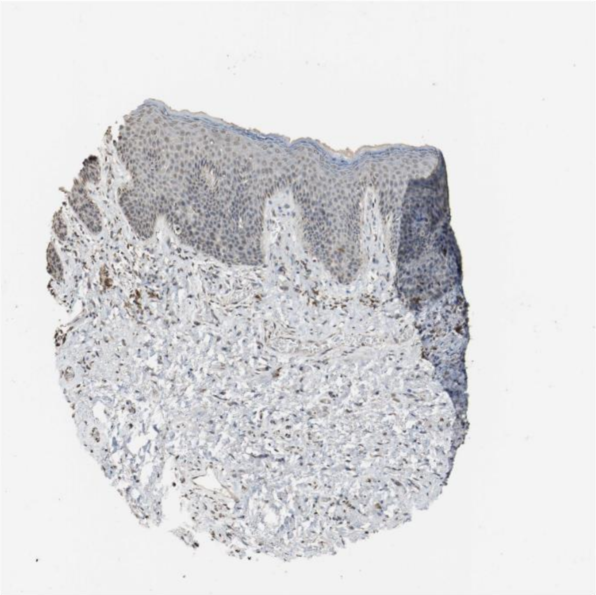
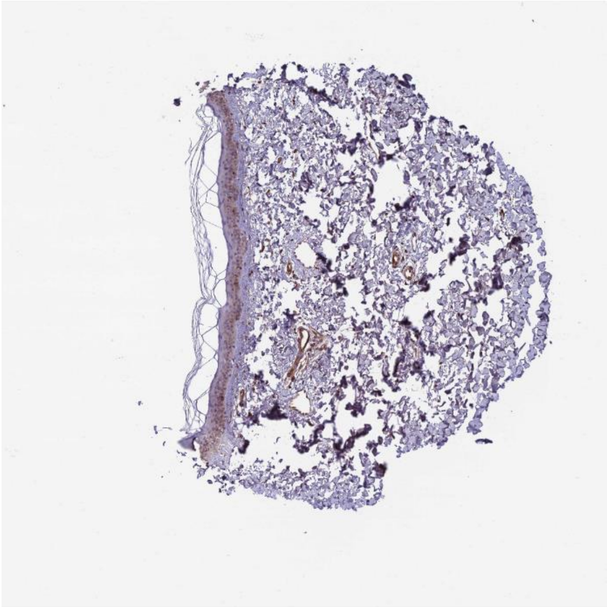

Calm1 Pde1b

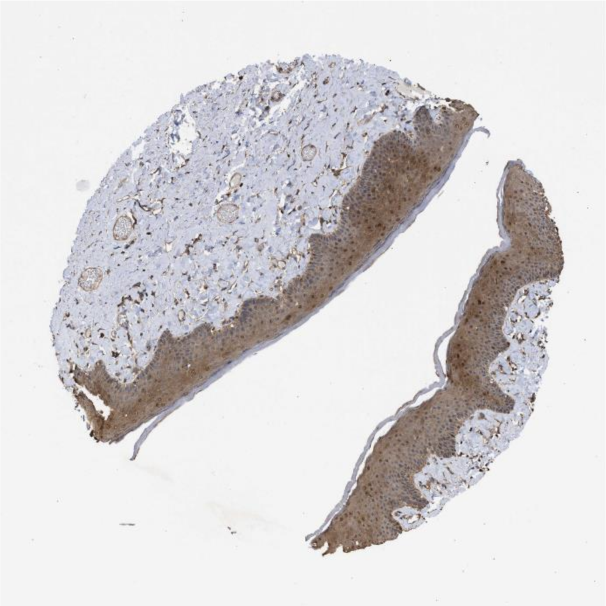
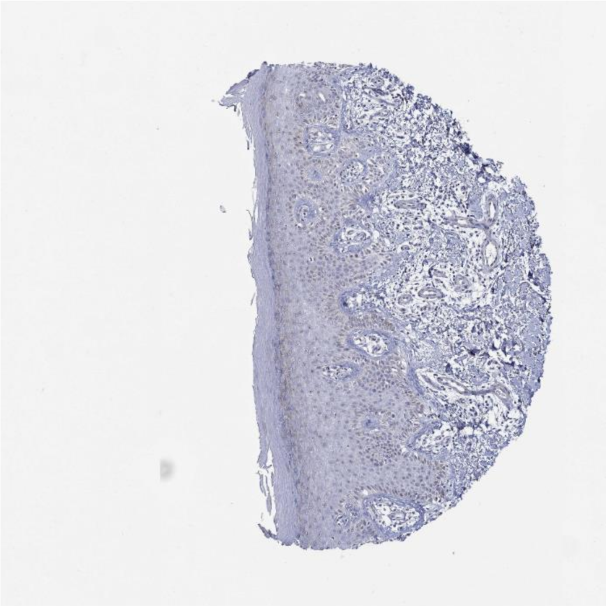

Lrpap1 Sort1

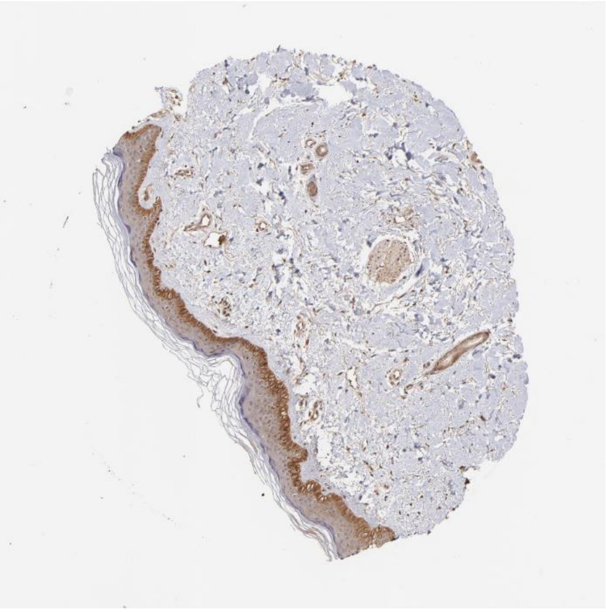

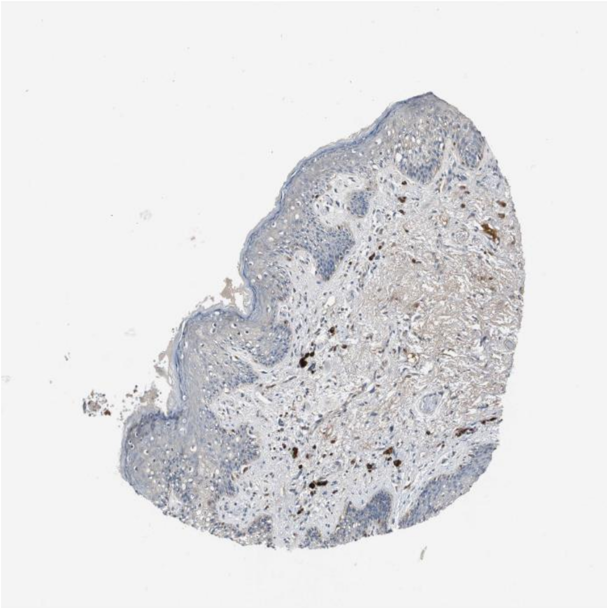

Timp2 Itgb1

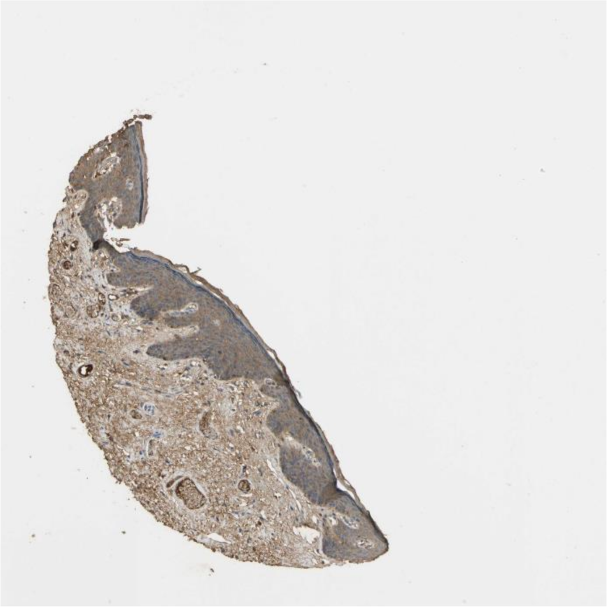

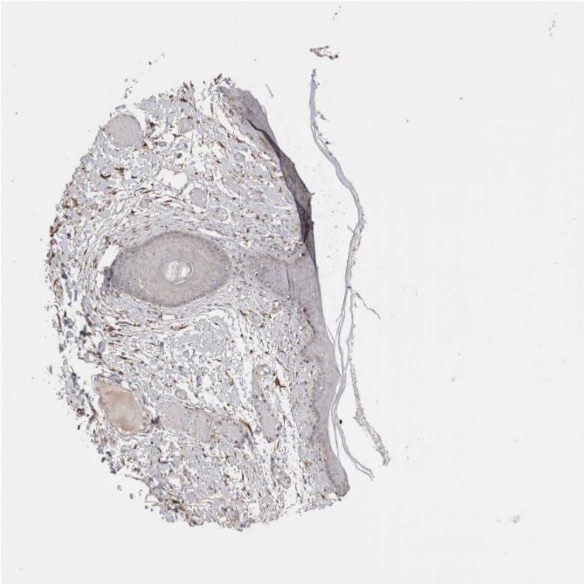

Efna4 Epha4

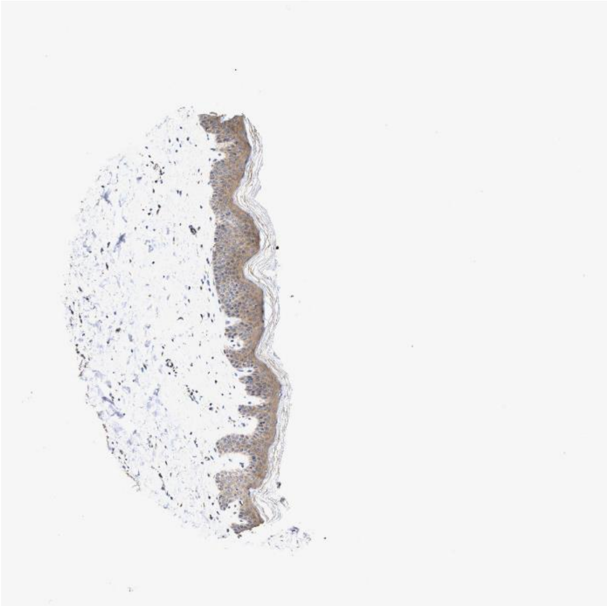 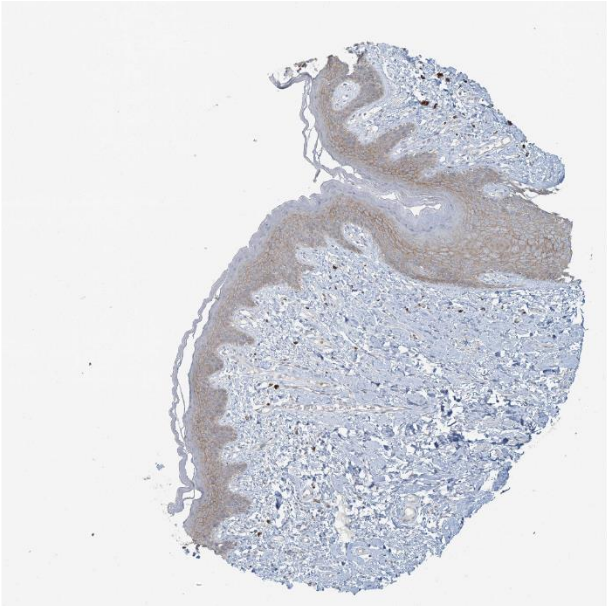

Efnb1 Ephb3

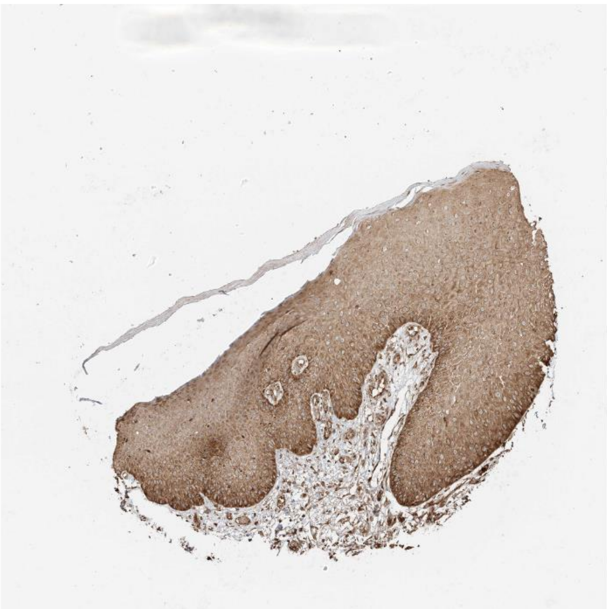 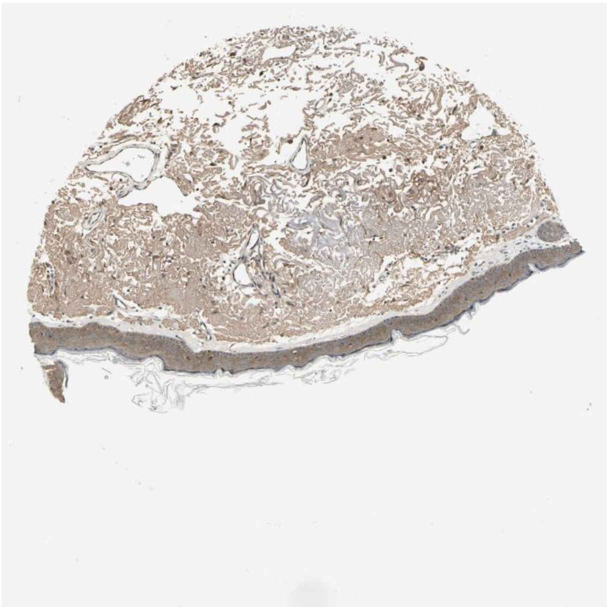

Psap Lrp1

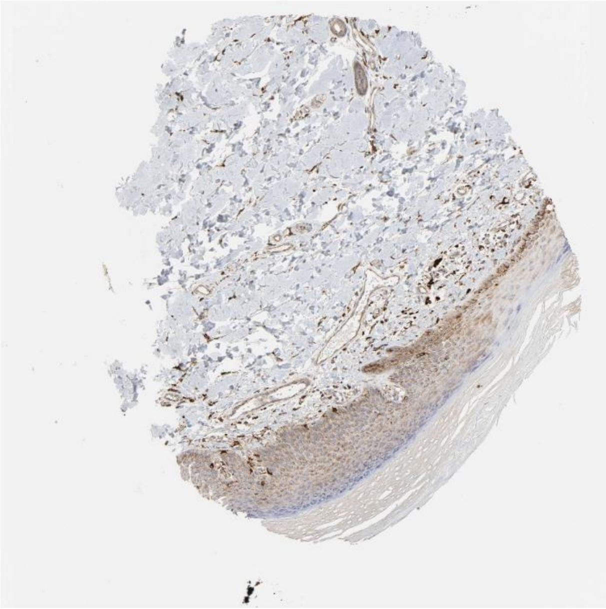

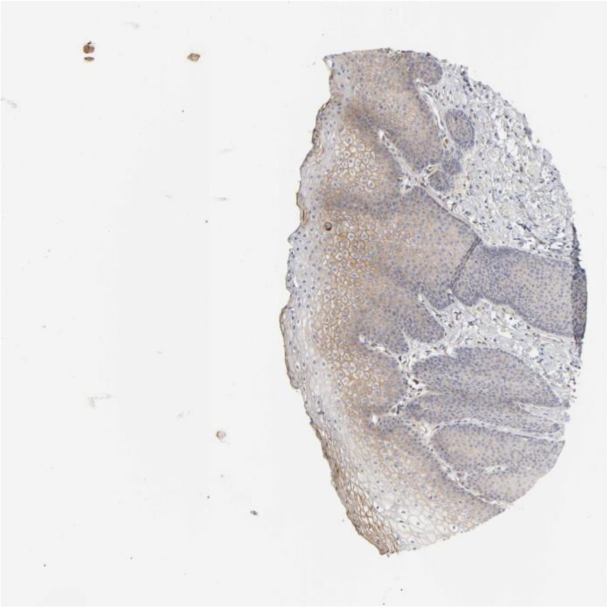

Tgfa Erbb3

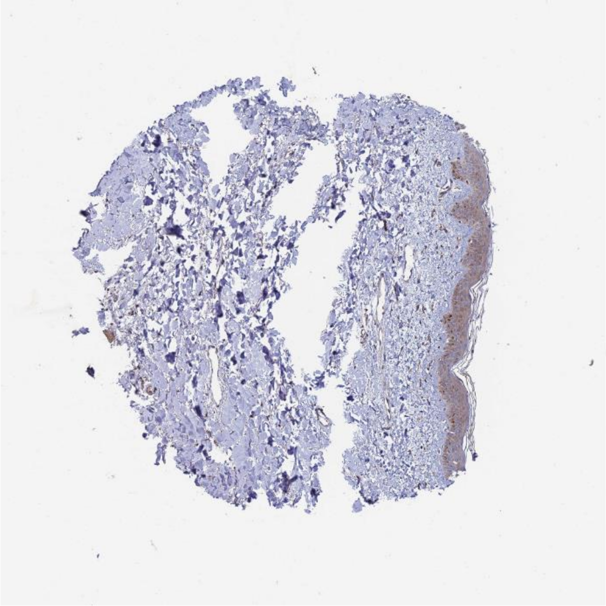

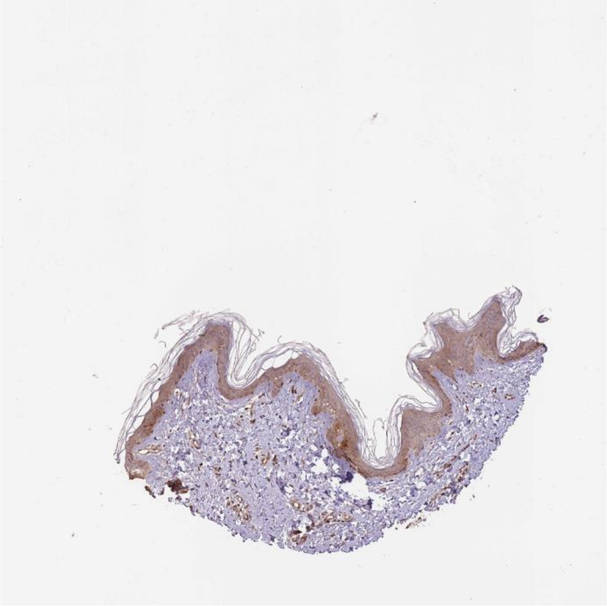

Calr Itga3

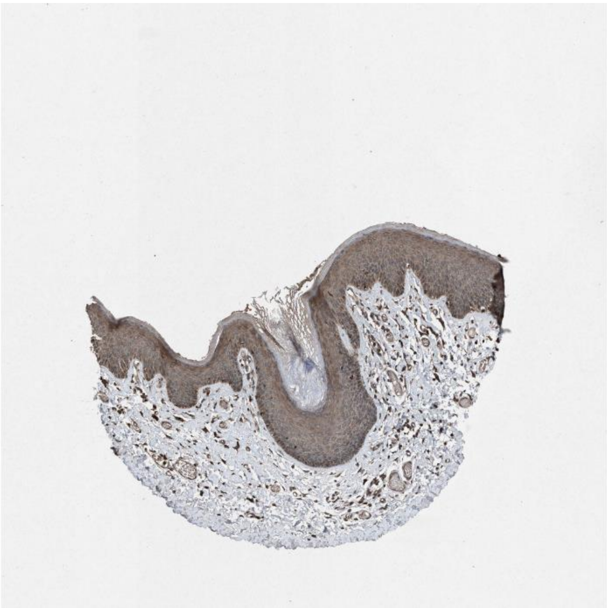

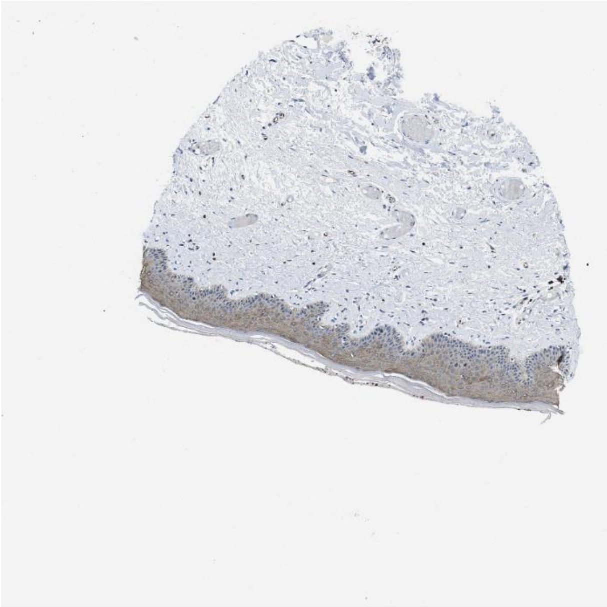

Adam17 Itgb1

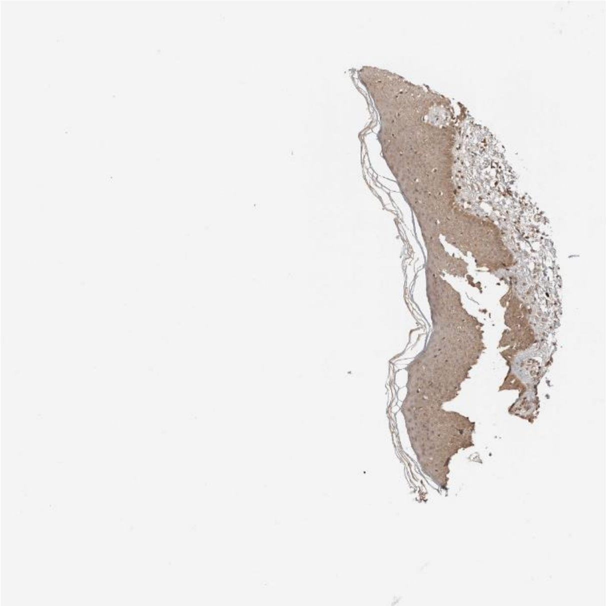

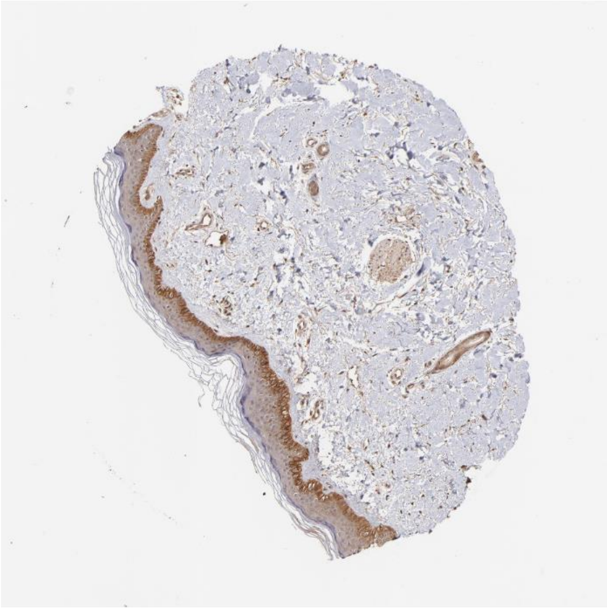

Tln1 Itgb5

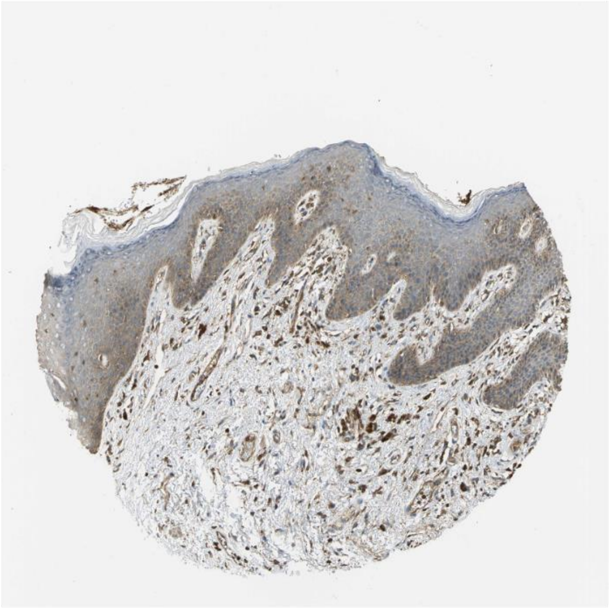

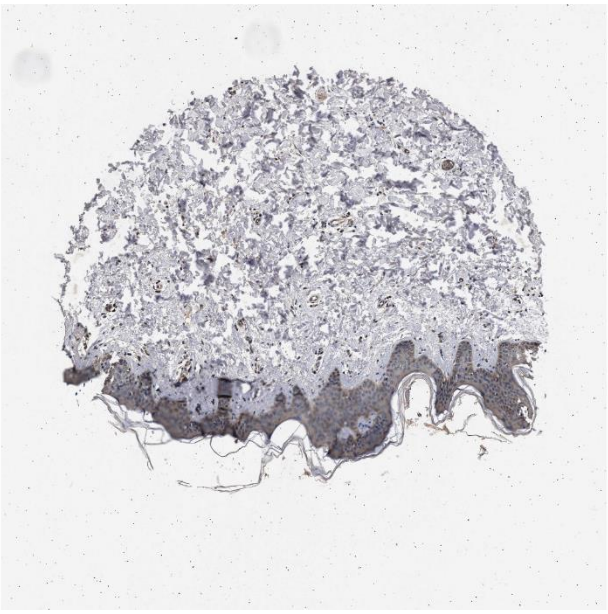

Calm2 Pde1b

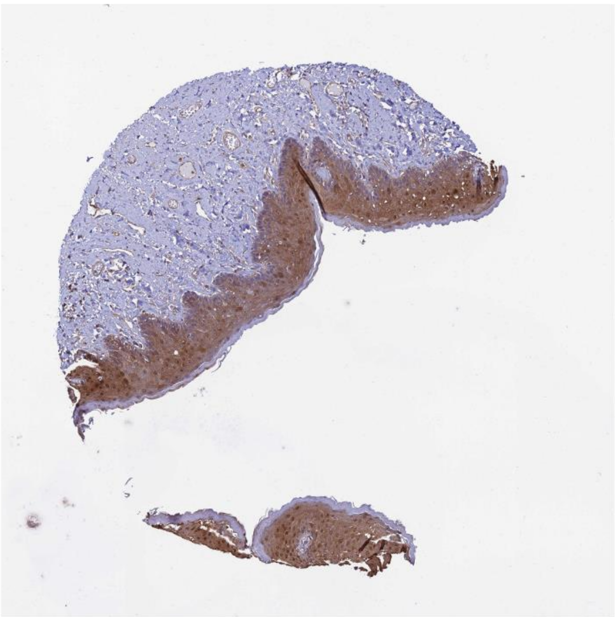

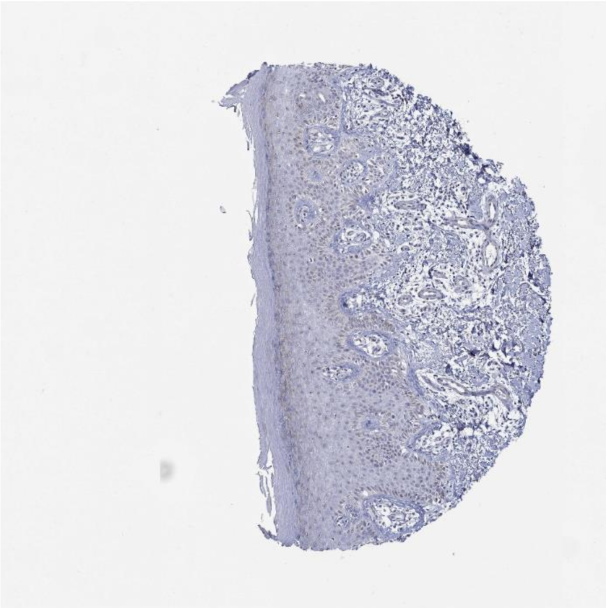

Pros1 Tyro3

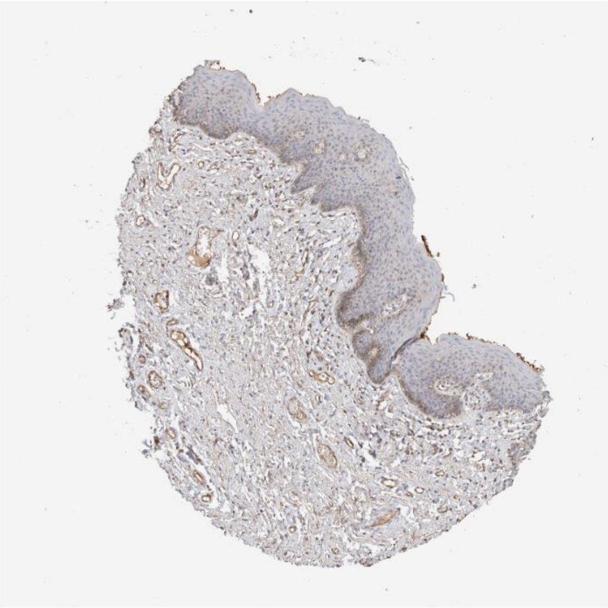

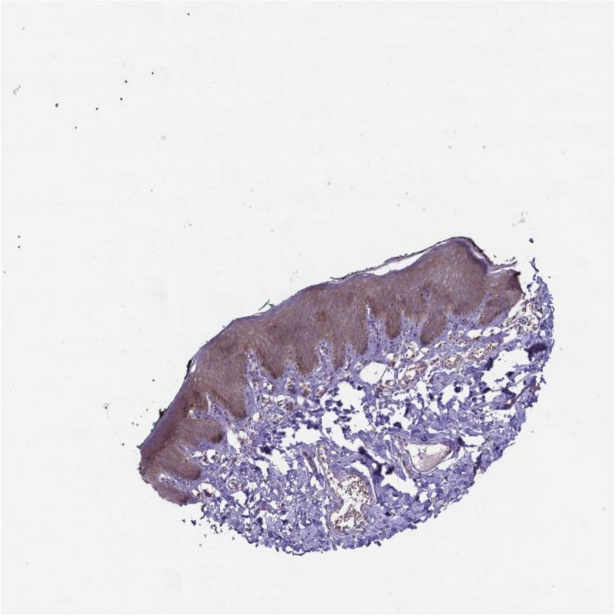

Dsc3 Dsg2

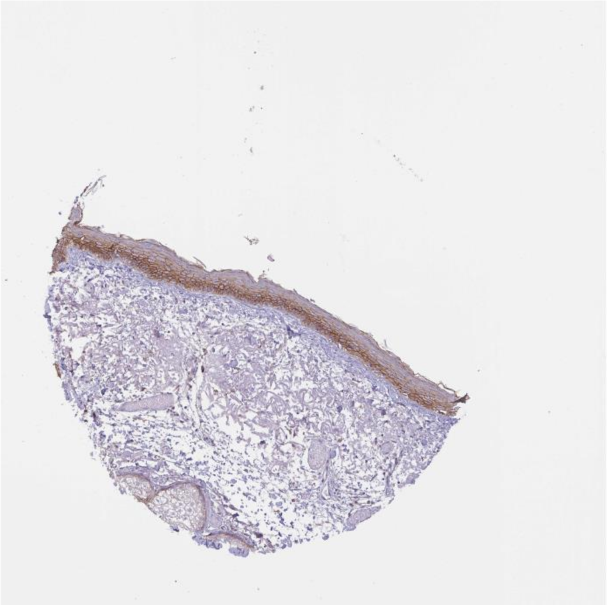

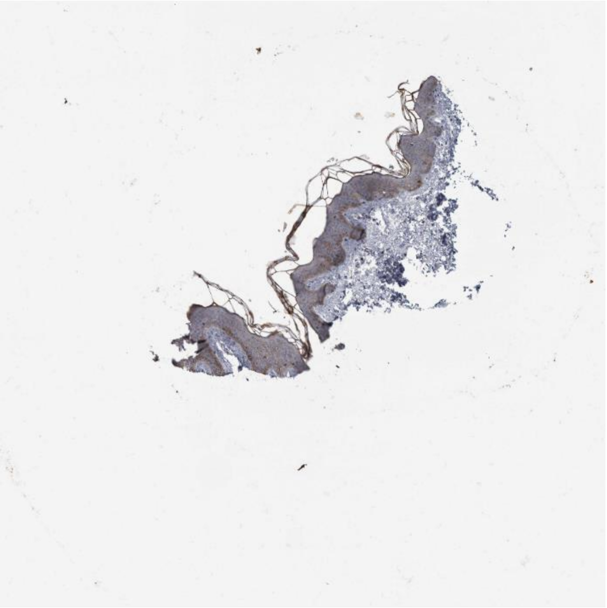

Dusp18 Itga3

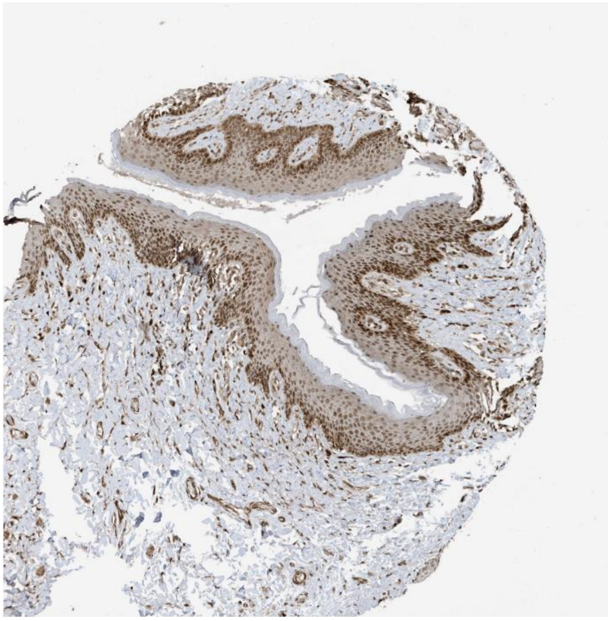

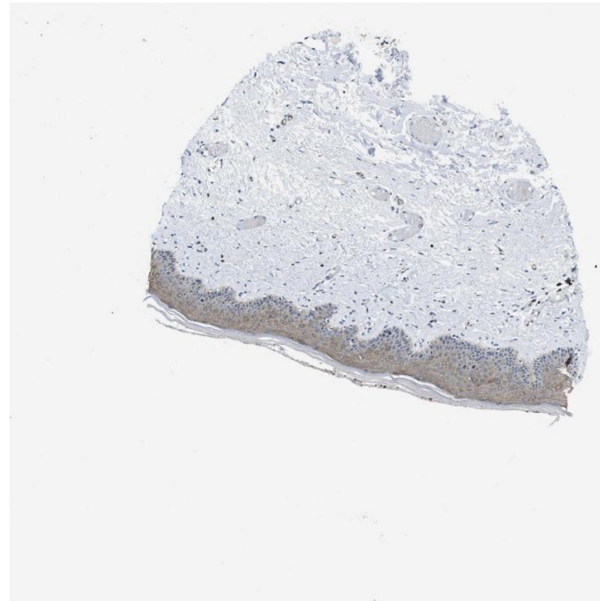

Adam10 Axl

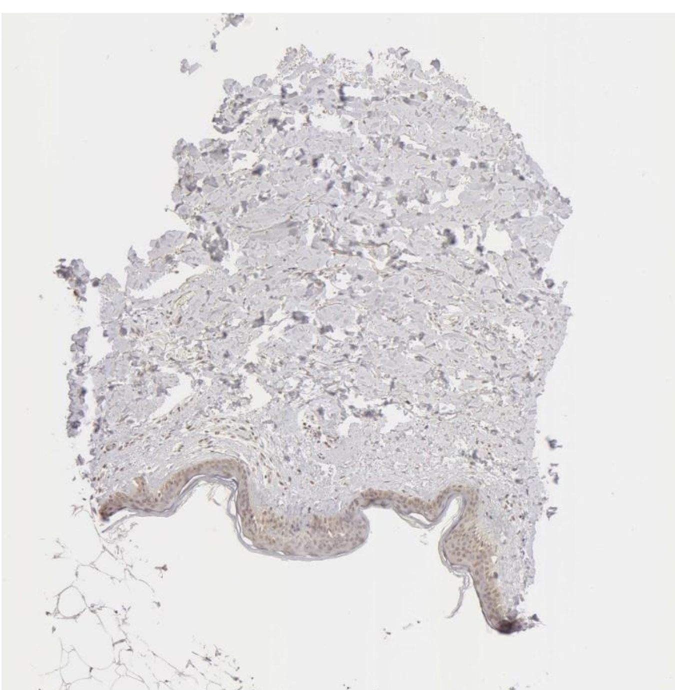

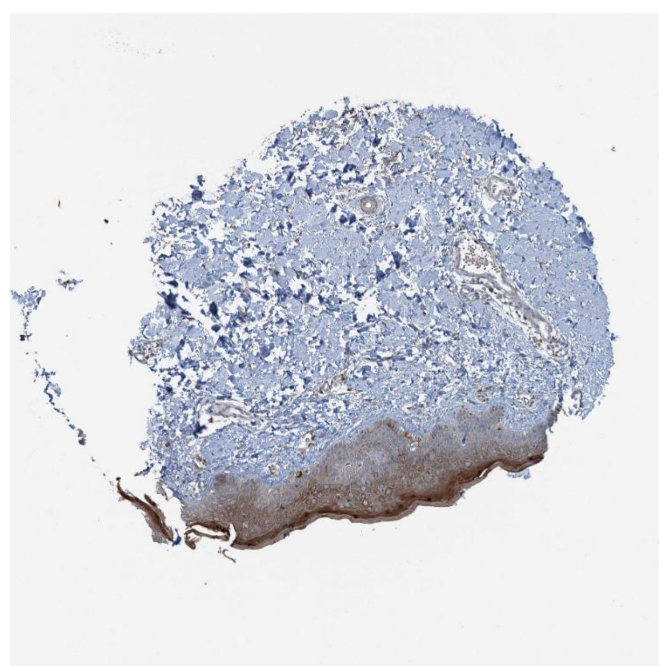

Arf1 Pld2

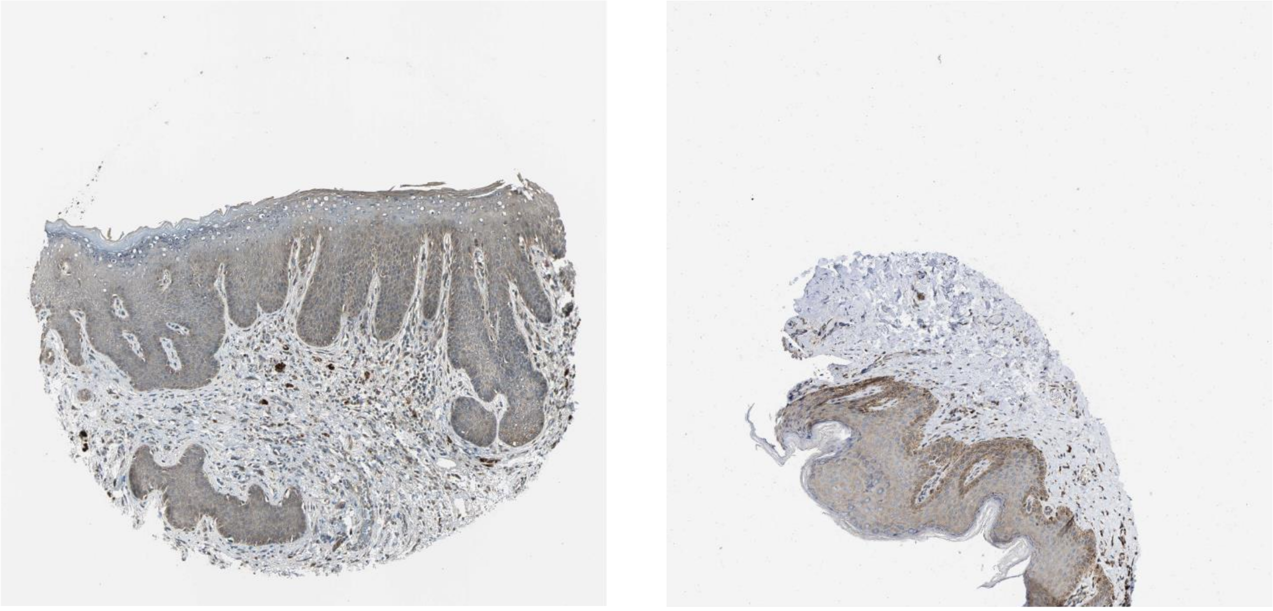

B2m Cd3d

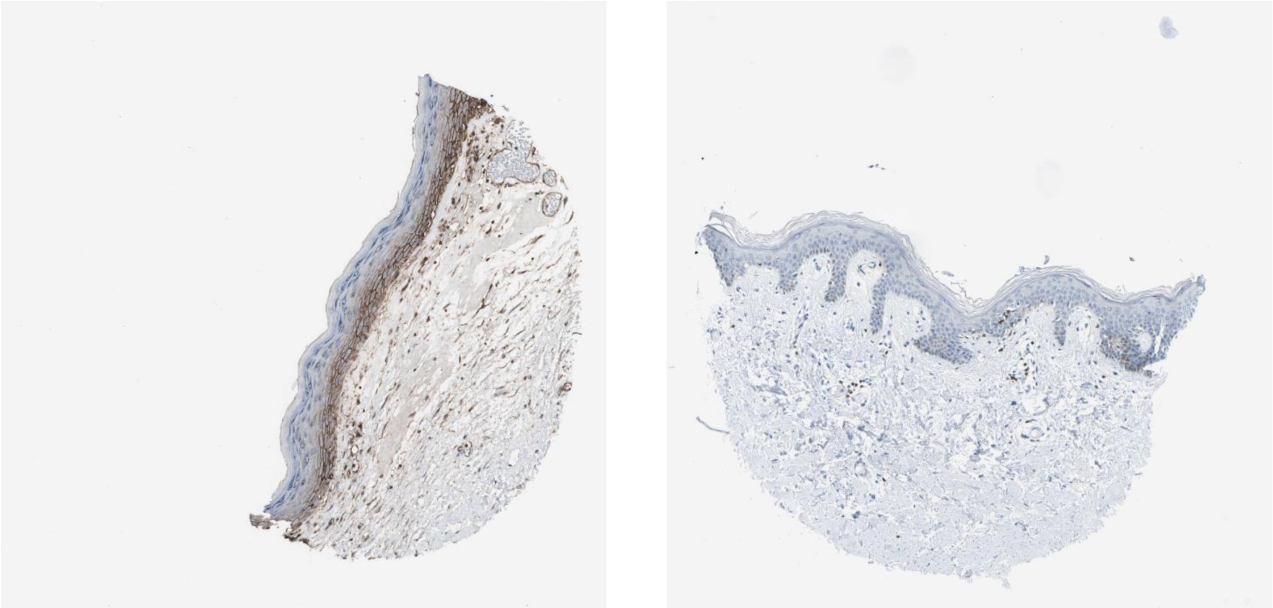

Adam15 Itgav

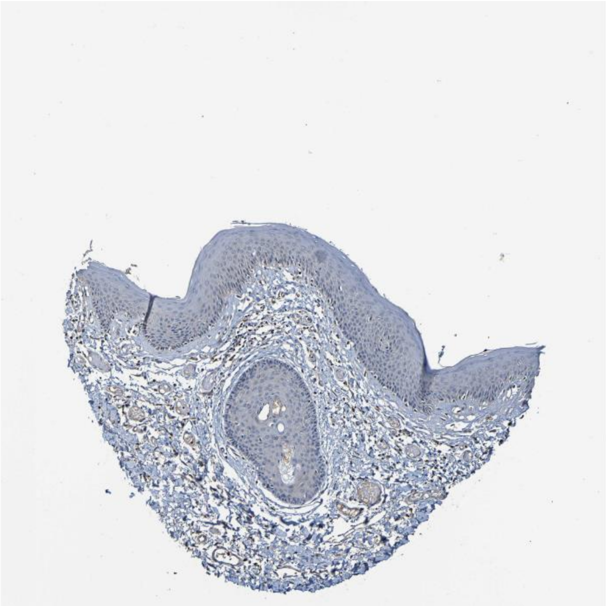

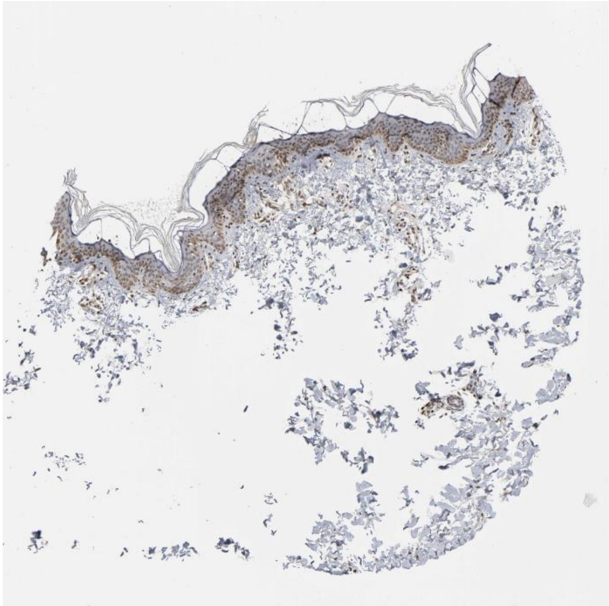

Calm1 Mylk

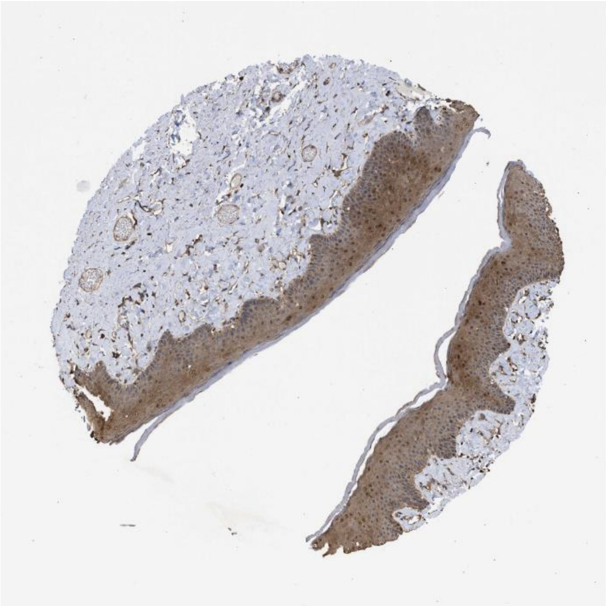

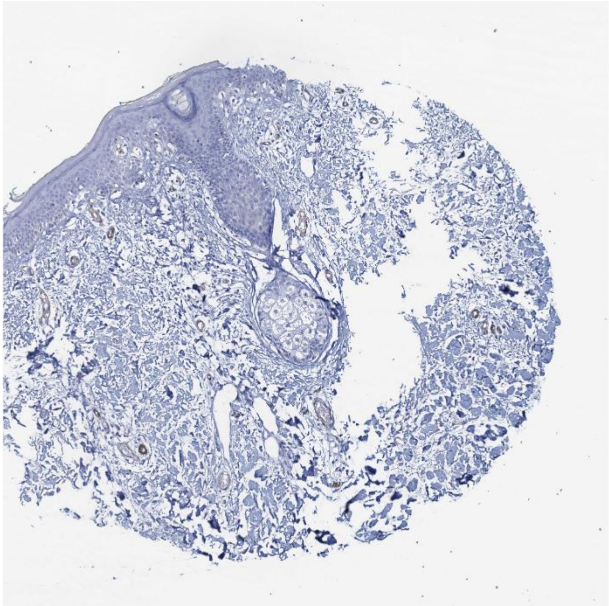

### Pdgfb Itgav

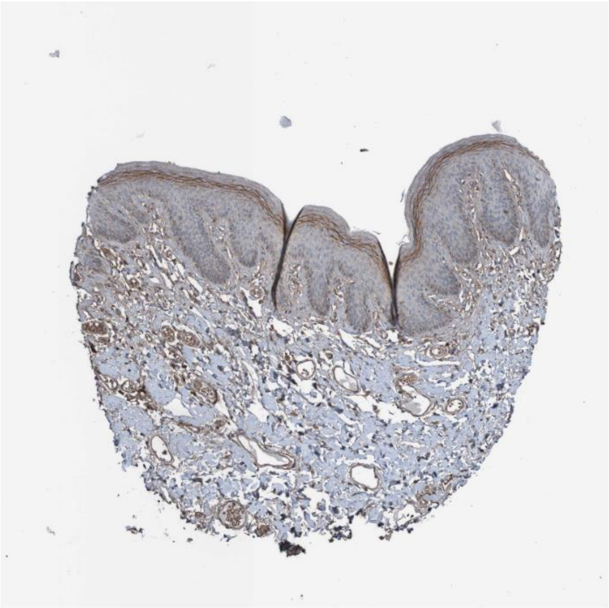

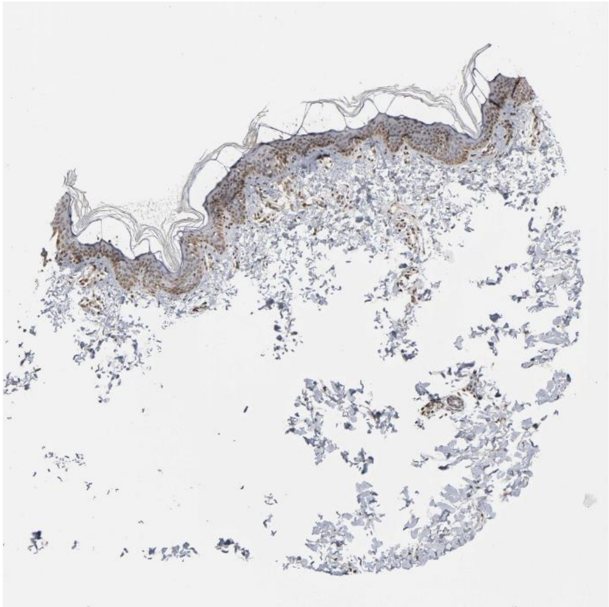

### Tnc Sdc1

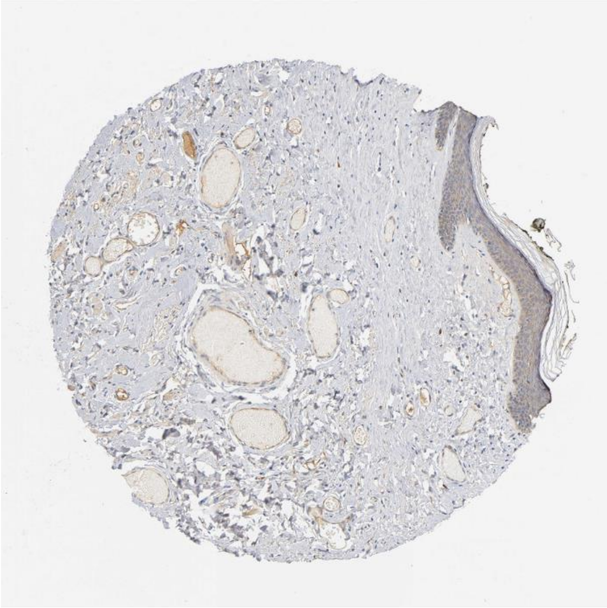

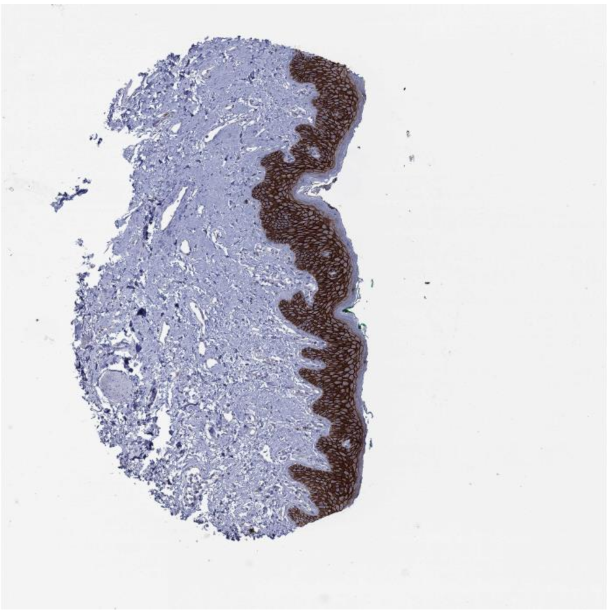

Lrpap1 Lrp1

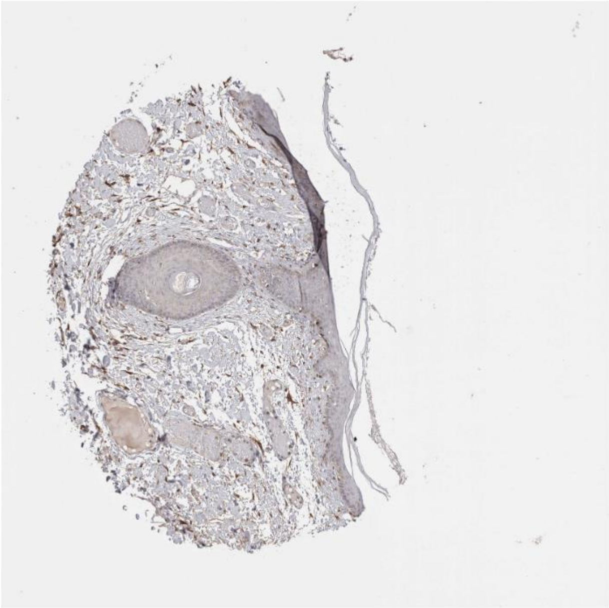

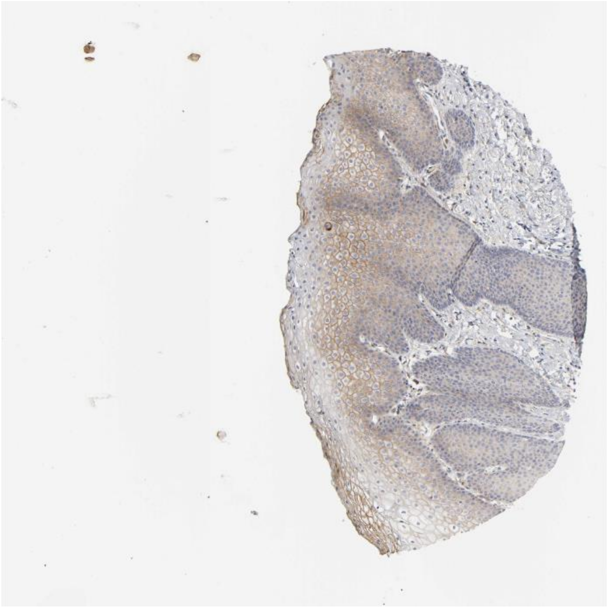

Dlk2 Notch1

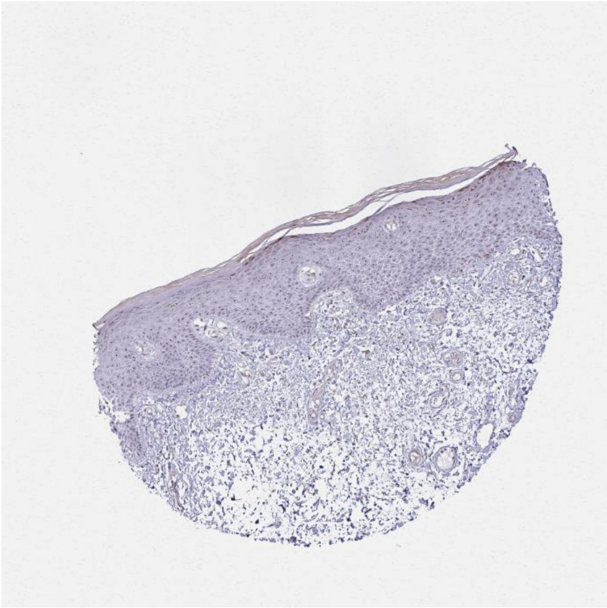

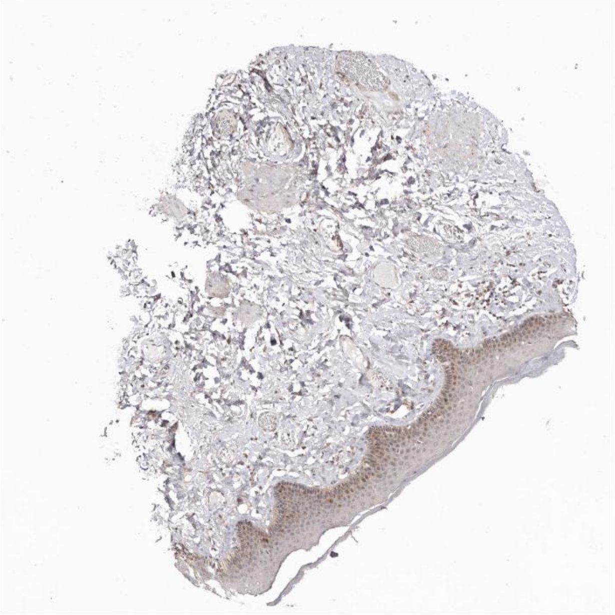

Itgb3bp Itgb5

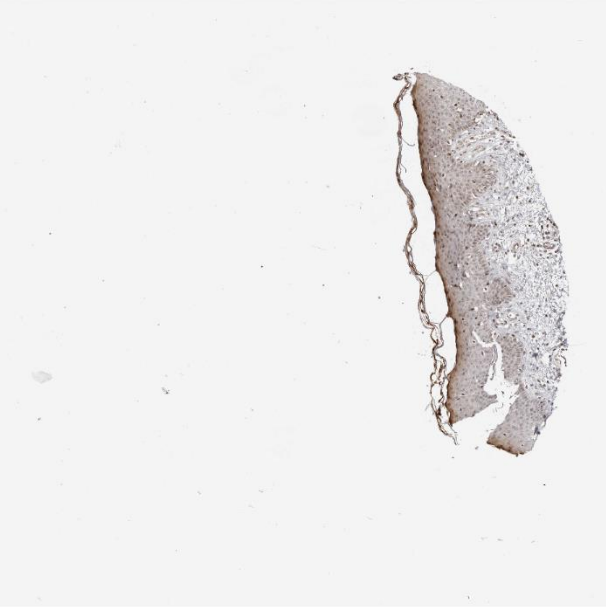
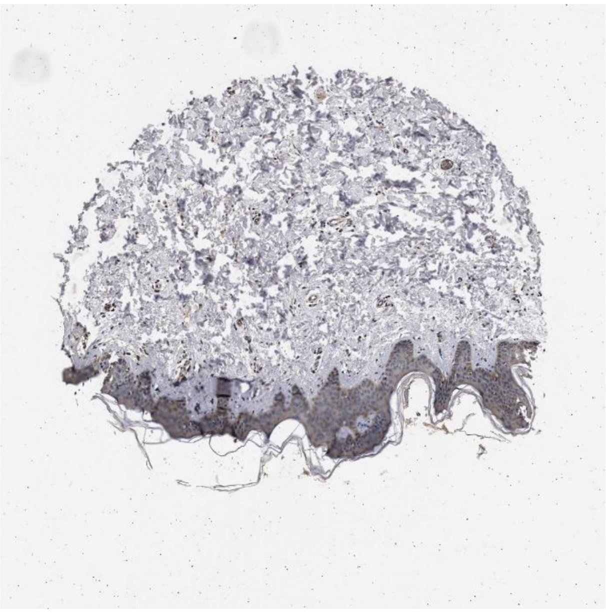

Rtn4 Gjb2

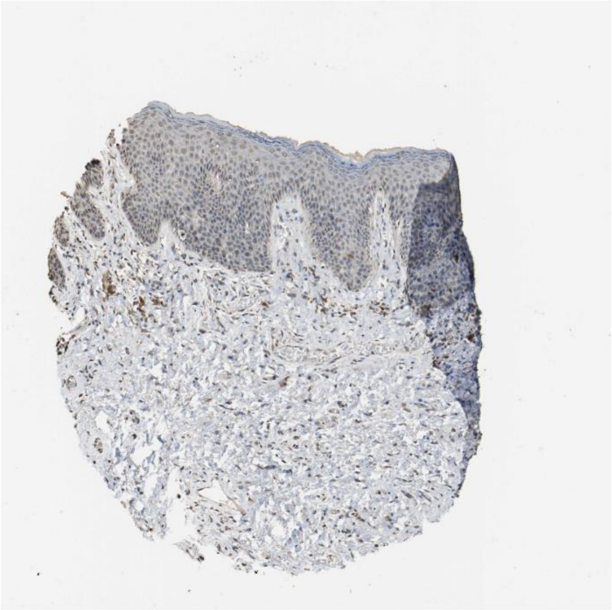
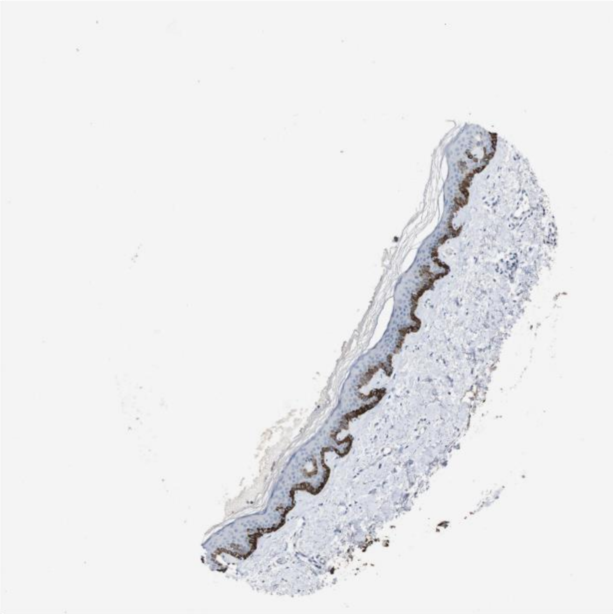

Lamb3 Cd151

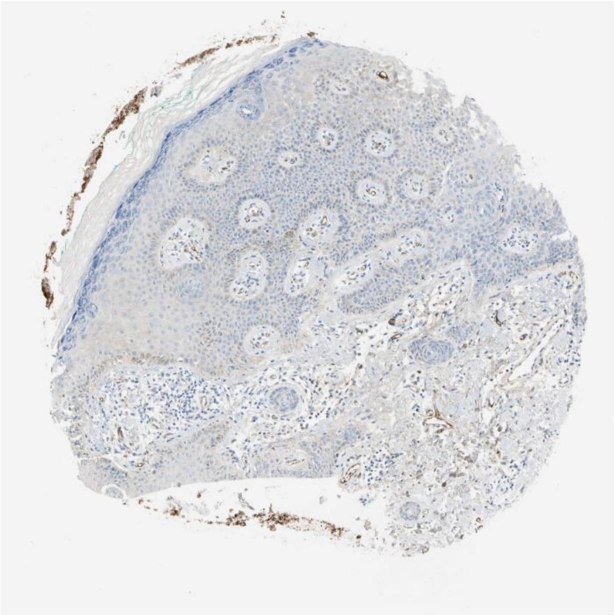

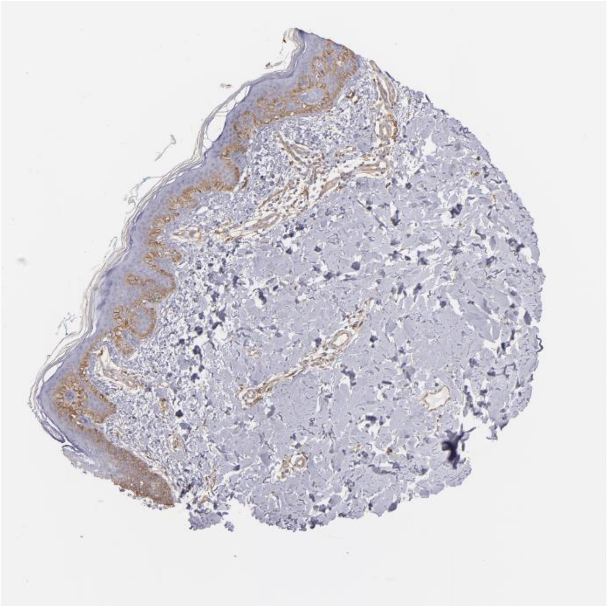

Lamb3 Itgb4

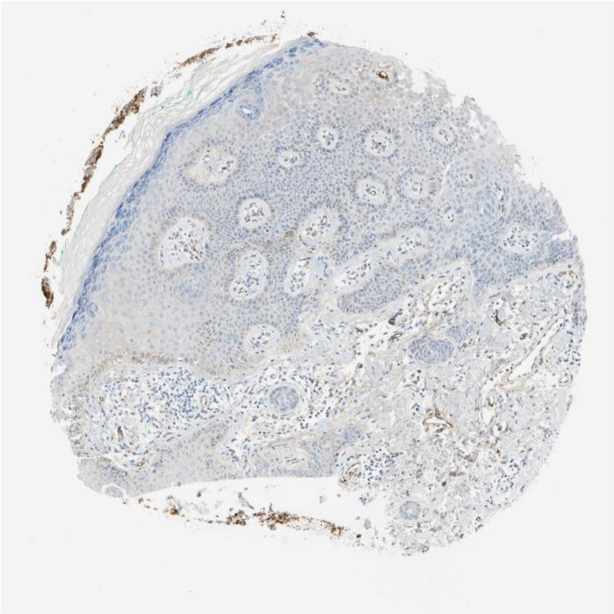

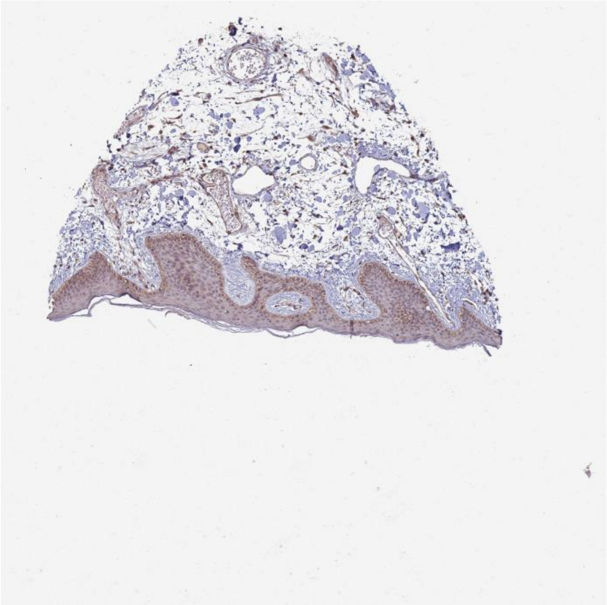

Agrn Lrp1

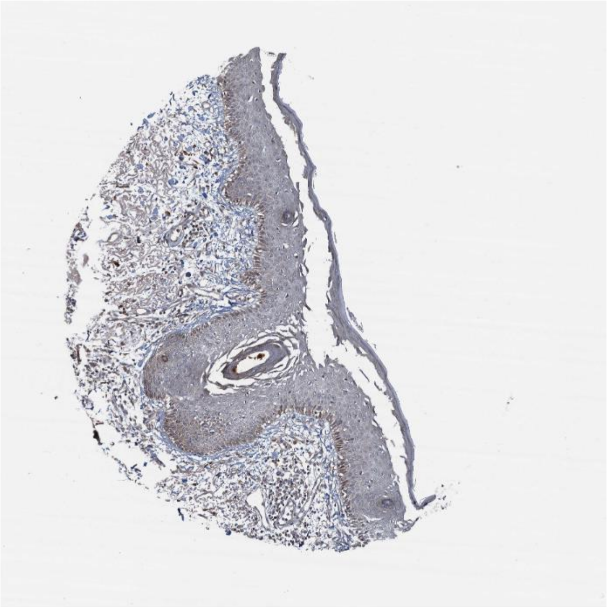

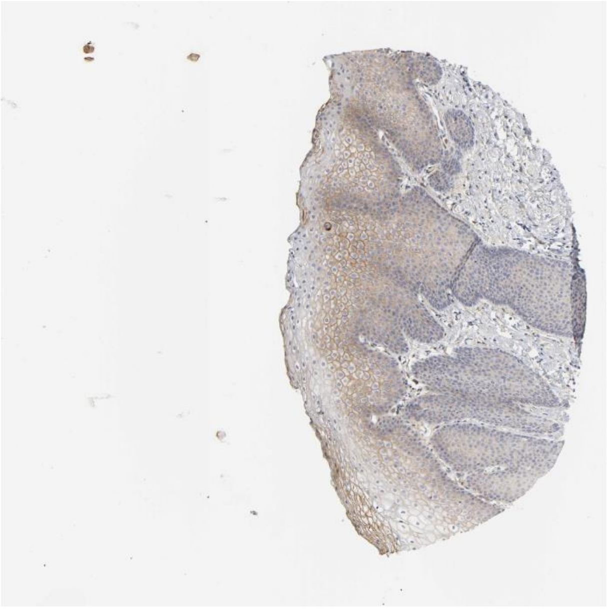

Vcl Itgb5

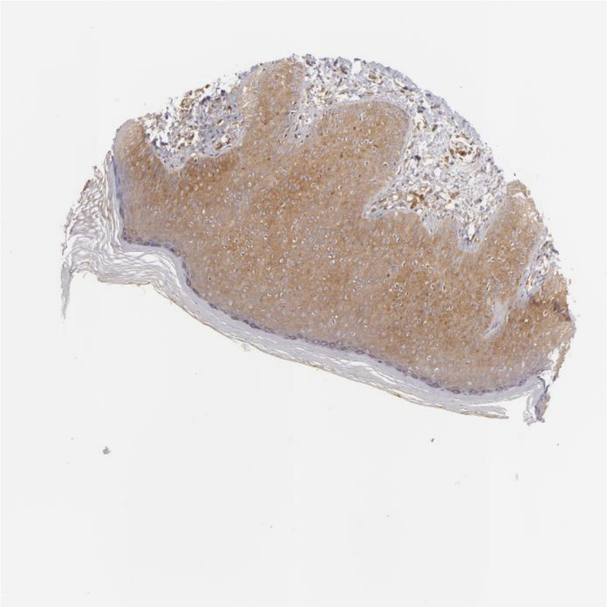

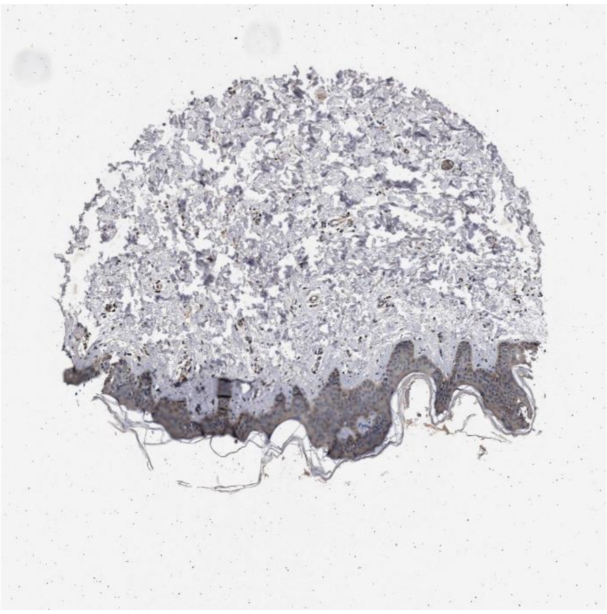

Gnas Adcy1

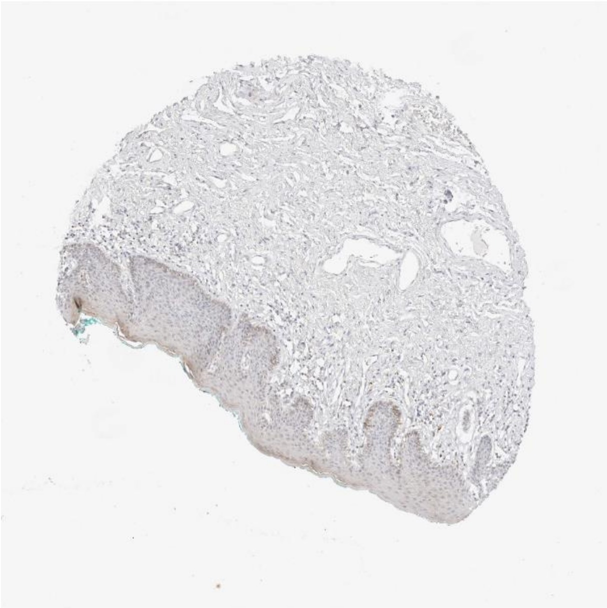
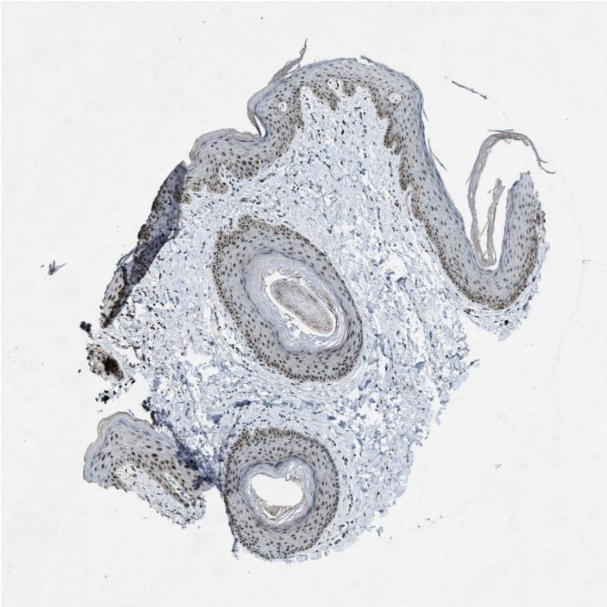

Calm2 Mylk

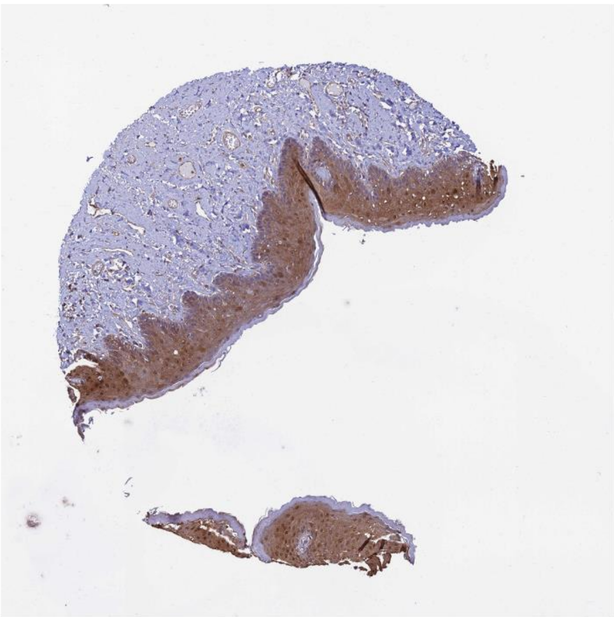
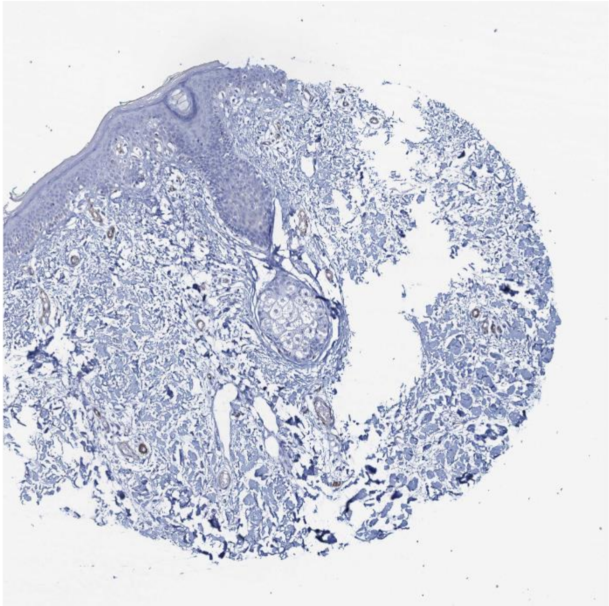

Calm3 Egfr

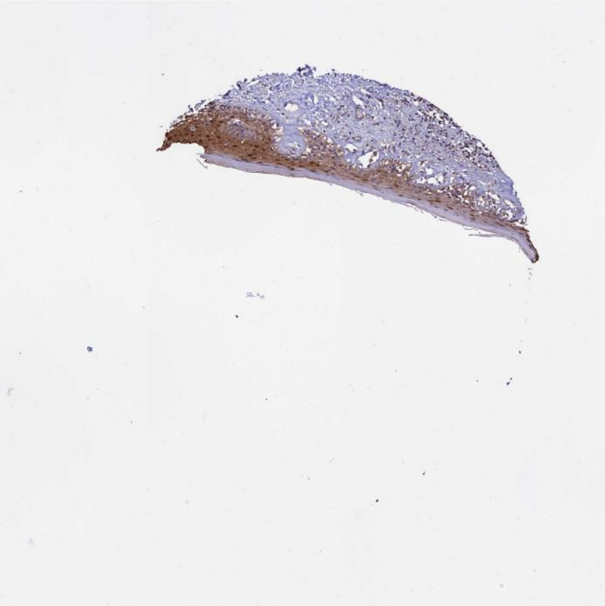

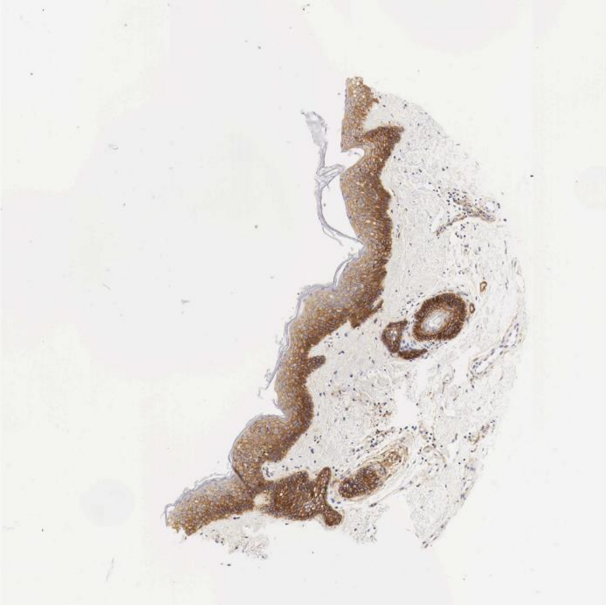

Dsc1 Dsg2

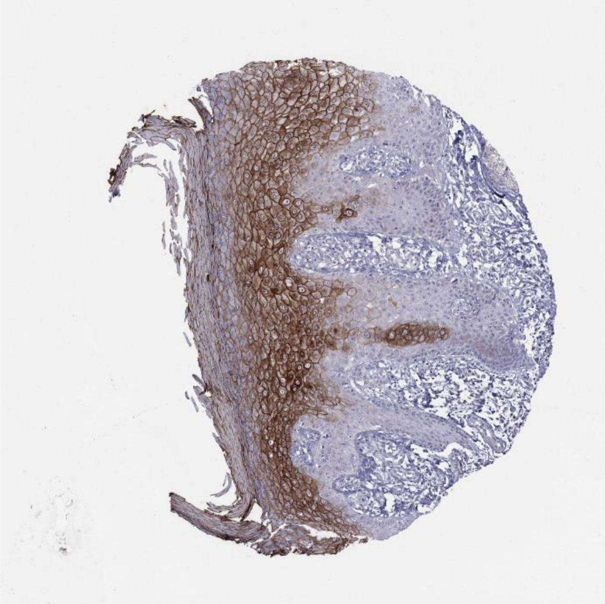

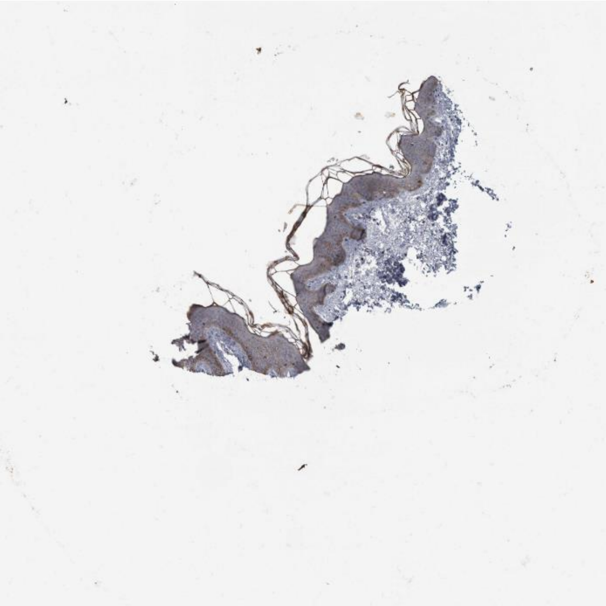

Sema6a Plxna2

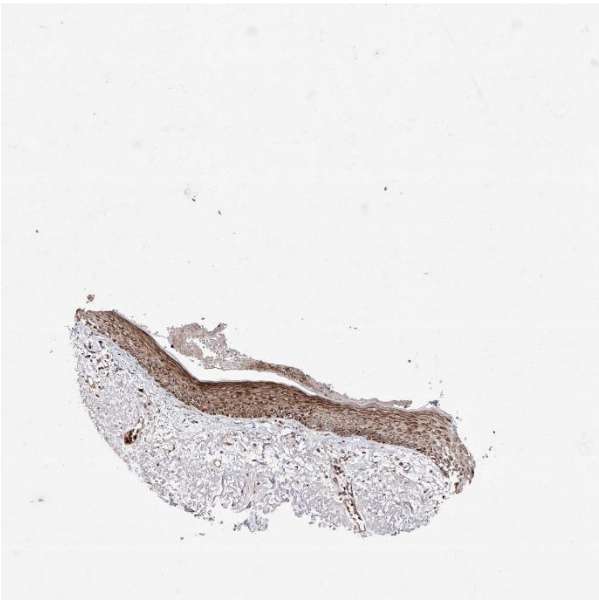

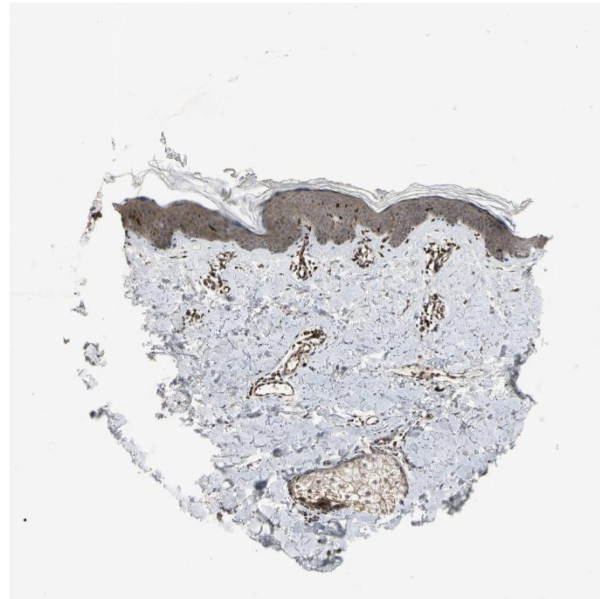

Efnb1 Ephb4

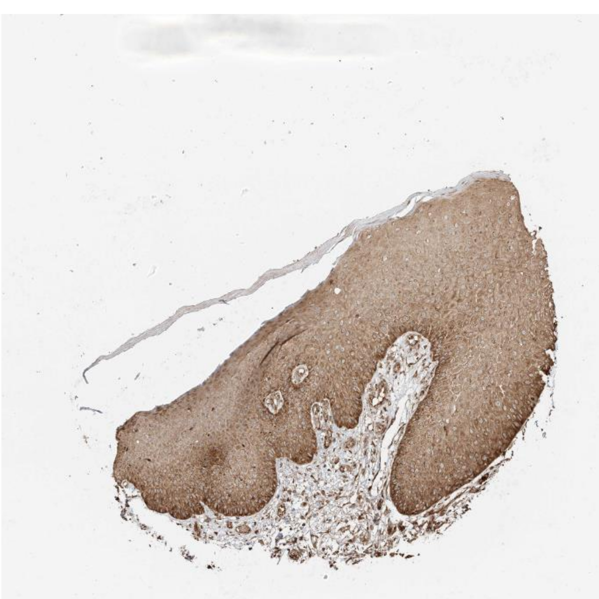

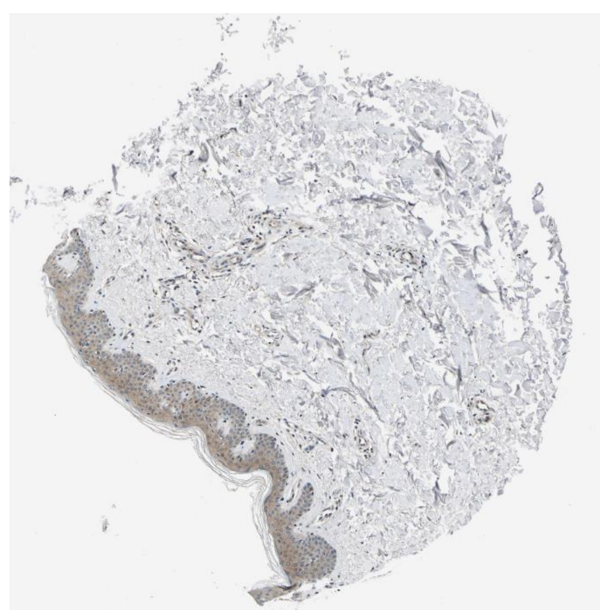

Ubc Smad3

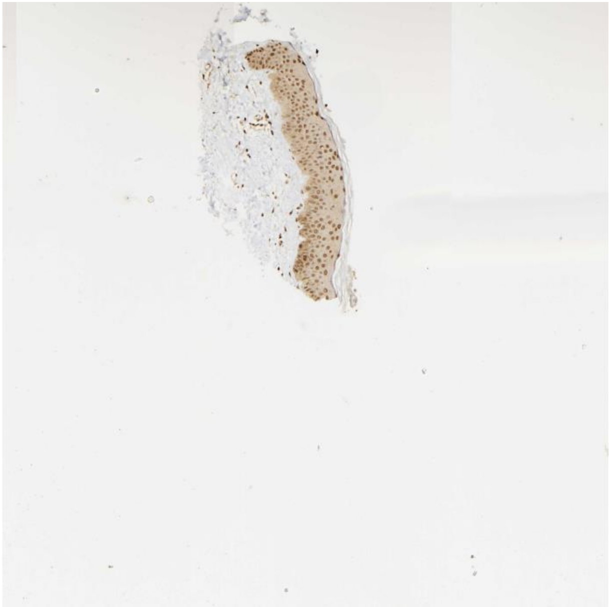

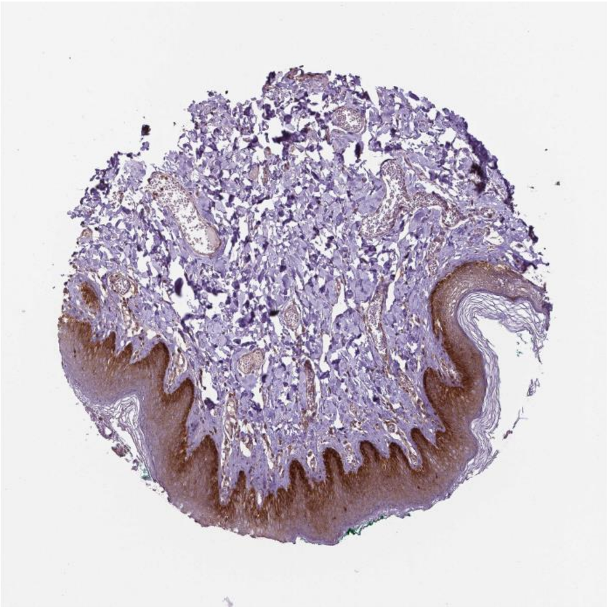

Gstp1 Traf2

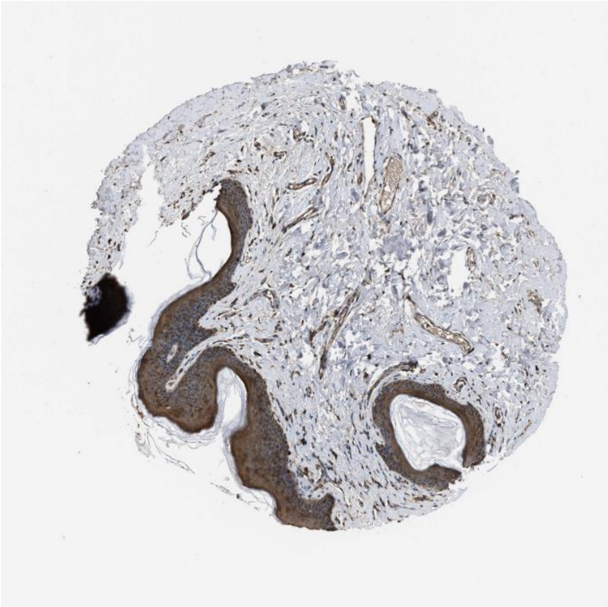

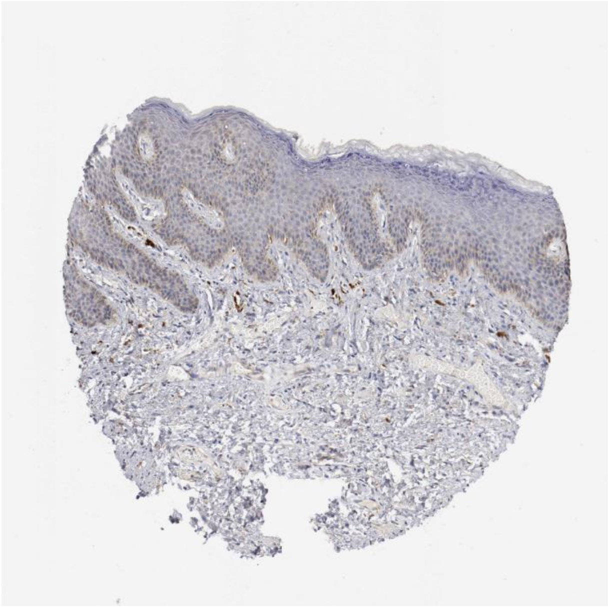

Areg Erbb3

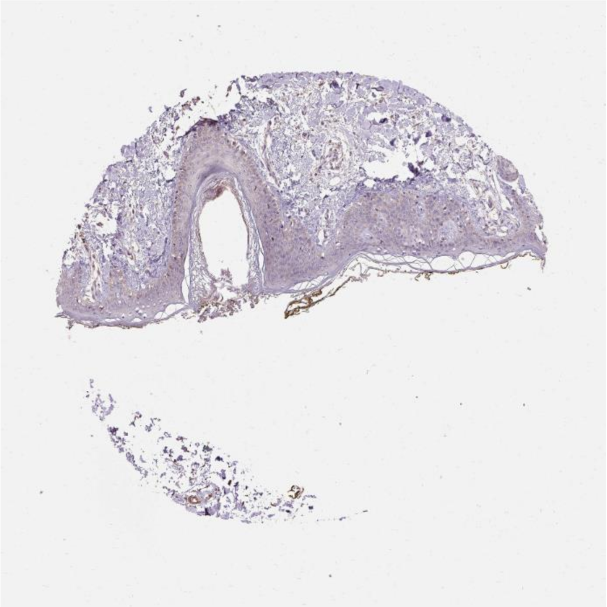

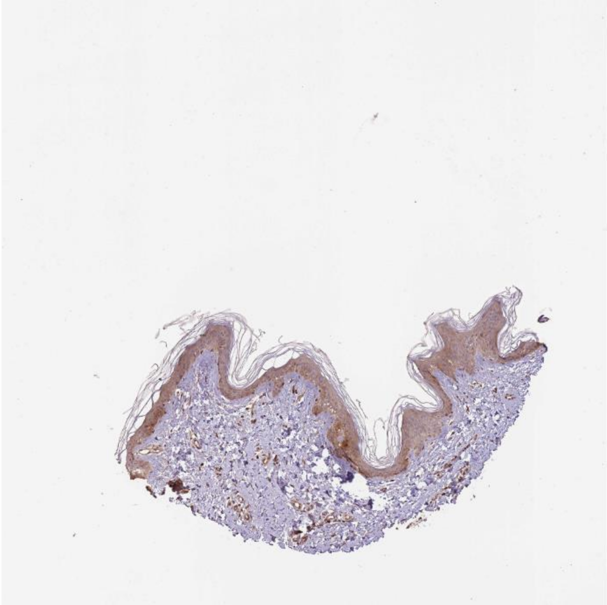

Calm1 Hmmr

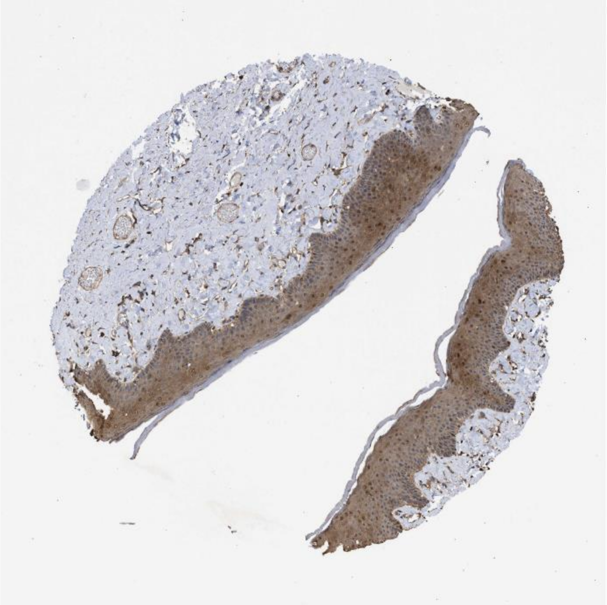

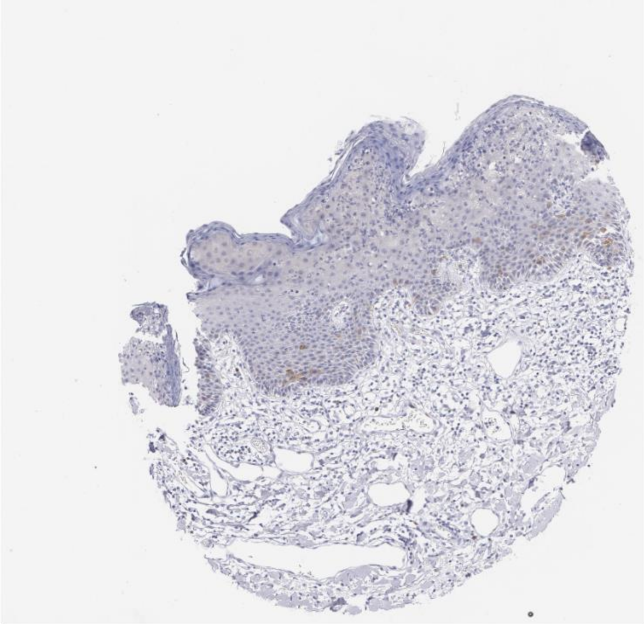

Col18a1 Itgb5

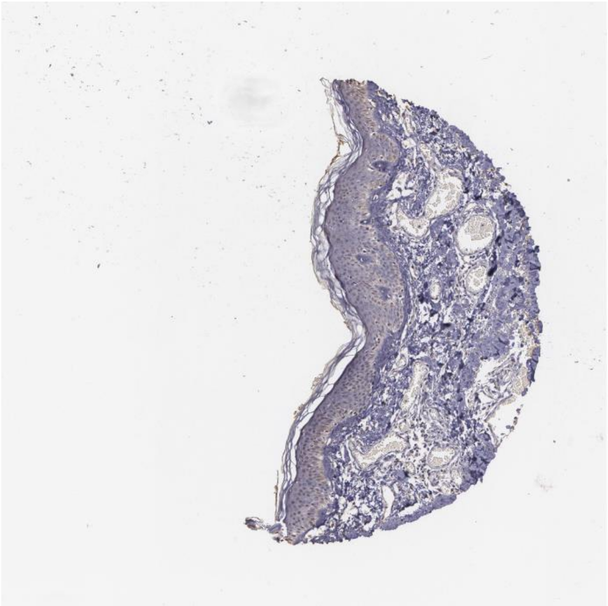 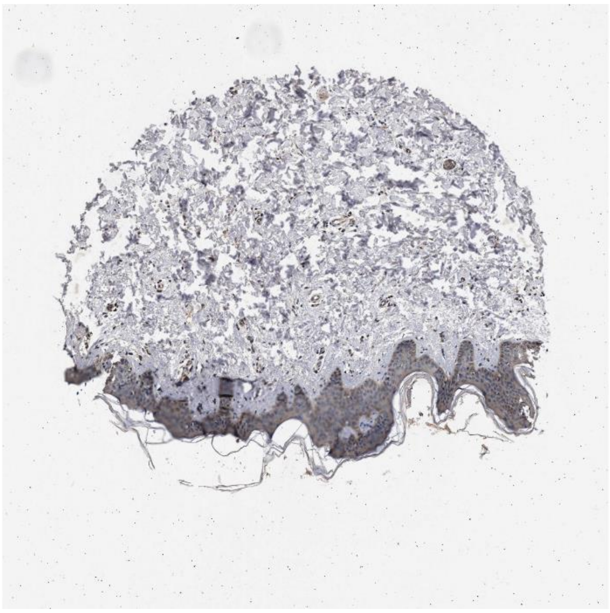

Il18 Il1rl2

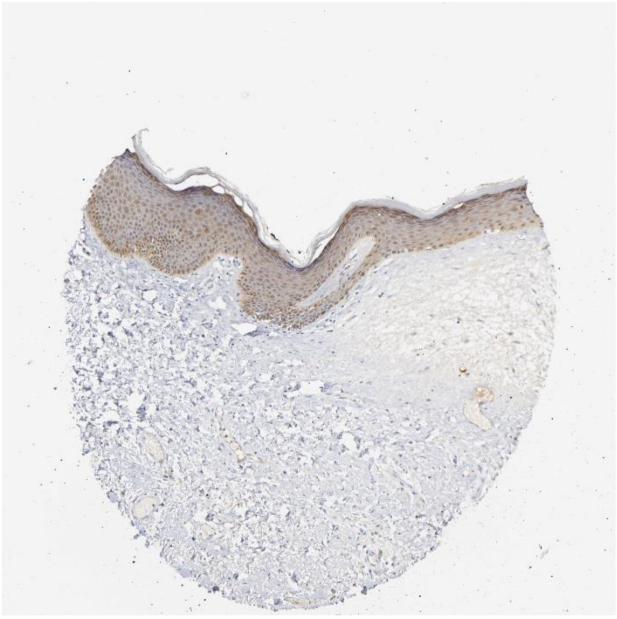 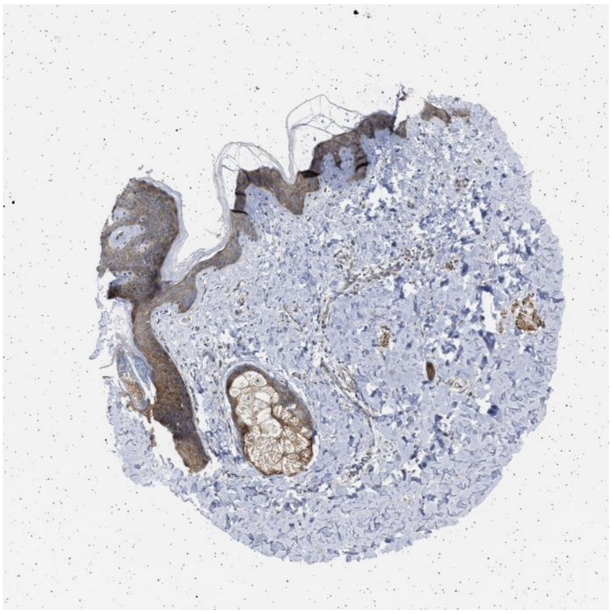

B2m Klrd1

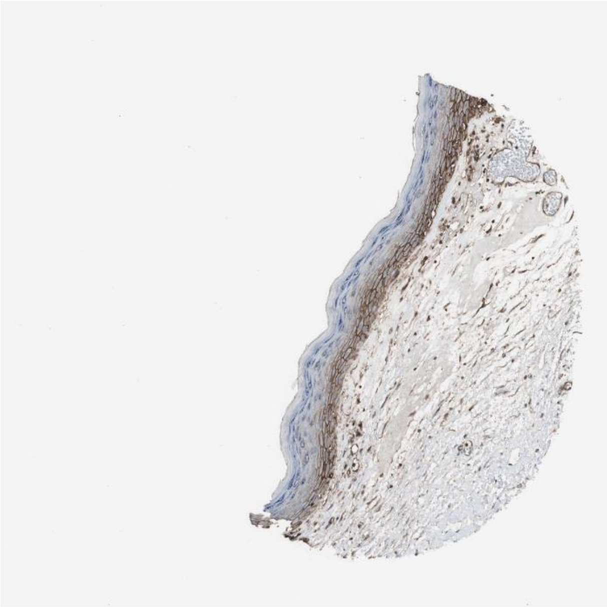

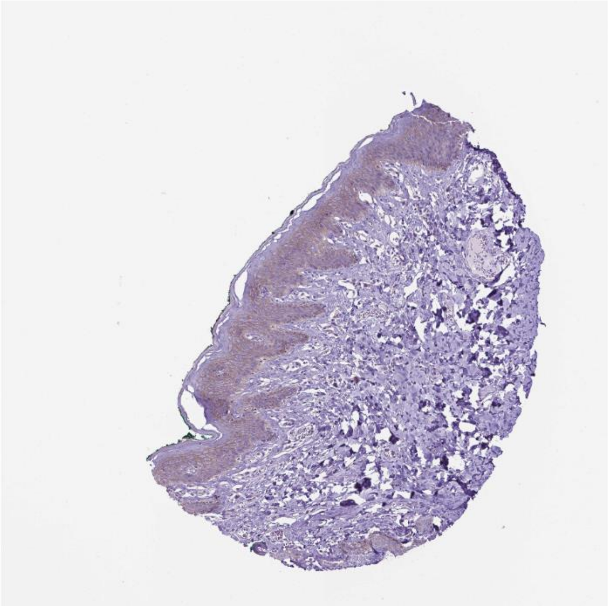

Thbs1 Itga6

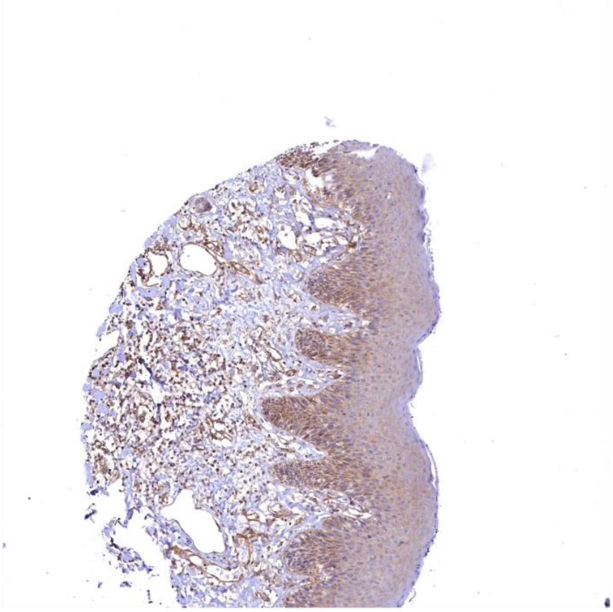

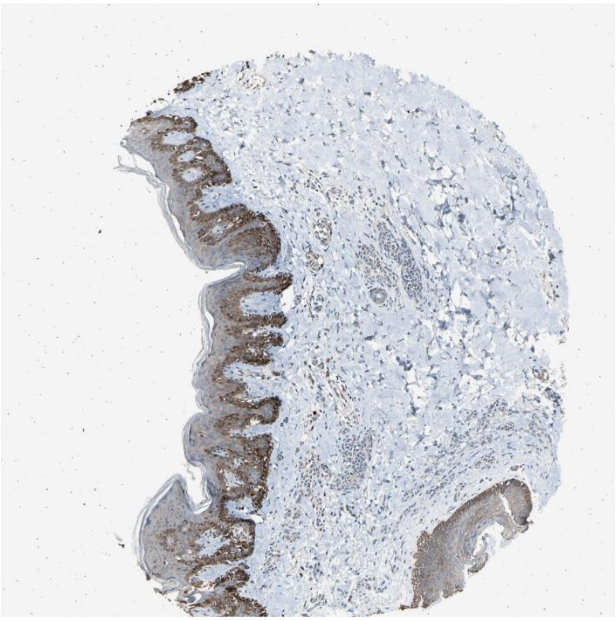

Liph Lpar2

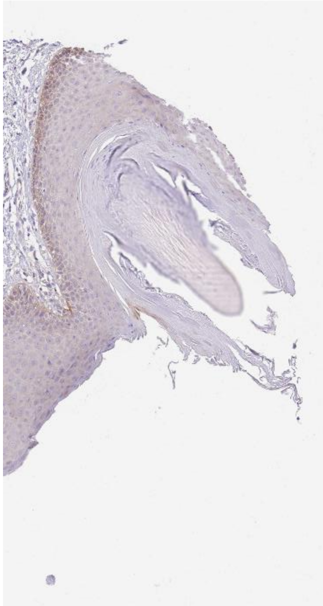

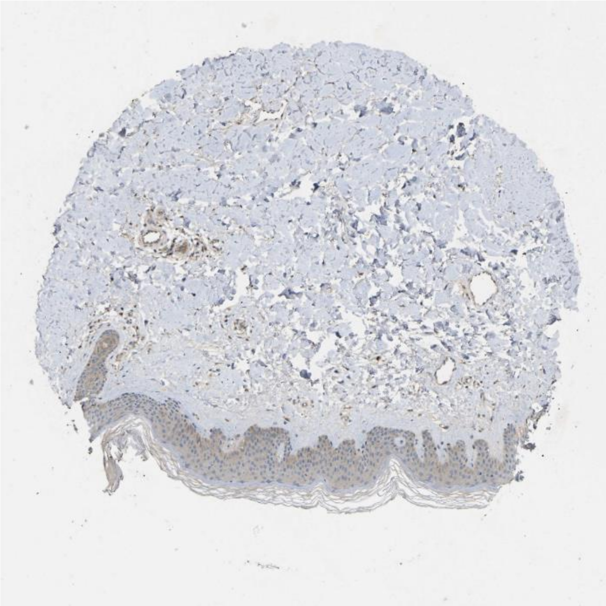

Sfrp1 Fzd6

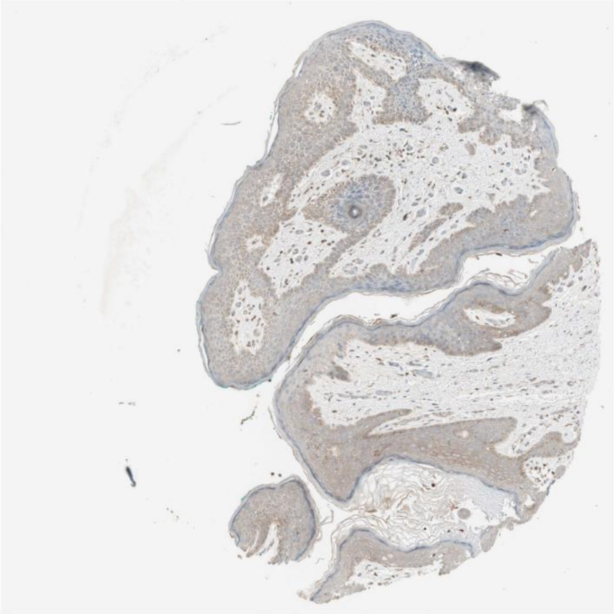

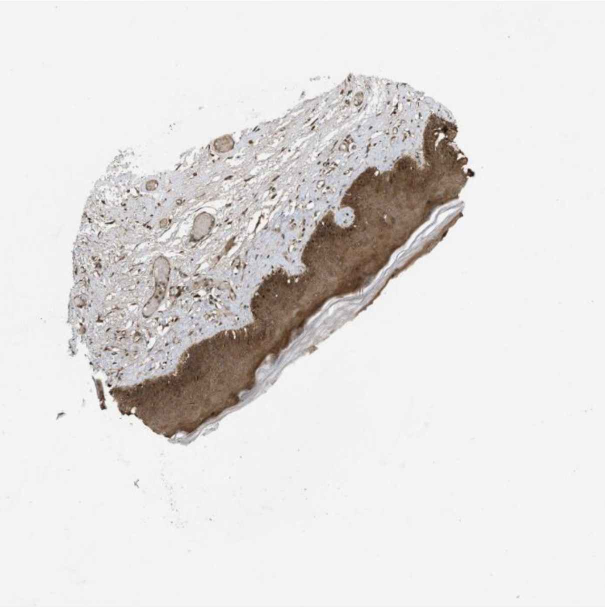

Pdgfb Lrp1

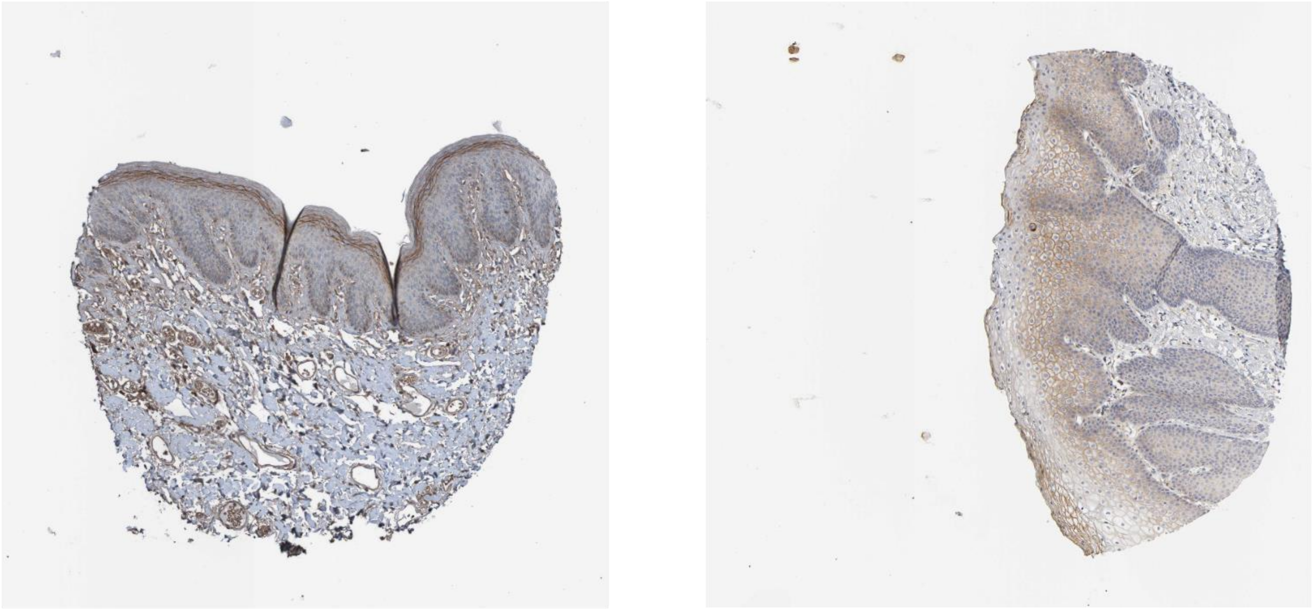

Timp2 Itga3

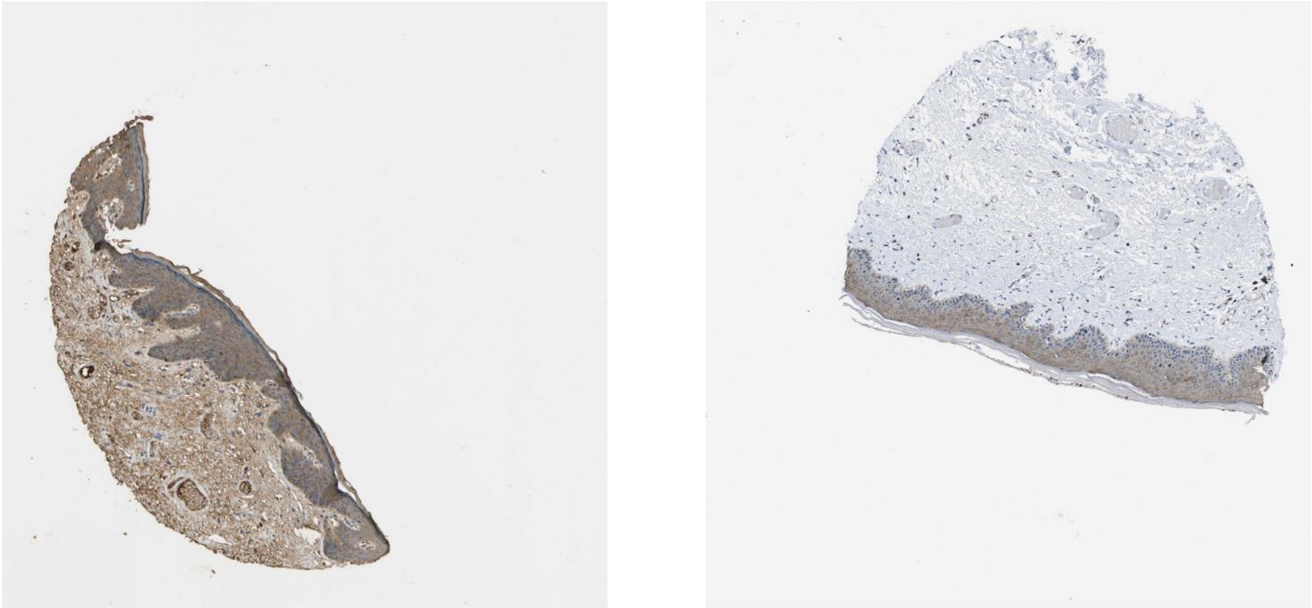

Thbs1 Cd47

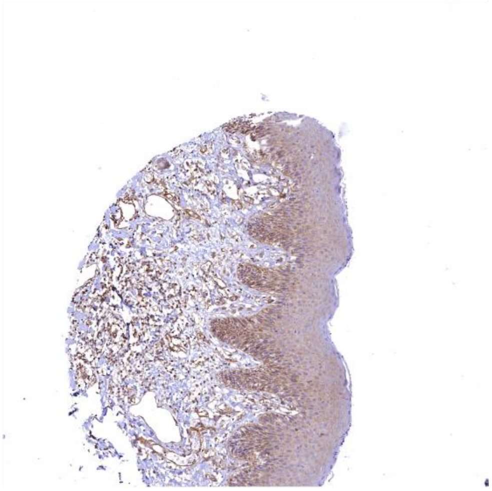 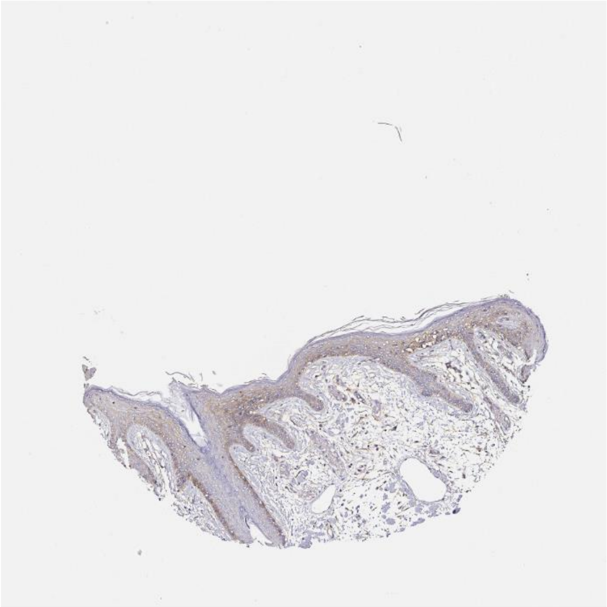

Lama5 Itga6

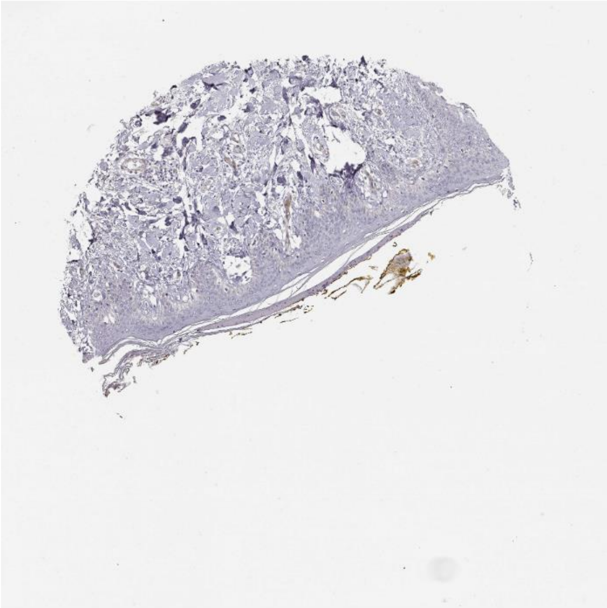 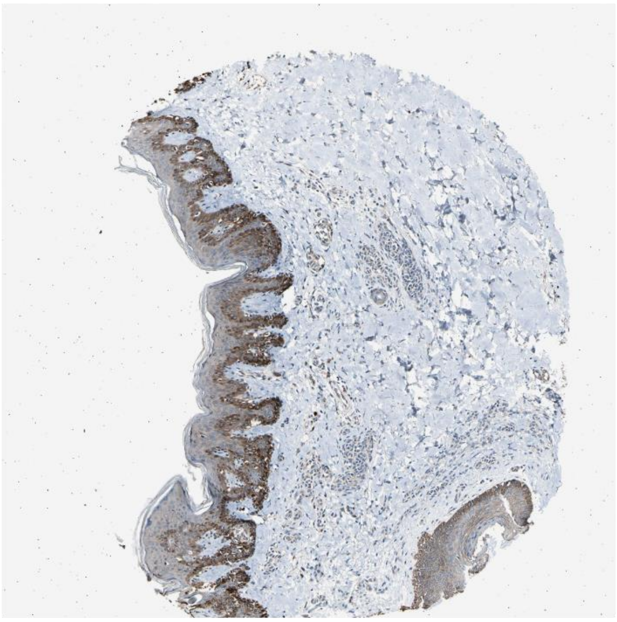

Hspg2 Lrp1

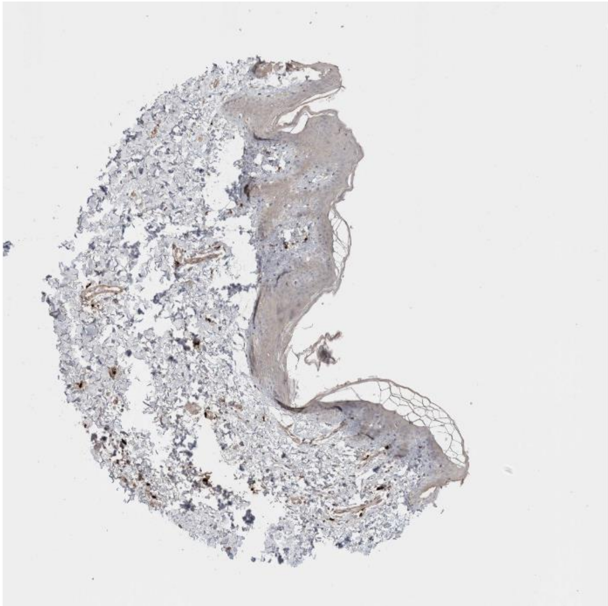

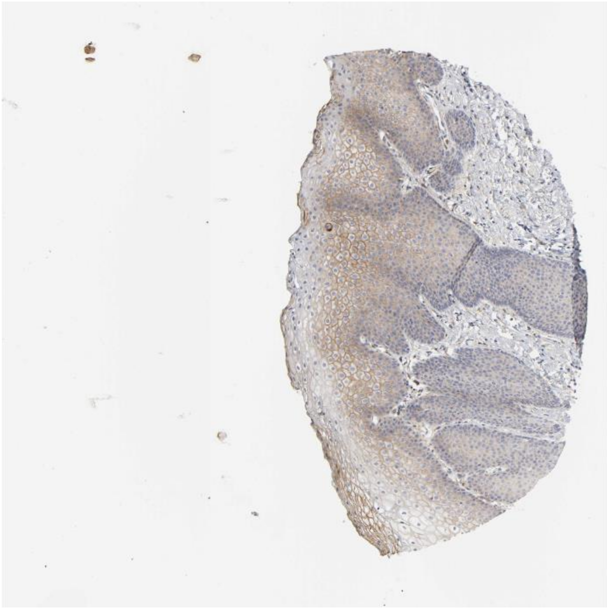

Thbs2 Itga6

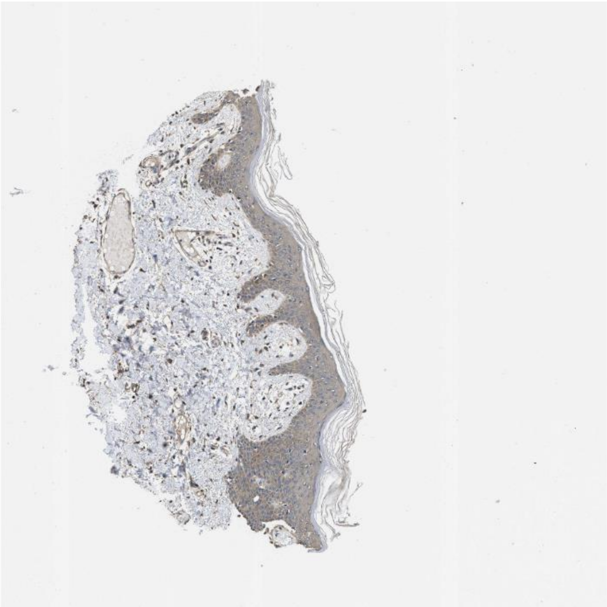

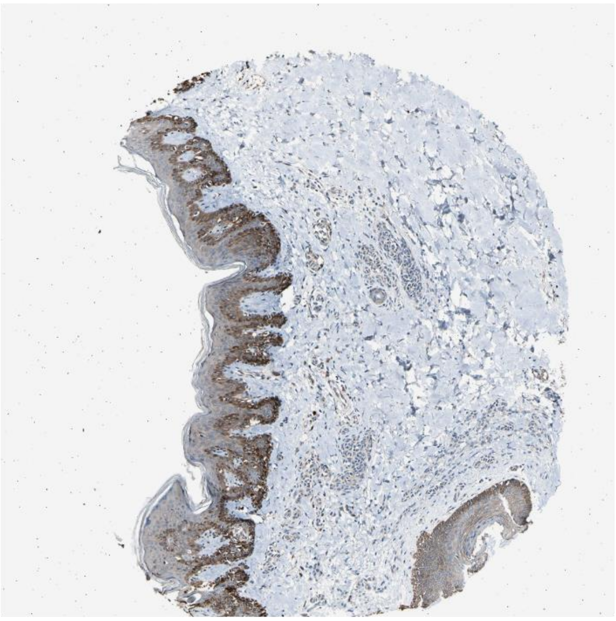

Fgf18 Fgfr1

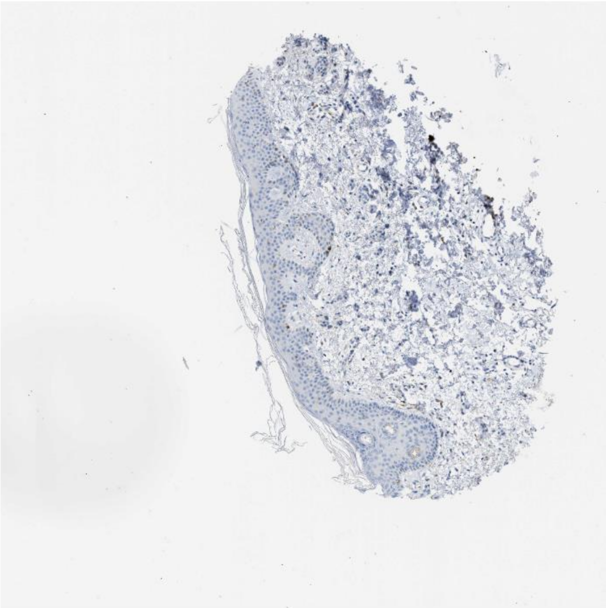

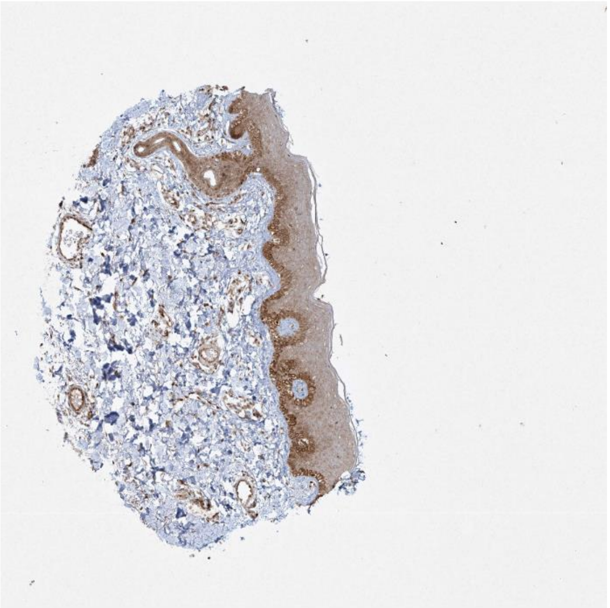

Pros1 Axl

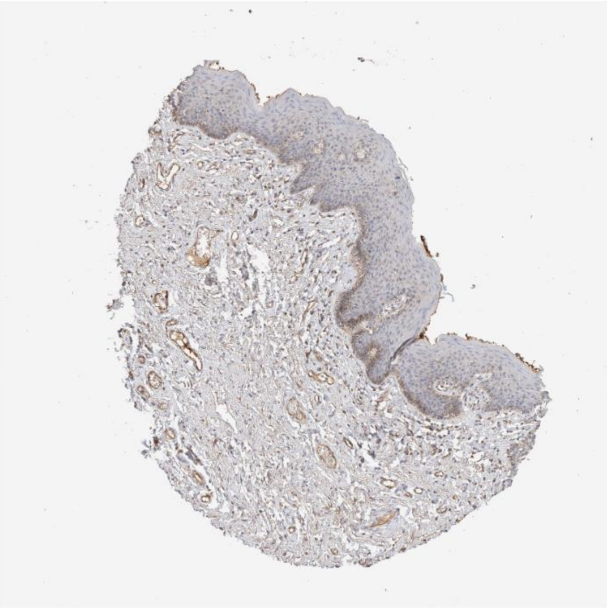

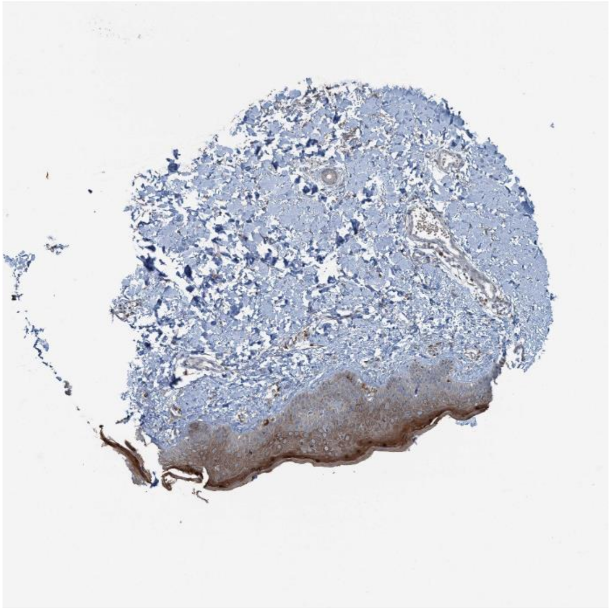

Ltbp1 Itgb5

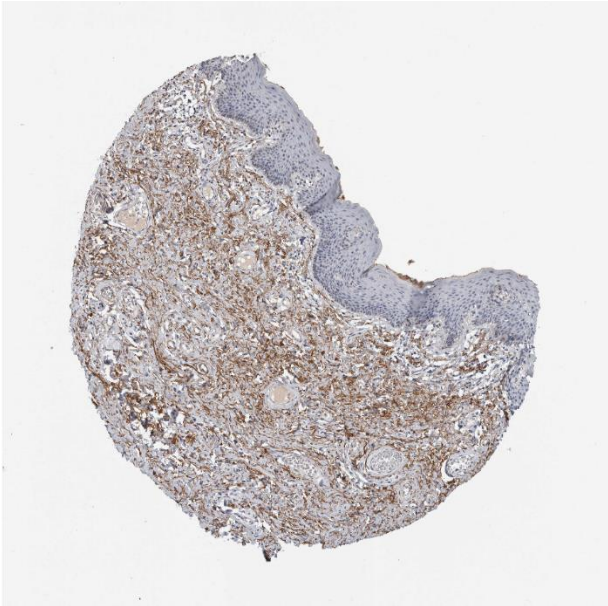

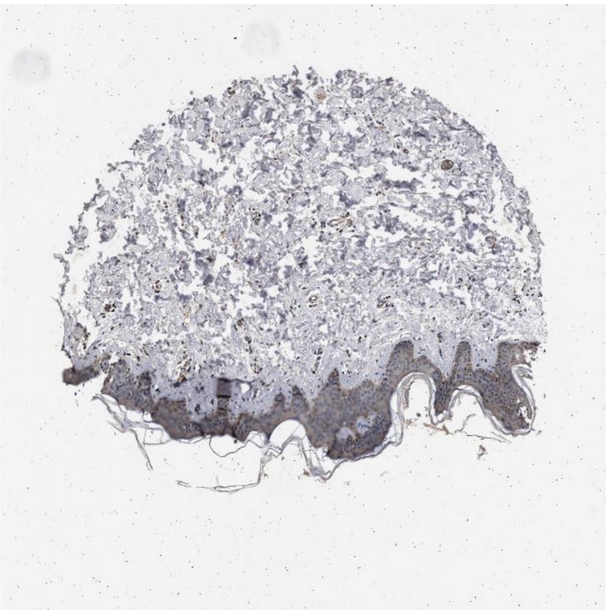

Nampt Insr

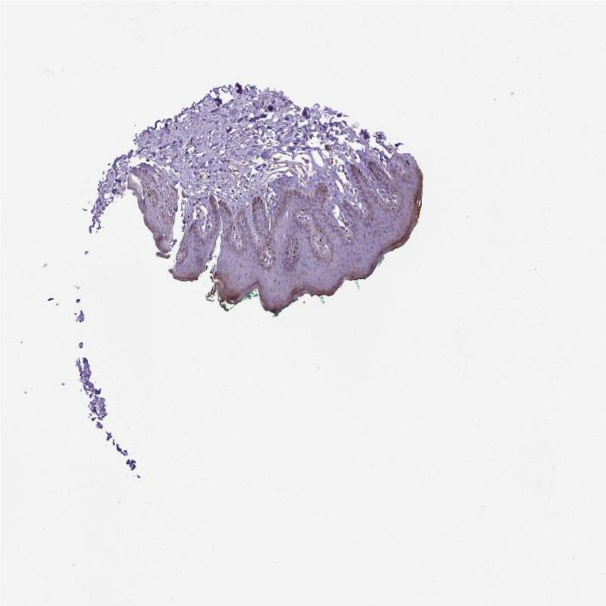

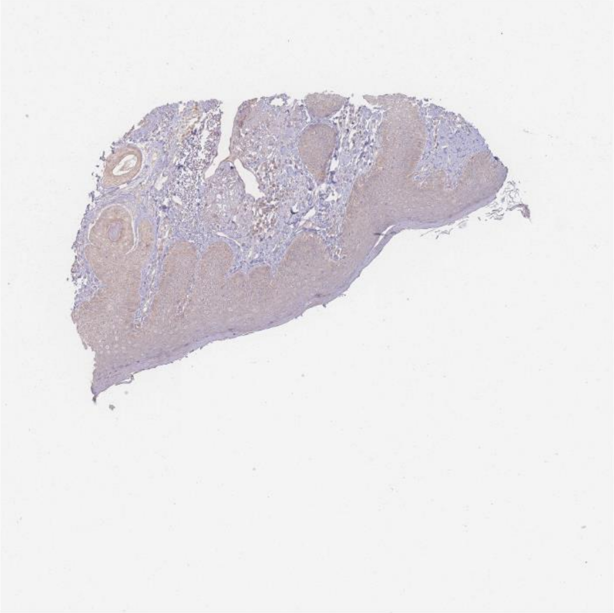

Rps27a Smad3

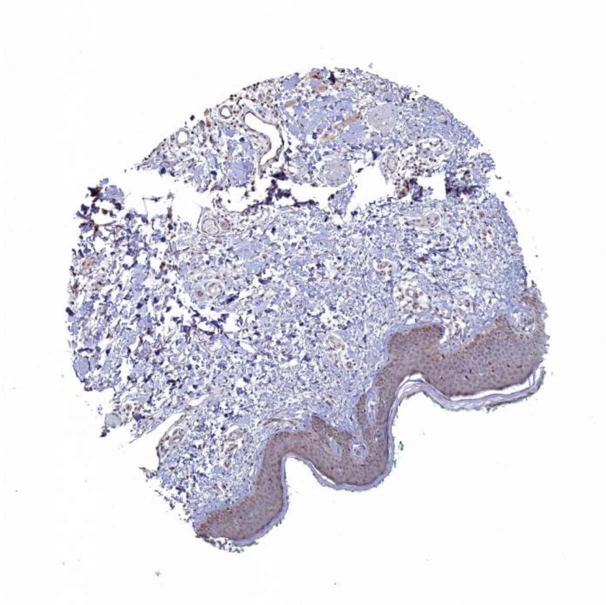

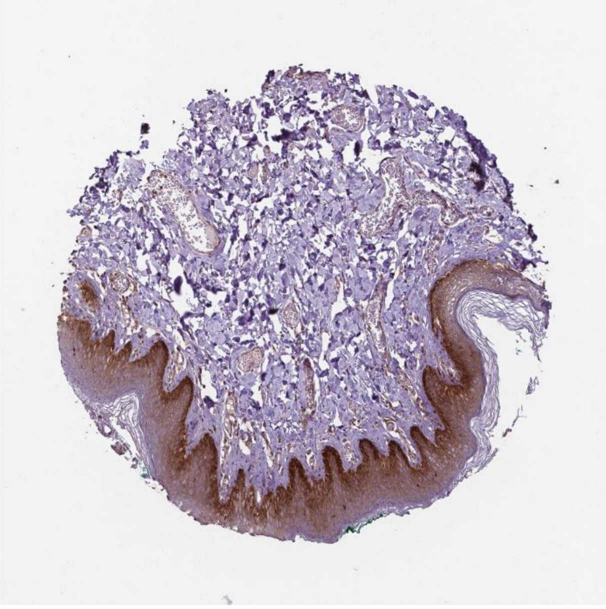

Thbs2 Cd47

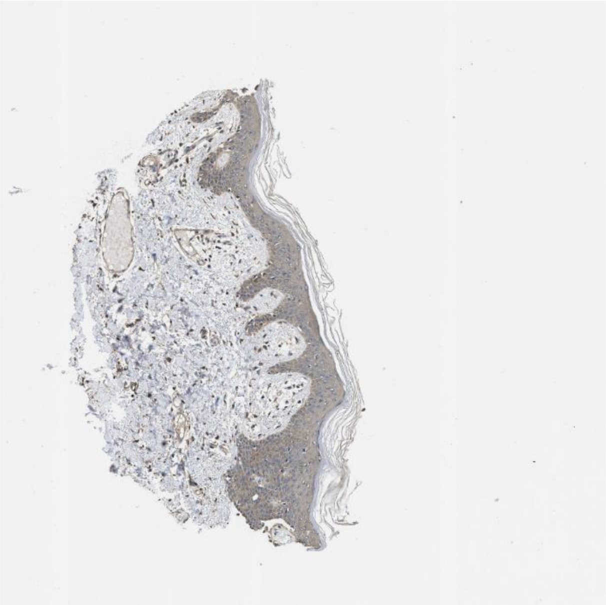

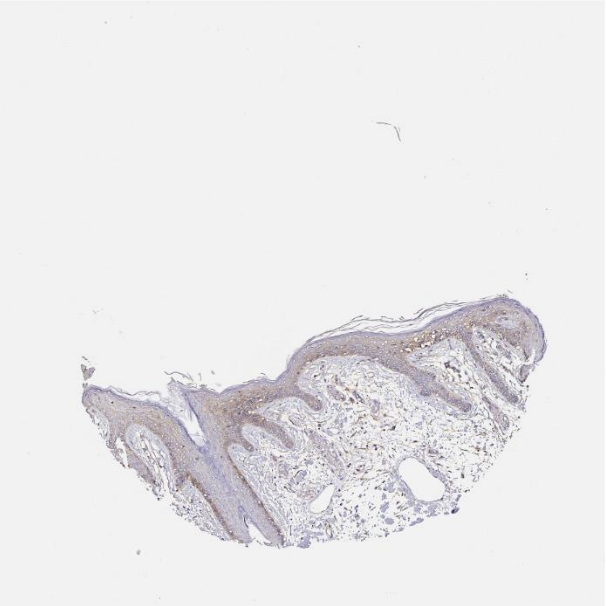
