## Supplemental file 1 - Supplemental material for "SingleCellSignalR: Inference of intercellular networks from single-cell transcriptomics"

**Supplementary Figures & Tables**

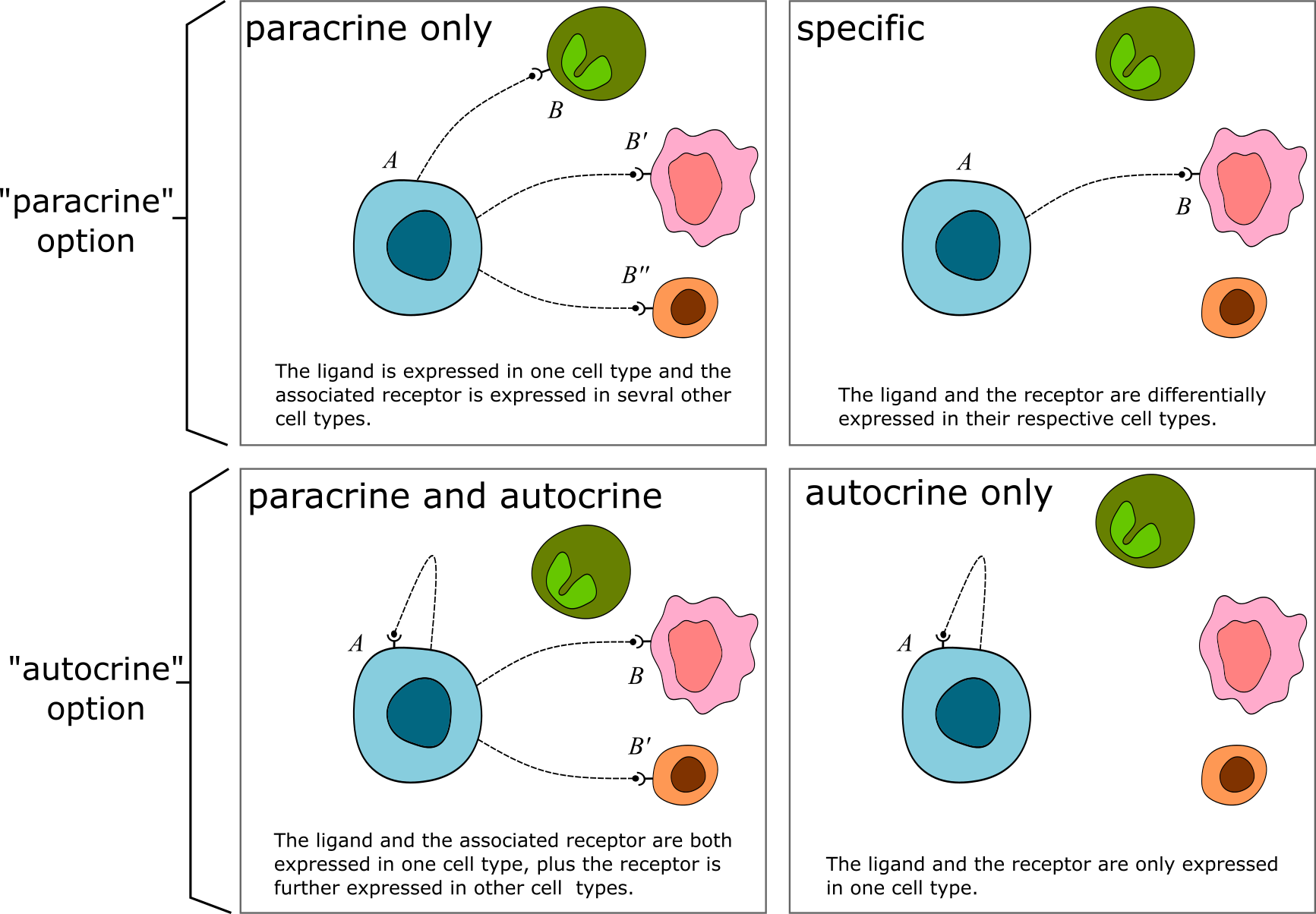

**Supplementary Figure 1.** Different types of cellular interactions between cell populations $A$ and $B$ can be inferred by SingleCellSignalR depending on the chosen options. The *specific* case is a particular case of the paracrine interaction.

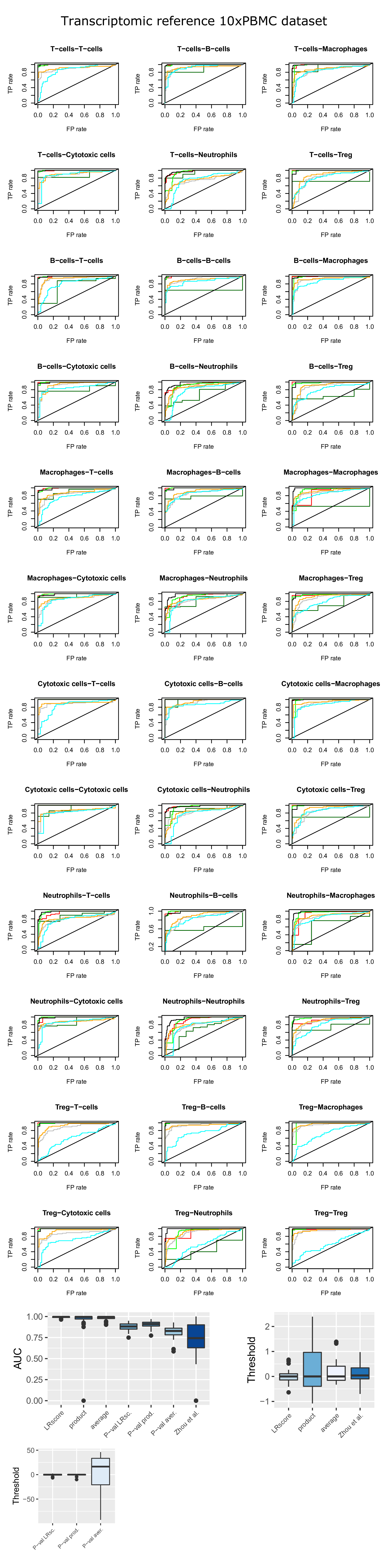

**Supplementary Figure 2.** ROC curves of LR pairs across cell populations of the 10xPBMC data set (1) with respect to the transcriptomic reference (2).

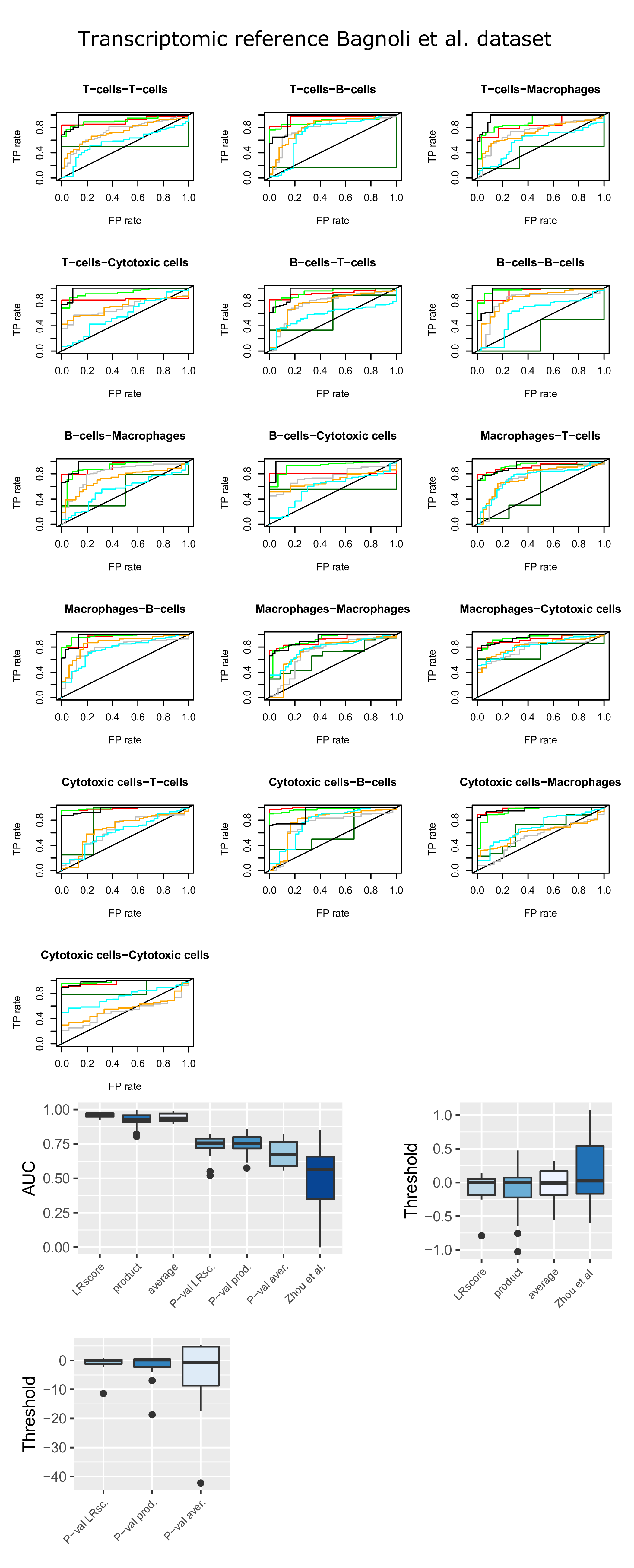

**Supplementary Figure 3.** ROC curves of LR pairs across cell populations of the PBMC data set (3) with respect to the transcriptomic reference (2).

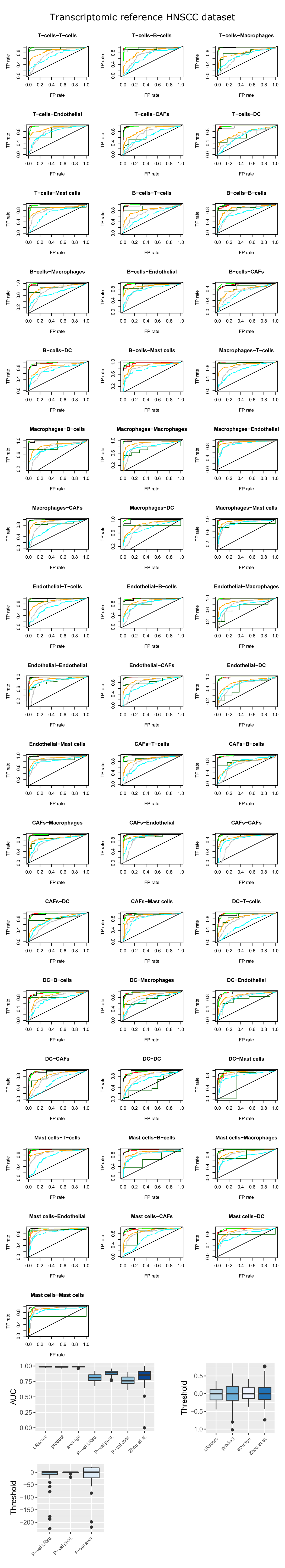

**Supplementary Figure 4.** ROC curves of LR pairs across cell populations of the HNSCC data set (4) with respect to the transcriptomic reference (2).

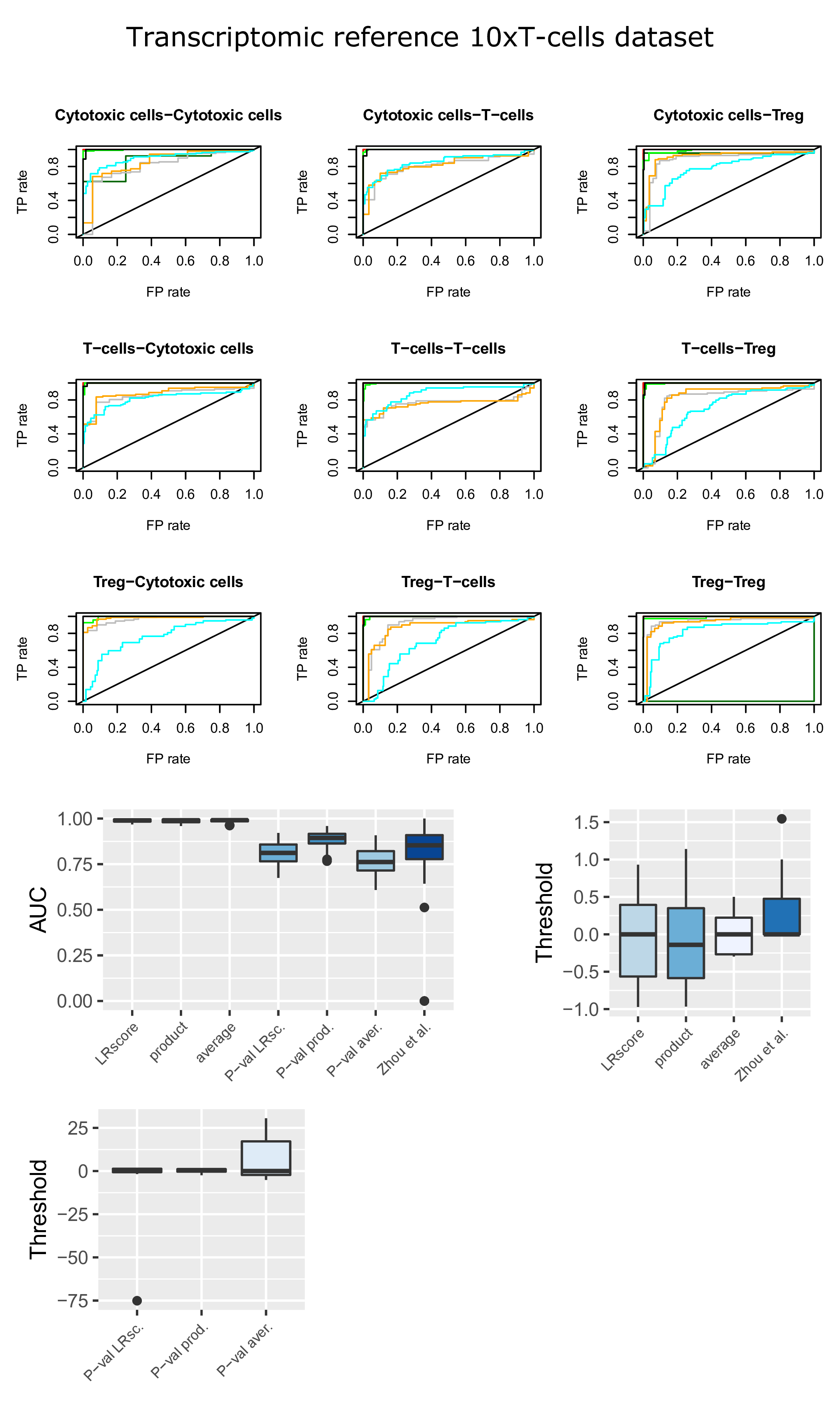

**Supplementary Figure 5.** ROC curves of LR pairs across cell populations of the 10xT data set (5) with respect to the transcriptomic reference (2).

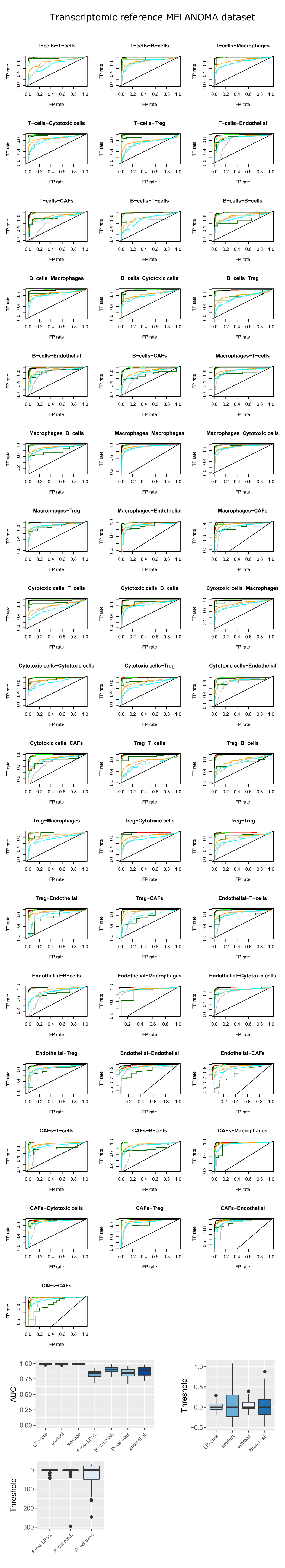

**Supplementary Figure 6.** ROC curves of LR pairs across cell populations of the MELANOMA data set (6) with respect to the transcriptomic reference (2).

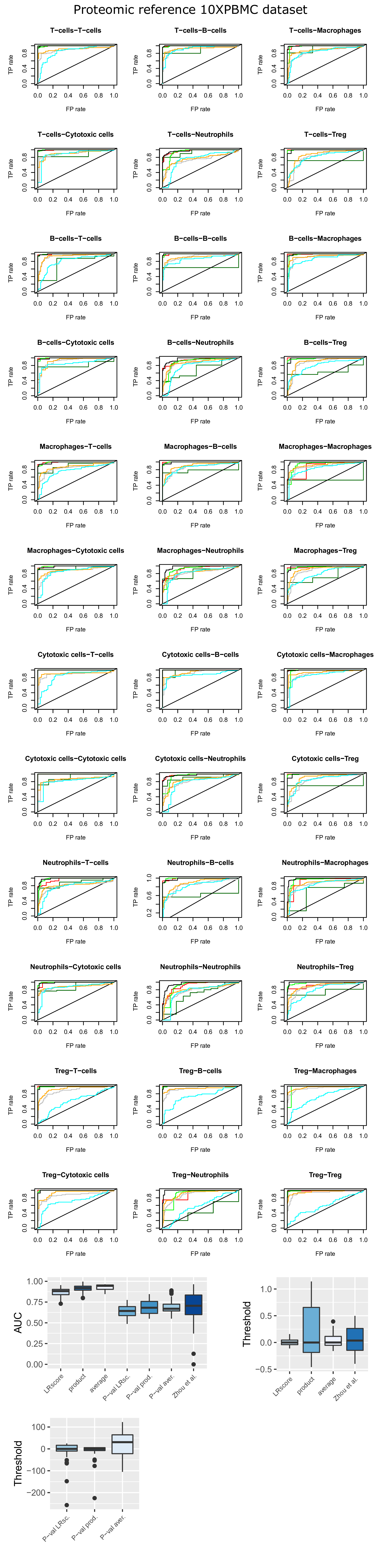

**Supplementary Figure 7.** ROC curves of LR pairs across cell populations of the 10xPBMC data set (1) with respect to the proteomic reference (7).

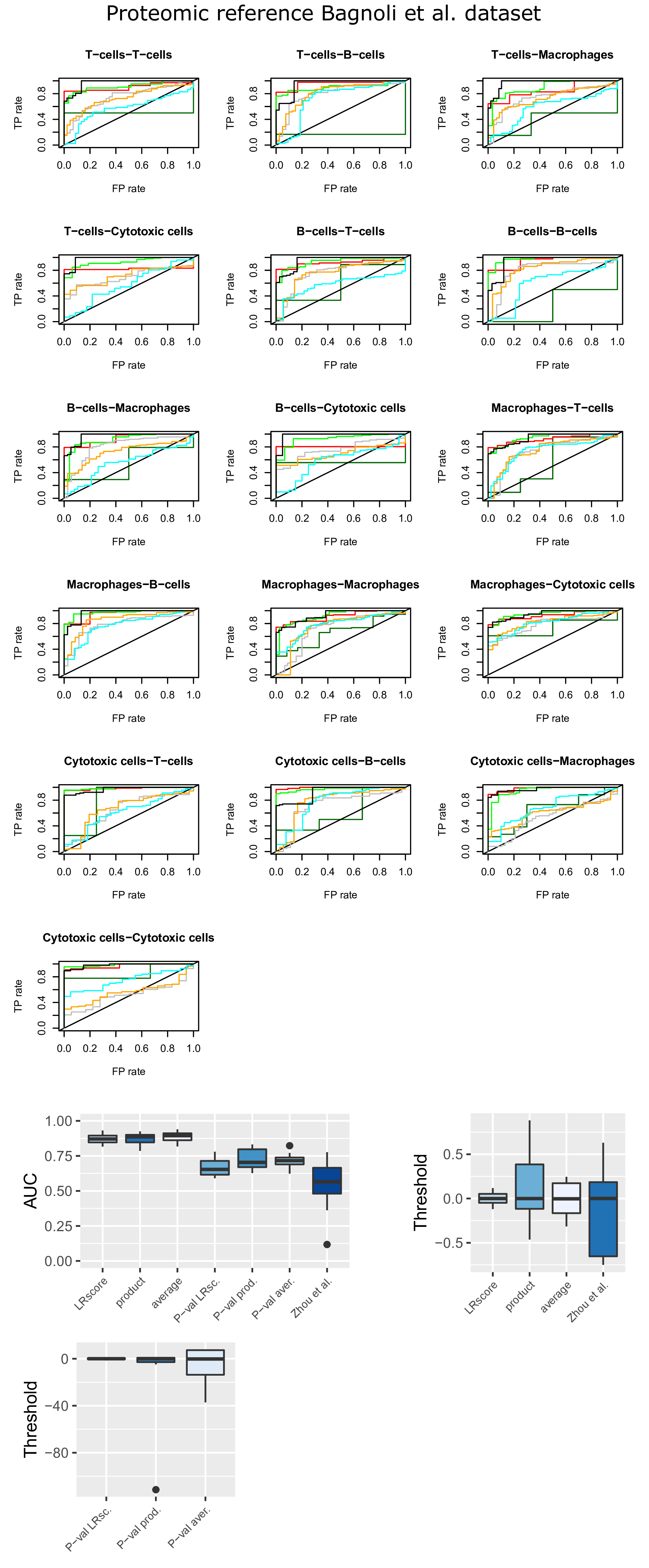

**Supplementary Figure 8.** ROC curves of LR pairs across cell populations of the PBMC data set (3) with respect to the proteomic reference (7).

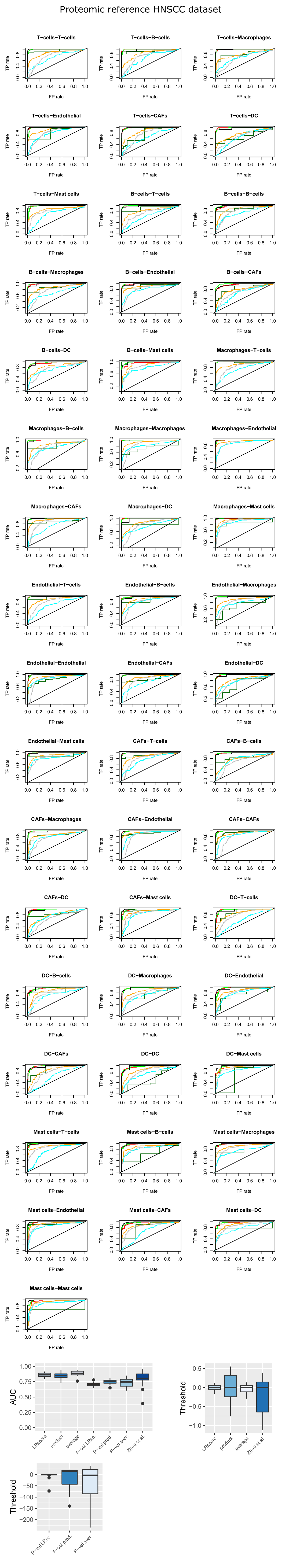

**Supplementary Figure 9.** ROC curves of LR pairs across cell populations of the HNSCC data set (4) with respect to the transcriptomic reference (7).

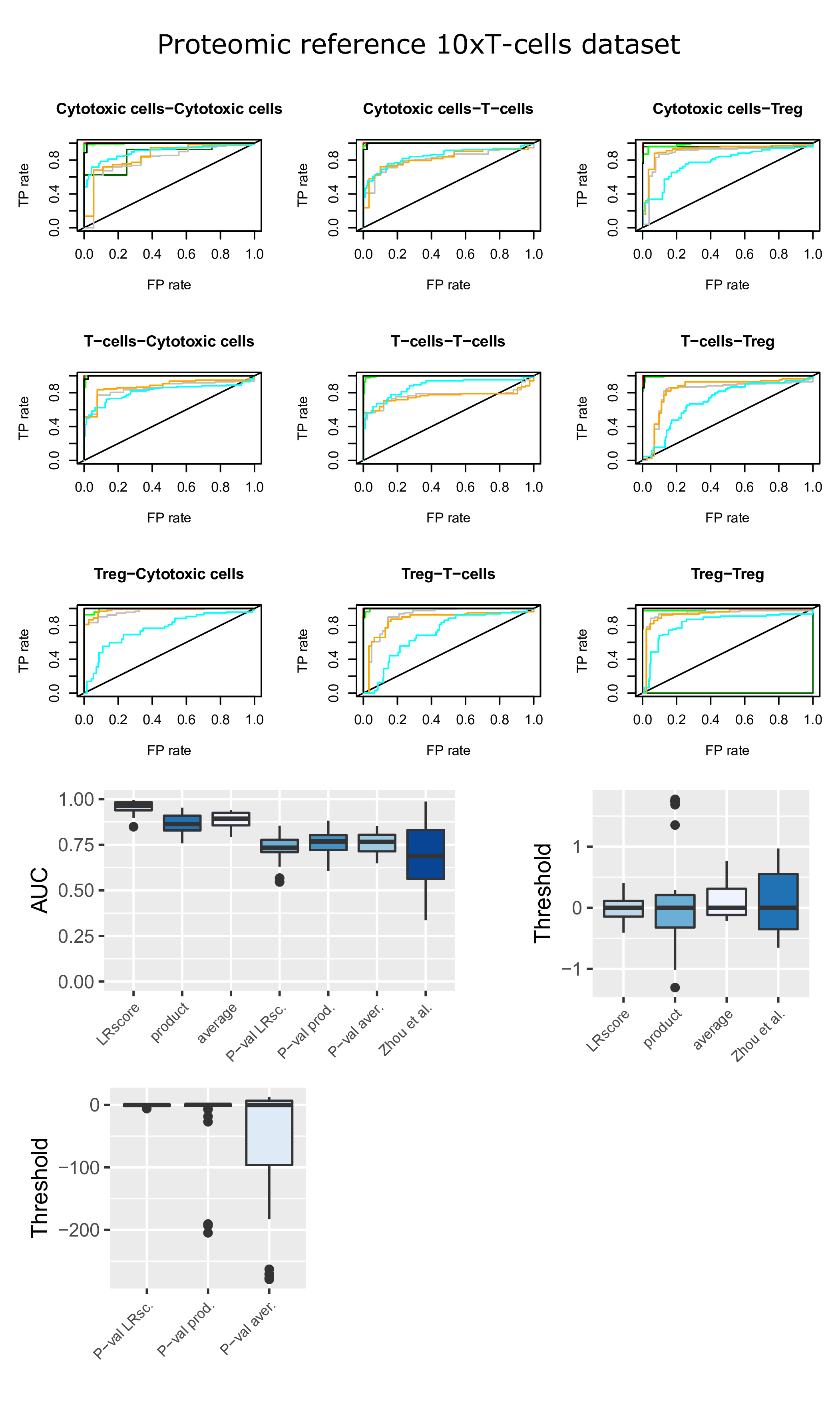

**Supplementary Figure 10.** ROC curves of LR pairs across cell populations of the 10xT data set (5) with respect to the transcriptomic reference (2).

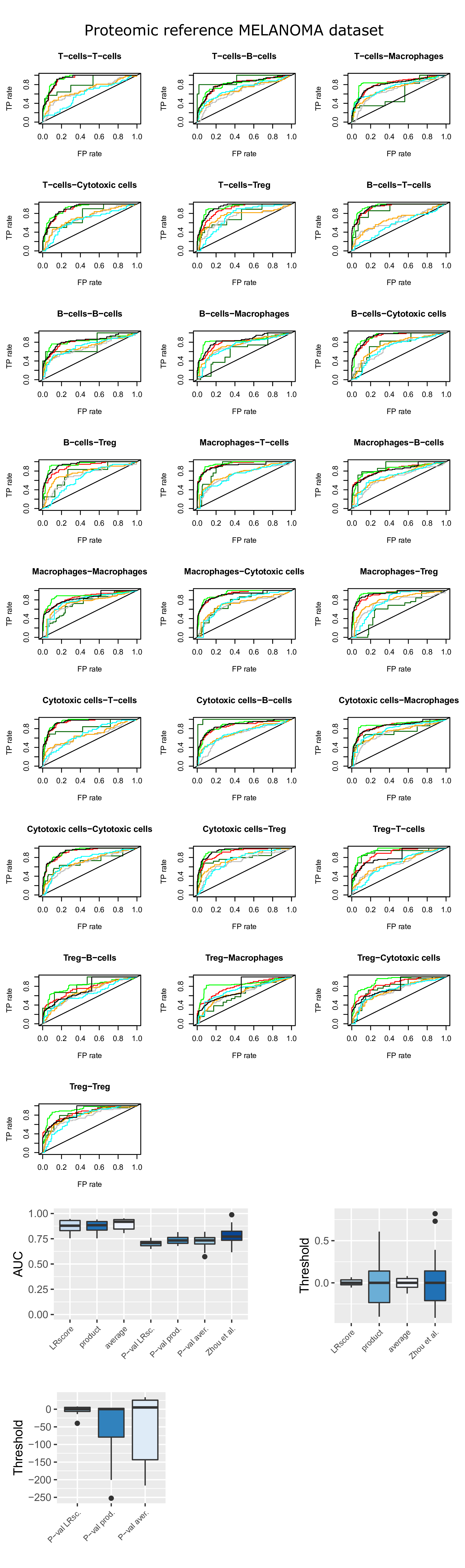

**Supplementary Figure 11.** ROC curves of LR pairs across cell populations of the MELANOMA data set (6) with respect to the proteomic reference (7).

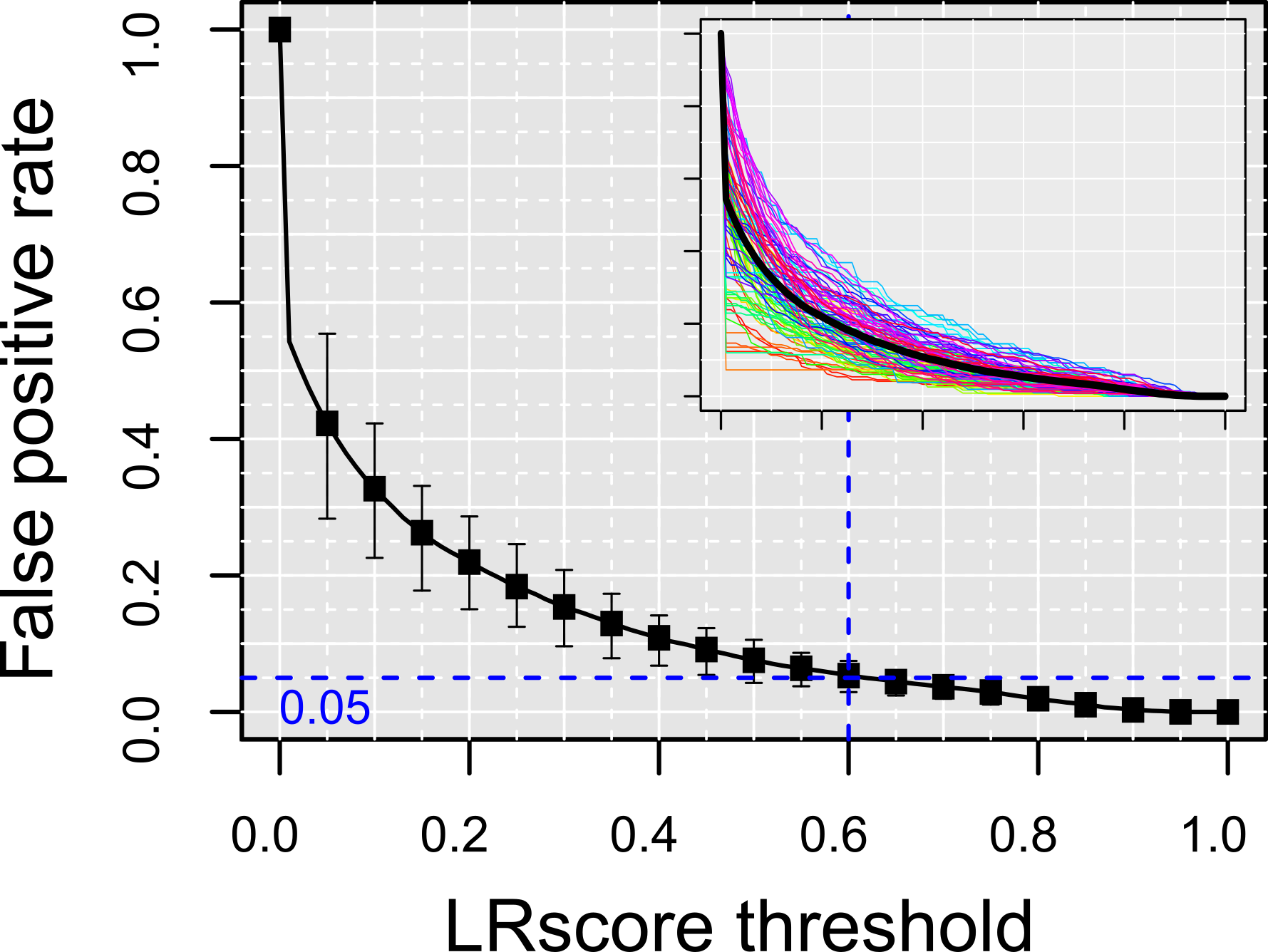

**Supplementary Figure 12.** Selection of the LRscore threshold requiring maximum 5% FP rate against the proteomic reference data set (7) for at least 75% of the ROC curves. Compare with Fig. 2D in the main text.

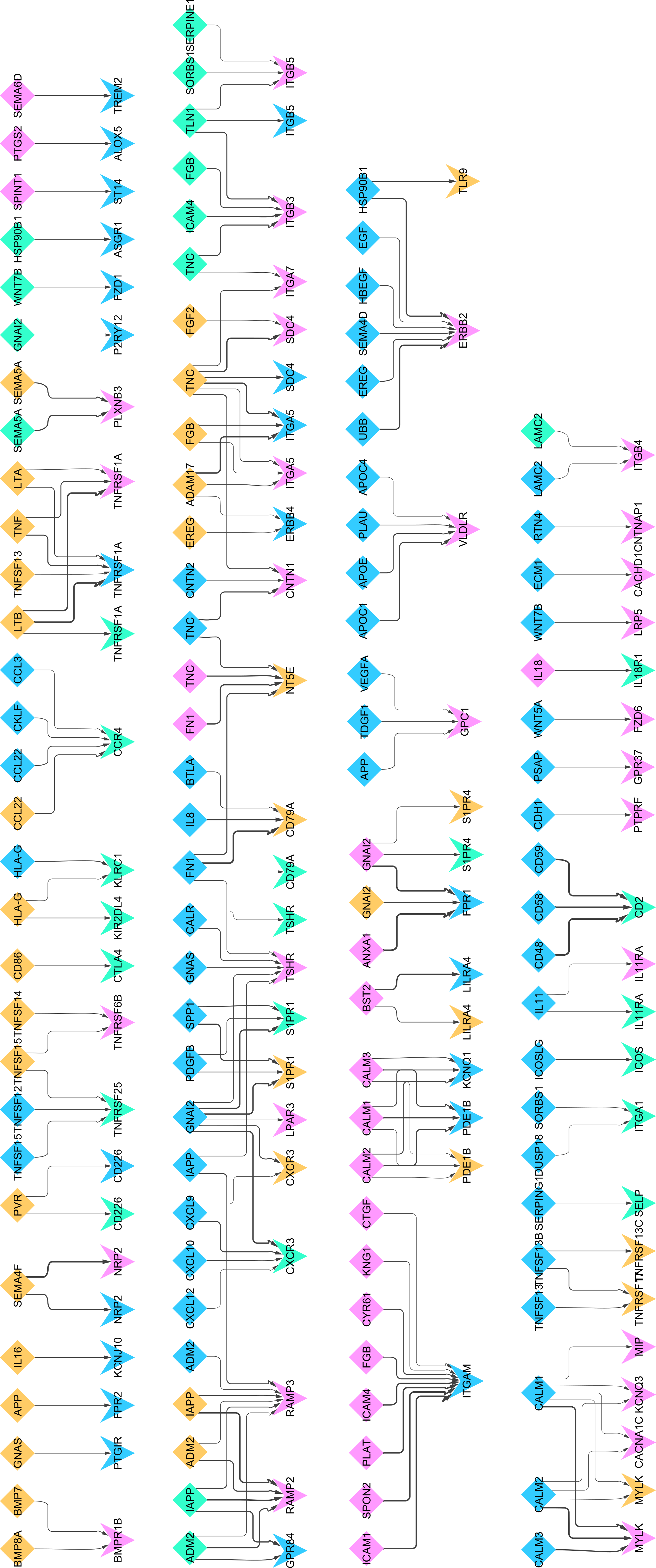

**Supplementary Figure 13.** Metastatic melanoma intercellular network, MELANOMA data set (6), patient 89.

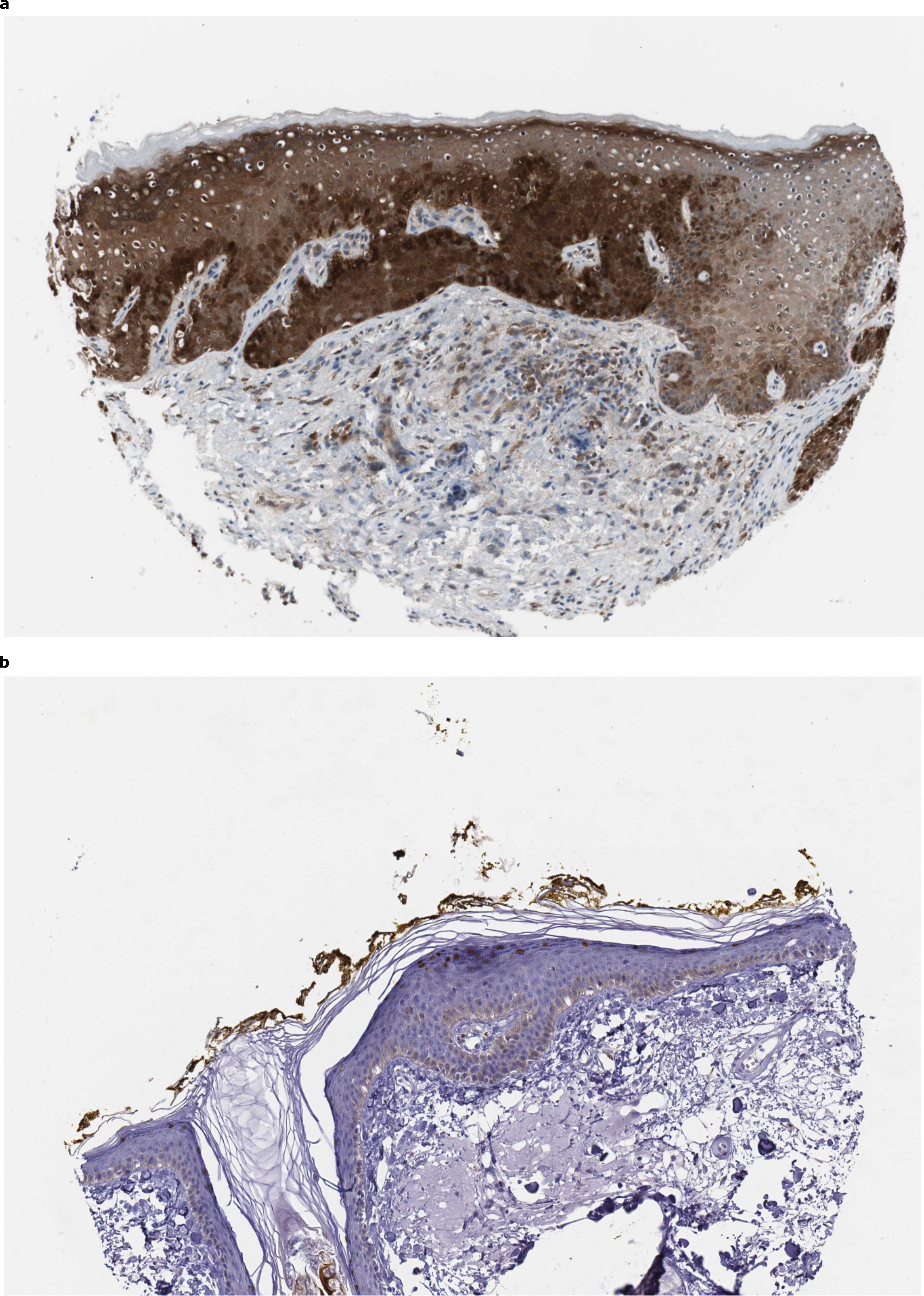

**Supplementary Figure 14.** Human Protein Atlas epidermis staining. (**a**) IL18 immunostaining in epidermal cells shows high expression in keratinocytes (CAB007772 andtibody). (**b**) PRDM1 immunostaining in epidermal cells shows medium expression in keratinocytes (HPA030033 andtibody).

**Supplementary Table 1.** Confirmation of mouse keratinized and suprabasal layer interactions. L/R in HPA indicate presence status in Human Protein Atlas skin data of the ligand, respectively the receptor (NA stands for absent, TRUE stands for present in the correct layer, FALSE for present but in the wrong layer). HPA is the HPA confirmation status. IF indicates the result of our IF mouse validation experiment. Literature indicates evidence from the literature. Confirmed is the final status. We considered that mouse IF or literature could supersede HPA. Low expression in HPA was considered positive. All HPA pictures are in supplementary data.

| **Keratinised** | **Suprabasal** | **LRscore** | **L in HPA** | **R in HPA** | **HPA** | **IF** | **Literature** | **Confirmed** |
| --- | --- | --- | --- | --- | --- | --- | --- | --- |
| Cdh1 | Ptprf | 0.905 | TRUE | TRUE | yes | - | - | yes |
| Hbegf | Cd9 | 0.903 | FALSE | TRUE | no | - | yes | yes |
| Calm1 | Ptpra | 0.880 | TRUE | TRUE | yes | - | - | yes |
| Calm1 | Egfr | 0.862 | TRUE | TRUE | yes | - | - | yes |
| Ubc | Ldlr | 0.857 | TRUE | TRUE | yes | - | - | yes |
| Ereg | Erbb2 | 0.855 | NA | TRUE | NA | - | - | NA |
| Efnb2 | Ephb6 | 0.852 | TRUE | NA | NA | - | - | NA |
| Calm2 | Egfr | 0.851 | TRUE | TRUE | yes | - | - | yes |
| Dsc3 | Dsg1b | 0.851 | TRUE | NA | NA | - | - | NA |
| Ubc | Erbb2 | 0.849 | TRUE | TRUE | yes | - | - | yes |
| Ubc | Adrb2 | 0.846 | TRUE | NA | NA | - | - | NA |
| Btc | Erbb2 | 0.843 | NA | TRUE | NA | - | - | NA |
| Rps27a | Ldlr | 0.843 | TRUE | TRUE | yes | - | - | yes |
| B2m | Tfrc | 0.842 | TRUE | TRUE | yes | - | - | yes |
| Apoe | Ldlr | 0.840 | TRUE | TRUE | yes | - | - | yes |
| Calm1 | Insr | 0.838 | TRUE | TRUE | yes | - | - | yes |
| Anxa1 | Egfr | 0.837 | TRUE | TRUE | yes | - | - | yes |
| Ptn | Sdc1 | 0.833 | FALSE | TRUE | no | - | - | no |
| Rps27a | Erbb2 | 0.833 | TRUE | TRUE | yes | - | - | yes |
| Rps27a | Adrb2 | 0.830 | TRUE | NA | NA | - | - | NA |
| Cgn | F11r | 0.829 | TRUE | TRUE | yes | - | - | yes |
| Dsc1 | Dsg1b | 0.829 | TRUE | NA | NA | - | - | NA |
| Hspa8 | Ldlr | 0.828 | TRUE | TRUE | yes | - | - | yes |
| Calm2 | Insr | 0.826 | TRUE | TRUE | yes | - | - | yes |
| Efna1 | Ephb6 | 0.825 | TRUE | NA | NA | - | - | NA |
| Ereg | Egfr | 0.824 | NA | TRUE | NA | - | - | NA |
| Efnb2 | Epha4 | 0.822 | TRUE | TRUE | yes | - | - | yes |
| Ubc | Fgfr2 | 0.822 | TRUE | TRUE | yes | - | - | yes |
| Hsp90b1 | Erbb2 | 0.819 | TRUE | TRUE | yes | - | - | yes |
| Fabp5 | Rxra | 0.817 | TRUE | TRUE | yes | - | - | yes |
| App | Ncstn | 0.815 | NA | TRUE | NA | - | - | NA |
| Hspa8 | Adrb2 | 0.815 | TRUE | NA | NA | - | - | NA |
| Ubb | Ldlr | 0.813 | TRUE | TRUE | yes | - | - | yes |
| App | Cav1 | 0.813 | NA | TRUE | NA | - | - | NA |
| Actr2 | Ldlr | 0.812 | TRUE | TRUE | yes | - | - | yes |
| Hbegf | **Cd44** | 0.810 | FALSE | TRUE | no | yes | yes | yes |
| Btc | Egfr | 0.809 | NA | TRUE | NA | - | - | NA |
| App | Lrp1 | 0.807 | NA | TRUE | NA | - | - | NA |
| Hbegf | Cd82 | 0.804 | FALSE | TRUE | no | - | yes | yes |
| Rps27a | Fgfr2 | 0.804 | TRUE | TRUE | yes | - | - | yes |
| Ubb | Erbb2 | 0.802 | TRUE | TRUE | yes | - | - | yes |
| Arpc5 | Ldlr | 0.799 | NA | TRUE | NA | - | - | NA |
| **Psen1** | Notch1 | 0.799 | FALSE | TRUE | no | yes | - | yes |
| Ubb | Adrb2 | 0.798 | TRUE | NA | NA | - | - | NA |
| Dusp18 | Itga6 | 0.798 | TRUE | TRUE | yes | - | - | yes |
| Actr2 | Adrb2 | 0.798 | TRUE | NA | NA | - | - | NA |
| Efna5 | Ephb6 | 0.797 | TRUE | NA | NA | - | - | NA |
| Hbegf | Erbb2 | 0.792 | FALSE | TRUE | no | - | yes | yes |
| Cgn | Ocln | 0.792 | TRUE | TRUE | yes | - | - | yes |
| Efna1 | Epha4 | 0.790 | TRUE | TRUE | yes | - | - | yes |
| Arpc5 | Adrb2 | 0.784 | NA | NA | NA | - | - | NA |
| Calm1 | Abca1 | 0.784 | TRUE | TRUE | yes | - | - | yes |
| Cdh1 | Egfr | 0.784 | TRUE | TRUE | yes | - | - | yes |
| **Psen1** | **Cd44** | **0.780** | **FALSE** | **TRUE** | **no** | yes | - | yes |
| Adam17 | Notch1 | 0.778 | TRUE | TRUE | yes | - | - | yes |
| App | Gpc1 | 0.778 | NA | TRUE | NA | - | - | NA |
| Apoe | Lrp1 | 0.776 | TRUE | TRUE | yes | - | - | yes |
| Hsp90aa1 | Fgfr3 | 0.776 | NA | TRUE | NA | - | - | NA |
| Calr | Itgav | 0.775 | TRUE | TRUE | yes | - | - | yes |
| Cdh1 | Igf1r | 0.772 | TRUE | TRUE | yes | - | - | yes |
| Hsp90aa1 | Erbb2 | 0.771 | NA | TRUE | NA | - | - | NA |
| Sorbs1 | Itgb5 | 0.770 | TRUE | TRUE | yes | - | - | yes |
| Efnb1 | Ephb6 | 0.770 | TRUE | NA | NA | - | - | NA |
| Ubb | Fgfr2 | 0.769 | TRUE | TRUE | yes | - | - | yes |
| Calm2 | Abca1 | 0.769 | TRUE | TRUE | yes | - | - | yes |
| Pkm | **Cd44** | 0.764 | TRUE | TRUE | yes | yes | - | yes |
| Sema4a | Plxnb2 | 0.764 | NA | TRUE | NA | - | - | NA |
| Calcb | Adrb2 | 0.762 | NA | NA | NA | - | - | NA |
| Hsp90b1 | Lrp1 | 0.762 | TRUE | TRUE | yes | - | - | yes |
| Efna5 | Epha4 | 0.759 | TRUE | TRUE | yes | - | - | yes |
| Tgfa | Erbb2 | 0.759 | TRUE | TRUE | yes | - | - | yes |
| Sptan1 | Ptpra | 0.758 | FALSE | TRUE | no | - | - | no |
| Fst | Bmpr2 | 0.756 | NA | FALSE | no | - | - | no |
| Arf1 | Insr | 0.755 | TRUE | TRUE | yes | - | - | yes |
| Sptbn2 | Ptpra | 0.753 | TRUE | TRUE | yes | - | - | yes |
| Lamb3 | Col17a1 | 0.753 | TRUE | TRUE | yes | - | - | yes |
| Hbegf | Egfr | 0.751 | FALSE | TRUE | no | - | yes | yes |
| Timp3 | Adam17 | 0.750 | NA | TRUE | NA | - | - | NA |
| Ecm1 | Cachd1 | 0.750 | TRUE | TRUE | yes | - | - | yes |
| **Psen1** | Notch2 | 0.750 | FALSE | TRUE | no | yes | - | yes |
| Hspg2 | Sdc1 | 0.742 | TRUE | TRUE | yes | - | - | yes |
| Ubc | Ripk1 | 0.741 | TRUE | TRUE | yes | - | - | yes |
| Adam10 | Notch1 | 0.741 | TRUE | TRUE | yes | - | - | yes |
| Calr | Lrp1 | 0.740 | TRUE | TRUE | yes | - | - | yes |
| Ereg | Erbb3 | 0.734 | NA | TRUE | NA | - | - | NA |
| Ptn | Plxnb2 | 0.732 | NA | TRUE | NA | - | - | NA |
| Gas6 | Tyro3 | 0.732 | FALSE | TRUE | no | - | yes | yes |
| Dusp18 | Cd151 | 0.728 | TRUE | TRUE | yes | - | - | yes |
| Dusp18 | Itgb4 | 0.728 | TRUE | TRUE | yes | - | - | yes |
| Hsp90aa1 | Egfr | 0.727 | NA | TRUE | NA | - | - | NA |
| Efnb2 | Ephb3 | 0.721 | TRUE | TRUE | yes | - | - | yes |
| Sorbs1 | Insr | 0.721 | TRUE | TRUE | yes | - | - | yes |
| Efna1 | Epha2 | 0.719 | TRUE | NA | NA | - | - | NA |
| Rps27a | Ripk1 | 0.718 | TRUE | TRUE | yes | - | - | yes |
| Rgmb | Bmpr2 | 0.716 | NA | FALSE | no | - | - | no |
| Btc | Erbb3 | 0.715 | NA | TRUE | NA | - | - | NA |
| Efna1 | Epha1 | 0.714 | TRUE | TRUE | yes | - | - | yes |
| Tgfa | Egfr | 0.713 | TRUE | TRUE | yes | - | - | yes |
| Efnb1 | Erbb2 | 0.710 | TRUE | TRUE | yes | - | - | yes |
| Lamc2 | Col17a1 | 0.709 | NA | TRUE | NA | - | - | NA |
| Mapk1 | Fgfr2 | 0.708 | TRUE | TRUE | yes | - | - | yes |
| Jag1 | Notch1 | 0.705 | TRUE | TRUE | yes | - | - | yes |
| Il18 | Il18r1 | 0.704 | TRUE | TRUE | yes | - | - | yes |
| **Psen1** | Ncstn | 0.704 | FALSE | TRUE | no | yes | - | yes |
| B2m | Cd3g | 0.696 | TRUE | TRUE | yes | - | - | yes |
| Spint1 | St14 | 0.695 | TRUE | NA | NA | - | - | NA |
| Ubc | Tgfbr1 | 0.692 | TRUE | TRUE | yes | - | - | yes |
| Fgf18 | Fgfr3 | 0.690 | FALSE | TRUE | no | - | - | no |
| Hbegf | Prlr | 0.689 | FALSE | NA | no | - | yes | yes |
| Gpi1 | Amfr | 0.687 | TRUE | TRUE | yes | - | - | yes |
| App | Tnfrsf21 | 0.685 | NA | TRUE | NA | - | - | NA |
| Cdh1 | Erbb3 | 0.682 | TRUE | TRUE | yes | - | - | yes |
| Efna5 | Epha2 | 0.681 | TRUE | NA | NA | - | - | NA |
| Il1rn | Il1r2 | 0.677 | TRUE | TRUE | yes | - | - | yes |
| Ubc | Tgfbr2 | 0.677 | TRUE | NA | NA | - | - | NA |
| Bmp2 | Bmpr2 | 0.676 | NA | FALSE | no | - | - | no |
| Efna5 | Epha1 | 0.676 | TRUE | TRUE | yes | - | - | yes |
| Ubb | Ripk1 | 0.674 | TRUE | TRUE | yes | - | - | yes |
| Thbs1 | Sdc1 | 0.673 | TRUE | TRUE | yes | - | - | yes |
| Apoe | Lrp5 | 0.670 | TRUE | TRUE | yes | - | - | yes |
| Apoe | Sorl1 | 0.669 | TRUE | TRUE | yes | - | - | yes |
| Rps27a | Tgfbr1 | 0.667 | TRUE | TRUE | yes | - | - | yes |
| Efnb2 | Rhbdl2 | 0.667 | TRUE | TRUE | yes | - | - | yes |
| Gas6 | Axl | 0.666 | FALSE | TRUE | no | - | yes | yes |
| Dusp18 | Itgb1 | 0.666 | TRUE | TRUE | yes | - | - | yes |
| Efnb2 | Ephb4 | 0.666 | TRUE | TRUE | yes | - | - | yes |
| Tgs1 | Rxra | 0.665 | TRUE | TRUE | yes | - | - | yes |
| Gnai2 | Egfr | 0.664 | TRUE | TRUE | yes | - | - | yes |
| Lama5 | Sdc1 | 0.664 | TRUE | TRUE | yes | - | - | yes |
| Bmp7 | Bmpr2 | 0.662 | NA | FALSE | no | - | - | no |
| Cgn | Tgfbr1 | 0.661 | TRUE | TRUE | yes | - | - | yes |
| Areg | Egfr | 0.659 | TRUE | TRUE | yes | - | - | yes |
| Lrpap1 | Ldlr | 0.659 | TRUE | TRUE | yes | - | - | yes |
| B2m | Hfe | 0.654 | TRUE | TRUE | yes | - | - | yes |
| Lpl | Sdc1 | 0.654 | NA | TRUE | NA | - | - | NA |
| Cdh1 | Lrp5 | 0.654 | TRUE | TRUE | yes | - | - | yes |
| Ptdss1 | Jmjd6 | 0.653 | FALSE | TRUE | no | - | - | no |
| Sema4a | Plxnb1 | 0.652 | NA | TRUE | NA | - | - | NA |
| Rps27a | Tgfbr2 | 0.651 | TRUE | NA | NA | - | - | NA |
| Gnai2 | Igf1r | 0.649 | TRUE | TRUE | yes | - | - | yes |
| Gnai2 | Cav1 | 0.647 | TRUE | TRUE | yes | - | - | yes |
| Lamb3 | Itga6 | 0.646 | TRUE | TRUE | yes | - | - | yes |
| Cgn | Tgfbr2 | 0.645 | TRUE | NA | NA | - | - | NA |
| Agrn | Lrp4 | 0.644 | TRUE | TRUE | yes | - | - | yes |
| Jag1 | Notch2 | 0.643 | TRUE | TRUE | yes | - | - | yes |
| Fgf18 | Fgfr2 | 0.641 | FALSE | TRUE | no | - | - | no |
| Psap | Sort1 | 0.638 | TRUE | TRUE | yes | - | - | yes |
| Il1a | Il1r2 | 0.633 | NA | TRUE | NA | - | - | NA |
| B2m | Cd247 | 0.631 | TRUE | TRUE | yes | - | - | yes |
| Fgf22 | Fgfr2 | 0.630 | NA | TRUE | NA | - | - | NA |
| Gnai2 | S1pr5 | 0.626 | TRUE | TRUE | yes | - | - | yes |
| Psap | Celsr1 | 0.622 | TRUE | NA | NA | - | - | NA |
| Il1rn | Il1r1 | 0.621 | TRUE | TRUE | yes | - | - | yes |
| Ubb | Tgfbr1 | 0.619 | TRUE | TRUE | yes | - | - | yes |
| Ltbp3 | Itgb5 | 0.617 | NA | TRUE | NA | - | - | NA |
| Tnf | Tnfrsf1a | 0.617 | NA | TRUE | NA | - | - | NA |
| Lin7c | Abca1 | 0.616 | TRUE | TRUE | yes | - | - | yes |
| App | Cd74 | 0.614 | NA | FALSE | no | - | - | no |
| Vim | **Cd44** | 0.613 | TRUE | TRUE | yes | - | - | yes |
| Adam9 | Itga6 | 0.612 | NA | TRUE | NA | - | - | NA |
| Col6a1 | Itga6 | 0.611 | TRUE | TRUE | yes | - | - | yes |
| Rgma | Bmpr2 | 0.610 | TRUE | FALSE | no | - | - | no |
| Rtn4 | Rtn4rl1 | 0.606 | TRUE | TRUE | yes | - | - | yes |
| Calm1 | Pde1b | 0.606 | TRUE | TRUE | yes | - | - | yes |
| Wnt7b | Fzd10 | 0.604 | NA | TRUE | NA | - | - | NA |
| Lrpap1 | Sort1 | 0.604 | TRUE | TRUE | yes | - | - | yes |
| Ubb | Tgfbr2 | 0.602 | TRUE | NA | NA | - | - | NA |
| Timp2 | Itgb1 | 0.602 | TRUE | TRUE | yes | - | - | yes |
| Efna4 | Epha4 | 0.600 | TRUE | TRUE | yes | - | - | yes |
| Efnb1 | Ephb3 | 0.600 | TRUE | TRUE | yes | - | - | yes |
| Epgn | Egfr | 0.597 | NA | TRUE | NA | - | - | NA |
| Ptn | Ptprs | 0.597 | FALSE | NA | no | - | - | no |
| Psap | Lrp1 | 0.596 | TRUE | TRUE | yes | - | - | yes |
| Tgfa | Erbb3 | 0.595 | TRUE | TRUE | yes | - | - | yes |
| Calr | Itga3 | 0.594 | TRUE | TRUE | yes | - | - | yes |
| Adam17 | Itgb1 | 0.594 | TRUE | TRUE | yes | - | - | yes |
| Lamc2 | Itga6 | 0.593 | NA | TRUE | NA | - | - | NA |
| Tln1 | Itgb5 | 0.588 | TRUE | TRUE | yes | - | - | yes |
| Calm2 | Pde1b | 0.585 | TRUE | TRUE | yes | - | - | yes |
| Pros1 | Tyro3 | 0.585 | TRUE | TRUE | yes | - | - | yes |
| Pthlh | Adrb2 | 0.584 | TRUE | NA | NA | - | - | NA |
| Dsc3 | Dsg2 | 0.584 | TRUE | TRUE | yes | - | - | yes |
| Dusp18 | Itga3 | 0.583 | TRUE | TRUE | yes | - | - | yes |
| Bmp2 | Bmpr1a | 0.582 | NA | TRUE | NA | - | - | NA |
| Adam10 | Axl | 0.579 | TRUE | TRUE | yes | - | - | yes |
| Il1a | Il1r1 | 0.575 | NA | TRUE | NA | - | - | NA |
| Arf1 | Pld2 | 0.573 | TRUE | TRUE | yes | - | - | yes |
| Rgmb | Neo1 | 0.571 | NA | TRUE | NA | - | - | NA |
| B2m | Cd3d | 0.570 | TRUE | FALSE | no | - | - | no |
| Adam15 | Itgav | 0.569 | FALSE | TRUE | no | - | - | no |
| Adam9 | Itgav | 0.567 | NA | TRUE | NA | - | - | NA |
| Calm1 | Mylk | 0.567 | TRUE | FALSE | no | - | - | no |
| Bmp7 | Bmpr1a | 0.567 | NA | TRUE | NA | - | - | NA |
| Adam9 | Itgb5 | 0.565 | NA | TRUE | NA | - | - | NA |
| Col4a5 | Cd47 | 0.564 | NA | TRUE | NA | - | - | NA |
| Pdgfb | Itgav | 0.562 | TRUE | TRUE | yes | - | - | yes |
| Tnc | Sdc1 | 0.562 | TRUE | TRUE | yes | - | - | yes |
| Lrpap1 | Lrp1 | 0.560 | TRUE | TRUE | yes | - | - | yes |
| Wnt3 | Ryk | 0.558 | NA | TRUE | NA | - | - | NA |
| Dlk2 | Notch1 | 0.557 | TRUE | TRUE | yes | - | - | yes |
| Itgb3bp | Itgb5 | 0.554 | TRUE | TRUE | yes | - | - | yes |
| Wnt7b | Fzd1 | 0.553 | NA | TRUE | NA | - | - | NA |
| Rtn4 | Gjb2 | 0.552 | TRUE | TRUE | yes | - | - | yes |
| Lamb3 | Cd151 | 0.552 | TRUE | TRUE | yes | - | - | yes |
| Lamb3 | Itgb4 | 0.552 | TRUE | TRUE | yes | - | - | yes |
| Agrn | Lrp1 | 0.551 | TRUE | TRUE | yes | - | - | yes |
| Vcl | Itgb5 | 0.549 | TRUE | TRUE | yes | - | - | yes |
| Gnas | Adcy1 | 0.549 | TRUE | TRUE | yes | - | - | yes |
| Calm2 | Mylk | 0.546 | TRUE | FALSE | no | - | - | no |
| Calm3 | Egfr | 0.545 | TRUE | TRUE | yes | - | - | yes |
| Dsc1 | Dsg2 | 0.544 | TRUE | TRUE | yes | - | - | yes |
| Bmp7 | Acvr2a | 0.540 | NA | TRUE | NA | - | - | NA |
| App | Slc45a3 | 0.539 | NA | TRUE | NA | - | - | NA |
| Sema6a | Plxna2 | 0.538 | TRUE | TRUE | yes | - | - | yes |
| Efnb1 | Ephb4 | 0.536 | TRUE | TRUE | yes | - | - | yes |
| Ubc | Smad3 | 0.536 | TRUE | TRUE | yes | - | - | yes |
| Gstp1 | Traf2 | 0.536 | TRUE | TRUE | yes | - | - | yes |
| Rtn4 | Rtn4r | 0.535 | TRUE | NA | NA | - | - | NA |
| Areg | Erbb3 | 0.533 | TRUE | TRUE | yes | - | - | yes |
| Inhbb | Acvr2a | 0.532 | NA | TRUE | NA | - | - | NA |
| Calm1 | Hmmr | 0.529 | TRUE | TRUE | yes | - | - | yes |
| Col18a1 | Itgb5 | 0.526 | TRUE | TRUE | yes | - | - | yes |
| Il1a | Il1rap | 0.526 | NA | TRUE | NA | - | - | NA |
| Il18 | Il1rl2 | 0.526 | TRUE | TRUE | yes | - | - | yes |
| Col4a5 | Itgav | 0.526 | NA | TRUE | NA | - | - | NA |
| Egf | Ldlr | 0.522 | NA | TRUE | NA | - | - | NA |
| B2m | Klrd1 | 0.521 | TRUE | TRUE | yes | - | - | yes |
| Wnt7b | Lrp5 | 0.521 | NA | TRUE | NA | - | - | NA |
| Thbs1 | Itga6 | 0.520 | TRUE | TRUE | yes | - | - | yes |
| Liph | Lpar2 | 0.519 | TRUE | TRUE | yes | - | - | yes |
| Lpl | **Cd44** | 0.518 | NA | TRUE | NA | yes | - | NA |
| Sfrp1 | Fzd6 | 0.515 | TRUE | TRUE | yes | - | - | yes |
| Pdgfb | Lrp1 | 0.514 | TRUE | TRUE | yes | - | - | yes |
| Timp2 | Itga3 | 0.514 | TRUE | TRUE | yes | - | - | yes |
| Thbs1 | Cd47 | 0.512 | TRUE | TRUE | yes | - | - | yes |
| Lama5 | Itga6 | 0.509 | TRUE | TRUE | yes | - | - | yes |
| Hspg2 | Lrp1 | 0.509 | TRUE | TRUE | yes | - | - | yes |
| Thbs2 | Itga6 | 0.509 | TRUE | TRUE | yes | - | - | yes |
| Fgf18 | Fgfr1 | 0.508 | FALSE | TRUE | no | - | - | no |
| Pros1 | Axl | 0.508 | TRUE | TRUE | yes | - | - | yes |
| Ltbp1 | Itgb5 | 0.508 | TRUE | TRUE | yes | - | - | yes |
| Nampt | Insr | 0.507 | TRUE | TRUE | yes | - | - | yes |
| Rps27a | Smad3 | 0.507 | TRUE | TRUE | yes | - | - | yes |
| Egf | Erbb2 | 0.505 | NA | TRUE | NA | - | - | NA |
| Efna4 | Epha2 | 0.504 | TRUE | NA | NA | - | - | NA |
| Il1f6 | Il1f5 | 0.502 | NA | NA | NA | - | - | NA |
| Scgb1a1 | Lmbr1l | 0.502 | FALSE | NA | no | - | - | no |
| Thbs2 | Cd47 | 0.501 | TRUE | TRUE | yes | - | - | yes |

**Supplementary Table 2.** scTensor comparison results.

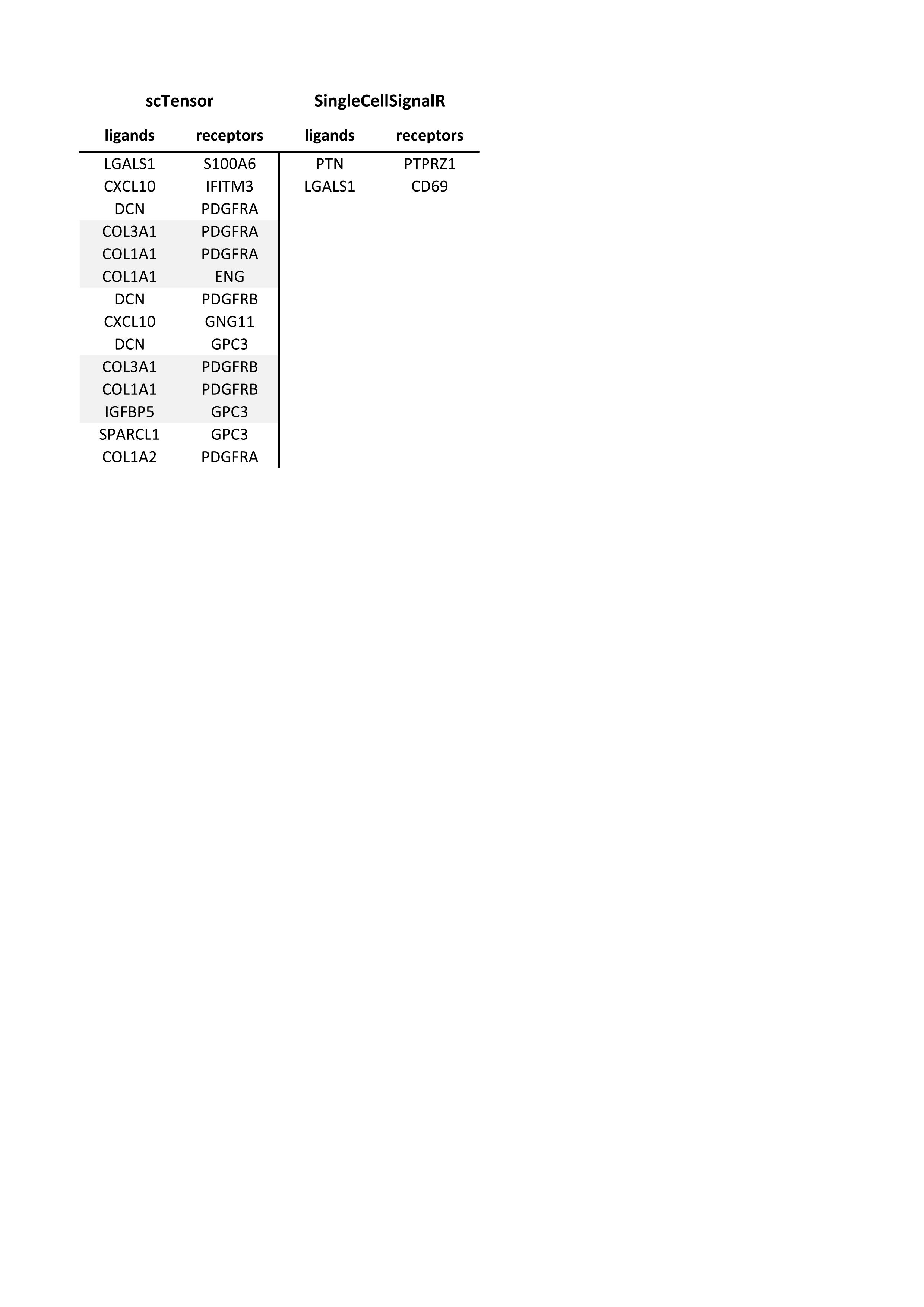

**Supplementary Table 3.** Paracrine interactions, comparison with PyMINEr on 10xPBMC data. SingleCellSignalR LRscore > 0.5.

|  | **Number of inferred LR interactions** | | |
| --- | --- | --- | --- |
| **Cell types** | **SCSignalR** | **PyMINEr** | **Shared** |
| T-cells-B-cells | 10 | 211 | 0 |
| T-cells-Macrophages | 17 | 669 | 0 |
| T-cells-Cytotoxic cells | 15 | 378 | 2 |
| T-cells-Neutrophils | 24 | 269 | 3 |
| B-cells-T-cells | 10 | 211 | 1 |
| B-cells-Macrophages | 28 | 721 | 0 |
| B-cells-Cytotoxic cells | 22 | 294 | 0 |
| B-cells-Neutrophils | 27 | 286 | 0 |
| Macrophages-T-cells | 17 | 669 | 1 |
| Macrophages-B-cells | 16 | 721 | 2 |
| Macrophages-Cytotoxic cells | 33 | 983 | 6 |
| Macrophages-Neutrophils | 9 | 1055 | 1 |
| Cytotoxic cells-T-cells | 19 | 378 | 0 |
| Cytotoxic cells-B-cells | 28 | 294 | 0 |
| Cytotoxic cells-Macrophages | 64 | 983 | 2 |
| Cytotoxic cells-Neutrophils | 50 | 399 | 5 |
| Neutrophils-T-cells | 38 | 269 | 2 |
| Neutrophils-B-cells | 42 | 286 | 4 |
| Neutrophils-Macrophages | 43 | 1055 | 3 |
| Neutrophils-Cytotoxic cells | 56 | 399 | 4 |

**Supplementary Table 4.** Paracrine and autocrine interactions, comparison with PyMINEr on 10xPBMC data. SingleCellSignalR LRscore > 0.5.

|  | **Number of inferred LR interactions** | | |
| --- | --- | --- | --- |
| **Cell types** | **SCSignalR** | **PyMINEr** | **Shared** |
| T-cells-T-cells | 136 | 358 | 6 |
| T-cells-B-cells | 138 | 211 | 5 |
| T-cells-Macrophages | 202 | 669 | 7 |
| T-cells-Cytotoxic cells | 158 | 378 | 13 |
| T-cells-Neutrophils | 166 | 269 | 7 |
| B-cells-T-cells | 128 | 211 | 4 |
| B-cells-B-cells | 127 | 352 | 3 |
| B-cells-Macrophages | 193 | 721 | 5 |
| B-cells-Cytotoxic cells | 149 | 294 | 3 |
| B-cells-Neutrophils | 158 | 286 | 3 |
| Macrophages-T-cells | 179 | 669 | 12 |
| Macrophages-B-cells | 170 | 721 | 7 |
| Macrophages-Macrophages | 245 | 2800 | 41 |
| Macrophages-Cytotoxic cells | 189 | 983 | 17 |
| Macrophages-Neutrophils | 209 | 1055 | 19 |
| Cytotoxic cells-T-cells | 150 | 378 | 7 |
| Cytotoxic cells-B-cells | 148 | 294 | 6 |
| Cytotoxic cells-Macrophages | 216 | 983 | 15 |
| Cytotoxic cells-Cytotoxic cells | 165 | 512 | 17 |
| Cytotoxic cells-Neutrophils | 175 | 399 | 10 |
| Neutrophils-T-cells | 158 | 269 | 5 |
| Neutrophils-B-cells | 143 | 286 | 5 |
| Neutrophils-Macrophages | 228 | 1055 | 21 |
| Neutrophils-Cytotoxic cells | 179 | 399 | 8 |
| Neutrophils-Neutrophils | 195 | 384 | 9 |

**Supplementary Table 5.** Paracrine and autocrine interactions, comparison with CellPhoneDB on the 10xPBMC data. SingleCellSignalR LRscore > 0.5. Since CellPhoneDB returns its predictions without directionality, we put our results in the same format and each combination of cell types appears once only.

|  | **Number of inferred LR interactions** | | |
| --- | --- | --- | --- |
| **Cell types** | **SCSignalR** | **CellPhoneDB** | **Shared** |
| T-cells-T-cells | 110 | 6 | 4 |
| T-cells-B-cells | 158 | 15 | 10 |
| T-cells-Macrophages | 224 | 28 | 15 |
| T-cells-Cytotoxic cells | 156 | 15 | 8 |
| T-cells-Neutrophils | 219 | 26 | 13 |
| T-cells-Treg | 147 | 8 | 6 |
| B-cells-B-cells | 125 | 8 | 3 |
| B-cells-Macrophages | 234 | 32 | 14 |
| B-cells-Cytotoxic cells | 186 | 20 | 8 |
| B-cells-Neutrophils | 229 | 30 | 14 |
| B-cells-Treg | 168 | 17 | 10 |
| Macrophages-Macrophages | 225 | 34 | 14 |
| Macrophages-Cytotoxic cells | 269 | 41 | 20 |
| Macrophages-Neutrophils | 242 | 37 | 17 |
| Macrophages-Treg | 234 | 34 | 16 |
| Cytotoxic cells-Cytotoxic cells | 153 | 15 | 6 |
| Cytotoxic cells-Neutrophils | 252 | 37 | 23 |
| Cytotoxic cells-Treg | 182 | 14 | 9 |
| Neutrophils-Neutrophils | 209 | 28 | 16 |
| Neutrophils-Treg | 227 | 30 | 14 |
| Treg-Treg | 111 | 4 | 3 |

**Supplementary Table 6.** Paracrine and autocrine interactions, comparison with iTALK on 10xPBMC data. 10xPBMC dada taken from iTALK GitHub for this particular case to match iTALK manuscript. SingleCellSignalR LRscore > 0.5.

|  | **Number of inferred LR interactions** | | |
| --- | --- | --- | --- |
| **Cell types** | **SCSignalR** | **iTALK** | **Shared** |
| cd56_nk-cd14_monocytes | 134 | 118 | 97 |
| cd56_nk-b_cells | 100 | 118 | 68 |
| cd56_nk-cytotoxic_t | 140 | 118 | 100 |
| cd56_nk-regulatory_t | 122 | 118 | 88 |
| cd56_nk-memory_t | 122 | 118 | 94 |
| cd56_nk-naive_t | 93 | 118 | 68 |
| cd14_monocytes-cd56_nk | 157 | 118 | 105 |
| cd14_monocytes-b_cells | 107 | 118 | 73 |
| cd14_monocytes-cytotoxic_t | 137 | 118 | 97 |
| cd14_monocytes-regulatory_t | 120 | 118 | 88 |
| cd14_monocytes-memory_t | 128 | 118 | 99 |
| cd14_monocytes-naive_t | 97 | 118 | 71 |
| b_cells-cd56_nk | 141 | 118 | 89 |
| b_cells-cd14_monocytes | 119 | 118 | 83 |
| b_cells-cytotoxic_t | 124 | 118 | 84 |
| b_cells-regulatory_t | 119 | 118 | 86 |
| b_cells-memory_t | 111 | 118 | 85 |
| b_cells-naive_t | 88 | 118 | 66 |
| cytotoxic_t-cd56_nk | 148 | 118 | 98 |
| cytotoxic_t-cd14_monocytes | 114 | 118 | 80 |
| cytotoxic_t-b_cells | 88 | 118 | 56 |
| cytotoxic_t-regulatory_t | 106 | 118 | 76 |
| cytotoxic_t-memory_t | 107 | 118 | 81 |
| cytotoxic_t-naive_t | 81 | 118 | 58 |
| regulatory_t-cd56_nk | 150 | 118 | 99 |
| regulatory_t-cd14_monocytes | 125 | 118 | 93 |
| regulatory_t-b_cells | 99 | 118 | 66 |
| regulatory_t-cytotoxic_t | 136 | 118 | 96 |
| regulatory_t-memory_t | 114 | 118 | 86 |
| regulatory_t-naive_t | 89 | 118 | 65 |
| memory_t-cd56_nk | 160 | 118 | 106 |
| memory_t-cd14_monocytes | 130 | 118 | 95 |
| memory_t-b_cells | 106 | 118 | 70 |
| memory_t-cytotoxic_t | 136 | 118 | 94 |
| memory_t-regulatory_t | 116 | 118 | 81 |
| memory_t-naive_t | 94 | 118 | 68 |
| naive_t-cd56_nk | 144 | 118 | 94 |
| naive_t-cd14_monocytes | 119 | 118 | 86 |
| naive_t-b_cells | 92 | 118 | 61 |
| naive_t-cytotoxic_t | 126 | 118 | 87 |
| naive_t-regulatory_t | 106 | 118 | 74 |
| naive_t-memory_t | 103 | 118 | 79 |

**Supplementary Box 1.** Example of SingleCellSIgnalR usage.

```
library(SingleCellSignalR)

# Define your working directory
setwd("~/example/")

# Define the file of interest you want to work with
file = "example_dataset.txt"

# Prepare the data for the analysis
data = data_prepare(file = file)
genes = rownames(data)

# Proceed to clustering
clust = clustering(data = data,n = 10, method = "simlr")
cluster = clust$cluster
tsne = clust$`t-SNE`

# Cell classification
my.markers = markers(c("immune"))
class = cell_classifier(data=data, genes=genes, markers = my.markers)

# Cluster analysis
clust.ana = cluster_analysis(data = data, genes = genes, cluster = cluster, markers = my.markers)

# Proceed to cell signaling
signal = cell_signaling(data = data, genes = genes, cluster = cluster,species = "homo sapiens")

# Visualization
visualize(inter = signal)
visualize(inter = signal, show.in = c(5))

expression.plot(data = data, name = "CD14", tsne = tsne)
expression.plot.2(data = data, name.1 = "CD40LG", name.2 = "CD40", tsne = tsne)

# Create interface network
inter.net = inter_network(data = data, signal = signal, genes = genes, cluster = cluster)

# Show interactions downstream a specific receptor
intra = intra_network(goi = "S1PR1",data = data,genes = genes,cluster = cluster,coi="cluster 3",signal=signal)
```

**Supplementary Box 2.** Example of integration with an external tool (Seurat).

```
library(SingleCellSignalR)
library(Seurat)

# Define your working directory
setwd("~/example/")

# Pre-processing using Seurat (https://satijalab.org/seurat/)
pbmc.data <- Read10X(data.dir = "./filtered_feature_bc_matrix/")
pbmc <- CreateSeuratObject(counts = pbmc.data, project = "pbmc1k")

# Data filtering and normalization
pbmc[["percent.mt"]] <- PercentageFeatureSet(pbmc, pattern = "^MT-")
pbmc <- subset(pbmc, subset = nFeature_RNA > 40)
pbmc <- NormalizeData(pbmc,scale.factor = 10000)
pbmc <- FindVariableFeatures(pbmc, selection.method = "vst", nfeatures = 2000)

# Data clustering
all.genes <- rownames(pbmc)
pbmc <- ScaleData(pbmc, features = all.genes)

pbmc <- RunPCA(pbmc, features = VariableFeatures(object = pbmc))
pbmc <- FindNeighbors(pbmc, dims = 1:10)
pbmc <- FindClusters(pbmc, resolution = 0.1)

# Retreiving the results of the preprocessing from the Seurat object
cluster = as.numeric(Idents(pbmc))
data = data.frame(pbmc[["RNA"]]@data)

# Ligand/Receptor analysis using SingleCellSignalR
signal = cell_signaling(data=data,genes=all.genes,cluster=cluster)

# Visualization
visualize(signal)
intra = intra_network("S1PR1",data,all.genes,cluster,"cluster 3",signal = signal)
```

**Supplementary Methods**

**Built-in call type calling algorithm**

In case SingleCellSignalR users do not want to use a specialized tool to infer the cell types corresponding to the cell clusters, we implemented the following algorithm.

A limited number of curated gene signatures are integrated with SingleCellSignalR that cover common cell types for our laboratory applications:

- Immune: T cells, B cells, macrophages, cytotoxic cells, dendritic cells, mast cells, neutrophils, natural killer cells, regulatory T cells;
- Tumor microenvironment: endothelial cells, cancer-associated fibroblasts;
- Melanoma: melanoma cancer cells;
- Breast cancer: triple-negative, HER+, and ER+ breast cancer cells.

They are stored in a format identical to PanglaoDB (9) exports such that users can easily add cell types from this rich source or provide their own. A cell type $t$ signature is comprised of genes $g_{t,1},\cdots,g_{t,k_{t}}$and its average expression in each cell $j$ is $a_{t,j}=\frac{1}{k_{t}}\sum_{i=1}^{k_{t}} c_{\text{row}\left( g_{t,i} \right),j}$, with $\text{row}\left( g_{t,i} \right)$ the row in $C$ representing gene $g_{t,i}$. $C$ is the preprocessed gene expression matrix, i.e., the normalized and clustered read counts. A matrix $A=(a_{t,j})$ is obtained with the average expression of all the signatures, which we normalize imposing that columns sum to 1. The normalized matrix is called $\tilde{A}$. We adjust a threshold $a^{*}$ such that the condition $\tilde{a}_{t,j}>a^{*}$ maximizes the number of cells assigned to a single cell type.

We illustrate the application of the algorithm above to 10xPBMC data (1) comprised of 7,857 cells. After data normalization with our default procedure (Materials and Methods), SIMLR applied with default parameters identified 6 cell clusters (**Suppl. Fig. 15A**). Using the predefined “immune” gene signatures revealed the predominant subpopulations (>10 cells). Two SIMLR clusters were homogeneous (B cells in cluster 1 and cytotoxic cells in cluster 6), while the other 4 clusters were mixed (**Suppl. Fig. 15B**). At the individual transcriptome level, cell type calling was unambiguous for 94% of the cells (**Suppl. Fig. 15C**). This classification was projected in the t-SNE coordinates (**Suppl. Fig. 15D**). Less than 2% of cells were assigned to more than one type. They corresponded to lymphoid and myeloid cells in an intermediate state with a dual assignment: T cells and cytotoxic T cells, or macrophages and neutrophils. In the remaining 4%, SingleCellSignalR cell type caller failed to attribute any cell type. Cells assigned to multiple or no cell type are featured as gray dots in **Suppl. Fig. 15D**. Cluster 2 was refined by cell type calling to unravel a gradient from neutrophils to macrophages, which come from the same lineage. Similarly, cluster 4 featured a gradient from cytotoxic cells to T cells, with a small number of regulatory T in the middle. Cluster 5 was made of B cells in a different transcriptomic state compared to cluster 1. The large cluster 4 that we found to cover T cells and cytotoxic cells could be further decomposed in multiple cytotoxic subpopulations. No NK cells were found in clusters 4 and 6.

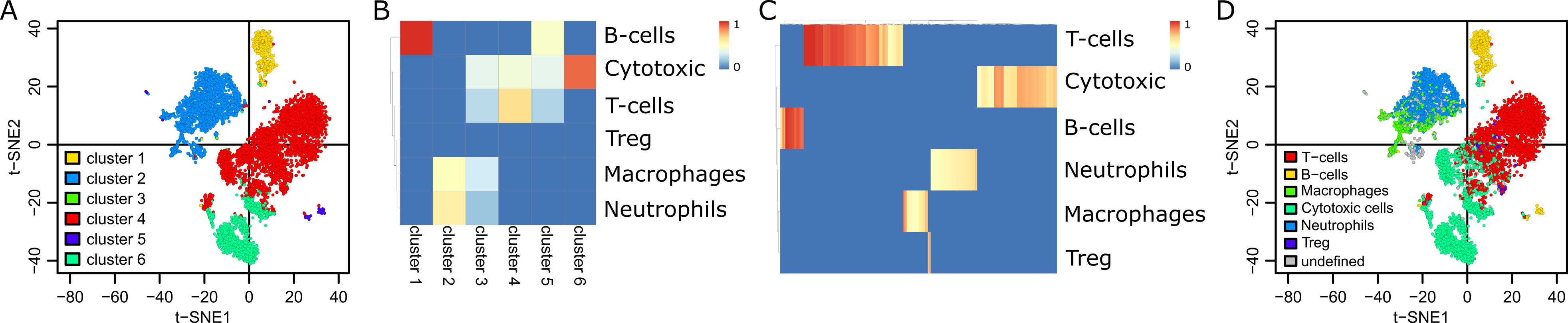

**Supplementary Figure 15.** Illustration of cell type calling algorithm.

A second illustration was obtained re-classifying the cells of the MELANOMA data set (6) and comparing with the authors original calling. We obtained very similar results as shown in **Suppl. Table 7** below.

**Supplementary Table 7.** Comparison of cell type calling between original MELANOMA data set values (patients 79, 80, 88, and 89) and SingleCellSignalR algorithm predictions.

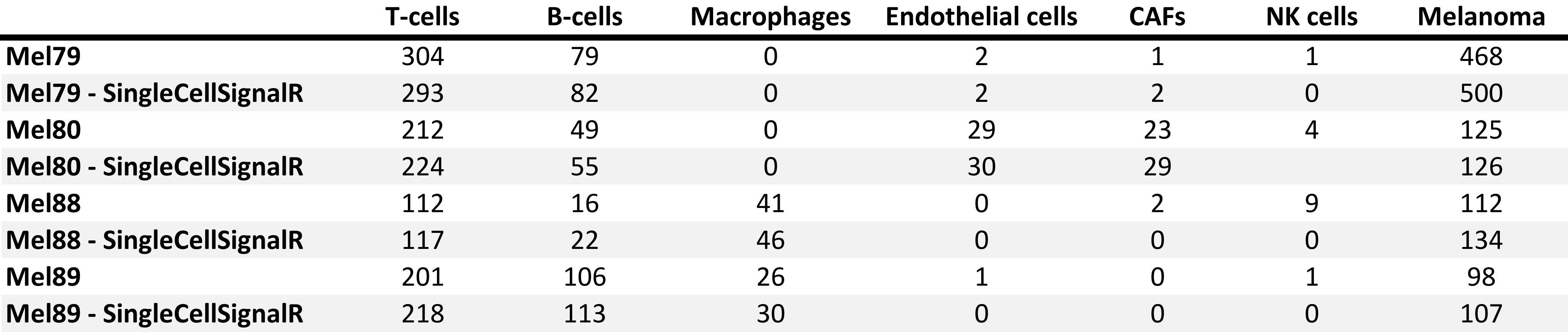

**Autocrine *versus* paracrine interactions**

By default, an interaction between two cell types A and B, with ligand in A and receptor in B, is considered paracrine if the A cells do not express the receptor (and *vice versa*). This cutoff can be raised, e.g., to 2%, to accommodate slight expression of the receptor in A (or the ligand in B) and maintain the paracrine classification. In this work, we used 0% everywhere but for the mouse interfollicular epidermis data (2%). In every case, all the LR pairs considered significant (LR score above threshold) will be classified as paracrine or autocrine.
